# Biological point-light displays scanning by the principal eyes of a jumping spider

**DOI:** 10.1101/2025.06.23.661007

**Authors:** Massimo De Agrò, Alex M. Winsor, Wes Walsh, Paul Shamble, Elizabeth Jakob

**Affiliations:** Center for Mind/Brain Sciences, University of Trento, Italy; Graduate Program in Organismic and Evolutionary Biology, University of Massachusetts Amherst; Kavli Institute for Neuroscience, Department of Neuroscience, Yale University School of Medicine, USA; Department of Biology, University of Massachusetts Amherst

**Keywords:** psychophysics, life detector, shape from motion, invertebrate, vision

## Abstract

The semi-rigid structure of bodies forces mobile animals to move in rhythmic patterns shared by all creatures with skeletons, exoskeletons, or limb connections. This pattern, known as biological motion, is instantly recognizable and conveys “animacy,” even when body shape is removed and only a cloud of moving dots is shown. Indeed, motion alone is so informative that some animals can infer the original shape based on the dots’ concurrent activity. Jumping spiders, highly visual arthropods, divide motion detection and shape recognition between their four pairs of eyes. Previous studies found that they can distinguish biological from non-biological motion using only their motion-detecting anterior lateral eyes yet seemed unable to extract shape from motion. In this study, we examined how the anterior medial eyes of jumping spiders—which are used for shape recognition—respond to dot clouds depicting biological or non-biological motion. Using a custom eye tracker, we monitored retinal movements during presentation of static and moving stimuli. We found spiders change their retinal shifting pattern based on both the target’s motion (biological or not) and implied structure (i.e., whether dots suggest a coherent shape). These results reveal that jumping spiders analyze motion with more complexity than previously thought, suggesting a deeper integration of motion and form processing within their minuscule, modular brains.

## 1 Introduction

Vision is an amazing biological feat—by capturing light from highly organized sensors, animals can meaningfully interpret a scene, segregating objects from the background, and categorizing and recognizing these objects. Yet while most research on vision focuses on static images much can be understood about objects in the environment through motion alone, even in the absence of a recognizable shape. This motion-based vision is particularly useful when detecting other animals in the environment, and distinguishing them from non-living objects. For example, animals are “self-propelled”, meaning that they are the only objects in the environment that can initiate, modulate, and cease locomotion without the intervention of an external agent (Di Giorgio et al., 2016; Rosa-Salva et al., 2016; Scholl and Tremoulet, 2000)—thus, movement can provide important clues about object identity.

However, this is not to say that object (or creature) shape is irrelevant. On the contrary, specifically due to their construction, animal bodies are spatially constrained, and therefore conform to semi-rigid motion patterns—for example, a wrist may never change its distance from an elbow, and can only move so far from a knee. Even in an arm which completely lacks rigid skeletal elements (e.g., in an octopus) these constraints are at work, as sections of the body move coherently with neighboring sections. This characteristic structure, and consequently this semi-rigid, idiosyncratic motion pattern, is shared across most animals and is typically referred to as “biological motion” (Johansson, 1975).

The ability to recognize biological motion in the absence of any structural information was first described in humans by Gunnar Johansson in the 1970s (Johansson, 1973; Johansson, 1975; Johansson, 1976). The author produced stimuli composed of just 11 dots, moving in correspondence with the major joints of a human body during locomotion. These deeply impoverished stimuli, termed point-light displays, were capable of eliciting the impression of a walking human in the observers. The effect is not limited to the recognition of a conspecific: observers can guess from the moving points of light the kind of animals they are looking at, which action is being performed, the direction of motion, or even subtler differences like the individual’s sex (Troje, 2002; Troje, 2008; Troje, 2013).

Curiously, even in point-light displays in which the initial location of each point is “scrambled” but preserve the same local trajectory, observers perceive “something alive”, even if no recognizable identity can be inferred (Bardi et al., 2011; Regolin et al., 2000; Troje, 2013; Troje and Westhoff, 2005). Animals seem capable of inferring whether an object is moving biologically, and use this information as a form of “life detector” (Vallortigara, 2021; Vallortigara et al., 2005). Only then can the motion pattern be used to extract the object shape (i.e., “shape from motion”), by virtually connecting the dots through their synchrony. Finally, subtleties of the dots’ motion and directionality can be used to infer characteristics of the depicted action or agent.

The ability to use biological motion as a life detector has been demonstrated to be innate and widespread across a plethora of vertebrate taxa (Blake, 1993; Nakayasu and Watanabe, 2014; Simion et al., 2008; Vallortigara and Regolin, 2006). However, the reconstruction of shape from motion, meaning this ability to infer the structure subtending the point-light display, has been more sparsely studied (Mascalzoni et al., 2009; Troje, 2013). In invertebrates, there is some indirect evidence regarding the use of general motion cues used to visually detect living organisms (De Agrò et al., 2024b), yet, the ability to perceive biological motion has been described before only in jumping spiders (Family: Salticidae) (De Agrò et al., 2021; De Agrò et al., 2024a).

This family of arachnids has recently become a powerful model for the study of visual perception (Cross and Jackson, 2018; De Agrò, 2020; Dolev and Nelson, 2014; Dolev and Nelson, 2016; Jakob et al., 2009; Menda et al., 2014; Shamble et al., 2017). Their highly developed visual system (Winsor et al., 2023) is modularly organized in 4 separate eye-pairs, each with its own dedicated brain region and each pair specialized for a given visual task. The anterior medial eyes (AMEs), or principal eyes, are the largest pair and are characterized by the highest visual acuity (Harland et al., 2012; Land, 1969b)—surpassing the resolution of any insect, and even exceeding many vertebrates. Their boomerang shaped visual fields are quite small but the retinas are situated at the rear of movable tubes (Land, 1969a; Winsor et al., 2021), allowing for the coverage of around 60 degrees of visual span directly in front of the animal. The other three pairs of eyes, anterior lateral, posterior medial and posterior lateral (ALEs, PMEs, PLEs) are collectively referred to as secondary eyes. These are characterized by their lower resolution compared to the AME and their retinas cannot be moved. However, their combined visual field is quite large, spanning across almost 360 degrees all around the spider (Land, 1985).

Crucially, different visual computations are thought to be segregated across different eye pairs, with the secondary eyes specialized in motion perception, and the principal eyes in figure recognition. This is shown by the spiders’ behavior: when a moving target shifts across the secondary eyes visual field, the animal suddenly pivots its body in order to face the object frontally (Land, 1972; Zurek and Nelson, 2012b; Zurek et al., 2010); thereafter, the AME muscles are engaged and the small visual fields start to move across the contours of the object, seemingly reconstructing its shape, in a process dubbed “scanning” (Jakob et al., 2018; Land, 1969a; Zurek and Nelson, 2012a). The movements of the AME visual fields during scanning are stereotyped and can be divided into two types (even if they can happen simultaneously) (Land, 1969a): (i) torsional, where the retinas are rotated around the center of the two fields; (ii) translational, where the retinas move back-and-forth across the target. Horizontal movement can be semi-independent for the two eyes, with each eye focused on different points in the visual scene during this scanning process.

Secondary eyes do not indiscriminately trigger a body pivot for any moving object. Rather, they are selective, often having to choose between many possible targets, not all worthy of attention (Beydizada et al., 2024; Bruce et al., 2021; Loconsole et al., 2024; Spano et al., 2012). Such selective responding requires the ability to not only detect moving objects, but to also discriminate different types of motion. Indeed, recent work has shown that jumping spiders can discriminate biological from non-biological motion (De Agrò et al., 2021) and do so using only their ALEs (De Agrò et al., 2024a). However, unlike in other systems, the spiders’ discrimination ability seems to work only as a “life detector”, without any figure (i.e., structure from motion) recognition, as they performed full body pivots to point-light displays depicting a walking spider and scrambled stimuli (De Agrò et al., 2021) equally.

As stated above, the figure and motion interpretation are split across separate eyes in these creatures, with ALEs only focused on the latter task. This may explain why the secondary eyes could discriminate biological from non-biological motion while remain unable to recognize to which creature they belonged to. The motion-specific eyes may be unable to extract “shape from motion” from the point-light displays, as the neural substrate required for shape perception is not present in the network of these eyes. However, it has been demonstrated that AMEs’ scanning movements are directed by ALEs (Jakob et al., 2018; Zurek and Nelson, 2012a), most likely using object motion to inform the required shifting of principal eyes. This suggests that ALEs may be capable of extracting shape information from biological motion stimuli, but this difference may not be relevant for orientation behavior though it may be crucial for AMEs’ scanning patterns. Observing the shape from motion ability in jumping spiders—along with the previously demonstrated life detector—would provide an immensely powerful model species for studying the two levels (“life detector” and “shape from motion”) of biological motion perception. Jumping spiders specifically offer the unique possibility of testing each module of the visual system alone by simply covering sets of eyes, and allow for deep inferences on the underlying computation that allows for the decoding of biological motion, both in its separate components and as a whole.

In this paper, we measured the scanning patterns produced by AMEs when presented with point-light displays depicting biological and non-biological motion. We placed *Phidippus audax* spiders into a specialized eye-tracker, capable of recording retinal movements of the animals’ AMEs while they are observing stimuli back projected onto a screen (Winsor et al., 2021). We used the same five stimuli described in (De Agrò et al., 2021): (i) a point-light display depicting a walking spiders; (ii) a scrambled point-light display, still biologically moving but with no recognizable figure; (iii) a random point-light display, with the same amount of dot movement, but with no biological motion; (iv) a spider silhouette, with legs moving as in the walking spider point-light stimulus (intended as a control, and providing a traceable figure contour); (v) a static ellipse with no motion information, but with a traceable contour. Every stimulus was presented alone, appearing and remaining static on screen for 12 seconds, then moving in place for 12 seconds, then static again for 12 seconds and finally disappearing. For all three sections we measured retinal velocity, the distance between the two retina centers, the rate of change of the latter (convergence/divergence of the retinas) and the distance between the retinas and the center of the visual field (Figure 1). We hypothesize that spiders’ retinal movement patterns will be indicative of structural exploration for biological displays compared to scrambled or random displays. This would be different than what has been observed when studying the secondary eyes only (De Agrò et al., 2021; De Agrò et al., 2024a), where the spiders shown an identical reaction to biological and scrambled displays.

**Figure 1.**
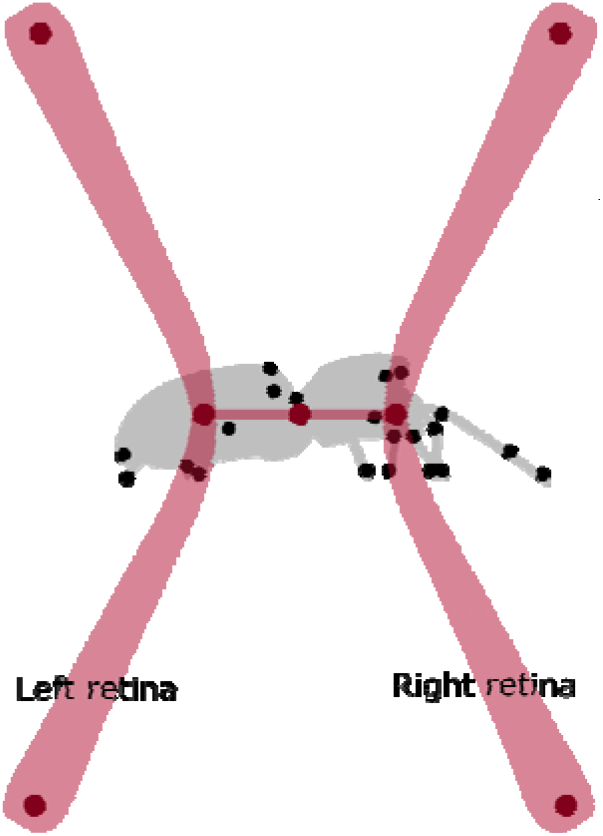
Schematic representation of the AMEs’ retinas. Retinas are depicted in semi-transparent red, overlapped with an example stimulus (the gray spider silhouette is here depicted to show the structure underlying the points, but was not presented together in the experiment). Each retina was tracked in DeepLabCut using three dots, one for each extreme of the “boomerang” shape, and one at the center. During analysis, we drew an imaginary line connecting these two center dots. The center of this line was defined as the center of the full spider visual field, and used to calculate retinal velocity and the distance from the stimulus center. The length of this line was instead defined as the distance between retinas centers, and its change across successive frame was used to determined convergence/divergence speed.

## 2 Materials and methods

### 2.1 Subjects

We collected *Phidippus audax* in the fall of 2022 from buildings, fences, and other artificial structures in Amherst, MA, USA. Spiders were housed individually in plastic boxes (18 × 13 × 10 cm) in a temperature-controlled room set to 25°C with a 16L:8D light cycle. Each enclosure contained a stick, a hollow black tube, and plastic foliage for enrichment. Spiders were fed one *Acheta domesticus* cricket per week and had access to water ad libitum.

For the experiment, we randomly selected 31 individuals from the lab colony using a random number generator. Each spider was isolated and starved for seven days prior to testing. On the day of the experiment, spiders were anesthetized on ice for 10 minutes, secured in Parafilm around a Styrofoam ball with only the anterior eyes exposed, and painted black (Stuart Semple Black Paint) to minimize IR glare during our experiment. The mounted spider was then aligned and calibrated in the eye tracker (see section 4.3).

Each spider completed two trials on the same day. Before each trial, we presented a blank screen for 300 seconds for habituation. Trials consisted of 10 stimulus bouts, each about 35 seconds long and separated by 30-second breaks (see section 4.2). Stimulus identity and order were randomized in Processing (version 3.5.4).

### 2.2 Stimuli

For a complete description of the experimental stimuli used in this experiment, please see (De Agrò et al., 2021). In brief, we employed 5 different stimuli, 3 of which were point light displays, and 2 were complete shapes.

The first stimulus was a “biological motion” point-light display, depicting a spider walking, as viewed directly from the side. To construct it, we collected 72 frames of a 60fps video of a *Salticus scenicus* walking on a twig, where three full steps were visible. Frame by frame, we located the coordinates of the three major joints of the 8 legs, plus the frontal eyes position, the pedicel and the spinneret, totaling 27 points. These coordinates were extracted, rescaled, and shifted such that the points were centered around the pedicel location. This produced a point cloud depicting a spider walking “on the spot”, allowing for seamless video looping.

To produce the second stimulus, the “scrambled” version, every point of the biological motion stimulus was moved to a random position in the very first frame. The new location was selected to be inside the general area occupied by the points in the original stimulus. Thereafter, the points moved with the exact same trajectory that they followed in the original stimulus. We produced 4 different versions of the scrambled stimulus, each with a different randomized position of dots in the first frame. Each subject was randomly assigned to one of the 4 versions.

For the “random” stimulus, we started by randomizing the location of dots in the first frame, as per the scrambled version. Then, for every subsequent frame, we calculated the distance that each point traveled in the original stimulus. We then applied the same displacement distance to the points of this stimulus, but in a random direction. To avoid excessive drift (points moving out of the general shape-area), randomized directions that would have brought the dots too far from the general area occupied by the original stimulus were discarded, and a different randomized direction was assigned. This generated stimuli that had no resemblance to the original biological motion trajectories, but that maintained the same number of dots and the same overall visual displacement. Again, we produced 4 different versions of this stimuli.

The “silhouette” stimulus was constructed starting from the biological motion stimuli. Dots of the same leg were connected by a line, and the shape of the spider was superimposed and aligned to eye-pedicel-spinneret points. This resulted in a silhouette of the spider moving biologically, but with a simplified and recognizable shape.

The “ellipse” stimulus consisted of a static ellipse, with the same number of black pixels as the silhouette stimulus. This was the only stimulus that did not move, thus acting as a baseline for measuring AME eye activity in the presence of a static object.

### 2.3 Eye Tracker

We used a customized salticid-specific eye tracker to track the movement of the principal eye retinas as spiders viewed video stimuli. The system is a modified ophthalmoscope that directs upwelling infrared light (920 nm) through the spider’s carapace via a fiber optic cable (Thorlabs Inc., Newton, NJ, USA) to illuminate the retinas. Spiders were restrained (see section 4.1), mounted on a micromanipulator, and positioned in front of the eye tracker. Alignment of the visual field was achieved using a calibration routine described in detail elsewhere (Jakob et al., 2018). Briefly, we used a moving target along the screen perimeter while checking for saccades and precise retinal tracking. By noting the coordinates of both the retinas and the moving target together, we could virtually superimpose the two coordinate systems.

Video stimuli were back-projected onto a diffusion screen (Roscolux Gel/116) using a Pico projector (AAXA P4-X). Retinal reflections were recorded with an infrared camera through a series of lenses in the eye tracker during stimulus presentation. Stimulus display and retinal position were monitored simultaneously in real time and recorded on separate windows on a large monitor.

### 2.4 Experimental procedure

First, the spider was appropriately positioned in the eye-tracker setup. The projector was left blank, showing a white background for 300 seconds, to allow the animal to habituate and revert back to normal behavior after manipulation. Then, the first stimulus appeared. This was one of the 5 stimuli described above, chosen at random. The image appeared as static, and remained so for a total of 12 seconds. After this time, the stimulus would start moving (all but the ellipse, which instead remained as static) for the following 12 seconds. After this second block elapsed, the stimulus stopped, remaining static for another 12 seconds before disappearing completely. After 30 seconds of blank, the next stimulus (again, randomly assigned) would appear, following the same pattern just described. During the full trial, the spiders were exposed to all 5 types of stimuli, one time oriented right and one time oriented left (left and right being arbitrary, as scrambled and random displays don’t necessarily have a “direction”), for a total of 10 stimuli presentation. The order was randomized, as well as the stimulus version for the scrambled and random point-light displays.

### 2.5 Scoring

Once the videos were collected, we used the software DeepLabCut (Mathis et al., 2018) to automatically track the position of both AMEs retinas in each frame. Using the calibration routine section, we centered and scaled the tracking data onto the stimulus size and position. The center of the stimuli were set as the origin, and the edges of the available visual field were set as ±1 (both in the x and y axes). The stimuli occupied the center section of this visual field, roughly between ±0.45x and ±0.15y. Given the empirical alignment procedure, the level of precision achievable may not be sufficient to exactly know which portion of the stimulus is being attended at any given frame, but can give a general idea on whether the retinas are in the general vicinity of the stimulus or in the periphery of the visual scene. Then, for every frame, we derived the center point between the two retinas, and measured how many pixels it moved with respect to the previous frame. This was set as the eyes shifting speed. We also measured the distance between the two retinas centers in every frame. Then, we measured how much this distance changed between one frame and the next, recorded as the speed of retinal distance change /. Lastly, we measured in each frame the distance of the center point between the two retinas to the stimulus center.

All of the aforementioned measures were recorded temporally matched (with a frame-by-frame precision) with the spiders’ observation: whether there was currently a stimulus projected, whether it was in motion or static, and what stimulus it was. All calculations performed on the coordinates’ data were performed using Python 3.10 (Van Rossum and Drake, 2009), with the packages pandas (Reback et al., 2020), scipy (Virtanen et al., 2020), numpy (Oliphant, 2006; van der Walt et al., 2011), opencv (Bradski, 2000).

### 2.6 Statistical analysis

All analysis were performed in R, with the libraries data.table, glmmTMB, emmeans, car, DHARMa. We also used the library ggplot2 for fast plotting, and the library parallel to accelerate the analysis. We started by focusing on the time between stimulus appearance and motion initiation, sub-setting our full data frame to only these first 12 seconds of every subject and every stimulus presentation. We also selected for every trial 10 more 12 seconds bouts where no stimulus was presented. This acted as a baseline with which compare the eye movement to the various stimuli.

In order to investigate not only the difference across stimuli, but also how responses change over time, we performed a time series, sliding window GLMM analysis. We divided the full 12 seconds into partially overlapping 1-second-long sections, taken every 0.2 seconds. For each, we performed 4 different generalized linear mixed model, one for each dependent variable we selected (eye displacement, inter-eye distance, eye-distance change speed, distance from center).

The stimulus identity (including no stimulus) was used as independent variable as well as the random slope, while individual identity was used as a random intercept. We performed an analysis of deviance for all of the generated models, and recorded the p-values linked with the stimulus identity. We then applied a Bonferroni correction, adjusting for the number of models performed. We then selected only the windows where the p-value was <0.01, considering them sections where a true observable difference between the stimuli was appreciable. As a last preparatory step, we dropped any significant section shorter that 5 consecutive windows, to avoid discussing possible spurious correlations. Only on the selected significant windows, we proceeded with a post-hoc analysis, comparing each stimulus with the no-stimulus (intended to test the difference with rest state) and with the biological motion stimulus (as the focus of the experiment).

We followed the same exact procedure for the next 24 seconds of stimulus presence, from the initiation of motion, to stopping, to then disappearance. This time however, we did not include the no-stimulus data, using instead the ellipse stimulus as a baseline (as it is the only one that does not start moving).

## 3 Results

Only general results are reported here. For the whole result section, we report the maximal p-value for all the time windows that resulted significantly different between stimuli. This means that across the windows considered many could have resulted with a much lower p-value and therefore higher significance. For the full R script, the output of all the models and results, raw data, see (Massimo De Agro’, 2026). For the raw P values for each window, see Supplementary Tables.

### 3.1 Retinal speed

When stimuli first appeared but before they began moving (Figure 2, left panel), we saw significant effects of stimulus type on retinal speed, but only for the initial 3.9s of observation (GLMM analysis of deviance, Bonferroni corrected over the number of windows. *P*<0.004; white sections in graph). The post-hoc revealed that retinas moved faster compared to a no-stimulus control when spiders were viewing either the ellipse (GLMM post-hoc with Bonferroni correction, *P*<0.02) or the spider silhouette (*P*<0.03). The three point-light displays also elicited faster retinal movement compared to the no-stimulus control, but differences faded more quickly, with the biological motion display maintaining a difference until 2s (*P*<0.02), the random display until 1.7s (*P*<0.03) and the scrambled display until 1.4s (*P*<0.03). We found no significant differences between the biological stimulus and the other four stimuli.

**Figure 2.**
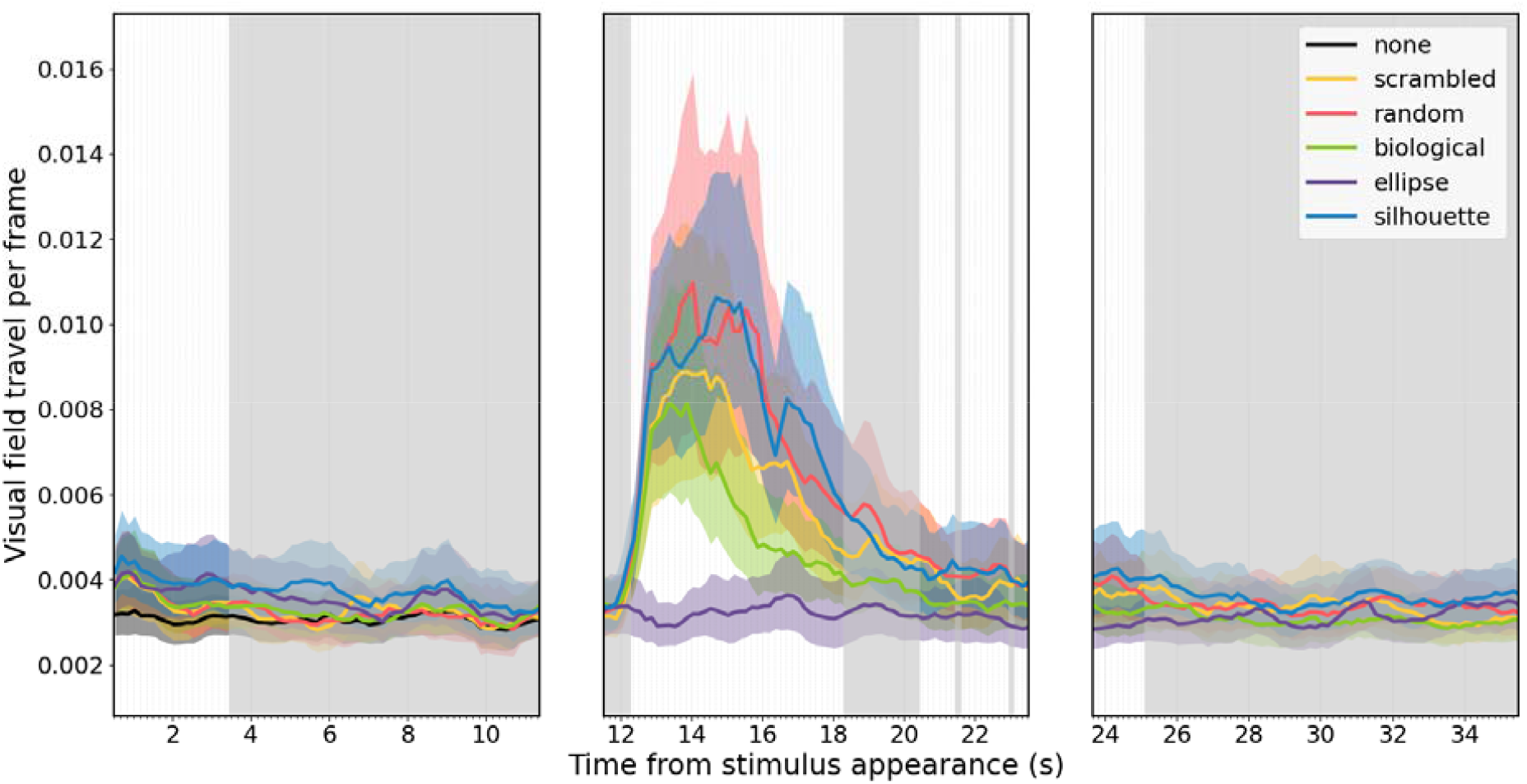
Results describing retinal speed. On the x-axis, the time from initial stimulus appearance in seconds. Every tick of the x-axis represents the center of one window as selected by the sliding window GLMM 1 second wide. This means that the very first window is the one centered on 0.5 seconds (from second 0 to 1). On the y-axis, the distance travelled by the imaginary point centered between the two retinas in each frame expressed as a portion of the visual field size, going from −1 to +1 both in x and y coordinates in the video recording. The plot is divided into three sections: from appearance of the stimulus to motion start, from motion start to motion stop, and from motion stop to disappearance. The 6 lines in the plot represent the 5 different stimuli, plus the no-stimulus control. The latter is shown only for the first section, acting as a comparison for the effect of a static stimulus appearance. Shaded area around the lines represents confidence interval. Across the plot, greyed areas represent windows for which ANOVA did not report any difference between the stimuli.

When stimuli began moving (Figure 2, middle panel), differences in retinal speed became more apparent, with a significant difference across all stimuli from 11.9-18.7s (GLMM analysis of deviance: *P*<0.009). The static ellipse evoked less movement than the silhouette, random or scrambled stimuli (GLMM post-hoc tests with Bonferroni correction; silhouette: *P*<.0001; random: *P*<.0001; scrambled: P<0.015). The biological display consistently evoked faster movement compared to the ellipse only until 16.7s (*P*<0.004), then the difference became intermittent. Across all moving stimuli, retinal speed remained roughly the same until the 13.7-14.7s window. After this, we observed a higher speed for the silhouette over the biological stimulus, lasting until the end of the section (*P*<0.05). A similar effect was observed for the random stimulus, starting at second 13.9 (*P*<0.04). In addition, the scrambled stimulus elicited faster retinal movement in respect to the biological one, but significantly only from 15.4-17.7s (*P*<0.04).

We observed a second section of significant difference between the stimuli, from 20 to 25.9s, intermixed with a few 1-window long breaks (GLMM analysis of deviance: *P*<0.01). In pairwise tests, random and silhouette stimuli resulted significantly different from the ellipse for the whole duration (both P<0.04). The scrambled and biological stimuli also resulted significantly different from the ellipse, but did so intermittently across the section. The difference between the biological stimulus with the random and the silhouette was also intermittent, but appreciable until the end of the section (see supplement for the exact windows). The difference between the biological and the scrambled was instead only observed for the very start of the section, from 20 to 21.4 (P<0.03), and then for a single window towards the end (24.5-25.5s: p=0.046).

After movement stopped (Figure 2, right panel), retinal movement quickly subsided for all stimuli.

### 3.2 Inter-retinal distance

When stimuli first appeared but before they began moving (Figure 3, left panel), inter-retinal distance did not differ across stimuli.

**Figure 3.**
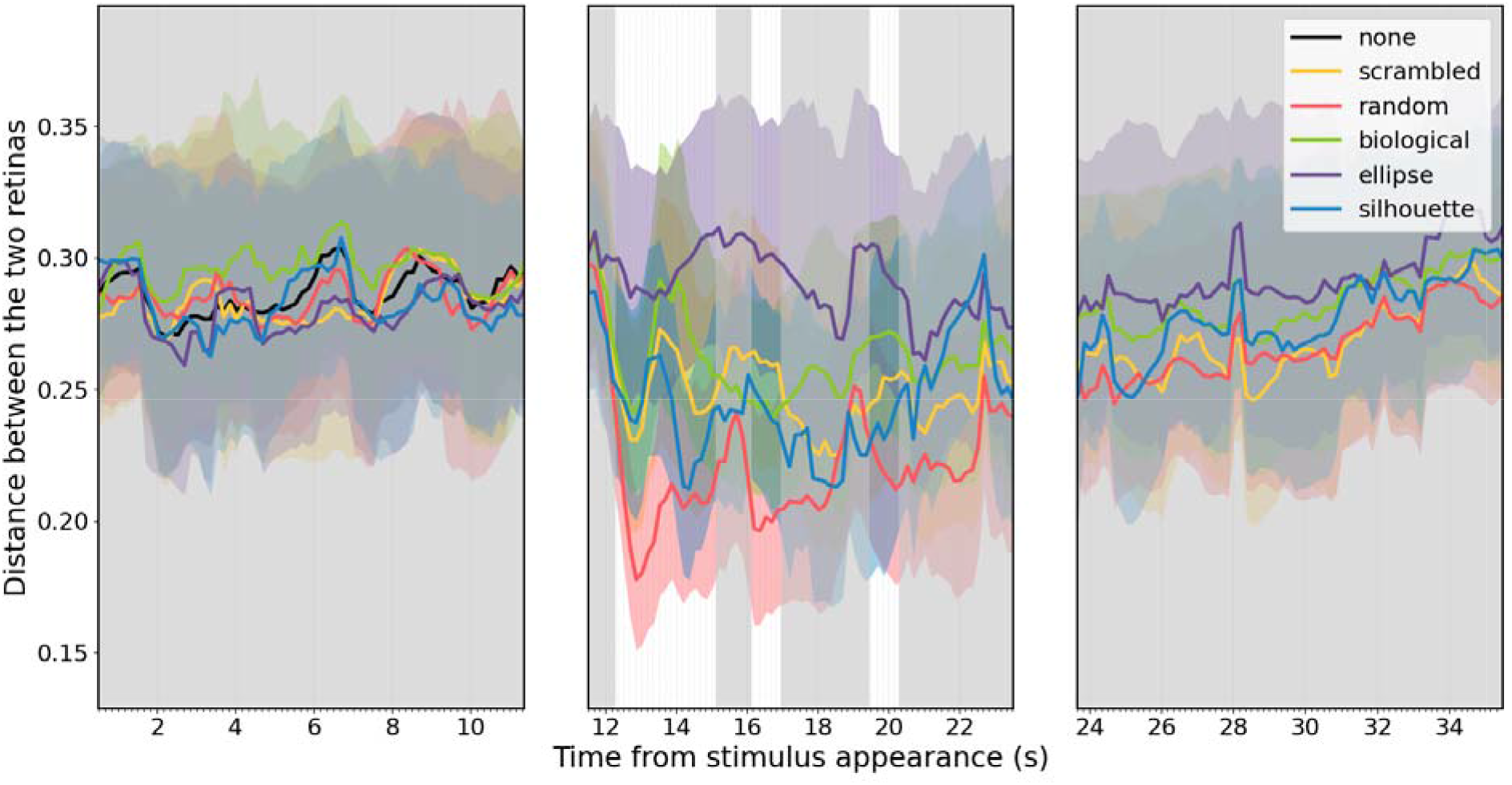
Results describing the distance between two retinas. On the x-axis, the time from initial stimulus appearance in seconds. On the y-axis, the distance between the centers of the two retinas in every frame. The structure of the plot follows the same described in Figure 2.

Upon movement initiation, we observed a significant difference between all stimuli from 11.9 until 15.5s (Figure 3, middle panel; GLMM analysis of deviance: *P*<0.0004). The retinas were significantly closer when spiders viewed the random stimulus compared to the ellipse (*P*<0.0001). The scrambled stimuli and silhouette elicited significantly smaller inter-retinal differences from the ellipse only at the start and end of the section (*P*<0.03 for scrambled and *P*<0.04 for the silhouette). Finally, the biological stimulus was significantly different from the ellipse only at the start of the section (*P*<0.04). We also observed that the distance between the retinas was significantly smaller for random over biological for the whole duration of the section (*P*<=0.04).

Two additional brief periods showed similar patterns of significance (GLMM analysis of deviance: 15.7-17.4s, *P*<0.002; 19-20.7s: *P*<0.008). Across both periods, the random stimulus elicited smaller inter-retinal differences compared to the ellipse (*P*<0.0001). We found significant differences between the ellipse and the silhouette (*P*<0.05), the biological stimulus (*P*<0.004), and the scrambled stimulus (*P*<0.007), but only for brief periods. We again found a difference between the random and the biological stimuli, with the former having a lower inter-retinal distance (*P*<0.04).

Shortly after stimulus movement stopped (Figure 3, right panel), there were no significant differences across stimuli.

### 3.3 Speed of change in inter-retinal distance

When stimuli first appeared but before they began moving (Figure 4, left panel), we saw significant effects of stimulus type on the speed of change of inter-retinal distance, but only for the initial 2.7s of observation (GLMM analysis of deviance P<0.0001). All stimuli differed significantly from the non-stimulus control (silhouette: *P*<0.04; biological motion: *P*<0.02 except for the last window in this section; random: *P*<0.035 except for first and last window; scrambled: *P*<0.055 until 2.2s; ellipse: *P*<0.012 from 0.4-2.7s).

**Figure 4.**
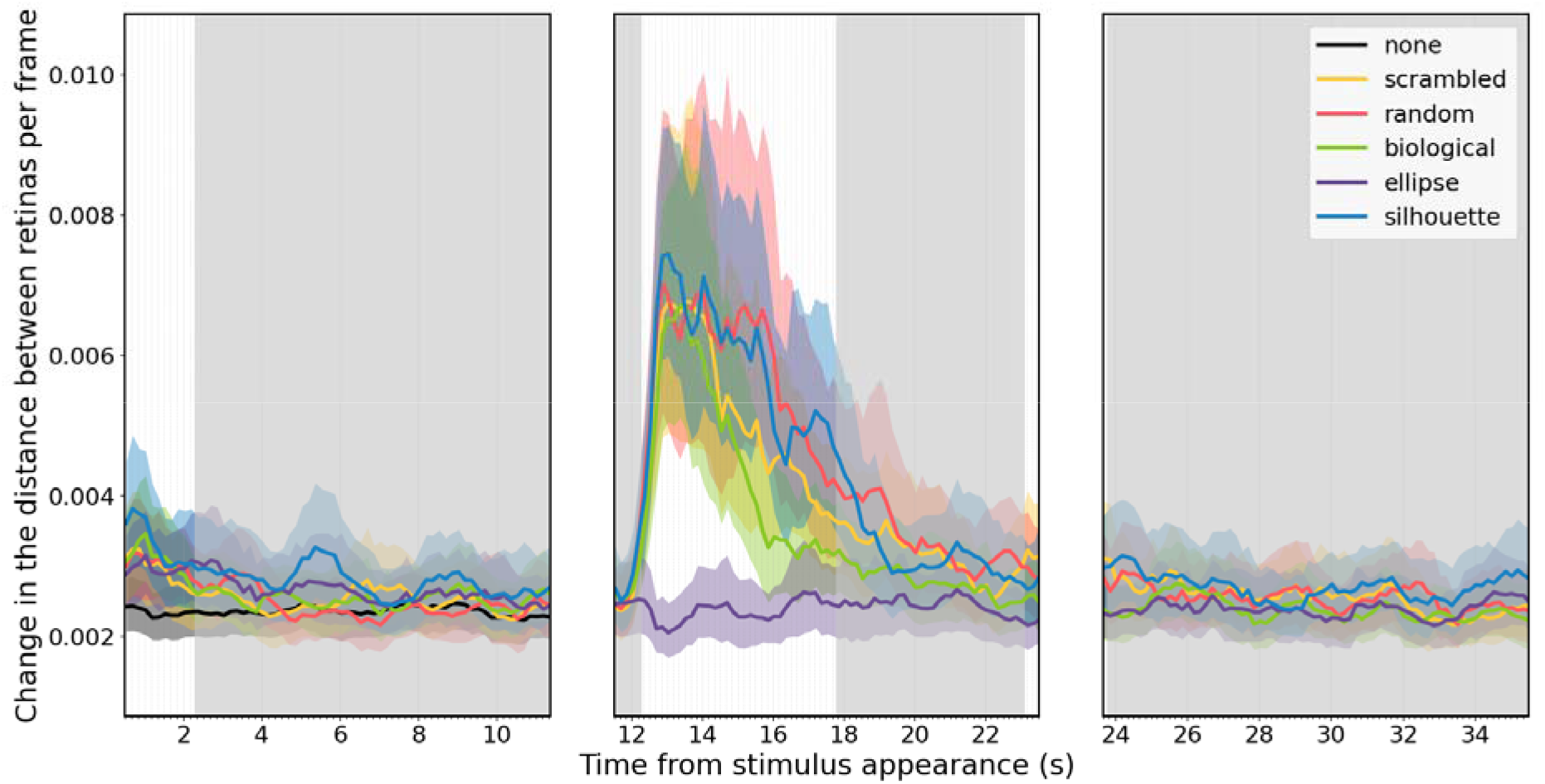
Results describing the distance between two retinas. On the x-axis, the time from initial stimulus appearance in seconds. On the y-axis, the change in the distance between the centers of the two retinas across every frame. The structure of the plot follows the same described in Figure 2.

Upon movement initiation, we observed a significant difference between all stimuli from 11.9 until 18.2s (Figure 4, middle panel; GLMM analysis of: *P*<0.0004). Across this time period, inter-retinal distance changed more rapidly, compared to the ellipse, for the random stimulus (*P*<0.0001), scrambled stimulus (*P*<0.001), spider silhouette (*P*<0.0001), and biological motion stimulus (*P*<0.004). Compared to the other stimuli, responses to the biological motion stimulus more rapidly returned to the same level as the ellipse, at 16.9s. We also observed towards the end of this time period that the speed of change in the distance between retinas was lower for biological motion with respect to random (*P*<0.02), scrambled (*P*<0.02), and silhouette (*P*<0.03).

We then observed a second short section of significant difference among stimuli, from 22.7 to 24.2s at the end of the movement period (GLMM analysis of deviance: *P*<0.005). The change in the distance between retinas was significantly higher for all moving stimuli compared to the ellipse, but only briefly (see Supplement for details).

When the movement stimulus stopped (Figure 4, right panel), this measure returned to baseline.

### 3.4 Distance of retinal midpoint from stimulus center

When stimuli first appeared but before they began moving (Figure 5, left panel), we saw no difference among stimuli for distance between the retinal midpoint to the stimulus center.

**Figure 5.**
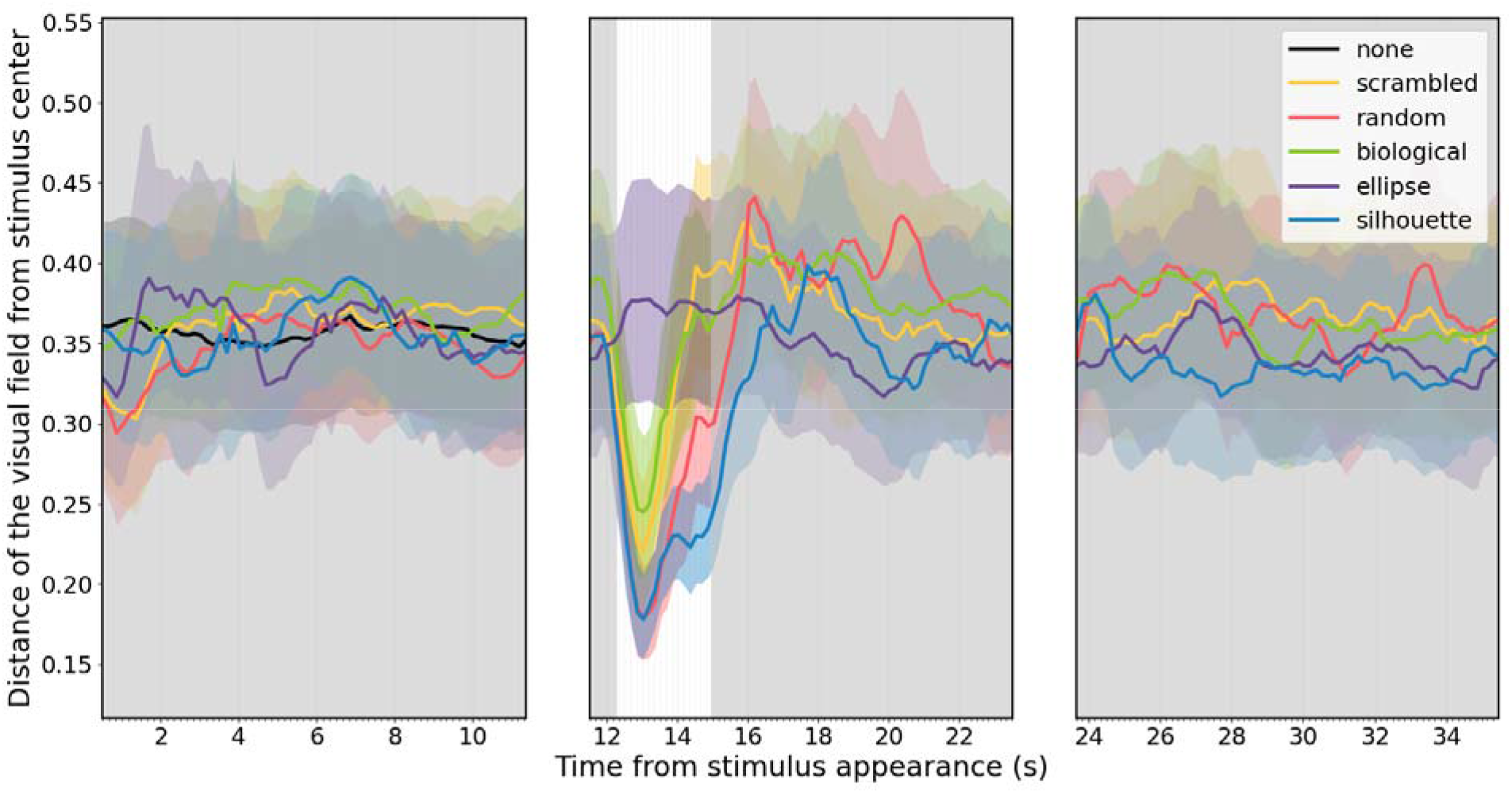
Results describing the distance that the imaginary midpoint between the two retinas has from the center of the stimulus position. On the x-axis, the time from initial stimulus appearance in seconds. On the y-axis, the distance from the stimulus center for each frame. Note that the stimulus has a width of 0.95 and a height of 0.3, expressed as portion of the presentation area going from ±1 both in x and y direction. The structure of the plot follows the same described in Figure 2.

Upon movement initiation, we observed a significant difference between all stimuli from 11.9 until 18.2s (Figure 5, middle panel; GLMM analysis of deviance: *P*<0.002). For each moving stimulus, the retinal midpoint was closer to the stimulus center than we observed for the ellipse (*P*<0.0001).

After the retinas returned to baseline position, they stayed there for the remainder of the trial.

## 4 Discussion

In this paper, we described the AMEs’ movement pattern when presented with biological and non-biological point light displays along with a silhouette and a shape control. We observed many significant differences in these eyes’ activity, as driven by stimulus type. While the data we collected goes much in depth into the activity over the full presentation time, to avoid overextending interpretations we will focus only on the main and evident results.

### 4.1 AMEs’ activity when attending to static stimuli may depend on their structure

When considering the overall gaze displacement, spiders show a significant activity with respect to the no-stimulus control. It could be imagined that this change in speed may reflect a sudden gaze shift from an initial state of quiescence, with the animals are shifting their gazes to fixate on the newly appeared stimulus. Such quick movements towards a target are already described in the literature and are termed “saccades”, since long been described as the typical response of the AMEs upon detection of an appearing target by the ALEs (Winsor et al., 2021; Zurek and Nelson, 2012a; Zurek et al., 2010). The data collected here however makes this interpretation uncertain. In fact, we also observed that the distance from the stimulus center does not change in respect to the no stimulus control for any of the tested targets. Without this evidence, we cannot conclude that the spiders are shifting their gaze toward the target, but only that they increase the eye movement speed. It is possible that the appearance of our targets did not trigger a saccade per se, but only increased the level of generalized activity in resting behavior, possibly as an attentional increase. We know from previous literature that appearing but not translating stimuli in the general visual field of the ALEs can trigger a shift in spatial attention in jumping spiders (Loconsole et al., 2024). It is also possible that our procedure lacked the spatial precision to detect saccades towards the stimuli position. In our setup, the stimuli occupied a wide portion of the available visual scene. Upon appearance, the spiders may have performed some saccades, but each in initially different location of the presented stimuli, increasing the chance of short movements lost to noise.

Regardless of the underlying behavior reflected in the change in retinal speed, we observed differences across stimuli. This initial AME speed change is identical for all 5 stimuli, suggesting that the nature of the target in our paradigm is irrelevant for triggering this behavior. Thereafter, the spiders maintain an activity significantly higher than the no-stimulus control only for the silhouette and ellipse, while the three-point light displays drop quickly down to baseline. A very similar pattern can be detected in the change of distance between the two retinas, with the difference to the no stimulus control maintained for longer for the ellipse and silhouette over the point-light displays. Fittingly, the first two are the only stimuli with a discernible shape, and as such they are the only ones for which the spiders may engage in scanning behavior (Klin and Jones, 2008; Land, 1969a). Indeed, for all three measures we collected of AME movement, we observed no differences between the three point-light displays between the appearance and the initiation of motion. This is in agreement with the literature, as the points cannot be connected into their discernible structure unless already in motion (Troje, 2013). Without motion, the three stimuli are identical sets of scattered dots. It is also worth noting however that the silhouette and ellipse stimuli both present a higher surface area over the point light displays, and therefore more strongly change the overall luminance of the visual scene. It is possible that the change of inter retinal distance and travel speed are a function of overall luminosity of the environment, and not linked to the stimuli shape at all. In general, without finer resolution of retina position, the results on the static images remain uncertain.

### 4.2 Difference between biological and random motion as evidence for the life detector

Upon the initiation of motion, we observed a rapid decrease in the distance from the stimulus center over the static ellipse for all moving stimuli. This confirms that the motion initiation causes a shift of attention and gaze over the stimulus location, and specifically towards its center ±0.15. The shift also allows us to interpret the other measured changes in retinal movement as performed after a saccade, and as such as part of the scanning behavior. For our other three measures the biological display appears to be different from the static ellipse control, as well as the random one. When considering the gaze movement speed, the spiders show a significantly slower velocity for the biological display when compared to all the other stimuli inter retinal distance decreases in respect to the ellipse for the biological, the scrambled and the silhouette, all three to the same level. The random stimulus instead was characterized by an even significantly lower inter-retinal distance. For the speed of change in retinal distance, the biological motion display sits again at the middle, between the slow speed observed for the static ellipse and the faster speed of the other stimuli. Overall, the behaviors produced when the spider is presented with the moving biological display is always different to the responses produced when it presented with the moving random stimulus.

This confirms the ability of jumping spiders to distinguish biological from non-biological types of motion, and that such distinction is influential to their subsequent scanning pattern. This represents the first level of biological motion perception, described in the introduction as the “life detector”. It is likely that this distinction is operated by ALEs alone, as previously demonstrated (De Agrò et al., 2021; De Agrò et al., 2024a). As we know from previous literature, the AMEs movement pattern is driven by the ALEs (Zurek and Nelson, 2012a), which may operate the biological vs non-biological distinction before the first AMEs fixation.

### 4.3 Responses to the scrambled display as evidence for shape-from-motion extraction

Scrambled point light displays fully maintain the biological motion attribute of their original counterpart, but do not depict any discernible shape. As such, they are treated the same by animals for whom discrimination is based on biological motion fidelity (i.e., the life detector) (Vallortigara et al., 2005). However, they appear different if the animal attempts to “connect the dots” as based on their motion pattern, in order to extract a discernible shape (i.e., the figure-from-motion). When previously tested, jumping spiders responded equally to the biological and the scrambled displays, suggesting that at least their secondary eyes are only acting as life detectors (De Agrò et al., 2021)– an expected result, as figure discrimination is a task of the AMEs in their modular visual system.

Here, we observed that both in terms of the visual field shifting speed and the distance change between the eyes, the scrambled display present values significantly higher than the biological and significantly lower than the random. This difference arose only after around 3 seconds of stimulus motion, suggesting that the difference between the three stimuli becomes evident after a minimal period of scanning of the stimulus. It is noteworthy that a similar time elapses before the centering of the gaze starts going back to baseline. We argue that the different retinal speeds observed for the biological, the scrambled, and the random point light displays reflect the differential availability of structural information in the stimulus, that is extracted (or attempted to be) over time through the AMEs’ scanning behavior. This would suggest that jumping spiders also possess the second level of biological motion perception: with ALEs acting as life detector and AMEs extracting the figure from motion. Crucially, the task of “connecting the dots” does not need to happens across gaze shifts. That is, we would not necessarily expect the spiders to move their gaze from one dot to the other, connecting the dots that move together. Indeed, this is improbable especially in our setup, where the relative size of the dots to the retinas field of view is such that more than one dot can be attended on at the same time. The reconstruction can be done directly in the brain, by virtually connecting dots that belong together into patterns of activation. The three point-light-displays presented in this experiment have been constructed in order to have identical low-level characteristics, and only differ in their biological motion component (random vs biological and scrambled) and implied structure (biological vs scrambled).

### 4.1 AMEs’ retinas convergence in response to dynamic stimuli

In this experiment we observed that the average distance between the two centers of the AMEs’ retinas decreases when spiders are presented with moving stimuli.

Indeed, the convergence of the AME in response to moving stimuli is known from earlier studies (Jakob et al., 2018; Land, 1969a) and is driven by ALEs (Jakob et al., 2018). That said, it remains unclear what function this behavior fulfills in the spider’s perception specifically for this class of stimuli, and what drives the difference in convergence across them. We may speculate that maintaining an average narrower AMEs’ fields span may help in following stimuli that are easy to lose sight of. However, unlike in previous work, the stimuli here presented are not translating in space, but instead its only their components, or “limbs” that shift on the spot. Thus, in this specific case the AMEs do not need to track the object, to maintain it in their field of view. However, a more erratically moving stimulus (i.e., the random one in respect to the biological one in this experiment) may have higher perceived probability of quickly shifting away. In other words, AMEs retinal convergence may be linked to the level of predictability of the observed stimulus. This may also explain why the biological and scrambled stimuli induce an equal level of retinal convergence: while they differ in their implied structure, their motion component is identical, which would make them equally “erratic”. In general, average retinal distance seems to be connected only to the motion type observed, thus the scrambled and biological display do not appear to elicit a different response. Regardless, understanding the characteristics of this behavior will require dedicated experiments, testing which stimuli can trigger it, and what its prevention causes in the spiders’ perception. Previous literature had already described a preference for objects with moving limbs (i.e., local motion) over ones simply translating (i.e., global motion), but in that case no eye-tracker was used, and as such no information about retinal convergence is available (Bednarski et al., 2012).

### 4.2 Responses to shaped stimuli

The observed retinal speed, convergence and positioning of the spiders when faced with the ellipse and silhouette stimuli are difficult to interpret. The latter especially appears for some of the measures to be equal to the random point light display, while for others it settles at the same level of the biological. Indeed, the silhouette stimulus moves biologically, following the same locomotion patterns that the point-light display presents. However, the presence of a trackable contour of the picture constitutes a confounding factor. We know from previous literature that during scanning the AMEs’ retinas follow the contour of the target object (Land, 1969a), which will definitely affect the translation speed as calculated in our first model. Here, we observed that upon movement initiation the presentation of the silhouette induces the highest retinal speed level, like owing to both its shape and its biological motion. A very similar pattern is observed for the speed in retinal distance change.

The effect we observed for retinal convergence further supports this hypothesis. As stated above, we hypothesize that this type of behavior is linked to the movement of the target being observed, independently from its shape. Here, the continuously static ellipse remains at baseline level, while the silhouette drops to the biological motion level–thus the shape is irrelevant and the behavior only depends on the motion characteristics.

### 4.3 Conclusions

Here, we tested jumping spiders’ AMEs’ retinal movement upon presentation of biologically and non-biologically moving stimuli. We identified two main behavioral categories: retina movement speed and inter-retinal distance. We hypothesized that the former is more linked to the scanning process, and as such to the object shape, while the latter dynamically changes depending on the target motion. The biological and random point-light displays elicit different responses, confirming the spiders can distinguish the two types of movements using a “life detector” system. On the other hand, biological and scrambled motion appear different for retinal speed (i.e., may be interpreted to be of different “shape”), but not for retinal convergence (i.e., may be interpreted to be moving the same way), suggesting that jumping spiders reach the second level of biological motion perception: i.e., “shape from motion”.

The unprecedented level of detail in the spiders’ discrimination abilities observed here opens up many new possibilities for vision research. Firstly, it brings the AMEs back into the discussion of how these animals accomplish the task of motion perception, while it has long been generally thought that these eyes were used solely for figure discrimination. Indeed, the change in scanning pattern suggests a function that needs to be experimentally explored. Perhaps more importantly, these results suggest a fineness in biological motion discrimination generally observed in “higher” animals (Troje, 2013), here performed by a modular visual system connected to a brain the size of a pinhead.

## 5 Data Availability

Raw data is available at (Massimo De Agro’, 2026)

## References

Bardi, L., Regolin, L. and Simion, F. (2011). Biological motion preference in humans at birth: role of dynamic and configural properties. Developmental Science 14, 353–359.

Beydizada, N. I., Cannone, F., Pekár, S., Baracchi, D. and De Agrò, M. (2024). Habituation to visual stimuli is independent of boldness in a jumping spider. Animal Behaviour 213, 61–70.

Blake, R. (1993). Cats Perceive Biological Motion. Psychol Sci 4, 54–57.

Bruce, M., Daye, D., Long, S. M., Winsor, A. M., Menda, G., Hoy, R. R. and Jakob, E. M. (2021). Attention and distraction in the modular visual system of a jumping spider. Journal of Experimental Biology 224,.

Cross, F. R. and Jackson, R. R. (2018). When it looks and walks like an ant. Learn Behav 46, 103–104.

De Agrò, M. (2020). SPiDbox: design and validation of an open-source “Skinner-box” system for the study of jumping spiders. Journal of Neuroscience Methods 346,.

De Agrò, M., Rößler, D. C., Kim, K. and Shamble, P. S. (2021). Perception of biological motion by jumping spiders. PLoS biology 19, e3001172.

De Agrò, M., Rößler, D. C. and Shamble, P. S. (2024a). Eye-specific detection and a multi-eye integration model of biological motion perception. Journal of Experimental Biology jeb.247061.

De Agrò, M., Galpayage Dona, H. S. and Vallortigara, G. (2024b). Seeing life in the teeming world: animacy perception in arthropods. Front. Psychol. 15,.

Di Giorgio, E., Lunghi, M., Simion, F. and Vallortigara, G. (2016). Visual cues of motion that trigger animacy perception at birth: The case of self-propulsion. Developmental Science 20, n/a-n/a.

Dolev, Y. and Nelson, X. J. (2014). Innate pattern recognition and categorization in a jumping spider. PLoS ONE 9, e97819.

Dolev, Y. and Nelson, X. J. (2016). Biological relevance affects object recognition in jumping spiders. New Zealand Journal of Zoology 43, 42–53.

Harland, D. P., Li, D. and Jackson, R. R. (2012). How Jumping Spiders See the World. In How Animals See the World: Comparative Behavior, Biology, and Evolution of Vision (ed. Lazareva, O.F.), Shimizu, T.), and Wasserman, E. A.), pp. 133–164. New York: Oxford University Press.

Jakob, E., D. Skow, C., Popson Haberman, M. and Plourde, A. (2009). Jumping Spiders Associate Food With Color Cues In A T-Maze. Journal of Arachnology 35, 487–492.

Jakob, E. M., Long, S. M., Harland, D. P., Jackson, R. R., Carey, A., Searles, M. E., Porter, A. H., Canavesi, C. and Rolland, J. P. (2018). Lateral eyes direct principal eyes as jumping spiders track objects. Current Biology 28, R1092–R1093.

Johansson, G. (1973). Visual perception of biological motion and a model for its analysis. Perception & Psychophysics 14, 201–211.

Johansson, G. (1975). Visual motion perception. Scientific American 232, 76–89.

Johansson, G. (1976). Spatio-temporal differentiation and integration in visual motion perception. Psychol. Res 38, 379–393.

Klin, A. and Jones, W. (2008). Altered face scanning and impaired recognition of biological motion in a 15-month-old infant with autism. Developmental Science 11, 40–46.

Land, M. F. (1969a). Movements of the retinae of jumping spiders (Salticidae: dendryphantinae) in response to visual stimuli. J. Exp. Biol. 51, 471–493.

Land, M. F. (1969b). Structure of the Retinae of the Principal Eyes of Jumping Spiders (Salticidae: Dendryphantinae) in Relation to Visual Optics. Journal of Experimental Biology 51, 443– 470.

Land, M. F. (1972). Stepping Movements Made by Jumping Spiders During Turns Mediated by the Lateral Eyes. Journal of Experimental Biology 57, 15–40.

Land, M. F. (1985). Short Communication: Fields of View of the Eyes of Primitive Jumping Spiders. Journal of Experimental Biology 119, 381–384.

Loconsole, M., Ferrante, F., Giacomazzi, D. and De Agrò, M. (2024). Independence and synergy of spatial attention in the two visual systems of jumping spiders. Journal of Experimental Biology 227, jeb246199.

Mascalzoni, E., Regolin, L. and Vallortigara, G. (2009). Mom’s shadow: structure-from-motion in newly hatched chicks as revealed by an imprinting procedure. Anim Cogn 12, 389–400.

Massimo De Agro’ (2026). massimodeagro/SI-Biological-point-light-displays-scanning-by-the-principal-eyes-of-a-jumping-spider: v0.9.

Menda, G., Shamble, P. S., Nitzany, E. I., Golden, J. R. and Hoy, R. R. (2014). Visual Perception in the Brain of a Jumping Spider. Current Biology 24, 2580–2585.

Nakayasu, T. and Watanabe, E. (2014). Biological motion stimuli are attractive to medaka fish. Anim Cogn 17, 559–575.

Regolin, L., Tommasi, L. and Vallortigara, G. (2000). Visual perception of biological motion in newly hatched chicks as revealed by an imprinting procedure. Anim Cogn 3, 53–60.

Rosa-Salva, O., Grassi, M., Lorenzi, E., Regolin, L. and Vallortigara, G. (2016). Spontaneous preference for visual cues of animacy in naïve domestic chicks: The case of speed changes. Cognition 157, 49–60.

Scholl, B. J. and Tremoulet, P. D. (2000). Perceptual causality and animacy. Trends in Cognitive Sciences 4, 299–309.

Shamble, P. S., Hoy, R. R., Cohen, I. and Beatus, T. (2017). Walking like an ant: a quantitative and experimental approach to understanding locomotor mimicry in the jumping spider Myrmarachne formicaria. Proceedings of the Royal Society B: Biological Sciences 284, 20170308.

Simion, F., Regolin, L. and Bulf, H. (2008). A predisposition for biological motion in the newborn baby. PNAS 105, 809–813.

Spano, L., Long, S. M. and Jakob, E. M. (2012). Secondary eyes mediate the response to looming objects in jumping spiders (Phidippus audax, Salticidae). Biol. Lett. 8, 949–951.

Troje, N. F. (2002). Decomposing biological motion: A framework for analysis and synthesis of human gait patterns. Journal of Vision 2, 2–2.

Troje, N. F. (2008). Biological motion perception. he senses: A comprehensive reference 2 18.

Troje, N. F. (2013). What Is Biological Motion? Definition, Stimuli, and Paradigms. In Social Perception (ed. Rutherford, M.D.) and Kuhlmeier, V. A.), pp. 13–36. The MIT Press.

Troje, N. F. and Westhoff, C. (2005). Detection of direction in scrambled motion: a simple “life detector”? Journal of Vision 5, 1058–1058.

Vallortigara, G. (2021). Born Knowing: Imprinting and the Origins of Knowledge. The MIT Press.

Vallortigara, G. and Regolin, L. (2006). Gravity bias in the interpretation of biological motion by inexperienced chicks. Curr Biol 16, R279–280.

Vallortigara, G., Regolin, L. and Marconato, F. (2005). Visually Inexperienced Chicks Exhibit Spontaneous Preference for Biological Motion Patterns. PLOS Biology 3, e208.

Winsor, A. M., Pagoti, G. F., Daye, D. J., Cheries, E. W., Cave, K. R. and Jakob, E. M. (2021). What gaze direction can tell us about cognitive processes in invertebrates. Biochemical and Biophysical Research Communications 564, 43–54.

Winsor, A. M., Morehouse, N. I. and Jakob, E. M. (2023). Distributed Vision in Spiders. In Distributed Vision: From Simple Sensors to Sophisticated Combination Eyes (ed. Buschbeck, E.) and Bok, M.), pp. 267–318. Cham: Springer International Publishing.

Zurek, D. B. and Nelson, X. J. (2012a). Saccadic tracking of targets mediated by the anterior-lateral eyes of jumping spiders. J Comp Physiol A 198, 411–417.

Zurek, D. B. and Nelson, X. J. (2012b). Hyperacute motion detection by the lateral eyes of jumping spiders. Vision Research 66, 26–30.

Zurek, D. B., Taylor, A. J., Evans, C. S. and Nelson, X. J. (2010). The role of the anterior lateral eyes in the vision-based behaviour of jumping spiders. Journal of Experimental Biology 213, 2372–2378.

