## Supplement for "Biological point-light displays scanning by the principal eyes of a jumping spider": analysis_windows.html


Code 

- Show All Code
- Hide All Code

### Biological point-light displays scanning by the principal eyes of a jumping spider - Analysis

#### Biological point-light displays scanning by the principal eyes of a jumping spider - Analysis

- 1 Prepare environment
  - 1.1 loading packages
  - 1.2 loading files
  - 1.3 Definitions
- 2 Analysis
  - 2.1 Whole time-series pass
    - 2.1.1 Eye displacement
    - 2.1.2 Inter-eye
      distance
    - 2.1.3 Inter-eye distance
      change
    - 2.1.4 Dist\_from\_center
  - 2.2 In depth analysis
    - 2.2.1 Eye displacement
    - 2.2.2 Inter-eye
      distance
    - 2.2.3 Inter-eye distance
      change
    - 2.2.4 Dist from Center
  - 2.3 Plots
    - 2.3.1 Eye displacement
    - 2.3.2 Inter-eye
      distance
    - 2.3.3 Inter-eye distance
      change
    - 2.3.4 Dist from Center

2026-02-19

#### Massimo De Agrò1, Alex M. Winsor2, Wes Walsh2, Paul Shamble3, Elizabeth Jakob4

###### 1 – Center for Mind/Brain Sciences, University of Trento, Italy

###### 2 – Graduate Program in Organismic and Evolutionary Biology, University of Massachusetts Amherst

###### 3 – Kavli Institute for Neuroscience, Department of Neuroscience, Yale University School of Medicine, USA

###### 4 – Department of Biology, University of Massachusetts Amherst

---

This document contains the statistical analysis output for the
manuscript. We also provide the entire raw data on which this analysis
is based. Note that much of the analysis is divided into sections using
tabs, found at the start of the sections. Click on the section you wish
to inspect, and scroll down for the full analysis.

---

### 1 Prepare environment

#### 1.1 loading packages

```
library(data.table) #To deal with big datasets
library(glmmTMB)
library(emmeans)
```

```
## Welcome to emmeans.
## Caution: You lose important information if you filter this package's results.
## See '? untidy'
```

```
library(car)
```

```
## Loading required package: carData
```

```
library(DHARMa)
```

```
## This is DHARMa 0.4.7. For overview type '?DHARMa'. For recent changes, type news(package = 'DHARMa')
```

```
library(ggplot2)
library(future.apply) # For cross-platform parallel processing. If not installed, run: install.packages('future.apply')
```

```
## Warning: package 'future.apply' was built under R version 4.4.3
```

```
## Loading required package: future
```

```
## Warning: package 'future' was built under R version 4.4.3
```

```
set.seed(12345)
```

#### 1.2 loading files

```
#### OPEN DATA ####

full_data <- fread(paste0(data_path, 'all.csv'))
head(full_data)
```

```
##    experimenter   subj     date LeftEye_Center_x LeftEye_Center_y
##          <char> <char>   <char>            <num>            <num>
## 1:         Alex  ID117 16_11_22              NaN              NaN
## 2:         Alex  ID117 16_11_22              NaN              NaN
## 3:         Alex  ID117 16_11_22              NaN              NaN
## 4:         Alex  ID117 16_11_22              NaN              NaN
## 5:         Alex  ID117 16_11_22              NaN              NaN
## 6:         Alex  ID117 16_11_22              NaN              NaN
##    LeftEye_pointShift RightEye_Center_x RightEye_Center_y RightEye_pointShift
##                 <num>             <num>             <num>               <num>
## 1:                NaN               NaN               NaN                 NaN
## 2:                NaN               NaN               NaN                 NaN
## 3:                NaN               NaN               NaN                 NaN
## 4:                NaN               NaN               NaN                 NaN
## 5:                NaN               NaN               NaN                 NaN
## 6:                NaN               NaN               NaN                 NaN
##    MidpointEyes_x MidpointEyes_y MidpointEyes_pointShift dist_from_center
##             <num>          <num>                   <num>            <num>
## 1:            NaN            NaN                     NaN              NaN
## 2:            NaN            NaN                     NaN              NaN
## 3:            NaN            NaN                     NaN              NaN
## 4:            NaN            NaN                     NaN              NaN
## 5:            NaN            NaN                     NaN              NaN
## 6:            NaN            NaN                     NaN              NaN
##    inter_eye_dist inter_eye_dist_change index stimtype stimn moving
##             <num>                 <num> <int>   <char> <int>  <int>
## 1:            NaN                   NaN  9311  ellipse     0      0
## 2:            NaN                   NaN  9312  ellipse     0      0
## 3:            NaN                   NaN  9313  ellipse     0      0
## 4:            NaN                   NaN  9314  ellipse     0      0
## 5:            NaN                   NaN  9315  ellipse     0      0
## 6:            NaN                   NaN  9316  ellipse     0      0
##    frame_from_start     time time_from_start
##               <int>    <num>           <num>
## 1:                1 310.3667      0.03333333
## 2:                2 310.4000      0.06666667
## 3:                3 310.4333      0.10000000
## 4:                4 310.4667      0.13333333
## 5:                5 310.5000      0.16666667
## 6:                6 310.5333      0.20000000
```

```
full_data$date <- as.factor(full_data$date)
full_data$subj <- as.factor(full_data$subj)
full_data$experimenter <- as.factor(full_data$experimenter)
full_data$stimtype <- as.factor(full_data$stimtype)
full_data$moving <- as.factor(full_data$moving)
full_data$displacement <- (full_data$LeftEye_pointShift+full_data$RightEye_pointShift)/2
full_data$midpoint_displacement <- full_data$MidpointEyes_pointShift

full_data
```

```
##         experimenter   subj     date LeftEye_Center_x LeftEye_Center_y
##               <fctr> <fctr>   <fctr>            <num>            <num>
##      1:         Alex  ID117 16_11_22              NaN              NaN
##      2:         Alex  ID117 16_11_22              NaN              NaN
##      3:         Alex  ID117 16_11_22              NaN              NaN
##      4:         Alex  ID117 16_11_22              NaN              NaN
##      5:         Alex  ID117 16_11_22              NaN              NaN
##     ---                                                               
## 886793:          Wes   ID25  7_12_22       -0.2996520        0.5454032
## 886794:          Wes   ID25  7_12_22       -0.2996490        0.5522659
## 886795:          Wes   ID25  7_12_22       -0.2989957        0.5525364
## 886796:          Wes   ID25  7_12_22       -0.2963548        0.5477848
## 886797:          Wes   ID25  7_12_22       -0.2917526        0.5518061
##         LeftEye_pointShift RightEye_Center_x RightEye_Center_y
##                      <num>             <num>             <num>
##      1:                NaN               NaN               NaN
##      2:                NaN               NaN               NaN
##      3:                NaN               NaN               NaN
##      4:                NaN               NaN               NaN
##      5:                NaN               NaN               NaN
##     ---                                                       
## 886793:       0.0013875010        0.06220320         0.5228413
## 886794:       0.0068627102        0.06751060         0.5225280
## 886795:       0.0007070078        0.06724111         0.5218944
## 886796:       0.0054361647        0.05826913         0.5271907
## 886797:       0.0061115610        0.05182759         0.5320094
##         RightEye_pointShift MidpointEyes_x MidpointEyes_y
##                       <num>          <num>          <num>
##      1:                 NaN            NaN            NaN
##      2:                 NaN            NaN            NaN
##      3:                 NaN            NaN            NaN
##      4:                 NaN            NaN            NaN
##      5:                 NaN            NaN            NaN
##     ---                                                  
## 886793:        0.0017485023     -0.1187244      0.5341222
## 886794:        0.0053166338     -0.1160692      0.5373970
## 886795:        0.0006885602     -0.1158773      0.5372154
## 886796:        0.0104186175     -0.1190428      0.5374878
## 886797:        0.0080444188     -0.1199625      0.5419077
##         MidpointEyes_pointShift dist_from_center inter_eye_dist
##                           <num>            <num>          <num>
##      1:                     NaN              NaN            NaN
##      2:                     NaN              NaN            NaN
##      3:                     NaN              NaN            NaN
##      4:                     NaN              NaN            NaN
##      5:                     NaN              NaN            NaN
##     ---                                                        
## 886793:            0.0003794173        0.5471582      0.3625579
## 886794:            0.0042159384        0.5497886      0.3683619
## 886795:            0.0002641490        0.5495707      0.3675165
## 886796:            0.0031772225        0.5505128      0.3552214
## 886797:            0.0045145962        0.5550270      0.3441500
##         inter_eye_dist_change index stimtype stimn moving frame_from_start
##                         <num> <int>   <fctr> <int> <fctr>            <int>
##      1:                   NaN  9311  ellipse     0      0                1
##      2:                   NaN  9312  ellipse     0      0                2
##      3:                   NaN  9313  ellipse     0      0                3
##      4:                   NaN  9314  ellipse     0      0                4
##      5:                   NaN  9315  ellipse     0      0                5
##     ---                                                                   
## 886793:         -0.0030484406 31073     none     9      0              355
## 886794:          0.0058039720 31074     none     9      0              356
## 886795:         -0.0008454018 31075     none     9      0              357
## 886796:         -0.0122950633 31076     none     9      0              358
## 886797:         -0.0110714197 31077     none     9      0              359
##              time time_from_start displacement midpoint_displacement
##             <num>           <num>        <num>                 <num>
##      1:  310.3667      0.03333333          NaN                   NaN
##      2:  310.4000      0.06666667          NaN                   NaN
##      3:  310.4333      0.10000000          NaN                   NaN
##      4:  310.4667      0.13333333          NaN                   NaN
##      5:  310.5000      0.16666667          NaN                   NaN
##     ---                                                             
## 886793: 1035.7667     11.83333333  0.001568002          0.0003794173
## 886794: 1035.8000     11.86666667  0.006089672          0.0042159384
## 886795: 1035.8333     11.90000000  0.000697784          0.0002641490
## 886796: 1035.8667     11.93333333  0.007927391          0.0031772225
## 886797: 1035.9000     11.96666667  0.007077990          0.0045145962
```

```
hist(log(full_data$displacement))
```

```
hist(log(full_data$midpoint_displacement))
```

```
hist(log(full_data$inter_eye_dist))
```

```
hist(log(full_data$inter_eye_dist_change))
```

```
## Warning in log(full_data$inter_eye_dist_change): NaNs produced
```

```
hist(log(full_data$dist_from_center))
```

as expected, log transform is needed for the analysis, as to make
value normal. Let’s drop 0s to avoid model failing

```
full_data$displacement[full_data$displacement==0]<-NA
full_data$midpoint_displacement[full_data$midpoint_displacement==0]<-NA
full_data$inter_eye_dist[full_data$inter_eye_dist==0]<-NA
full_data$inter_eye_dist_change[full_data$inter_eye_dist_change==0]<-NA
```

#### 1.3 Definitions

Define values

```
window_size <- 1 # The size of the analysis windows in seconds
step_size <- 5/30 # The stepsize, how many seconds it shifts

alpha <- 0.01 # The required alpha to consider a value acceptable

# Define the number of cores to use for parallel processing
# We use one less than the total available to keep the system responsive.
num_cores <- future::availableCores() - 4
```

then, I will define functions

```
# I am bad ad writing functions in R. I believe that it would be possible to just pass data and variables, but this is what i am going to do

windowed_model_displacement <-function(window_start){
  gc()
  window_end <- window_start + window_size

  window_data <- subset(data, data$time_from_start>window_start)
  window_data <- subset(window_data, window_data$time_from_start<window_end)

  full_tmb <- glmmTMB(log(midpoint_displacement) ~ stimtype + (stimtype|subj), family=gaussian, data=window_data)
  av <- Anova(full_tmb)
  stimtype_effect <- av$`Pr(>Chisq)`

  e <- emmeans(full_tmb, ~stimtype, type='response')
  e <- as.data.frame(e) # This is for later plotting

    return (c(stimtype_effect, e))

}


windowed_model_inter_eye_dist<-function(window_start){
  gc()
  window_end <- window_start + window_size

  window_data <- subset(data, data$time_from_start>window_start)
  window_data <- subset(window_data, window_data$time_from_start<window_end)

  full_tmb <- glmmTMB(log(inter_eye_dist) ~ stimtype + (stimtype|subj), family=gaussian, data=window_data)
  av <- Anova(full_tmb)
  stimtype_effect <- av$`Pr(>Chisq)`

  e <- emmeans(full_tmb, ~stimtype, type='response')
  e <- as.data.frame(e) # This is for later plotting

    return (c(stimtype_effect, e))

}


windowed_model_inter_eye_dist_change<-function(window_start){
  gc()
  window_end <- window_start + window_size

  window_data <- subset(data, data$time_from_start>window_start)
  window_data <- subset(window_data, window_data$time_from_start<window_end)

  full_tmb <- glmmTMB(log(inter_eye_dist_change) ~ stimtype + (stimtype|subj), family=gaussian, data=window_data)
  av <- Anova(full_tmb)
  stimtype_effect <- av$`Pr(>Chisq)`

  e <- emmeans(full_tmb, ~stimtype, type='response')
  e <- as.data.frame(e) # This is for later plotting

    return (c(stimtype_effect, e))

}


windowed_model_dist_from_center<-function(window_start){
  gc()
  window_end <- window_start + window_size

  window_data <- subset(data, data$time_from_start>window_start)
  window_data <- subset(window_data, window_data$time_from_start<window_end)

  full_tmb <- glmmTMB(log(dist_from_center) ~ stimtype + (stimtype|subj), family=gaussian, data=window_data)
  av <- Anova(full_tmb)
  stimtype_effect <- av$`Pr(>Chisq)`

  e <- emmeans(full_tmb, ~stimtype, type='response')
  e <- as.data.frame(e) # This is for later plotting

    return (c(stimtype_effect, e))

}
```

### 2 Analysis

We have three variables:

- The per frame eye displacement, meaning the speed.
- The distance between the two eyes
- The change of the distance between the two eyes (can be similar to
  eye displacement, but only has the inter eye component)

Let’s start the analysis one by one.

#### 2.1 Whole time-series pass

First, we do time series analysis. We will proceed with anova and
post-hoc only for windows that result significant.

##### 2.1.1 Eye displacement

Each analysis will proceed in two parts. first, I will analyze the
static section, before stimuli start moving. This is compared with a
section of no stimulus. This is to test weather the spider will scan the
static stimuli or not.

In the second phase, I will drop the no-stimulus control and test for
the whole stimulus duration.

###### 2.1.1.1 Static

```
onlystatic <- subset(full_data, full_data$moving == 0)
onlynonemaxframe <- max(onlystatic[onlystatic$stimtype=='none']$time_from_start)
onlystatic <- subset(onlystatic, onlystatic$time_from_start<onlynonemaxframe)

start_frame <- min(onlystatic$time_from_start)
end_frame <- max(onlystatic$time_from_start)

windows_starts <- seq(start_frame, end_frame-window_size, step_size)

data <- onlystatic
plan(multisession, workers = num_cores)
windows_p_static <- future_lapply(windows_starts, windowed_model_displacement, future.seed=TRUE)
```

```
## Warning: package 'future' was built under R version 4.4.3
## Warning: package 'future' was built under R version 4.4.3
## Warning: package 'future' was built under R version 4.4.3
## Warning: package 'future' was built under R version 4.4.3
## Warning: package 'future' was built under R version 4.4.3
## Warning: package 'future' was built under R version 4.4.3
## Warning: package 'future' was built under R version 4.4.3
```

```
## Warning in finalizeTMB(TMBStruc, obj, fit, h, data.tmb.old): Model convergence
## problem; non-positive-definite Hessian matrix. See vignette('troubleshooting')
## Warning in finalizeTMB(TMBStruc, obj, fit, h, data.tmb.old): Model convergence
## problem; non-positive-definite Hessian matrix. See vignette('troubleshooting')
```

```
## Warning: package 'future' was built under R version 4.4.3
```

Saving routine

```
static_results <- data.frame(list('win_starts'=NaN,
                                  'win_span'=NaN,
                                  'p'=NaN,
                                  'stimtype'=NaN,
'emmean'=NaN, 'SE'=NaN, 'CL.upper'=NaN, 'CL.lower'=NaN))

for (i in 1:length(windows_starts)){
  window_start <- windows_starts[i]
  p <- windows_p_static[[i]][[1]]
  for (stimtype in c('none', 'bio', 'ellipse', 'scramb', 'silhouette', 'rand')){
  newrow <- c(window_start, window_size, p, stimtype,
              windows_p_static[[i]]$response[windows_p_static[[i]]$stimtype==stimtype],
  windows_p_static[[i]]$SE[windows_p_static[[i]]$stimtype==stimtype],
              windows_p_static[[i]]$upper.CL[windows_p_static[[i]]$stimtype==stimtype],
  windows_p_static[[i]]$lower.CL[windows_p_static[[i]]$stimtype==stimtype])
    static_results <- rbind(static_results, newrow)
  }

}

static_results <- static_results[-1,]

write.csv(static_results, paste0(tmp_path, 'displacement_staticResults.csv'))
```

###### 2.1.1.2 Moving

```
onlymoving <- subset(full_data, full_data$stimtype != 'none')

start_frame <- min(onlymoving$time_from_start)
end_frame <- max(onlymoving$time_from_start)

windows_starts <- seq(start_frame, end_frame-window_size, step_size)
winsizes <- rep(window_size, length(windows_starts))

data <- onlymoving
plan(multisession, workers = num_cores)
windows_p_moving <- future_lapply(windows_starts, windowed_model_displacement, future.seed=TRUE)
```

```
## Warning in finalizeTMB(TMBStruc, obj, fit, h, data.tmb.old): Model convergence
## problem; non-positive-definite Hessian matrix. See vignette('troubleshooting')
```

```
## Warning in finalizeTMB(TMBStruc, obj, fit, h, data.tmb.old): Model convergence
## problem; singular convergence (7). See vignette('troubleshooting'),
## help('diagnose')
```

```
## Warning in finalizeTMB(TMBStruc, obj, fit, h, data.tmb.old): Model convergence
## problem; non-positive-definite Hessian matrix. See vignette('troubleshooting')
```

```
## Warning in finalizeTMB(TMBStruc, obj, fit, h, data.tmb.old): Model convergence
## problem; false convergence (8). See vignette('troubleshooting'),
## help('diagnose')
```

```
## Warning in finalizeTMB(TMBStruc, obj, fit, h, data.tmb.old): Model convergence
## problem; non-positive-definite Hessian matrix. See vignette('troubleshooting')
```

```
## Warning in finalizeTMB(TMBStruc, obj, fit, h, data.tmb.old): Model convergence
## problem; false convergence (8). See vignette('troubleshooting'),
## help('diagnose')
```

```
## Warning in finalizeTMB(TMBStruc, obj, fit, h, data.tmb.old): Model convergence
## problem; non-positive-definite Hessian matrix. See vignette('troubleshooting')
## Warning in finalizeTMB(TMBStruc, obj, fit, h, data.tmb.old): Model convergence
## problem; non-positive-definite Hessian matrix. See vignette('troubleshooting')
## Warning in finalizeTMB(TMBStruc, obj, fit, h, data.tmb.old): Model convergence
## problem; non-positive-definite Hessian matrix. See vignette('troubleshooting')
```

```
## Warning in finalizeTMB(TMBStruc, obj, fit, h, data.tmb.old): Model convergence
## problem; false convergence (8). See vignette('troubleshooting'),
## help('diagnose')
```

```
## Warning in finalizeTMB(TMBStruc, obj, fit, h, data.tmb.old): Model convergence
## problem; non-positive-definite Hessian matrix. See vignette('troubleshooting')
```

Saving routine

```
moving_results <- data.frame(list('win_starts'=NaN,
                                  'win_span'=NaN,
                                  'p'=NaN,
                                  'stimtype'=NaN,
'emmean'=NaN, 'SE'=NaN, 'CL.upper'=NaN, 'CL.lower'=NaN))
for (i in 1:length(windows_starts)){
  window_start <- windows_starts[i]
  p <- windows_p_moving[[i]][[1]]
  for (stimtype in c('bio', 'ellipse', 'scramb', 'silhouette', 'rand')){
  newrow <- c(window_start, window_size, p, stimtype,
              windows_p_moving[[i]]$response[windows_p_moving[[i]]$stimtype==stimtype],
  windows_p_moving[[i]]$SE[windows_p_moving[[i]]$stimtype==stimtype],
              windows_p_moving[[i]]$upper.CL[windows_p_moving[[i]]$stimtype==stimtype],
  windows_p_moving[[i]]$lower.CL[windows_p_moving[[i]]$stimtype==stimtype])
    moving_results <- rbind(moving_results, newrow)
  }

}

moving_results <- moving_results[-1,]

write.csv(moving_results, paste0(tmp_path, 'displacement_movingResults.csv'))
```

##### 2.1.2 Inter-eye distance

###### 2.1.2.1 Static

```
onlystatic <- subset(full_data, full_data$moving == 0)
onlynonemaxframe <- max(onlystatic[onlystatic$stimtype=='none']$time_from_start)
onlystatic <- subset(onlystatic, onlystatic$time_from_start<onlynonemaxframe)

start_frame <- min(onlystatic$time_from_start)
end_frame <- max(onlystatic$time_from_start)

windows_starts <- seq(start_frame, end_frame-window_size, step_size)

data <- onlystatic
plan(multisession, workers = num_cores)
windows_p_static <- future_lapply(windows_starts, windowed_model_inter_eye_dist, future.seed=TRUE)
```

Saving routine

```
static_results <- data.frame(list('win_starts'=NaN,
                                  'win_span'=NaN,
                                  'p'=NaN,
                                  'stimtype'=NaN,
'emmean'=NaN, 'SE'=NaN, 'CL.upper'=NaN, 'CL.lower'=NaN))

for (i in 1:length(windows_starts)){
  window_start <- windows_starts[i]
  p <- windows_p_static[[i]][[1]]
  for (stimtype in c('none', 'bio', 'ellipse', 'scramb', 'silhouette', 'rand')){
  newrow <- c(window_start, window_size, p, stimtype,
              windows_p_static[[i]]$response[windows_p_static[[i]]$stimtype==stimtype],
  windows_p_static[[i]]$SE[windows_p_static[[i]]$stimtype==stimtype],
              windows_p_static[[i]]$upper.CL[windows_p_static[[i]]$stimtype==stimtype],
  windows_p_static[[i]]$lower.CL[windows_p_static[[i]]$stimtype==stimtype])
    static_results <- rbind(static_results, newrow)
  }

}

static_results <- static_results[-1,]

write.csv(static_results, paste0(tmp_path, 'interEyeDist_staticResults.csv'))
```

###### 2.1.2.2 Moving

```
onlymoving <- subset(full_data, full_data$stimtype != 'none')

start_frame <- min(onlymoving$time_from_start)
end_frame <- max(onlymoving$time_from_start)

windows_starts <- seq(start_frame, end_frame-window_size, step_size)
winsizes <- rep(window_size, length(windows_starts))

data <- onlymoving
plan(multisession, workers = num_cores)
windows_p_moving <- future_lapply(windows_starts, windowed_model_inter_eye_dist, future.seed=TRUE)
```

Saving routine

```
moving_results <- data.frame(list('win_starts'=NaN,
                                  'win_span'=NaN,
                                  'p'=NaN,
                                  'stimtype'=NaN,
'emmean'=NaN, 'SE'=NaN, 'CL.upper'=NaN, 'CL.lower'=NaN))
for (i in 1:length(windows_starts)){
  window_start <- windows_starts[i]
  p <- windows_p_moving[[i]][[1]]
  for (stimtype in c('bio', 'ellipse', 'scramb', 'silhouette', 'rand')){
  newrow <- c(window_start, window_size, p, stimtype,
              windows_p_moving[[i]]$response[windows_p_moving[[i]]$stimtype==stimtype],
  windows_p_moving[[i]]$SE[windows_p_moving[[i]]$stimtype==stimtype],
              windows_p_moving[[i]]$upper.CL[windows_p_moving[[i]]$stimtype==stimtype],
  windows_p_moving[[i]]$lower.CL[windows_p_moving[[i]]$stimtype==stimtype])
    moving_results <- rbind(moving_results, newrow)
  }

}

moving_results <- moving_results[-1,]

write.csv(moving_results, paste0(tmp_path, 'interEyeDist_movingResults.csv'))
```

##### 2.1.3 Inter-eye distance change

###### 2.1.3.1 Static

```
onlystatic <- subset(full_data, full_data$moving == 0)
onlynonemaxframe <- max(onlystatic[onlystatic$stimtype=='none']$time_from_start)
onlystatic <- subset(onlystatic, onlystatic$time_from_start<onlynonemaxframe)

start_frame <- min(onlystatic$time_from_start)
end_frame <- max(onlystatic$time_from_start)

windows_starts <- seq(start_frame, end_frame-window_size, step_size)

data <- onlystatic
plan(multisession, workers = num_cores)
windows_p_static <- future_lapply(windows_starts, windowed_model_inter_eye_dist_change, future.seed=TRUE)
```

```
## Warning in log(inter_eye_dist_change): NaNs produced
## Warning in log(inter_eye_dist_change): NaNs produced
## Warning in log(inter_eye_dist_change): NaNs produced
## Warning in log(inter_eye_dist_change): NaNs produced
```

```
## Warning in finalizeTMB(TMBStruc, obj, fit, h, data.tmb.old): Model convergence
## problem; non-positive-definite Hessian matrix. See vignette('troubleshooting')
```

```
## Warning in finalizeTMB(TMBStruc, obj, fit, h, data.tmb.old): Model convergence
## problem; singular convergence (7). See vignette('troubleshooting'),
## help('diagnose')
```

```
## Warning in log(inter_eye_dist_change): NaNs produced
## Warning in log(inter_eye_dist_change): NaNs produced
```

```
## Warning in finalizeTMB(TMBStruc, obj, fit, h, data.tmb.old): Model convergence
## problem; singular convergence (7). See vignette('troubleshooting'),
## help('diagnose')
```

```
## Warning in log(inter_eye_dist_change): NaNs produced
## Warning in log(inter_eye_dist_change): NaNs produced
```

```
## Warning in finalizeTMB(TMBStruc, obj, fit, h, data.tmb.old): Model convergence
## problem; non-positive-definite Hessian matrix. See vignette('troubleshooting')
```

```
## Warning in finalizeTMB(TMBStruc, obj, fit, h, data.tmb.old): Model convergence
## problem; singular convergence (7). See vignette('troubleshooting'),
## help('diagnose')
```

```
## Warning in log(inter_eye_dist_change): NaNs produced
## Warning in log(inter_eye_dist_change): NaNs produced
## Warning in log(inter_eye_dist_change): NaNs produced
## Warning in log(inter_eye_dist_change): NaNs produced
```

```
## Warning in finalizeTMB(TMBStruc, obj, fit, h, data.tmb.old): Model convergence
## problem; non-positive-definite Hessian matrix. See vignette('troubleshooting')
```

```
## Warning in log(inter_eye_dist_change): NaNs produced
## Warning in log(inter_eye_dist_change): NaNs produced
```

```
## Warning in finalizeTMB(TMBStruc, obj, fit, h, data.tmb.old): Model convergence
## problem; singular convergence (7). See vignette('troubleshooting'),
## help('diagnose')
```

```
## Warning in log(inter_eye_dist_change): NaNs produced
## Warning in log(inter_eye_dist_change): NaNs produced
```

```
## Warning in finalizeTMB(TMBStruc, obj, fit, h, data.tmb.old): Model convergence
## problem; non-positive-definite Hessian matrix. See vignette('troubleshooting')
```

```
## Warning in log(inter_eye_dist_change): NaNs produced
## Warning in log(inter_eye_dist_change): NaNs produced
```

```
## Warning in finalizeTMB(TMBStruc, obj, fit, h, data.tmb.old): Model convergence
## problem; non-positive-definite Hessian matrix. See vignette('troubleshooting')
```

```
## Warning in finalizeTMB(TMBStruc, obj, fit, h, data.tmb.old): Model convergence
## problem; false convergence (8). See vignette('troubleshooting'),
## help('diagnose')
```

```
## Warning in log(inter_eye_dist_change): NaNs produced
## Warning in log(inter_eye_dist_change): NaNs produced
```

```
## Warning in finalizeTMB(TMBStruc, obj, fit, h, data.tmb.old): Model convergence
## problem; non-positive-definite Hessian matrix. See vignette('troubleshooting')
```

```
## Warning in finalizeTMB(TMBStruc, obj, fit, h, data.tmb.old): Model convergence
## problem; singular convergence (7). See vignette('troubleshooting'),
## help('diagnose')
```

```
## Warning in log(inter_eye_dist_change): NaNs produced
## Warning in log(inter_eye_dist_change): NaNs produced
```

```
## Warning in finalizeTMB(TMBStruc, obj, fit, h, data.tmb.old): Model convergence
## problem; non-positive-definite Hessian matrix. See vignette('troubleshooting')
```

```
## Warning in finalizeTMB(TMBStruc, obj, fit, h, data.tmb.old): Model convergence
## problem; singular convergence (7). See vignette('troubleshooting'),
## help('diagnose')
```

```
## Warning in log(inter_eye_dist_change): NaNs produced
## Warning in log(inter_eye_dist_change): NaNs produced
```

```
## Warning in finalizeTMB(TMBStruc, obj, fit, h, data.tmb.old): Model convergence
## problem; non-positive-definite Hessian matrix. See vignette('troubleshooting')
```

```
## Warning in finalizeTMB(TMBStruc, obj, fit, h, data.tmb.old): Model convergence
## problem; singular convergence (7). See vignette('troubleshooting'),
## help('diagnose')
```

```
## Warning in log(inter_eye_dist_change): NaNs produced
## Warning in log(inter_eye_dist_change): NaNs produced
```

```
## Warning in finalizeTMB(TMBStruc, obj, fit, h, data.tmb.old): Model convergence
## problem; non-positive-definite Hessian matrix. See vignette('troubleshooting')
```

```
## Warning in log(inter_eye_dist_change): NaNs produced
## Warning in log(inter_eye_dist_change): NaNs produced
```

```
## Warning in finalizeTMB(TMBStruc, obj, fit, h, data.tmb.old): Model convergence
## problem; non-positive-definite Hessian matrix. See vignette('troubleshooting')
```

```
## Warning in log(inter_eye_dist_change): NaNs produced
## Warning in log(inter_eye_dist_change): NaNs produced
## Warning in log(inter_eye_dist_change): NaNs produced
## Warning in log(inter_eye_dist_change): NaNs produced
```

```
## Warning in finalizeTMB(TMBStruc, obj, fit, h, data.tmb.old): Model convergence
## problem; non-positive-definite Hessian matrix. See vignette('troubleshooting')
```

```
## Warning in log(inter_eye_dist_change): NaNs produced
## Warning in log(inter_eye_dist_change): NaNs produced
## Warning in log(inter_eye_dist_change): NaNs produced
## Warning in log(inter_eye_dist_change): NaNs produced
## Warning in log(inter_eye_dist_change): NaNs produced
## Warning in log(inter_eye_dist_change): NaNs produced
## Warning in log(inter_eye_dist_change): NaNs produced
## Warning in log(inter_eye_dist_change): NaNs produced
```

```
## Warning in finalizeTMB(TMBStruc, obj, fit, h, data.tmb.old): Model convergence
## problem; non-positive-definite Hessian matrix. See vignette('troubleshooting')
```

```
## Warning in finalizeTMB(TMBStruc, obj, fit, h, data.tmb.old): Model convergence
## problem; singular convergence (7). See vignette('troubleshooting'),
## help('diagnose')
```

```
## Warning in log(inter_eye_dist_change): NaNs produced
## Warning in log(inter_eye_dist_change): NaNs produced
```

```
## Warning in finalizeTMB(TMBStruc, obj, fit, h, data.tmb.old): Model convergence
## problem; non-positive-definite Hessian matrix. See vignette('troubleshooting')
```

```
## Warning in finalizeTMB(TMBStruc, obj, fit, h, data.tmb.old): Model convergence
## problem; singular convergence (7). See vignette('troubleshooting'),
## help('diagnose')
```

```
## Warning in log(inter_eye_dist_change): NaNs produced
## Warning in log(inter_eye_dist_change): NaNs produced
```

```
## Warning in finalizeTMB(TMBStruc, obj, fit, h, data.tmb.old): Model convergence
## problem; non-positive-definite Hessian matrix. See vignette('troubleshooting')
```

```
## Warning in finalizeTMB(TMBStruc, obj, fit, h, data.tmb.old): Model convergence
## problem; singular convergence (7). See vignette('troubleshooting'),
## help('diagnose')
```

```
## Warning in log(inter_eye_dist_change): NaNs produced
## Warning in log(inter_eye_dist_change): NaNs produced
```

```
## Warning in finalizeTMB(TMBStruc, obj, fit, h, data.tmb.old): Model convergence
## problem; non-positive-definite Hessian matrix. See vignette('troubleshooting')
```

```
## Warning in finalizeTMB(TMBStruc, obj, fit, h, data.tmb.old): Model convergence
## problem; singular convergence (7). See vignette('troubleshooting'),
## help('diagnose')
```

```
## Warning in log(inter_eye_dist_change): NaNs produced
## Warning in log(inter_eye_dist_change): NaNs produced
## Warning in log(inter_eye_dist_change): NaNs produced
## Warning in log(inter_eye_dist_change): NaNs produced
```

```
## Warning in finalizeTMB(TMBStruc, obj, fit, h, data.tmb.old): Model convergence
## problem; non-positive-definite Hessian matrix. See vignette('troubleshooting')
```

```
## Warning in finalizeTMB(TMBStruc, obj, fit, h, data.tmb.old): Model convergence
## problem; false convergence (8). See vignette('troubleshooting'),
## help('diagnose')
```

```
## Warning in log(inter_eye_dist_change): NaNs produced
## Warning in log(inter_eye_dist_change): NaNs produced
```

```
## Warning in finalizeTMB(TMBStruc, obj, fit, h, data.tmb.old): Model convergence
## problem; non-positive-definite Hessian matrix. See vignette('troubleshooting')
```

```
## Warning in finalizeTMB(TMBStruc, obj, fit, h, data.tmb.old): Model convergence
## problem; singular convergence (7). See vignette('troubleshooting'),
## help('diagnose')
```

```
## Warning in log(inter_eye_dist_change): NaNs produced
## Warning in log(inter_eye_dist_change): NaNs produced
```

```
## Warning in finalizeTMB(TMBStruc, obj, fit, h, data.tmb.old): Model convergence
## problem; non-positive-definite Hessian matrix. See vignette('troubleshooting')
```

```
## Warning in finalizeTMB(TMBStruc, obj, fit, h, data.tmb.old): Model convergence
## problem; singular convergence (7). See vignette('troubleshooting'),
## help('diagnose')
```

```
## Warning in log(inter_eye_dist_change): NaNs produced
## Warning in log(inter_eye_dist_change): NaNs produced
```

```
## Warning in finalizeTMB(TMBStruc, obj, fit, h, data.tmb.old): Model convergence
## problem; singular convergence (7). See vignette('troubleshooting'),
## help('diagnose')
```

```
## Warning in log(inter_eye_dist_change): NaNs produced
## Warning in log(inter_eye_dist_change): NaNs produced
```

```
## Warning in finalizeTMB(TMBStruc, obj, fit, h, data.tmb.old): Model convergence
## problem; non-positive-definite Hessian matrix. See vignette('troubleshooting')
```

```
## Warning in log(inter_eye_dist_change): NaNs produced
## Warning in log(inter_eye_dist_change): NaNs produced
## Warning in log(inter_eye_dist_change): NaNs produced
## Warning in log(inter_eye_dist_change): NaNs produced
## Warning in log(inter_eye_dist_change): NaNs produced
## Warning in log(inter_eye_dist_change): NaNs produced
## Warning in log(inter_eye_dist_change): NaNs produced
## Warning in log(inter_eye_dist_change): NaNs produced
## Warning in log(inter_eye_dist_change): NaNs produced
## Warning in log(inter_eye_dist_change): NaNs produced
## Warning in log(inter_eye_dist_change): NaNs produced
## Warning in log(inter_eye_dist_change): NaNs produced
```

```
## Warning in finalizeTMB(TMBStruc, obj, fit, h, data.tmb.old): Model convergence
## problem; non-positive-definite Hessian matrix. See vignette('troubleshooting')
```

```
## Warning in finalizeTMB(TMBStruc, obj, fit, h, data.tmb.old): Model convergence
## problem; singular convergence (7). See vignette('troubleshooting'),
## help('diagnose')
```

```
## Warning in log(inter_eye_dist_change): NaNs produced
## Warning in log(inter_eye_dist_change): NaNs produced
```

```
## Warning in finalizeTMB(TMBStruc, obj, fit, h, data.tmb.old): Model convergence
## problem; non-positive-definite Hessian matrix. See vignette('troubleshooting')
```

```
## Warning in log(inter_eye_dist_change): NaNs produced
## Warning in log(inter_eye_dist_change): NaNs produced
```

```
## Warning in finalizeTMB(TMBStruc, obj, fit, h, data.tmb.old): Model convergence
## problem; singular convergence (7). See vignette('troubleshooting'),
## help('diagnose')
```

```
## Warning in log(inter_eye_dist_change): NaNs produced
## Warning in log(inter_eye_dist_change): NaNs produced
```

```
## Warning in finalizeTMB(TMBStruc, obj, fit, h, data.tmb.old): Model convergence
## problem; non-positive-definite Hessian matrix. See vignette('troubleshooting')
```

```
## Warning in finalizeTMB(TMBStruc, obj, fit, h, data.tmb.old): Model convergence
## problem; false convergence (8). See vignette('troubleshooting'),
## help('diagnose')
```

```
## Warning in log(inter_eye_dist_change): NaNs produced
## Warning in log(inter_eye_dist_change): NaNs produced
```

```
## Warning in finalizeTMB(TMBStruc, obj, fit, h, data.tmb.old): Model convergence
## problem; non-positive-definite Hessian matrix. See vignette('troubleshooting')
```

```
## Warning in finalizeTMB(TMBStruc, obj, fit, h, data.tmb.old): Model convergence
## problem; false convergence (8). See vignette('troubleshooting'),
## help('diagnose')
```

```
## Warning in log(inter_eye_dist_change): NaNs produced
## Warning in log(inter_eye_dist_change): NaNs produced
```

```
## Warning in finalizeTMB(TMBStruc, obj, fit, h, data.tmb.old): Model convergence
## problem; non-positive-definite Hessian matrix. See vignette('troubleshooting')
```

```
## Warning in log(inter_eye_dist_change): NaNs produced
## Warning in log(inter_eye_dist_change): NaNs produced
```

```
## Warning in finalizeTMB(TMBStruc, obj, fit, h, data.tmb.old): Model convergence
## problem; non-positive-definite Hessian matrix. See vignette('troubleshooting')
```

```
## Warning in finalizeTMB(TMBStruc, obj, fit, h, data.tmb.old): Model convergence
## problem; singular convergence (7). See vignette('troubleshooting'),
## help('diagnose')
```

```
## Warning in log(inter_eye_dist_change): NaNs produced
## Warning in log(inter_eye_dist_change): NaNs produced
```

```
## Warning in finalizeTMB(TMBStruc, obj, fit, h, data.tmb.old): Model convergence
## problem; non-positive-definite Hessian matrix. See vignette('troubleshooting')
```

```
## Warning in finalizeTMB(TMBStruc, obj, fit, h, data.tmb.old): Model convergence
## problem; false convergence (8). See vignette('troubleshooting'),
## help('diagnose')
```

```
## Warning in log(inter_eye_dist_change): NaNs produced
## Warning in log(inter_eye_dist_change): NaNs produced
```

```
## Warning in finalizeTMB(TMBStruc, obj, fit, h, data.tmb.old): Model convergence
## problem; non-positive-definite Hessian matrix. See vignette('troubleshooting')
```

```
## Warning in log(inter_eye_dist_change): NaNs produced
## Warning in log(inter_eye_dist_change): NaNs produced
```

```
## Warning in finalizeTMB(TMBStruc, obj, fit, h, data.tmb.old): Model convergence
## problem; non-positive-definite Hessian matrix. See vignette('troubleshooting')
```

```
## Warning in finalizeTMB(TMBStruc, obj, fit, h, data.tmb.old): Model convergence
## problem; false convergence (8). See vignette('troubleshooting'),
## help('diagnose')
```

```
## Warning in log(inter_eye_dist_change): NaNs produced
## Warning in log(inter_eye_dist_change): NaNs produced
```

```
## Warning in finalizeTMB(TMBStruc, obj, fit, h, data.tmb.old): Model convergence
## problem; non-positive-definite Hessian matrix. See vignette('troubleshooting')
```

```
## Warning in finalizeTMB(TMBStruc, obj, fit, h, data.tmb.old): Model convergence
## problem; singular convergence (7). See vignette('troubleshooting'),
## help('diagnose')
```

```
## Warning in log(inter_eye_dist_change): NaNs produced
## Warning in log(inter_eye_dist_change): NaNs produced
```

```
## Warning in finalizeTMB(TMBStruc, obj, fit, h, data.tmb.old): Model convergence
## problem; non-positive-definite Hessian matrix. See vignette('troubleshooting')
```

```
## Warning in finalizeTMB(TMBStruc, obj, fit, h, data.tmb.old): Model convergence
## problem; false convergence (8). See vignette('troubleshooting'),
## help('diagnose')
```

```
## Warning in log(inter_eye_dist_change): NaNs produced
## Warning in log(inter_eye_dist_change): NaNs produced
```

```
## Warning in finalizeTMB(TMBStruc, obj, fit, h, data.tmb.old): Model convergence
## problem; non-positive-definite Hessian matrix. See vignette('troubleshooting')
```

```
## Warning in log(inter_eye_dist_change): NaNs produced
## Warning in log(inter_eye_dist_change): NaNs produced
```

```
## Warning in finalizeTMB(TMBStruc, obj, fit, h, data.tmb.old): Model convergence
## problem; non-positive-definite Hessian matrix. See vignette('troubleshooting')
```

```
## Warning in finalizeTMB(TMBStruc, obj, fit, h, data.tmb.old): Model convergence
## problem; singular convergence (7). See vignette('troubleshooting'),
## help('diagnose')
```

```
## Warning in log(inter_eye_dist_change): NaNs produced
## Warning in log(inter_eye_dist_change): NaNs produced
```

```
## Warning in finalizeTMB(TMBStruc, obj, fit, h, data.tmb.old): Model convergence
## problem; non-positive-definite Hessian matrix. See vignette('troubleshooting')
```

```
## Warning in log(inter_eye_dist_change): NaNs produced
## Warning in log(inter_eye_dist_change): NaNs produced
```

```
## Warning in finalizeTMB(TMBStruc, obj, fit, h, data.tmb.old): Model convergence
## problem; non-positive-definite Hessian matrix. See vignette('troubleshooting')
```

```
## Warning in log(inter_eye_dist_change): NaNs produced
## Warning in log(inter_eye_dist_change): NaNs produced
```

```
## Warning in finalizeTMB(TMBStruc, obj, fit, h, data.tmb.old): Model convergence
## problem; non-positive-definite Hessian matrix. See vignette('troubleshooting')
```

```
## Warning in finalizeTMB(TMBStruc, obj, fit, h, data.tmb.old): Model convergence
## problem; false convergence (8). See vignette('troubleshooting'),
## help('diagnose')
```

```
## Warning in log(inter_eye_dist_change): NaNs produced
## Warning in log(inter_eye_dist_change): NaNs produced
## Warning in log(inter_eye_dist_change): NaNs produced
## Warning in log(inter_eye_dist_change): NaNs produced
```

```
## Warning in finalizeTMB(TMBStruc, obj, fit, h, data.tmb.old): Model convergence
## problem; non-positive-definite Hessian matrix. See vignette('troubleshooting')
```

```
## Warning in finalizeTMB(TMBStruc, obj, fit, h, data.tmb.old): Model convergence
## problem; singular convergence (7). See vignette('troubleshooting'),
## help('diagnose')
```

```
## Warning in log(inter_eye_dist_change): NaNs produced
## Warning in log(inter_eye_dist_change): NaNs produced
```

```
## Warning in finalizeTMB(TMBStruc, obj, fit, h, data.tmb.old): Model convergence
## problem; non-positive-definite Hessian matrix. See vignette('troubleshooting')
```

```
## Warning in log(inter_eye_dist_change): NaNs produced
## Warning in log(inter_eye_dist_change): NaNs produced
```

```
## Warning in finalizeTMB(TMBStruc, obj, fit, h, data.tmb.old): Model convergence
## problem; non-positive-definite Hessian matrix. See vignette('troubleshooting')
```

```
## Warning in finalizeTMB(TMBStruc, obj, fit, h, data.tmb.old): Model convergence
## problem; false convergence (8). See vignette('troubleshooting'),
## help('diagnose')
```

```
## Warning in log(inter_eye_dist_change): NaNs produced
## Warning in log(inter_eye_dist_change): NaNs produced
## Warning in log(inter_eye_dist_change): NaNs produced
## Warning in log(inter_eye_dist_change): NaNs produced
## Warning in log(inter_eye_dist_change): NaNs produced
## Warning in log(inter_eye_dist_change): NaNs produced
```

```
## Warning in finalizeTMB(TMBStruc, obj, fit, h, data.tmb.old): Model convergence
## problem; singular convergence (7). See vignette('troubleshooting'),
## help('diagnose')
```

```
## Warning in log(inter_eye_dist_change): NaNs produced
## Warning in log(inter_eye_dist_change): NaNs produced
## Warning in log(inter_eye_dist_change): NaNs produced
## Warning in log(inter_eye_dist_change): NaNs produced
```

```
## Warning in finalizeTMB(TMBStruc, obj, fit, h, data.tmb.old): Model convergence
## problem; non-positive-definite Hessian matrix. See vignette('troubleshooting')
```

```
## Warning in log(inter_eye_dist_change): NaNs produced
## Warning in log(inter_eye_dist_change): NaNs produced
```

```
## Warning in finalizeTMB(TMBStruc, obj, fit, h, data.tmb.old): Model convergence
## problem; non-positive-definite Hessian matrix. See vignette('troubleshooting')
```

```
## Warning in finalizeTMB(TMBStruc, obj, fit, h, data.tmb.old): Model convergence
## problem; singular convergence (7). See vignette('troubleshooting'),
## help('diagnose')
```

```
## Warning in log(inter_eye_dist_change): NaNs produced
## Warning in log(inter_eye_dist_change): NaNs produced
```

```
## Warning in finalizeTMB(TMBStruc, obj, fit, h, data.tmb.old): Model convergence
## problem; non-positive-definite Hessian matrix. See vignette('troubleshooting')
```

```
## Warning in finalizeTMB(TMBStruc, obj, fit, h, data.tmb.old): Model convergence
## problem; singular convergence (7). See vignette('troubleshooting'),
## help('diagnose')
```

```
## Warning in log(inter_eye_dist_change): NaNs produced
## Warning in log(inter_eye_dist_change): NaNs produced
```

```
## Warning in finalizeTMB(TMBStruc, obj, fit, h, data.tmb.old): Model convergence
## problem; non-positive-definite Hessian matrix. See vignette('troubleshooting')
```

```
## Warning in finalizeTMB(TMBStruc, obj, fit, h, data.tmb.old): Model convergence
## problem; singular convergence (7). See vignette('troubleshooting'),
## help('diagnose')
```

```
## Warning in log(inter_eye_dist_change): NaNs produced
## Warning in log(inter_eye_dist_change): NaNs produced
## Warning in log(inter_eye_dist_change): NaNs produced
## Warning in log(inter_eye_dist_change): NaNs produced
## Warning in log(inter_eye_dist_change): NaNs produced
## Warning in log(inter_eye_dist_change): NaNs produced
```

Saving routine

```
static_results <- data.frame(list('win_starts'=NaN,
                                  'win_span'=NaN,
                                  'p'=NaN,
                                  'stimtype'=NaN,
'emmean'=NaN, 'SE'=NaN, 'CL.upper'=NaN, 'CL.lower'=NaN))

for (i in 1:length(windows_starts)){
  window_start <- windows_starts[i]
  p <- windows_p_static[[i]][[1]]
  for (stimtype in c('none', 'bio', 'ellipse', 'scramb', 'silhouette', 'rand')){
  newrow <- c(window_start, window_size, p, stimtype,
              windows_p_static[[i]]$response[windows_p_static[[i]]$stimtype==stimtype],
  windows_p_static[[i]]$SE[windows_p_static[[i]]$stimtype==stimtype],
              windows_p_static[[i]]$upper.CL[windows_p_static[[i]]$stimtype==stimtype],
  windows_p_static[[i]]$lower.CL[windows_p_static[[i]]$stimtype==stimtype])
    static_results <- rbind(static_results, newrow)
  }

}

static_results <- static_results[-1,]

write.csv(static_results, paste0(tmp_path, 'interEyeDistChange_staticResults.csv'))
```

###### 2.1.3.2 Moving

```
onlymoving <- subset(full_data, full_data$stimtype != 'none')

start_frame <- min(onlymoving$time_from_start)
end_frame <- max(onlymoving$time_from_start)

windows_starts <- seq(start_frame, end_frame-window_size, step_size)
winsizes <- rep(window_size, length(windows_starts))

data <- onlymoving
plan(multisession, workers = num_cores)
windows_p_moving <- future_lapply(windows_starts, windowed_model_inter_eye_dist_change, future.seed=TRUE)
```

```
## Warning in log(inter_eye_dist_change): NaNs produced
## Warning in log(inter_eye_dist_change): NaNs produced
```

```
## Warning in finalizeTMB(TMBStruc, obj, fit, h, data.tmb.old): Model convergence
## problem; non-positive-definite Hessian matrix. See vignette('troubleshooting')
```

```
## Warning in log(inter_eye_dist_change): NaNs produced
## Warning in log(inter_eye_dist_change): NaNs produced
## Warning in log(inter_eye_dist_change): NaNs produced
## Warning in log(inter_eye_dist_change): NaNs produced
## Warning in log(inter_eye_dist_change): NaNs produced
## Warning in log(inter_eye_dist_change): NaNs produced
## Warning in log(inter_eye_dist_change): NaNs produced
## Warning in log(inter_eye_dist_change): NaNs produced
## Warning in log(inter_eye_dist_change): NaNs produced
## Warning in log(inter_eye_dist_change): NaNs produced
```

```
## Warning in finalizeTMB(TMBStruc, obj, fit, h, data.tmb.old): Model convergence
## problem; non-positive-definite Hessian matrix. See vignette('troubleshooting')
```

```
## Warning in finalizeTMB(TMBStruc, obj, fit, h, data.tmb.old): Model convergence
## problem; false convergence (8). See vignette('troubleshooting'),
## help('diagnose')
```

```
## Warning in log(inter_eye_dist_change): NaNs produced
## Warning in log(inter_eye_dist_change): NaNs produced
```

```
## Warning in finalizeTMB(TMBStruc, obj, fit, h, data.tmb.old): Model convergence
## problem; non-positive-definite Hessian matrix. See vignette('troubleshooting')
```

```
## Warning in log(inter_eye_dist_change): NaNs produced
## Warning in log(inter_eye_dist_change): NaNs produced
```

```
## Warning in finalizeTMB(TMBStruc, obj, fit, h, data.tmb.old): Model convergence
## problem; non-positive-definite Hessian matrix. See vignette('troubleshooting')
```

```
## Warning in finalizeTMB(TMBStruc, obj, fit, h, data.tmb.old): Model convergence
## problem; false convergence (8). See vignette('troubleshooting'),
## help('diagnose')
```

```
## Warning in log(inter_eye_dist_change): NaNs produced
## Warning in log(inter_eye_dist_change): NaNs produced
## Warning in log(inter_eye_dist_change): NaNs produced
## Warning in log(inter_eye_dist_change): NaNs produced
```

```
## Warning in finalizeTMB(TMBStruc, obj, fit, h, data.tmb.old): Model convergence
## problem; non-positive-definite Hessian matrix. See vignette('troubleshooting')
```

```
## Warning in log(inter_eye_dist_change): NaNs produced
## Warning in log(inter_eye_dist_change): NaNs produced
## Warning in log(inter_eye_dist_change): NaNs produced
## Warning in log(inter_eye_dist_change): NaNs produced
```

```
## Warning in finalizeTMB(TMBStruc, obj, fit, h, data.tmb.old): Model convergence
## problem; non-positive-definite Hessian matrix. See vignette('troubleshooting')
```

```
## Warning in finalizeTMB(TMBStruc, obj, fit, h, data.tmb.old): Model convergence
## problem; false convergence (8). See vignette('troubleshooting'),
## help('diagnose')
```

```
## Warning in log(inter_eye_dist_change): NaNs produced
## Warning in log(inter_eye_dist_change): NaNs produced
```

```
## Warning in finalizeTMB(TMBStruc, obj, fit, h, data.tmb.old): Model convergence
## problem; non-positive-definite Hessian matrix. See vignette('troubleshooting')
```

```
## Warning in log(inter_eye_dist_change): NaNs produced
## Warning in log(inter_eye_dist_change): NaNs produced
```

```
## Warning in finalizeTMB(TMBStruc, obj, fit, h, data.tmb.old): Model convergence
## problem; non-positive-definite Hessian matrix. See vignette('troubleshooting')
```

```
## Warning in log(inter_eye_dist_change): NaNs produced
## Warning in log(inter_eye_dist_change): NaNs produced
## Warning in log(inter_eye_dist_change): NaNs produced
## Warning in log(inter_eye_dist_change): NaNs produced
```

```
## Warning in finalizeTMB(TMBStruc, obj, fit, h, data.tmb.old): Model convergence
## problem; non-positive-definite Hessian matrix. See vignette('troubleshooting')
```

```
## Warning in finalizeTMB(TMBStruc, obj, fit, h, data.tmb.old): Model convergence
## problem; false convergence (8). See vignette('troubleshooting'),
## help('diagnose')
```

```
## Warning in log(inter_eye_dist_change): NaNs produced
## Warning in log(inter_eye_dist_change): NaNs produced
## Warning in log(inter_eye_dist_change): NaNs produced
## Warning in log(inter_eye_dist_change): NaNs produced
```

```
## Warning in finalizeTMB(TMBStruc, obj, fit, h, data.tmb.old): Model convergence
## problem; non-positive-definite Hessian matrix. See vignette('troubleshooting')
```

```
## Warning in finalizeTMB(TMBStruc, obj, fit, h, data.tmb.old): Model convergence
## problem; false convergence (8). See vignette('troubleshooting'),
## help('diagnose')
```

```
## Warning in log(inter_eye_dist_change): NaNs produced
## Warning in log(inter_eye_dist_change): NaNs produced
```

```
## Warning in finalizeTMB(TMBStruc, obj, fit, h, data.tmb.old): Model convergence
## problem; non-positive-definite Hessian matrix. See vignette('troubleshooting')
```

```
## Warning in finalizeTMB(TMBStruc, obj, fit, h, data.tmb.old): Model convergence
## problem; false convergence (8). See vignette('troubleshooting'),
## help('diagnose')
```

```
## Warning in log(inter_eye_dist_change): NaNs produced
## Warning in log(inter_eye_dist_change): NaNs produced
```

```
## Warning in finalizeTMB(TMBStruc, obj, fit, h, data.tmb.old): Model convergence
## problem; non-positive-definite Hessian matrix. See vignette('troubleshooting')
```

```
## Warning in finalizeTMB(TMBStruc, obj, fit, h, data.tmb.old): Model convergence
## problem; singular convergence (7). See vignette('troubleshooting'),
## help('diagnose')
```

```
## Warning in log(inter_eye_dist_change): NaNs produced
## Warning in log(inter_eye_dist_change): NaNs produced
```

```
## Warning in finalizeTMB(TMBStruc, obj, fit, h, data.tmb.old): Model convergence
## problem; non-positive-definite Hessian matrix. See vignette('troubleshooting')
```

```
## Warning in finalizeTMB(TMBStruc, obj, fit, h, data.tmb.old): Model convergence
## problem; singular convergence (7). See vignette('troubleshooting'),
## help('diagnose')
```

```
## Warning in log(inter_eye_dist_change): NaNs produced
## Warning in log(inter_eye_dist_change): NaNs produced
## Warning in log(inter_eye_dist_change): NaNs produced
## Warning in log(inter_eye_dist_change): NaNs produced
```

```
## Warning in finalizeTMB(TMBStruc, obj, fit, h, data.tmb.old): Model convergence
## problem; non-positive-definite Hessian matrix. See vignette('troubleshooting')
```

```
## Warning in finalizeTMB(TMBStruc, obj, fit, h, data.tmb.old): Model convergence
## problem; singular convergence (7). See vignette('troubleshooting'),
## help('diagnose')
```

```
## Warning in log(inter_eye_dist_change): NaNs produced
## Warning in log(inter_eye_dist_change): NaNs produced
```

```
## Warning in finalizeTMB(TMBStruc, obj, fit, h, data.tmb.old): Model convergence
## problem; non-positive-definite Hessian matrix. See vignette('troubleshooting')
```

```
## Warning in finalizeTMB(TMBStruc, obj, fit, h, data.tmb.old): Model convergence
## problem; singular convergence (7). See vignette('troubleshooting'),
## help('diagnose')
```

```
## Warning in log(inter_eye_dist_change): NaNs produced
## Warning in log(inter_eye_dist_change): NaNs produced
```

```
## Warning in finalizeTMB(TMBStruc, obj, fit, h, data.tmb.old): Model convergence
## problem; non-positive-definite Hessian matrix. See vignette('troubleshooting')
```

```
## Warning in log(inter_eye_dist_change): NaNs produced
## Warning in log(inter_eye_dist_change): NaNs produced
```

```
## Warning in finalizeTMB(TMBStruc, obj, fit, h, data.tmb.old): Model convergence
## problem; non-positive-definite Hessian matrix. See vignette('troubleshooting')
```

```
## Warning in log(inter_eye_dist_change): NaNs produced
## Warning in log(inter_eye_dist_change): NaNs produced
## Warning in log(inter_eye_dist_change): NaNs produced
## Warning in log(inter_eye_dist_change): NaNs produced
```

```
## Warning in finalizeTMB(TMBStruc, obj, fit, h, data.tmb.old): Model convergence
## problem; non-positive-definite Hessian matrix. See vignette('troubleshooting')
```

```
## Warning in log(inter_eye_dist_change): NaNs produced
## Warning in log(inter_eye_dist_change): NaNs produced
```

```
## Warning in finalizeTMB(TMBStruc, obj, fit, h, data.tmb.old): Model convergence
## problem; non-positive-definite Hessian matrix. See vignette('troubleshooting')
```

```
## Warning in finalizeTMB(TMBStruc, obj, fit, h, data.tmb.old): Model convergence
## problem; false convergence (8). See vignette('troubleshooting'),
## help('diagnose')
```

```
## Warning in log(inter_eye_dist_change): NaNs produced
## Warning in log(inter_eye_dist_change): NaNs produced
## Warning in log(inter_eye_dist_change): NaNs produced
## Warning in log(inter_eye_dist_change): NaNs produced
```

```
## Warning in finalizeTMB(TMBStruc, obj, fit, h, data.tmb.old): Model convergence
## problem; non-positive-definite Hessian matrix. See vignette('troubleshooting')
```

```
## Warning in finalizeTMB(TMBStruc, obj, fit, h, data.tmb.old): Model convergence
## problem; singular convergence (7). See vignette('troubleshooting'),
## help('diagnose')
```

```
## Warning in log(inter_eye_dist_change): NaNs produced
## Warning in log(inter_eye_dist_change): NaNs produced
## Warning in log(inter_eye_dist_change): NaNs produced
## Warning in log(inter_eye_dist_change): NaNs produced
```

```
## Warning in finalizeTMB(TMBStruc, obj, fit, h, data.tmb.old): Model convergence
## problem; non-positive-definite Hessian matrix. See vignette('troubleshooting')
```

```
## Warning in finalizeTMB(TMBStruc, obj, fit, h, data.tmb.old): Model convergence
## problem; singular convergence (7). See vignette('troubleshooting'),
## help('diagnose')
```

```
## Warning in log(inter_eye_dist_change): NaNs produced
## Warning in log(inter_eye_dist_change): NaNs produced
```

```
## Warning in finalizeTMB(TMBStruc, obj, fit, h, data.tmb.old): Model convergence
## problem; non-positive-definite Hessian matrix. See vignette('troubleshooting')
```

```
## Warning in log(inter_eye_dist_change): NaNs produced
## Warning in log(inter_eye_dist_change): NaNs produced
## Warning in log(inter_eye_dist_change): NaNs produced
## Warning in log(inter_eye_dist_change): NaNs produced
## Warning in log(inter_eye_dist_change): NaNs produced
## Warning in log(inter_eye_dist_change): NaNs produced
```

```
## Warning in finalizeTMB(TMBStruc, obj, fit, h, data.tmb.old): Model convergence
## problem; non-positive-definite Hessian matrix. See vignette('troubleshooting')
```

```
## Warning in log(inter_eye_dist_change): NaNs produced
## Warning in log(inter_eye_dist_change): NaNs produced
```

```
## Warning in finalizeTMB(TMBStruc, obj, fit, h, data.tmb.old): Model convergence
## problem; non-positive-definite Hessian matrix. See vignette('troubleshooting')
```

```
## Warning in finalizeTMB(TMBStruc, obj, fit, h, data.tmb.old): Model convergence
## problem; false convergence (8). See vignette('troubleshooting'),
## help('diagnose')
```

```
## Warning in log(inter_eye_dist_change): NaNs produced
## Warning in log(inter_eye_dist_change): NaNs produced
```

```
## Warning in finalizeTMB(TMBStruc, obj, fit, h, data.tmb.old): Model convergence
## problem; non-positive-definite Hessian matrix. See vignette('troubleshooting')
```

```
## Warning in log(inter_eye_dist_change): NaNs produced
## Warning in log(inter_eye_dist_change): NaNs produced
```

```
## Warning in finalizeTMB(TMBStruc, obj, fit, h, data.tmb.old): Model convergence
## problem; non-positive-definite Hessian matrix. See vignette('troubleshooting')
```

```
## Warning in log(inter_eye_dist_change): NaNs produced
## Warning in log(inter_eye_dist_change): NaNs produced
## Warning in log(inter_eye_dist_change): NaNs produced
## Warning in log(inter_eye_dist_change): NaNs produced
```

```
## Warning in finalizeTMB(TMBStruc, obj, fit, h, data.tmb.old): Model convergence
## problem; non-positive-definite Hessian matrix. See vignette('troubleshooting')
```

```
## Warning in finalizeTMB(TMBStruc, obj, fit, h, data.tmb.old): Model convergence
## problem; singular convergence (7). See vignette('troubleshooting'),
## help('diagnose')
```

```
## Warning in log(inter_eye_dist_change): NaNs produced
## Warning in log(inter_eye_dist_change): NaNs produced
```

```
## Warning in finalizeTMB(TMBStruc, obj, fit, h, data.tmb.old): Model convergence
## problem; non-positive-definite Hessian matrix. See vignette('troubleshooting')
```

```
## Warning in log(inter_eye_dist_change): NaNs produced
## Warning in log(inter_eye_dist_change): NaNs produced
```

```
## Warning in finalizeTMB(TMBStruc, obj, fit, h, data.tmb.old): Model convergence
## problem; non-positive-definite Hessian matrix. See vignette('troubleshooting')
```

```
## Warning in finalizeTMB(TMBStruc, obj, fit, h, data.tmb.old): Model convergence
## problem; singular convergence (7). See vignette('troubleshooting'),
## help('diagnose')
```

```
## Warning in log(inter_eye_dist_change): NaNs produced
## Warning in log(inter_eye_dist_change): NaNs produced
```

```
## Warning in finalizeTMB(TMBStruc, obj, fit, h, data.tmb.old): Model convergence
## problem; non-positive-definite Hessian matrix. See vignette('troubleshooting')
```

```
## Warning in log(inter_eye_dist_change): NaNs produced
## Warning in log(inter_eye_dist_change): NaNs produced
```

```
## Warning in finalizeTMB(TMBStruc, obj, fit, h, data.tmb.old): Model convergence
## problem; non-positive-definite Hessian matrix. See vignette('troubleshooting')
```

```
## Warning in log(inter_eye_dist_change): NaNs produced
## Warning in log(inter_eye_dist_change): NaNs produced
## Warning in log(inter_eye_dist_change): NaNs produced
## Warning in log(inter_eye_dist_change): NaNs produced
```

```
## Warning in finalizeTMB(TMBStruc, obj, fit, h, data.tmb.old): Model convergence
## problem; non-positive-definite Hessian matrix. See vignette('troubleshooting')
```

```
## Warning in log(inter_eye_dist_change): NaNs produced
## Warning in log(inter_eye_dist_change): NaNs produced
```

```
## Warning in finalizeTMB(TMBStruc, obj, fit, h, data.tmb.old): Model convergence
## problem; non-positive-definite Hessian matrix. See vignette('troubleshooting')
```

```
## Warning in finalizeTMB(TMBStruc, obj, fit, h, data.tmb.old): Model convergence
## problem; false convergence (8). See vignette('troubleshooting'),
## help('diagnose')
```

```
## Warning in log(inter_eye_dist_change): NaNs produced
## Warning in log(inter_eye_dist_change): NaNs produced
```

```
## Warning in finalizeTMB(TMBStruc, obj, fit, h, data.tmb.old): Model convergence
## problem; non-positive-definite Hessian matrix. See vignette('troubleshooting')
```

```
## Warning in log(inter_eye_dist_change): NaNs produced
## Warning in log(inter_eye_dist_change): NaNs produced
## Warning in log(inter_eye_dist_change): NaNs produced
## Warning in log(inter_eye_dist_change): NaNs produced
```

```
## Warning in finalizeTMB(TMBStruc, obj, fit, h, data.tmb.old): Model convergence
## problem; non-positive-definite Hessian matrix. See vignette('troubleshooting')
```

```
## Warning in finalizeTMB(TMBStruc, obj, fit, h, data.tmb.old): Model convergence
## problem; singular convergence (7). See vignette('troubleshooting'),
## help('diagnose')
```

```
## Warning in log(inter_eye_dist_change): NaNs produced
## Warning in log(inter_eye_dist_change): NaNs produced
## Warning in log(inter_eye_dist_change): NaNs produced
## Warning in log(inter_eye_dist_change): NaNs produced
```

```
## Warning in finalizeTMB(TMBStruc, obj, fit, h, data.tmb.old): Model convergence
## problem; non-positive-definite Hessian matrix. See vignette('troubleshooting')
```

```
## Warning in log(inter_eye_dist_change): NaNs produced
## Warning in log(inter_eye_dist_change): NaNs produced
```

```
## Warning in finalizeTMB(TMBStruc, obj, fit, h, data.tmb.old): Model convergence
## problem; non-positive-definite Hessian matrix. See vignette('troubleshooting')
```

```
## Warning in finalizeTMB(TMBStruc, obj, fit, h, data.tmb.old): Model convergence
## problem; false convergence (8). See vignette('troubleshooting'),
## help('diagnose')
```

```
## Warning in log(inter_eye_dist_change): NaNs produced
## Warning in log(inter_eye_dist_change): NaNs produced
## Warning in log(inter_eye_dist_change): NaNs produced
## Warning in log(inter_eye_dist_change): NaNs produced
```

```
## Warning in finalizeTMB(TMBStruc, obj, fit, h, data.tmb.old): Model convergence
## problem; non-positive-definite Hessian matrix. See vignette('troubleshooting')
```

```
## Warning in finalizeTMB(TMBStruc, obj, fit, h, data.tmb.old): Model convergence
## problem; false convergence (8). See vignette('troubleshooting'),
## help('diagnose')
```

```
## Warning in log(inter_eye_dist_change): NaNs produced
## Warning in log(inter_eye_dist_change): NaNs produced
```

```
## Warning in finalizeTMB(TMBStruc, obj, fit, h, data.tmb.old): Model convergence
## problem; non-positive-definite Hessian matrix. See vignette('troubleshooting')
```

```
## Warning in finalizeTMB(TMBStruc, obj, fit, h, data.tmb.old): Model convergence
## problem; singular convergence (7). See vignette('troubleshooting'),
## help('diagnose')
```

```
## Warning in log(inter_eye_dist_change): NaNs produced
## Warning in log(inter_eye_dist_change): NaNs produced
```

```
## Warning in finalizeTMB(TMBStruc, obj, fit, h, data.tmb.old): Model convergence
## problem; non-positive-definite Hessian matrix. See vignette('troubleshooting')
```

```
## Warning in finalizeTMB(TMBStruc, obj, fit, h, data.tmb.old): Model convergence
## problem; singular convergence (7). See vignette('troubleshooting'),
## help('diagnose')
```

```
## Warning in log(inter_eye_dist_change): NaNs produced
## Warning in log(inter_eye_dist_change): NaNs produced
```

```
## Warning in finalizeTMB(TMBStruc, obj, fit, h, data.tmb.old): Model convergence
## problem; non-positive-definite Hessian matrix. See vignette('troubleshooting')
```

```
## Warning in finalizeTMB(TMBStruc, obj, fit, h, data.tmb.old): Model convergence
## problem; singular convergence (7). See vignette('troubleshooting'),
## help('diagnose')
```

```
## Warning in log(inter_eye_dist_change): NaNs produced
## Warning in log(inter_eye_dist_change): NaNs produced
```

```
## Warning in finalizeTMB(TMBStruc, obj, fit, h, data.tmb.old): Model convergence
## problem; non-positive-definite Hessian matrix. See vignette('troubleshooting')
```

```
## Warning in finalizeTMB(TMBStruc, obj, fit, h, data.tmb.old): Model convergence
## problem; singular convergence (7). See vignette('troubleshooting'),
## help('diagnose')
```

```
## Warning in log(inter_eye_dist_change): NaNs produced
## Warning in log(inter_eye_dist_change): NaNs produced
```

```
## Warning in finalizeTMB(TMBStruc, obj, fit, h, data.tmb.old): Model convergence
## problem; non-positive-definite Hessian matrix. See vignette('troubleshooting')
```

```
## Warning in finalizeTMB(TMBStruc, obj, fit, h, data.tmb.old): Model convergence
## problem; singular convergence (7). See vignette('troubleshooting'),
## help('diagnose')
```

```
## Warning in log(inter_eye_dist_change): NaNs produced
## Warning in log(inter_eye_dist_change): NaNs produced
## Warning in log(inter_eye_dist_change): NaNs produced
## Warning in log(inter_eye_dist_change): NaNs produced
```

```
## Warning in finalizeTMB(TMBStruc, obj, fit, h, data.tmb.old): Model convergence
## problem; non-positive-definite Hessian matrix. See vignette('troubleshooting')
```

```
## Warning in finalizeTMB(TMBStruc, obj, fit, h, data.tmb.old): Model convergence
## problem; singular convergence (7). See vignette('troubleshooting'),
## help('diagnose')
```

```
## Warning in log(inter_eye_dist_change): NaNs produced
## Warning in log(inter_eye_dist_change): NaNs produced
```

```
## Warning in finalizeTMB(TMBStruc, obj, fit, h, data.tmb.old): Model convergence
## problem; non-positive-definite Hessian matrix. See vignette('troubleshooting')
```

```
## Warning in finalizeTMB(TMBStruc, obj, fit, h, data.tmb.old): Model convergence
## problem; singular convergence (7). See vignette('troubleshooting'),
## help('diagnose')
```

```
## Warning in log(inter_eye_dist_change): NaNs produced
## Warning in log(inter_eye_dist_change): NaNs produced
```

```
## Warning in finalizeTMB(TMBStruc, obj, fit, h, data.tmb.old): Model convergence
## problem; non-positive-definite Hessian matrix. See vignette('troubleshooting')
```

```
## Warning in finalizeTMB(TMBStruc, obj, fit, h, data.tmb.old): Model convergence
## problem; singular convergence (7). See vignette('troubleshooting'),
## help('diagnose')
```

```
## Warning in log(inter_eye_dist_change): NaNs produced
## Warning in log(inter_eye_dist_change): NaNs produced
```

```
## Warning in finalizeTMB(TMBStruc, obj, fit, h, data.tmb.old): Model convergence
## problem; non-positive-definite Hessian matrix. See vignette('troubleshooting')
```

```
## Warning in finalizeTMB(TMBStruc, obj, fit, h, data.tmb.old): Model convergence
## problem; singular convergence (7). See vignette('troubleshooting'),
## help('diagnose')
```

```
## Warning in log(inter_eye_dist_change): NaNs produced
## Warning in log(inter_eye_dist_change): NaNs produced
```

```
## Warning in finalizeTMB(TMBStruc, obj, fit, h, data.tmb.old): Model convergence
## problem; non-positive-definite Hessian matrix. See vignette('troubleshooting')
```

```
## Warning in finalizeTMB(TMBStruc, obj, fit, h, data.tmb.old): Model convergence
## problem; singular convergence (7). See vignette('troubleshooting'),
## help('diagnose')
```

```
## Warning in log(inter_eye_dist_change): NaNs produced
## Warning in log(inter_eye_dist_change): NaNs produced
```

```
## Warning in finalizeTMB(TMBStruc, obj, fit, h, data.tmb.old): Model convergence
## problem; non-positive-definite Hessian matrix. See vignette('troubleshooting')
```

```
## Warning in finalizeTMB(TMBStruc, obj, fit, h, data.tmb.old): Model convergence
## problem; singular convergence (7). See vignette('troubleshooting'),
## help('diagnose')
```

```
## Warning in log(inter_eye_dist_change): NaNs produced
## Warning in log(inter_eye_dist_change): NaNs produced
```

```
## Warning in finalizeTMB(TMBStruc, obj, fit, h, data.tmb.old): Model convergence
## problem; non-positive-definite Hessian matrix. See vignette('troubleshooting')
```

```
## Warning in finalizeTMB(TMBStruc, obj, fit, h, data.tmb.old): Model convergence
## problem; false convergence (8). See vignette('troubleshooting'),
## help('diagnose')
```

```
## Warning in log(inter_eye_dist_change): NaNs produced
## Warning in log(inter_eye_dist_change): NaNs produced
```

```
## Warning in finalizeTMB(TMBStruc, obj, fit, h, data.tmb.old): Model convergence
## problem; non-positive-definite Hessian matrix. See vignette('troubleshooting')
```

```
## Warning in finalizeTMB(TMBStruc, obj, fit, h, data.tmb.old): Model convergence
## problem; singular convergence (7). See vignette('troubleshooting'),
## help('diagnose')
```

```
## Warning in log(inter_eye_dist_change): NaNs produced
## Warning in log(inter_eye_dist_change): NaNs produced
## Warning in log(inter_eye_dist_change): NaNs produced
## Warning in log(inter_eye_dist_change): NaNs produced
```

```
## Warning in finalizeTMB(TMBStruc, obj, fit, h, data.tmb.old): Model convergence
## problem; non-positive-definite Hessian matrix. See vignette('troubleshooting')
```

```
## Warning in finalizeTMB(TMBStruc, obj, fit, h, data.tmb.old): Model convergence
## problem; false convergence (8). See vignette('troubleshooting'),
## help('diagnose')
```

```
## Warning in log(inter_eye_dist_change): NaNs produced
## Warning in log(inter_eye_dist_change): NaNs produced
```

```
## Warning in finalizeTMB(TMBStruc, obj, fit, h, data.tmb.old): Model convergence
## problem; non-positive-definite Hessian matrix. See vignette('troubleshooting')
```

```
## Warning in finalizeTMB(TMBStruc, obj, fit, h, data.tmb.old): Model convergence
## problem; singular convergence (7). See vignette('troubleshooting'),
## help('diagnose')
```

```
## Warning in log(inter_eye_dist_change): NaNs produced
## Warning in log(inter_eye_dist_change): NaNs produced
## Warning in log(inter_eye_dist_change): NaNs produced
## Warning in log(inter_eye_dist_change): NaNs produced
```

```
## Warning in finalizeTMB(TMBStruc, obj, fit, h, data.tmb.old): Model convergence
## problem; singular convergence (7). See vignette('troubleshooting'),
## help('diagnose')
```

```
## Warning in log(inter_eye_dist_change): NaNs produced
## Warning in log(inter_eye_dist_change): NaNs produced
## Warning in log(inter_eye_dist_change): NaNs produced
## Warning in log(inter_eye_dist_change): NaNs produced
## Warning in log(inter_eye_dist_change): NaNs produced
## Warning in log(inter_eye_dist_change): NaNs produced
## Warning in log(inter_eye_dist_change): NaNs produced
## Warning in log(inter_eye_dist_change): NaNs produced
## Warning in log(inter_eye_dist_change): NaNs produced
## Warning in log(inter_eye_dist_change): NaNs produced
## Warning in log(inter_eye_dist_change): NaNs produced
## Warning in log(inter_eye_dist_change): NaNs produced
## Warning in log(inter_eye_dist_change): NaNs produced
## Warning in log(inter_eye_dist_change): NaNs produced
```

```
## Warning in finalizeTMB(TMBStruc, obj, fit, h, data.tmb.old): Model convergence
## problem; non-positive-definite Hessian matrix. See vignette('troubleshooting')
```

```
## Warning in finalizeTMB(TMBStruc, obj, fit, h, data.tmb.old): Model convergence
## problem; singular convergence (7). See vignette('troubleshooting'),
## help('diagnose')
```

```
## Warning in log(inter_eye_dist_change): NaNs produced
## Warning in log(inter_eye_dist_change): NaNs produced
```

```
## Warning in finalizeTMB(TMBStruc, obj, fit, h, data.tmb.old): Model convergence
## problem; non-positive-definite Hessian matrix. See vignette('troubleshooting')
```

```
## Warning in finalizeTMB(TMBStruc, obj, fit, h, data.tmb.old): Model convergence
## problem; singular convergence (7). See vignette('troubleshooting'),
## help('diagnose')
```

```
## Warning in log(inter_eye_dist_change): NaNs produced
## Warning in log(inter_eye_dist_change): NaNs produced
## Warning in log(inter_eye_dist_change): NaNs produced
## Warning in log(inter_eye_dist_change): NaNs produced
## Warning in log(inter_eye_dist_change): NaNs produced
## Warning in log(inter_eye_dist_change): NaNs produced
## Warning in log(inter_eye_dist_change): NaNs produced
## Warning in log(inter_eye_dist_change): NaNs produced
## Warning in log(inter_eye_dist_change): NaNs produced
## Warning in log(inter_eye_dist_change): NaNs produced
```

```
## Warning in finalizeTMB(TMBStruc, obj, fit, h, data.tmb.old): Model convergence
## problem; non-positive-definite Hessian matrix. See vignette('troubleshooting')
```

```
## Warning in finalizeTMB(TMBStruc, obj, fit, h, data.tmb.old): Model convergence
## problem; singular convergence (7). See vignette('troubleshooting'),
## help('diagnose')
```

```
## Warning in log(inter_eye_dist_change): NaNs produced
## Warning in log(inter_eye_dist_change): NaNs produced
```

```
## Warning in finalizeTMB(TMBStruc, obj, fit, h, data.tmb.old): Model convergence
## problem; non-positive-definite Hessian matrix. See vignette('troubleshooting')
```

```
## Warning in finalizeTMB(TMBStruc, obj, fit, h, data.tmb.old): Model convergence
## problem; singular convergence (7). See vignette('troubleshooting'),
## help('diagnose')
```

```
## Warning in log(inter_eye_dist_change): NaNs produced
## Warning in log(inter_eye_dist_change): NaNs produced
```

```
## Warning in finalizeTMB(TMBStruc, obj, fit, h, data.tmb.old): Model convergence
## problem; non-positive-definite Hessian matrix. See vignette('troubleshooting')
```

```
## Warning in finalizeTMB(TMBStruc, obj, fit, h, data.tmb.old): Model convergence
## problem; singular convergence (7). See vignette('troubleshooting'),
## help('diagnose')
```

```
## Warning in log(inter_eye_dist_change): NaNs produced
## Warning in log(inter_eye_dist_change): NaNs produced
```

```
## Warning in finalizeTMB(TMBStruc, obj, fit, h, data.tmb.old): Model convergence
## problem; non-positive-definite Hessian matrix. See vignette('troubleshooting')
```

```
## Warning in finalizeTMB(TMBStruc, obj, fit, h, data.tmb.old): Model convergence
## problem; singular convergence (7). See vignette('troubleshooting'),
## help('diagnose')
```

```
## Warning in log(inter_eye_dist_change): NaNs produced
## Warning in log(inter_eye_dist_change): NaNs produced
```

```
## Warning in finalizeTMB(TMBStruc, obj, fit, h, data.tmb.old): Model convergence
## problem; non-positive-definite Hessian matrix. See vignette('troubleshooting')
```

```
## Warning in finalizeTMB(TMBStruc, obj, fit, h, data.tmb.old): Model convergence
## problem; singular convergence (7). See vignette('troubleshooting'),
## help('diagnose')
```

```
## Warning in log(inter_eye_dist_change): NaNs produced
## Warning in log(inter_eye_dist_change): NaNs produced
```

```
## Warning in finalizeTMB(TMBStruc, obj, fit, h, data.tmb.old): Model convergence
## problem; non-positive-definite Hessian matrix. See vignette('troubleshooting')
```

```
## Warning in finalizeTMB(TMBStruc, obj, fit, h, data.tmb.old): Model convergence
## problem; singular convergence (7). See vignette('troubleshooting'),
## help('diagnose')
```

```
## Warning in log(inter_eye_dist_change): NaNs produced
## Warning in log(inter_eye_dist_change): NaNs produced
## Warning in log(inter_eye_dist_change): NaNs produced
## Warning in log(inter_eye_dist_change): NaNs produced
## Warning in log(inter_eye_dist_change): NaNs produced
## Warning in log(inter_eye_dist_change): NaNs produced
## Warning in log(inter_eye_dist_change): NaNs produced
## Warning in log(inter_eye_dist_change): NaNs produced
## Warning in log(inter_eye_dist_change): NaNs produced
## Warning in log(inter_eye_dist_change): NaNs produced
## Warning in log(inter_eye_dist_change): NaNs produced
## Warning in log(inter_eye_dist_change): NaNs produced
## Warning in log(inter_eye_dist_change): NaNs produced
## Warning in log(inter_eye_dist_change): NaNs produced
## Warning in log(inter_eye_dist_change): NaNs produced
## Warning in log(inter_eye_dist_change): NaNs produced
## Warning in log(inter_eye_dist_change): NaNs produced
## Warning in log(inter_eye_dist_change): NaNs produced
## Warning in log(inter_eye_dist_change): NaNs produced
## Warning in log(inter_eye_dist_change): NaNs produced
## Warning in log(inter_eye_dist_change): NaNs produced
## Warning in log(inter_eye_dist_change): NaNs produced
## Warning in log(inter_eye_dist_change): NaNs produced
## Warning in log(inter_eye_dist_change): NaNs produced
## Warning in log(inter_eye_dist_change): NaNs produced
## Warning in log(inter_eye_dist_change): NaNs produced
## Warning in log(inter_eye_dist_change): NaNs produced
## Warning in log(inter_eye_dist_change): NaNs produced
## Warning in log(inter_eye_dist_change): NaNs produced
## Warning in log(inter_eye_dist_change): NaNs produced
## Warning in log(inter_eye_dist_change): NaNs produced
## Warning in log(inter_eye_dist_change): NaNs produced
## Warning in log(inter_eye_dist_change): NaNs produced
## Warning in log(inter_eye_dist_change): NaNs produced
## Warning in log(inter_eye_dist_change): NaNs produced
## Warning in log(inter_eye_dist_change): NaNs produced
## Warning in log(inter_eye_dist_change): NaNs produced
## Warning in log(inter_eye_dist_change): NaNs produced
## Warning in log(inter_eye_dist_change): NaNs produced
## Warning in log(inter_eye_dist_change): NaNs produced
## Warning in log(inter_eye_dist_change): NaNs produced
## Warning in log(inter_eye_dist_change): NaNs produced
## Warning in log(inter_eye_dist_change): NaNs produced
## Warning in log(inter_eye_dist_change): NaNs produced
## Warning in log(inter_eye_dist_change): NaNs produced
## Warning in log(inter_eye_dist_change): NaNs produced
## Warning in log(inter_eye_dist_change): NaNs produced
## Warning in log(inter_eye_dist_change): NaNs produced
## Warning in log(inter_eye_dist_change): NaNs produced
## Warning in log(inter_eye_dist_change): NaNs produced
## Warning in log(inter_eye_dist_change): NaNs produced
## Warning in log(inter_eye_dist_change): NaNs produced
## Warning in log(inter_eye_dist_change): NaNs produced
## Warning in log(inter_eye_dist_change): NaNs produced
## Warning in log(inter_eye_dist_change): NaNs produced
## Warning in log(inter_eye_dist_change): NaNs produced
## Warning in log(inter_eye_dist_change): NaNs produced
## Warning in log(inter_eye_dist_change): NaNs produced
## Warning in log(inter_eye_dist_change): NaNs produced
## Warning in log(inter_eye_dist_change): NaNs produced
```

```
## Warning in finalizeTMB(TMBStruc, obj, fit, h, data.tmb.old): Model convergence
## problem; singular convergence (7). See vignette('troubleshooting'),
## help('diagnose')
```

```
## Warning in log(inter_eye_dist_change): NaNs produced
## Warning in log(inter_eye_dist_change): NaNs produced
```

```
## Warning in finalizeTMB(TMBStruc, obj, fit, h, data.tmb.old): Model convergence
## problem; non-positive-definite Hessian matrix. See vignette('troubleshooting')
```

```
## Warning in finalizeTMB(TMBStruc, obj, fit, h, data.tmb.old): Model convergence
## problem; singular convergence (7). See vignette('troubleshooting'),
## help('diagnose')
```

```
## Warning in log(inter_eye_dist_change): NaNs produced
## Warning in log(inter_eye_dist_change): NaNs produced
## Warning in log(inter_eye_dist_change): NaNs produced
## Warning in log(inter_eye_dist_change): NaNs produced
```

```
## Warning in finalizeTMB(TMBStruc, obj, fit, h, data.tmb.old): Model convergence
## problem; non-positive-definite Hessian matrix. See vignette('troubleshooting')
```

```
## Warning in finalizeTMB(TMBStruc, obj, fit, h, data.tmb.old): Model convergence
## problem; singular convergence (7). See vignette('troubleshooting'),
## help('diagnose')
```

```
## Warning in log(inter_eye_dist_change): NaNs produced
## Warning in log(inter_eye_dist_change): NaNs produced
```

```
## Warning in finalizeTMB(TMBStruc, obj, fit, h, data.tmb.old): Model convergence
## problem; non-positive-definite Hessian matrix. See vignette('troubleshooting')
```

```
## Warning in finalizeTMB(TMBStruc, obj, fit, h, data.tmb.old): Model convergence
## problem; singular convergence (7). See vignette('troubleshooting'),
## help('diagnose')
```

```
## Warning in log(inter_eye_dist_change): NaNs produced
## Warning in log(inter_eye_dist_change): NaNs produced
```

```
## Warning in finalizeTMB(TMBStruc, obj, fit, h, data.tmb.old): Model convergence
## problem; non-positive-definite Hessian matrix. See vignette('troubleshooting')
```

```
## Warning in finalizeTMB(TMBStruc, obj, fit, h, data.tmb.old): Model convergence
## problem; singular convergence (7). See vignette('troubleshooting'),
## help('diagnose')
```

```
## Warning in log(inter_eye_dist_change): NaNs produced
## Warning in log(inter_eye_dist_change): NaNs produced
```

```
## Warning in finalizeTMB(TMBStruc, obj, fit, h, data.tmb.old): Model convergence
## problem; non-positive-definite Hessian matrix. See vignette('troubleshooting')
```

```
## Warning in log(inter_eye_dist_change): NaNs produced
## Warning in log(inter_eye_dist_change): NaNs produced
## Warning in log(inter_eye_dist_change): NaNs produced
## Warning in log(inter_eye_dist_change): NaNs produced
```

```
## Warning in finalizeTMB(TMBStruc, obj, fit, h, data.tmb.old): Model convergence
## problem; non-positive-definite Hessian matrix. See vignette('troubleshooting')
```

```
## Warning in finalizeTMB(TMBStruc, obj, fit, h, data.tmb.old): Model convergence
## problem; singular convergence (7). See vignette('troubleshooting'),
## help('diagnose')
```

```
## Warning in log(inter_eye_dist_change): NaNs produced
## Warning in log(inter_eye_dist_change): NaNs produced
## Warning in log(inter_eye_dist_change): NaNs produced
## Warning in log(inter_eye_dist_change): NaNs produced
## Warning in log(inter_eye_dist_change): NaNs produced
## Warning in log(inter_eye_dist_change): NaNs produced
## Warning in log(inter_eye_dist_change): NaNs produced
## Warning in log(inter_eye_dist_change): NaNs produced
```

```
## Warning in finalizeTMB(TMBStruc, obj, fit, h, data.tmb.old): Model convergence
## problem; singular convergence (7). See vignette('troubleshooting'),
## help('diagnose')
```

```
## Warning in log(inter_eye_dist_change): NaNs produced
## Warning in log(inter_eye_dist_change): NaNs produced
```

```
## Warning in finalizeTMB(TMBStruc, obj, fit, h, data.tmb.old): Model convergence
## problem; non-positive-definite Hessian matrix. See vignette('troubleshooting')
```

```
## Warning in finalizeTMB(TMBStruc, obj, fit, h, data.tmb.old): Model convergence
## problem; false convergence (8). See vignette('troubleshooting'),
## help('diagnose')
```

```
## Warning in log(inter_eye_dist_change): NaNs produced
## Warning in log(inter_eye_dist_change): NaNs produced
```

```
## Warning in finalizeTMB(TMBStruc, obj, fit, h, data.tmb.old): Model convergence
## problem; non-positive-definite Hessian matrix. See vignette('troubleshooting')
```

```
## Warning in finalizeTMB(TMBStruc, obj, fit, h, data.tmb.old): Model convergence
## problem; singular convergence (7). See vignette('troubleshooting'),
## help('diagnose')
```

```
## Warning in log(inter_eye_dist_change): NaNs produced
## Warning in log(inter_eye_dist_change): NaNs produced
```

```
## Warning in finalizeTMB(TMBStruc, obj, fit, h, data.tmb.old): Model convergence
## problem; non-positive-definite Hessian matrix. See vignette('troubleshooting')
```

```
## Warning in finalizeTMB(TMBStruc, obj, fit, h, data.tmb.old): Model convergence
## problem; false convergence (8). See vignette('troubleshooting'),
## help('diagnose')
```

```
## Warning in log(inter_eye_dist_change): NaNs produced
## Warning in log(inter_eye_dist_change): NaNs produced
## Warning in log(inter_eye_dist_change): NaNs produced
## Warning in log(inter_eye_dist_change): NaNs produced
## Warning in log(inter_eye_dist_change): NaNs produced
## Warning in log(inter_eye_dist_change): NaNs produced
```

```
## Warning in finalizeTMB(TMBStruc, obj, fit, h, data.tmb.old): Model convergence
## problem; non-positive-definite Hessian matrix. See vignette('troubleshooting')
```

```
## Warning in log(inter_eye_dist_change): NaNs produced
## Warning in log(inter_eye_dist_change): NaNs produced
```

```
## Warning in finalizeTMB(TMBStruc, obj, fit, h, data.tmb.old): Model convergence
## problem; non-positive-definite Hessian matrix. See vignette('troubleshooting')
```

```
## Warning in finalizeTMB(TMBStruc, obj, fit, h, data.tmb.old): Model convergence
## problem; singular convergence (7). See vignette('troubleshooting'),
## help('diagnose')
```

```
## Warning in log(inter_eye_dist_change): NaNs produced
## Warning in log(inter_eye_dist_change): NaNs produced
## Warning in log(inter_eye_dist_change): NaNs produced
## Warning in log(inter_eye_dist_change): NaNs produced
## Warning in log(inter_eye_dist_change): NaNs produced
## Warning in log(inter_eye_dist_change): NaNs produced
```

```
## Warning in finalizeTMB(TMBStruc, obj, fit, h, data.tmb.old): Model convergence
## problem; non-positive-definite Hessian matrix. See vignette('troubleshooting')
```

```
## Warning in finalizeTMB(TMBStruc, obj, fit, h, data.tmb.old): Model convergence
## problem; singular convergence (7). See vignette('troubleshooting'),
## help('diagnose')
```

```
## Warning in log(inter_eye_dist_change): NaNs produced
## Warning in log(inter_eye_dist_change): NaNs produced
## Warning in log(inter_eye_dist_change): NaNs produced
## Warning in log(inter_eye_dist_change): NaNs produced
## Warning in log(inter_eye_dist_change): NaNs produced
## Warning in log(inter_eye_dist_change): NaNs produced
```

```
## Warning in finalizeTMB(TMBStruc, obj, fit, h, data.tmb.old): Model convergence
## problem; non-positive-definite Hessian matrix. See vignette('troubleshooting')
```

```
## Warning in finalizeTMB(TMBStruc, obj, fit, h, data.tmb.old): Model convergence
## problem; false convergence (8). See vignette('troubleshooting'),
## help('diagnose')
```

```
## Warning in log(inter_eye_dist_change): NaNs produced
## Warning in log(inter_eye_dist_change): NaNs produced
## Warning in log(inter_eye_dist_change): NaNs produced
## Warning in log(inter_eye_dist_change): NaNs produced
```

```
## Warning in finalizeTMB(TMBStruc, obj, fit, h, data.tmb.old): Model convergence
## problem; non-positive-definite Hessian matrix. See vignette('troubleshooting')
```

```
## Warning in finalizeTMB(TMBStruc, obj, fit, h, data.tmb.old): Model convergence
## problem; singular convergence (7). See vignette('troubleshooting'),
## help('diagnose')
```

```
## Warning in log(inter_eye_dist_change): NaNs produced
## Warning in log(inter_eye_dist_change): NaNs produced
```

```
## Warning in finalizeTMB(TMBStruc, obj, fit, h, data.tmb.old): Model convergence
## problem; non-positive-definite Hessian matrix. See vignette('troubleshooting')
```

```
## Warning in log(inter_eye_dist_change): NaNs produced
## Warning in log(inter_eye_dist_change): NaNs produced
## Warning in log(inter_eye_dist_change): NaNs produced
## Warning in log(inter_eye_dist_change): NaNs produced
```

```
## Warning in finalizeTMB(TMBStruc, obj, fit, h, data.tmb.old): Model convergence
## problem; non-positive-definite Hessian matrix. See vignette('troubleshooting')
```

```
## Warning in finalizeTMB(TMBStruc, obj, fit, h, data.tmb.old): Model convergence
## problem; singular convergence (7). See vignette('troubleshooting'),
## help('diagnose')
```

```
## Warning in log(inter_eye_dist_change): NaNs produced
## Warning in log(inter_eye_dist_change): NaNs produced
```

```
## Warning in finalizeTMB(TMBStruc, obj, fit, h, data.tmb.old): Model convergence
## problem; non-positive-definite Hessian matrix. See vignette('troubleshooting')
```

```
## Warning in finalizeTMB(TMBStruc, obj, fit, h, data.tmb.old): Model convergence
## problem; singular convergence (7). See vignette('troubleshooting'),
## help('diagnose')
```

```
## Warning in log(inter_eye_dist_change): NaNs produced
## Warning in log(inter_eye_dist_change): NaNs produced
```

```
## Warning in finalizeTMB(TMBStruc, obj, fit, h, data.tmb.old): Model convergence
## problem; non-positive-definite Hessian matrix. See vignette('troubleshooting')
```

```
## Warning in log(inter_eye_dist_change): NaNs produced
## Warning in log(inter_eye_dist_change): NaNs produced
```

```
## Warning in finalizeTMB(TMBStruc, obj, fit, h, data.tmb.old): Model convergence
## problem; non-positive-definite Hessian matrix. See vignette('troubleshooting')
```

```
## Warning in log(inter_eye_dist_change): NaNs produced
## Warning in log(inter_eye_dist_change): NaNs produced
```

```
## Warning in finalizeTMB(TMBStruc, obj, fit, h, data.tmb.old): Model convergence
## problem; non-positive-definite Hessian matrix. See vignette('troubleshooting')
```

```
## Warning in finalizeTMB(TMBStruc, obj, fit, h, data.tmb.old): Model convergence
## problem; singular convergence (7). See vignette('troubleshooting'),
## help('diagnose')
```

```
## Warning in log(inter_eye_dist_change): NaNs produced
## Warning in log(inter_eye_dist_change): NaNs produced
```

```
## Warning in finalizeTMB(TMBStruc, obj, fit, h, data.tmb.old): Model convergence
## problem; non-positive-definite Hessian matrix. See vignette('troubleshooting')
```

```
## Warning in log(inter_eye_dist_change): NaNs produced
## Warning in log(inter_eye_dist_change): NaNs produced
```

```
## Warning in finalizeTMB(TMBStruc, obj, fit, h, data.tmb.old): Model convergence
## problem; non-positive-definite Hessian matrix. See vignette('troubleshooting')
```

```
## Warning in log(inter_eye_dist_change): NaNs produced
## Warning in log(inter_eye_dist_change): NaNs produced
## Warning in log(inter_eye_dist_change): NaNs produced
## Warning in log(inter_eye_dist_change): NaNs produced
## Warning in log(inter_eye_dist_change): NaNs produced
## Warning in log(inter_eye_dist_change): NaNs produced
```

```
## Warning in finalizeTMB(TMBStruc, obj, fit, h, data.tmb.old): Model convergence
## problem; non-positive-definite Hessian matrix. See vignette('troubleshooting')
```

```
## Warning in finalizeTMB(TMBStruc, obj, fit, h, data.tmb.old): Model convergence
## problem; singular convergence (7). See vignette('troubleshooting'),
## help('diagnose')
```

```
## Warning in log(inter_eye_dist_change): NaNs produced
## Warning in log(inter_eye_dist_change): NaNs produced
```

```
## Warning in finalizeTMB(TMBStruc, obj, fit, h, data.tmb.old): Model convergence
## problem; non-positive-definite Hessian matrix. See vignette('troubleshooting')
```

```
## Warning in finalizeTMB(TMBStruc, obj, fit, h, data.tmb.old): Model convergence
## problem; singular convergence (7). See vignette('troubleshooting'),
## help('diagnose')
```

```
## Warning in log(inter_eye_dist_change): NaNs produced
## Warning in log(inter_eye_dist_change): NaNs produced
```

```
## Warning in finalizeTMB(TMBStruc, obj, fit, h, data.tmb.old): Model convergence
## problem; non-positive-definite Hessian matrix. See vignette('troubleshooting')
```

```
## Warning in log(inter_eye_dist_change): NaNs produced
## Warning in log(inter_eye_dist_change): NaNs produced
```

```
## Warning in finalizeTMB(TMBStruc, obj, fit, h, data.tmb.old): Model convergence
## problem; non-positive-definite Hessian matrix. See vignette('troubleshooting')
```

```
## Warning in log(inter_eye_dist_change): NaNs produced
## Warning in log(inter_eye_dist_change): NaNs produced
```

```
## Warning in finalizeTMB(TMBStruc, obj, fit, h, data.tmb.old): Model convergence
## problem; non-positive-definite Hessian matrix. See vignette('troubleshooting')
```

```
## Warning in finalizeTMB(TMBStruc, obj, fit, h, data.tmb.old): Model convergence
## problem; singular convergence (7). See vignette('troubleshooting'),
## help('diagnose')
```

```
## Warning in log(inter_eye_dist_change): NaNs produced
## Warning in log(inter_eye_dist_change): NaNs produced
## Warning in log(inter_eye_dist_change): NaNs produced
## Warning in log(inter_eye_dist_change): NaNs produced
```

```
## Warning in finalizeTMB(TMBStruc, obj, fit, h, data.tmb.old): Model convergence
## problem; non-positive-definite Hessian matrix. See vignette('troubleshooting')
```

```
## Warning in log(inter_eye_dist_change): NaNs produced
## Warning in log(inter_eye_dist_change): NaNs produced
## Warning in log(inter_eye_dist_change): NaNs produced
## Warning in log(inter_eye_dist_change): NaNs produced
```

```
## Warning in finalizeTMB(TMBStruc, obj, fit, h, data.tmb.old): Model convergence
## problem; non-positive-definite Hessian matrix. See vignette('troubleshooting')
```

```
## Warning in log(inter_eye_dist_change): NaNs produced
## Warning in log(inter_eye_dist_change): NaNs produced
## Warning in log(inter_eye_dist_change): NaNs produced
## Warning in log(inter_eye_dist_change): NaNs produced
```

```
## Warning in finalizeTMB(TMBStruc, obj, fit, h, data.tmb.old): Model convergence
## problem; non-positive-definite Hessian matrix. See vignette('troubleshooting')
```

```
## Warning in finalizeTMB(TMBStruc, obj, fit, h, data.tmb.old): Model convergence
## problem; singular convergence (7). See vignette('troubleshooting'),
## help('diagnose')
```

```
## Warning in log(inter_eye_dist_change): NaNs produced
## Warning in log(inter_eye_dist_change): NaNs produced
```

```
## Warning in finalizeTMB(TMBStruc, obj, fit, h, data.tmb.old): Model convergence
## problem; non-positive-definite Hessian matrix. See vignette('troubleshooting')
```

```
## Warning in finalizeTMB(TMBStruc, obj, fit, h, data.tmb.old): Model convergence
## problem; false convergence (8). See vignette('troubleshooting'),
## help('diagnose')
```

```
## Warning in log(inter_eye_dist_change): NaNs produced
## Warning in log(inter_eye_dist_change): NaNs produced
```

```
## Warning in finalizeTMB(TMBStruc, obj, fit, h, data.tmb.old): Model convergence
## problem; non-positive-definite Hessian matrix. See vignette('troubleshooting')
```

```
## Warning in log(inter_eye_dist_change): NaNs produced
## Warning in log(inter_eye_dist_change): NaNs produced
## Warning in log(inter_eye_dist_change): NaNs produced
## Warning in log(inter_eye_dist_change): NaNs produced
## Warning in log(inter_eye_dist_change): NaNs produced
## Warning in log(inter_eye_dist_change): NaNs produced
## Warning in log(inter_eye_dist_change): NaNs produced
## Warning in log(inter_eye_dist_change): NaNs produced
```

```
## Warning in finalizeTMB(TMBStruc, obj, fit, h, data.tmb.old): Model convergence
## problem; non-positive-definite Hessian matrix. See vignette('troubleshooting')
```

```
## Warning in log(inter_eye_dist_change): NaNs produced
## Warning in log(inter_eye_dist_change): NaNs produced
```

```
## Warning in finalizeTMB(TMBStruc, obj, fit, h, data.tmb.old): Model convergence
## problem; non-positive-definite Hessian matrix. See vignette('troubleshooting')
```

```
## Warning in log(inter_eye_dist_change): NaNs produced
## Warning in log(inter_eye_dist_change): NaNs produced
## Warning in log(inter_eye_dist_change): NaNs produced
## Warning in log(inter_eye_dist_change): NaNs produced
```

```
## Warning in finalizeTMB(TMBStruc, obj, fit, h, data.tmb.old): Model convergence
## problem; singular convergence (7). See vignette('troubleshooting'),
## help('diagnose')
```

```
## Warning in log(inter_eye_dist_change): NaNs produced
## Warning in log(inter_eye_dist_change): NaNs produced
## Warning in log(inter_eye_dist_change): NaNs produced
## Warning in log(inter_eye_dist_change): NaNs produced
```

```
## Warning in finalizeTMB(TMBStruc, obj, fit, h, data.tmb.old): Model convergence
## problem; singular convergence (7). See vignette('troubleshooting'),
## help('diagnose')
```

```
## Warning in log(inter_eye_dist_change): NaNs produced
## Warning in log(inter_eye_dist_change): NaNs produced
```

```
## Warning in finalizeTMB(TMBStruc, obj, fit, h, data.tmb.old): Model convergence
## problem; non-positive-definite Hessian matrix. See vignette('troubleshooting')
```

```
## Warning in finalizeTMB(TMBStruc, obj, fit, h, data.tmb.old): Model convergence
## problem; singular convergence (7). See vignette('troubleshooting'),
## help('diagnose')
```

```
## Warning in log(inter_eye_dist_change): NaNs produced
## Warning in log(inter_eye_dist_change): NaNs produced
```

```
## Warning in finalizeTMB(TMBStruc, obj, fit, h, data.tmb.old): Model convergence
## problem; non-positive-definite Hessian matrix. See vignette('troubleshooting')
```

```
## Warning in log(inter_eye_dist_change): NaNs produced
## Warning in log(inter_eye_dist_change): NaNs produced
```

```
## Warning in finalizeTMB(TMBStruc, obj, fit, h, data.tmb.old): Model convergence
## problem; non-positive-definite Hessian matrix. See vignette('troubleshooting')
```

```
## Warning in finalizeTMB(TMBStruc, obj, fit, h, data.tmb.old): Model convergence
## problem; singular convergence (7). See vignette('troubleshooting'),
## help('diagnose')
```

```
## Warning in log(inter_eye_dist_change): NaNs produced
## Warning in log(inter_eye_dist_change): NaNs produced
## Warning in log(inter_eye_dist_change): NaNs produced
## Warning in log(inter_eye_dist_change): NaNs produced
```

```
## Warning in finalizeTMB(TMBStruc, obj, fit, h, data.tmb.old): Model convergence
## problem; non-positive-definite Hessian matrix. See vignette('troubleshooting')
```

```
## Warning in finalizeTMB(TMBStruc, obj, fit, h, data.tmb.old): Model convergence
## problem; singular convergence (7). See vignette('troubleshooting'),
## help('diagnose')
```

```
## Warning in log(inter_eye_dist_change): NaNs produced
## Warning in log(inter_eye_dist_change): NaNs produced
```

```
## Warning in finalizeTMB(TMBStruc, obj, fit, h, data.tmb.old): Model convergence
## problem; non-positive-definite Hessian matrix. See vignette('troubleshooting')
```

```
## Warning in finalizeTMB(TMBStruc, obj, fit, h, data.tmb.old): Model convergence
## problem; singular convergence (7). See vignette('troubleshooting'),
## help('diagnose')
```

```
## Warning in log(inter_eye_dist_change): NaNs produced
## Warning in log(inter_eye_dist_change): NaNs produced
```

```
## Warning in finalizeTMB(TMBStruc, obj, fit, h, data.tmb.old): Model convergence
## problem; non-positive-definite Hessian matrix. See vignette('troubleshooting')
```

```
## Warning in finalizeTMB(TMBStruc, obj, fit, h, data.tmb.old): Model convergence
## problem; singular convergence (7). See vignette('troubleshooting'),
## help('diagnose')
```

```
## Warning in log(inter_eye_dist_change): NaNs produced
## Warning in log(inter_eye_dist_change): NaNs produced
## Warning in log(inter_eye_dist_change): NaNs produced
## Warning in log(inter_eye_dist_change): NaNs produced
## Warning in log(inter_eye_dist_change): NaNs produced
## Warning in log(inter_eye_dist_change): NaNs produced
## Warning in log(inter_eye_dist_change): NaNs produced
## Warning in log(inter_eye_dist_change): NaNs produced
## Warning in log(inter_eye_dist_change): NaNs produced
## Warning in log(inter_eye_dist_change): NaNs produced
## Warning in log(inter_eye_dist_change): NaNs produced
## Warning in log(inter_eye_dist_change): NaNs produced
## Warning in log(inter_eye_dist_change): NaNs produced
## Warning in log(inter_eye_dist_change): NaNs produced
## Warning in log(inter_eye_dist_change): NaNs produced
## Warning in log(inter_eye_dist_change): NaNs produced
```

```
## Warning in finalizeTMB(TMBStruc, obj, fit, h, data.tmb.old): Model convergence
## problem; non-positive-definite Hessian matrix. See vignette('troubleshooting')
```

```
## Warning in log(inter_eye_dist_change): NaNs produced
## Warning in log(inter_eye_dist_change): NaNs produced
```

```
## Warning in finalizeTMB(TMBStruc, obj, fit, h, data.tmb.old): Model convergence
## problem; non-positive-definite Hessian matrix. See vignette('troubleshooting')
```

```
## Warning in finalizeTMB(TMBStruc, obj, fit, h, data.tmb.old): Model convergence
## problem; false convergence (8). See vignette('troubleshooting'),
## help('diagnose')
```

```
## Warning in log(inter_eye_dist_change): NaNs produced
## Warning in log(inter_eye_dist_change): NaNs produced
```

```
## Warning in finalizeTMB(TMBStruc, obj, fit, h, data.tmb.old): Model convergence
## problem; non-positive-definite Hessian matrix. See vignette('troubleshooting')
```

```
## Warning in finalizeTMB(TMBStruc, obj, fit, h, data.tmb.old): Model convergence
## problem; false convergence (8). See vignette('troubleshooting'),
## help('diagnose')
```

```
## Warning in log(inter_eye_dist_change): NaNs produced
## Warning in log(inter_eye_dist_change): NaNs produced
```

```
## Warning in finalizeTMB(TMBStruc, obj, fit, h, data.tmb.old): Model convergence
## problem; non-positive-definite Hessian matrix. See vignette('troubleshooting')
```

```
## Warning in log(inter_eye_dist_change): NaNs produced
## Warning in log(inter_eye_dist_change): NaNs produced
```

```
## Warning in finalizeTMB(TMBStruc, obj, fit, h, data.tmb.old): Model convergence
## problem; non-positive-definite Hessian matrix. See vignette('troubleshooting')
```

```
## Warning in finalizeTMB(TMBStruc, obj, fit, h, data.tmb.old): Model convergence
## problem; singular convergence (7). See vignette('troubleshooting'),
## help('diagnose')
```

```
## Warning in log(inter_eye_dist_change): NaNs produced
## Warning in log(inter_eye_dist_change): NaNs produced
```

```
## Warning in finalizeTMB(TMBStruc, obj, fit, h, data.tmb.old): Model convergence
## problem; non-positive-definite Hessian matrix. See vignette('troubleshooting')
```

```
## Warning in finalizeTMB(TMBStruc, obj, fit, h, data.tmb.old): Model convergence
## problem; singular convergence (7). See vignette('troubleshooting'),
## help('diagnose')
```

```
## Warning in log(inter_eye_dist_change): NaNs produced
## Warning in log(inter_eye_dist_change): NaNs produced
```

```
## Warning in finalizeTMB(TMBStruc, obj, fit, h, data.tmb.old): Model convergence
## problem; non-positive-definite Hessian matrix. See vignette('troubleshooting')
```

```
## Warning in finalizeTMB(TMBStruc, obj, fit, h, data.tmb.old): Model convergence
## problem; false convergence (8). See vignette('troubleshooting'),
## help('diagnose')
```

```
## Warning in log(inter_eye_dist_change): NaNs produced
## Warning in log(inter_eye_dist_change): NaNs produced
```

```
## Warning in finalizeTMB(TMBStruc, obj, fit, h, data.tmb.old): Model convergence
## problem; non-positive-definite Hessian matrix. See vignette('troubleshooting')
```

```
## Warning in finalizeTMB(TMBStruc, obj, fit, h, data.tmb.old): Model convergence
## problem; singular convergence (7). See vignette('troubleshooting'),
## help('diagnose')
```

```
## Warning in log(inter_eye_dist_change): NaNs produced
## Warning in log(inter_eye_dist_change): NaNs produced
```

```
## Warning in finalizeTMB(TMBStruc, obj, fit, h, data.tmb.old): Model convergence
## problem; non-positive-definite Hessian matrix. See vignette('troubleshooting')
```

```
## Warning in log(inter_eye_dist_change): NaNs produced
## Warning in log(inter_eye_dist_change): NaNs produced
## Warning in log(inter_eye_dist_change): NaNs produced
## Warning in log(inter_eye_dist_change): NaNs produced
```

```
## Warning in finalizeTMB(TMBStruc, obj, fit, h, data.tmb.old): Model convergence
## problem; non-positive-definite Hessian matrix. See vignette('troubleshooting')
```

```
## Warning in finalizeTMB(TMBStruc, obj, fit, h, data.tmb.old): Model convergence
## problem; singular convergence (7). See vignette('troubleshooting'),
## help('diagnose')
```

```
## Warning in log(inter_eye_dist_change): NaNs produced
## Warning in log(inter_eye_dist_change): NaNs produced
```

```
## Warning in finalizeTMB(TMBStruc, obj, fit, h, data.tmb.old): Model convergence
## problem; non-positive-definite Hessian matrix. See vignette('troubleshooting')
```

```
## Warning in finalizeTMB(TMBStruc, obj, fit, h, data.tmb.old): Model convergence
## problem; singular convergence (7). See vignette('troubleshooting'),
## help('diagnose')
```

```
## Warning in log(inter_eye_dist_change): NaNs produced
## Warning in log(inter_eye_dist_change): NaNs produced
```

```
## Warning in finalizeTMB(TMBStruc, obj, fit, h, data.tmb.old): Model convergence
## problem; non-positive-definite Hessian matrix. See vignette('troubleshooting')
```

```
## Warning in log(inter_eye_dist_change): NaNs produced
## Warning in log(inter_eye_dist_change): NaNs produced
```

```
## Warning in finalizeTMB(TMBStruc, obj, fit, h, data.tmb.old): Model convergence
## problem; non-positive-definite Hessian matrix. See vignette('troubleshooting')
```

```
## Warning in finalizeTMB(TMBStruc, obj, fit, h, data.tmb.old): Model convergence
## problem; singular convergence (7). See vignette('troubleshooting'),
## help('diagnose')
```

Saving routine

```
moving_results <- data.frame(list('win_starts'=NaN,
                                  'win_span'=NaN,
                                  'p'=NaN,
                                  'stimtype'=NaN,
'emmean'=NaN, 'SE'=NaN, 'CL.upper'=NaN, 'CL.lower'=NaN))
for (i in 1:length(windows_starts)){
  window_start <- windows_starts[i]
  p <- windows_p_moving[[i]][[1]]
  for (stimtype in c('bio', 'ellipse', 'scramb', 'silhouette', 'rand')){
  newrow <- c(window_start, window_size, p, stimtype,
              windows_p_moving[[i]]$response[windows_p_moving[[i]]$stimtype==stimtype],
  windows_p_moving[[i]]$SE[windows_p_moving[[i]]$stimtype==stimtype],
              windows_p_moving[[i]]$upper.CL[windows_p_moving[[i]]$stimtype==stimtype],
  windows_p_moving[[i]]$lower.CL[windows_p_moving[[i]]$stimtype==stimtype])
    moving_results <- rbind(moving_results, newrow)
  }

}

moving_results <- moving_results[-1,]

write.csv(moving_results, paste0(tmp_path, 'interEyeDistChange_movingResults.csv'))
```

##### 2.1.4 Dist\_from\_center

###### 2.1.4.1 Static

```
onlystatic <- subset(full_data, full_data$moving == 0)
onlynonemaxframe <- max(onlystatic[onlystatic$stimtype=='none']$time_from_start)
onlystatic <- subset(onlystatic, onlystatic$time_from_start<onlynonemaxframe)

start_frame <- min(onlystatic$time_from_start)
end_frame <- max(onlystatic$time_from_start)

windows_starts <- seq(start_frame, end_frame-window_size, step_size)

data <- onlystatic
plan(multisession, workers = num_cores)
windows_p_static <- future_lapply(windows_starts, windowed_model_dist_from_center, future.seed=TRUE)
```

Saving routine

```
static_results <- data.frame(list('win_starts'=NaN,
                                  'win_span'=NaN,
                                  'p'=NaN,
                                  'stimtype'=NaN,
'emmean'=NaN, 'SE'=NaN, 'CL.upper'=NaN, 'CL.lower'=NaN))

for (i in 1:length(windows_starts)){
  window_start <- windows_starts[i]
  p <- windows_p_static[[i]][[1]]
  for (stimtype in c('none', 'bio', 'ellipse', 'scramb', 'silhouette', 'rand')){
  newrow <- c(window_start, window_size, p, stimtype,
              windows_p_static[[i]]$response[windows_p_static[[i]]$stimtype==stimtype],
  windows_p_static[[i]]$SE[windows_p_static[[i]]$stimtype==stimtype],
              windows_p_static[[i]]$upper.CL[windows_p_static[[i]]$stimtype==stimtype],
  windows_p_static[[i]]$lower.CL[windows_p_static[[i]]$stimtype==stimtype])
    static_results <- rbind(static_results, newrow)
  }

}

static_results <- static_results[-1,]

write.csv(static_results, paste0(tmp_path, 'distFromCenter_staticResults.csv'))
```

###### 2.1.4.2 Moving

```
onlymoving <- subset(full_data, full_data$stimtype != 'none')

start_frame <- min(onlymoving$time_from_start)
end_frame <- max(onlymoving$time_from_start)

windows_starts <- seq(start_frame, end_frame-window_size, step_size)
winsizes <- rep(window_size, length(windows_starts))

data <- onlymoving
plan(multisession, workers = num_cores)
windows_p_moving <- future_lapply(windows_starts, windowed_model_dist_from_center, future.seed=TRUE)
```

Saving routine

```
moving_results <- data.frame(list('win_starts'=NaN,
                                  'win_span'=NaN,
                                  'p'=NaN,
                                  'stimtype'=NaN,
'emmean'=NaN, 'SE'=NaN, 'CL.upper'=NaN, 'CL.lower'=NaN))
for (i in 1:length(windows_starts)){
  window_start <- windows_starts[i]
  p <- windows_p_moving[[i]][[1]]
  for (stimtype in c('bio', 'ellipse', 'scramb', 'silhouette', 'rand')){
  newrow <- c(window_start, window_size, p, stimtype,
              windows_p_moving[[i]]$response[windows_p_moving[[i]]$stimtype==stimtype],
  windows_p_moving[[i]]$SE[windows_p_moving[[i]]$stimtype==stimtype],
              windows_p_moving[[i]]$upper.CL[windows_p_moving[[i]]$stimtype==stimtype],
  windows_p_moving[[i]]$lower.CL[windows_p_moving[[i]]$stimtype==stimtype])
    moving_results <- rbind(moving_results, newrow)
  }

}

moving_results <- moving_results[-1,]

write.csv(moving_results, paste0(tmp_path, 'distFromCenter_movingResults.csv'))
```

#### 2.2 In depth analysis

Now that we have the whole time series analysis, we can select
section that resulted significant (including the multiple comparison
correction) and do pairwise post-hoc

##### 2.2.1 Eye displacement

###### 2.2.1.1 Static

remaking specific dataframe

```
onlystatic <- subset(full_data, full_data$moving == 0)

data <- onlystatic
```

reloading of the data and ps

```
static <- read.csv(paste0(tmp_path, 'displacement_staticResults.csv'))

static$adjustedp <- static$p*sum(static$stimtype=='bio') # create bonferroni corrected p
static$adjustedp[static$adjustedp>1] <- 1
significant_static <- subset(static, static$stimtype=='bio') #subset to have only 1 row per model

significant_static <- subset(significant_static, significant_static$adjustedp < alpha)

significant_static_wins <- significant_static$win_starts #identify the windows that are significant
```

###### 2.2.1.1.1 Detailed

now redo the models

```
static_results_posthoc <- data.frame(list('win_starts'=NaN,'win_span'=NaN,
                                          'bio/ellipse'=NaN, 'bio/none'=NaN,
                                          'bio/rand'=NaN, 'bio/scramb'=NaN,
                                          'bio/silhouette'=NaN, 'ellipse/none'=NaN,
                                          'ellipse/rand'=NaN, 'ellipse/scramb'=NaN,
                                          'ellipse/silhouette'=NaN, 'none/rand'=NaN,
                                          'none/scramb'=NaN, 'none/silhouette'=NaN,
                                          'rand/scramb'=NaN, 'rand/silhouette'=NaN,
                                          'scramb/silhouette'=NaN))

for (window_start in significant_static_wins){
  window_end <- window_start + window_size

  window_data <- subset(data, data$time_from_start>window_start)
  window_data <- subset(window_data, window_data$time_from_start<window_end)

  full_tmb <- glmmTMB(log(midpoint_displacement) ~ stimtype + (stimtype|subj), family=gaussian, data=window_data)
  Anova(full_tmb)
  e <- emmeans(full_tmb, ~stimtype, type='response')
  prs <- as.data.frame(pairs(e))
  newrow <- c(window_start, window_size, prs$p.value[1], prs$p.value[2],
              prs$p.value[3], prs$p.value[4], prs$p.value[5], prs$p.value[6],
              prs$p.value[7], prs$p.value[8], prs$p.value[9], prs$p.value[10],
              prs$p.value[11], prs$p.value[12], prs$p.value[13], prs$p.value[14],
              prs$p.value[15])
  static_results_posthoc <- rbind(static_results_posthoc, newrow)
  print(window_start)
  print(pairs(e))
}
```

```
## [1] 0.03333333
##  contrast             ratio     SE    df null t.ratio p.value
##  bio / ellipse        0.989 0.0543 24648    1  -0.199  1.0000
##  bio / none           1.202 0.0588 24648    1   3.774  0.0022
##  bio / rand           0.980 0.0691 24648    1  -0.288  0.9997
##  bio / scramb         0.998 0.0505 24648    1  -0.031  1.0000
##  bio / silhouette     0.921 0.0658 24648    1  -1.153  0.8590
##  ellipse / none       1.216 0.0560 24648    1   4.241  0.0003
##  ellipse / rand       0.991 0.0598 24648    1  -0.156  1.0000
##  ellipse / scramb     1.009 0.0619 24648    1   0.152  1.0000
##  ellipse / silhouette 0.931 0.0545 24648    1  -1.221  0.8266
##  none / rand          0.815 0.0528 24648    1  -3.162  0.0196
##  none / scramb        0.830 0.0493 24648    1  -3.131  0.0216
##  none / silhouette    0.766 0.0471 24648    1  -4.340  0.0002
##  rand / scramb        1.019 0.0583 24648    1   0.328  0.9995
##  rand / silhouette    0.940 0.0621 24648    1  -0.939  0.9365
##  scramb / silhouette  0.922 0.0691 24648    1  -1.080  0.8896
## 
## P value adjustment: tukey method for comparing a family of 6 estimates 
## Tests are performed on the log scale 
## [1] 0.2
##  contrast             ratio     SE    df null t.ratio p.value
##  bio / ellipse        0.973 0.0551 23577    1  -0.476  0.9970
##  bio / none           1.277 0.0687 23577    1   4.540  0.0001
##  bio / rand           0.978 0.0737 23577    1  -0.292  0.9997
##  bio / scramb         1.005 0.0593 23577    1   0.091  1.0000
##  bio / silhouette     0.891 0.0654 23577    1  -1.571  0.6175
##  ellipse / none       1.312 0.0763 23577    1   4.663  <.0001
##  ellipse / rand       1.005 0.0680 23577    1   0.073  1.0000
##  ellipse / scramb     1.033 0.0704 23577    1   0.474  0.9970
##  ellipse / silhouette 0.915 0.0670 23577    1  -1.208  0.8331
##  none / rand          0.766 0.0530 23577    1  -3.848  0.0017
##  none / scramb        0.788 0.0534 23577    1  -3.524  0.0057
##  none / silhouette    0.698 0.0473 23577    1  -5.301  <.0001
##  rand / scramb        1.028 0.0588 23577    1   0.479  0.9969
##  rand / silhouette    0.911 0.0587 23577    1  -1.449  0.6967
##  scramb / silhouette  0.886 0.0697 23577    1  -1.536  0.6411
## 
## P value adjustment: tukey method for comparing a family of 6 estimates 
## Tests are performed on the log scale 
## [1] 0.3666667
##  contrast             ratio     SE    df null t.ratio p.value
##  bio / ellipse        0.996 0.0651 24423    1  -0.061  1.0000
##  bio / none           1.285 0.0740 24423    1   4.352  0.0002
##  bio / rand           0.998 0.0706 24423    1  -0.023  1.0000
##  bio / scramb         1.013 0.0652 24423    1   0.205  1.0000
##  bio / silhouette     0.947 0.0650 24423    1  -0.788  0.9696
##  ellipse / none       1.290 0.0764 24423    1   4.303  0.0002
##  ellipse / rand       1.002 0.0648 24423    1   0.037  1.0000
##  ellipse / scramb     1.017 0.0691 24423    1   0.253  0.9999
##  ellipse / silhouette 0.951 0.0698 24423    1  -0.683  0.9839
##  none / rand          0.777 0.0513 24423    1  -3.818  0.0019
##  none / scramb        0.789 0.0528 24423    1  -3.550  0.0052
##  none / silhouette    0.737 0.0432 24423    1  -5.207  <.0001
##  rand / scramb        1.015 0.0607 24423    1   0.247  0.9999
##  rand / silhouette    0.949 0.0627 24423    1  -0.795  0.9685
##  scramb / silhouette  0.935 0.0745 24423    1  -0.845  0.9591
## 
## P value adjustment: tukey method for comparing a family of 6 estimates 
## Tests are performed on the log scale 
## [1] 0.5333333
##  contrast             ratio     SE    df null t.ratio p.value
##  bio / ellipse        1.001 0.0671 24455    1   0.011  1.0000
##  bio / none           1.268 0.0745 24455    1   4.034  0.0008
##  bio / rand           1.023 0.0741 24455    1   0.313  0.9996
##  bio / scramb         1.049 0.0736 24455    1   0.677  0.9845
##  bio / silhouette     0.937 0.0696 24455    1  -0.878  0.9518
##  ellipse / none       1.267 0.0772 24455    1   3.882  0.0015
##  ellipse / rand       1.022 0.0660 24455    1   0.340  0.9994
##  ellipse / scramb     1.048 0.0728 24455    1   0.674  0.9848
##  ellipse / silhouette 0.936 0.0723 24455    1  -0.853  0.9574
##  none / rand          0.807 0.0538 24455    1  -3.220  0.0162
##  none / scramb        0.827 0.0561 24455    1  -2.799  0.0576
##  none / silhouette    0.739 0.0433 24455    1  -5.160  <.0001
##  rand / scramb        1.025 0.0544 24455    1   0.469  0.9972
##  rand / silhouette    0.916 0.0657 24455    1  -1.226  0.8245
##  scramb / silhouette  0.893 0.0733 24455    1  -1.373  0.7431
## 
## P value adjustment: tukey method for comparing a family of 6 estimates 
## Tests are performed on the log scale 
## [1] 0.7
##  contrast             ratio     SE    df null t.ratio p.value
##  bio / ellipse        1.017 0.0756 23584    1   0.224  0.9999
##  bio / none           1.241 0.0724 23584    1   3.700  0.0030
##  bio / rand           1.008 0.0753 23584    1   0.104  1.0000
##  bio / scramb         1.021 0.0820 23584    1   0.262  0.9998
##  bio / silhouette     0.929 0.0653 23584    1  -1.044  0.9030
##  ellipse / none       1.221 0.0679 23584    1   3.580  0.0046
##  ellipse / rand       0.991 0.0615 23584    1  -0.143  1.0000
##  ellipse / scramb     1.004 0.0704 23584    1   0.063  1.0000
##  ellipse / silhouette 0.914 0.0661 23584    1  -1.244  0.8153
##  none / rand          0.812 0.0551 23584    1  -3.070  0.0261
##  none / scramb        0.823 0.0568 23584    1  -2.825  0.0536
##  none / silhouette    0.749 0.0428 23584    1  -5.060  <.0001
##  rand / scramb        1.013 0.0577 23584    1   0.233  0.9999
##  rand / silhouette    0.922 0.0622 23584    1  -1.201  0.8364
##  scramb / silhouette  0.910 0.0746 23584    1  -1.150  0.8603
## 
## P value adjustment: tukey method for comparing a family of 6 estimates 
## Tests are performed on the log scale 
## [1] 0.8666667
##  contrast             ratio     SE    df null t.ratio p.value
##  bio / ellipse        1.007 0.0720 24314    1   0.098  1.0000
##  bio / none           1.220 0.0756 24314    1   3.210  0.0168
##  bio / rand           1.013 0.0726 24314    1   0.182  1.0000
##  bio / scramb         1.030 0.0838 24314    1   0.365  0.9992
##  bio / silhouette     0.929 0.0715 24314    1  -0.963  0.9296
##  ellipse / none       1.212 0.0613 24314    1   3.796  0.0020
##  ellipse / rand       1.006 0.0635 24314    1   0.095  1.0000
##  ellipse / scramb     1.023 0.0749 24314    1   0.310  0.9996
##  ellipse / silhouette 0.922 0.0637 24314    1  -1.174  0.8495
##  none / rand          0.830 0.0571 24314    1  -2.702  0.0748
##  none / scramb        0.844 0.0599 24314    1  -2.385  0.1614
##  none / silhouette    0.761 0.0455 24314    1  -4.572  0.0001
##  rand / scramb        1.017 0.0653 24314    1   0.260  0.9998
##  rand / silhouette    0.917 0.0648 24314    1  -1.232  0.8215
##  scramb / silhouette  0.901 0.0817 24314    1  -1.146  0.8620
## 
## P value adjustment: tukey method for comparing a family of 6 estimates 
## Tests are performed on the log scale 
## [1] 1.033333
##  contrast             ratio     SE    df null t.ratio p.value
##  bio / ellipse        1.015 0.0764 24277    1   0.201  1.0000
##  bio / none           1.216 0.0754 24277    1   3.161  0.0196
##  bio / rand           1.010 0.0705 24277    1   0.140  1.0000
##  bio / scramb         1.018 0.0834 24277    1   0.219  0.9999
##  bio / silhouette     0.939 0.0649 24277    1  -0.904  0.9455
##  ellipse / none       1.198 0.0681 24277    1   3.178  0.0186
##  ellipse / rand       0.995 0.0655 24277    1  -0.082  1.0000
##  ellipse / scramb     1.003 0.0755 24277    1   0.037  1.0000
##  ellipse / silhouette 0.925 0.0577 24277    1  -1.245  0.8147
##  none / rand          0.830 0.0578 24277    1  -2.673  0.0808
##  none / scramb        0.837 0.0572 24277    1  -2.606  0.0958
##  none / silhouette    0.772 0.0422 24277    1  -4.731  <.0001
##  rand / scramb        1.008 0.0644 24277    1   0.128  1.0000
##  rand / silhouette    0.930 0.0648 24277    1  -1.036  0.9058
##  scramb / silhouette  0.923 0.0799 24277    1  -0.928  0.9393
## 
## P value adjustment: tukey method for comparing a family of 6 estimates 
## Tests are performed on the log scale 
## [1] 1.2
##  contrast             ratio     SE    df null t.ratio p.value
##  bio / ellipse        0.958 0.0654 23554    1  -0.628  0.9890
##  bio / none           1.165 0.0655 23554    1   2.709  0.0735
##  bio / rand           0.991 0.0630 23554    1  -0.144  1.0000
##  bio / scramb         1.021 0.0784 23554    1   0.276  0.9998
##  bio / silhouette     0.902 0.0548 23554    1  -1.698  0.5331
##  ellipse / none       1.216 0.0615 23554    1   3.861  0.0016
##  ellipse / rand       1.034 0.0655 23554    1   0.531  0.9949
##  ellipse / scramb     1.066 0.0811 23554    1   0.841  0.9599
##  ellipse / silhouette 0.941 0.0454 23554    1  -1.252  0.8109
##  none / rand          0.851 0.0612 23554    1  -2.247  0.2164
##  none / scramb        0.877 0.0589 23554    1  -1.953  0.3700
##  none / silhouette    0.774 0.0375 23554    1  -5.279  <.0001
##  rand / scramb        1.031 0.0707 23554    1   0.442  0.9979
##  rand / silhouette    0.910 0.0618 23554    1  -1.385  0.7363
##  scramb / silhouette  0.883 0.0744 23554    1  -1.475  0.6805
## 
## P value adjustment: tukey method for comparing a family of 6 estimates 
## Tests are performed on the log scale 
## [1] 1.366667
##  contrast             ratio     SE    df null t.ratio p.value
##  bio / ellipse        0.901 0.0588 24429    1  -1.598  0.5998
##  bio / none           1.131 0.0624 24429    1   2.227  0.2253
##  bio / rand           0.985 0.0686 24429    1  -0.216  0.9999
##  bio / scramb         1.012 0.0778 24429    1   0.158  1.0000
##  bio / silhouette     0.865 0.0598 24429    1  -2.090  0.2923
##  ellipse / none       1.255 0.0587 24429    1   4.862  <.0001
##  ellipse / rand       1.093 0.0639 24429    1   1.528  0.6461
##  ellipse / scramb     1.124 0.0858 24429    1   1.525  0.6482
##  ellipse / silhouette 0.961 0.0419 24429    1  -0.920  0.9414
##  none / rand          0.871 0.0612 24429    1  -1.963  0.3636
##  none / scramb        0.895 0.0596 24429    1  -1.663  0.5564
##  none / silhouette    0.765 0.0420 24429    1  -4.874  <.0001
##  rand / scramb        1.028 0.0729 24429    1   0.384  0.9989
##  rand / silhouette    0.879 0.0560 24429    1  -2.031  0.3247
##  scramb / silhouette  0.855 0.0745 24429    1  -1.798  0.4668
## 
## P value adjustment: tukey method for comparing a family of 6 estimates 
## Tests are performed on the log scale 
## [1] 1.533333
##  contrast             ratio     SE    df null t.ratio p.value
##  bio / ellipse        0.896 0.0667 24435    1  -1.472  0.6825
##  bio / none           1.143 0.0680 24435    1   2.245  0.2175
##  bio / rand           0.993 0.0678 24435    1  -0.096  1.0000
##  bio / scramb         1.027 0.0833 24435    1   0.333  0.9995
##  bio / silhouette     0.862 0.0638 24435    1  -2.012  0.3353
##  ellipse / none       1.275 0.0604 24435    1   5.134  <.0001
##  ellipse / rand       1.108 0.0620 24435    1   1.842  0.4388
##  ellipse / scramb     1.146 0.0867 24435    1   1.805  0.4626
##  ellipse / silhouette 0.961 0.0388 24435    1  -0.980  0.9245
##  none / rand          0.869 0.0558 24435    1  -2.182  0.2460
##  none / scramb        0.899 0.0558 24435    1  -1.717  0.5204
##  none / silhouette    0.754 0.0412 24435    1  -5.165  <.0001
##  rand / scramb        1.034 0.0743 24435    1   0.466  0.9973
##  rand / silhouette    0.867 0.0526 24435    1  -2.351  0.1739
##  scramb / silhouette  0.839 0.0668 24435    1  -2.209  0.2335
## 
## P value adjustment: tukey method for comparing a family of 6 estimates 
## Tests are performed on the log scale 
## [1] 1.7
##  contrast             ratio     SE    df null t.ratio p.value
##  bio / ellipse        0.888 0.0671 23655    1  -1.575  0.6154
##  bio / none           1.134 0.0656 23655    1   2.166  0.2538
##  bio / rand           1.006 0.0639 23655    1   0.090  1.0000
##  bio / scramb         1.033 0.0793 23655    1   0.425  0.9982
##  bio / silhouette     0.856 0.0670 23655    1  -1.981  0.3534
##  ellipse / none       1.277 0.0671 23655    1   4.651  <.0001
##  ellipse / rand       1.133 0.0615 23655    1   2.296  0.1955
##  ellipse / scramb     1.164 0.0826 23655    1   2.137  0.2684
##  ellipse / silhouette 0.965 0.0431 23655    1  -0.804  0.9669
##  none / rand          0.887 0.0502 23655    1  -2.115  0.2794
##  none / scramb        0.911 0.0512 23655    1  -1.650  0.5651
##  none / silhouette    0.755 0.0397 23655    1  -5.333  <.0001
##  rand / scramb        1.027 0.0680 23655    1   0.407  0.9986
##  rand / silhouette    0.852 0.0515 23655    1  -2.659  0.0838
##  scramb / silhouette  0.829 0.0594 23655    1  -2.620  0.0926
## 
## P value adjustment: tukey method for comparing a family of 6 estimates 
## Tests are performed on the log scale 
## [1] 1.866667
##  contrast             ratio     SE    df null t.ratio p.value
##  bio / ellipse        0.879 0.0728 24444    1  -1.558  0.6261
##  bio / none           1.141 0.0653 24444    1   2.304  0.1922
##  bio / rand           1.045 0.0704 24444    1   0.654  0.9867
##  bio / scramb         1.062 0.0842 24444    1   0.760  0.9741
##  bio / silhouette     0.875 0.0655 24444    1  -1.787  0.4741
##  ellipse / none       1.298 0.0792 24444    1   4.280  0.0003
##  ellipse / rand       1.189 0.0734 24444    1   2.807  0.0563
##  ellipse / scramb     1.209 0.0821 24444    1   2.789  0.0592
##  ellipse / silhouette 0.995 0.0463 24444    1  -0.098  1.0000
##  none / rand          0.916 0.0494 24444    1  -1.627  0.5806
##  none / scramb        0.931 0.0505 24444    1  -1.318  0.7753
##  none / silhouette    0.767 0.0395 24444    1  -5.148  <.0001
##  rand / scramb        1.016 0.0641 24444    1   0.256  0.9999
##  rand / silhouette    0.837 0.0454 24444    1  -3.279  0.0134
##  scramb / silhouette  0.824 0.0480 24444    1  -3.330  0.0112
## 
## P value adjustment: tukey method for comparing a family of 6 estimates 
## Tests are performed on the log scale 
## [1] 2.033333
##  contrast             ratio     SE    df null t.ratio p.value
##  bio / ellipse        0.875 0.0782 24483    1  -1.499  0.6651
##  bio / none           1.142 0.0736 24483    1   2.062  0.3076
##  bio / rand           1.068 0.0788 24483    1   0.886  0.9499
##  bio / scramb         1.052 0.0879 24483    1   0.608  0.9905
##  bio / silhouette     0.872 0.0760 24483    1  -1.567  0.6203
##  ellipse / none       1.306 0.0746 24483    1   4.672  <.0001
##  ellipse / rand       1.221 0.0680 24483    1   3.580  0.0046
##  ellipse / scramb     1.203 0.0730 24483    1   3.049  0.0279
##  ellipse / silhouette 0.998 0.0459 24483    1  -0.054  1.0000
##  none / rand          0.935 0.0488 24483    1  -1.289  0.7911
##  none / scramb        0.921 0.0447 24483    1  -1.691  0.5377
##  none / silhouette    0.764 0.0405 24483    1  -5.083  <.0001
##  rand / scramb        0.985 0.0622 24483    1  -0.232  0.9999
##  rand / silhouette    0.817 0.0476 24483    1  -3.471  0.0069
##  scramb / silhouette  0.829 0.0490 24483    1  -3.167  0.0192
## 
## P value adjustment: tukey method for comparing a family of 6 estimates 
## Tests are performed on the log scale 
## [1] 2.2
##  contrast             ratio     SE    df null t.ratio p.value
##  bio / ellipse        0.845 0.0735 23658    1  -1.938  0.3791
##  bio / none           1.130 0.0769 23658    1   1.790  0.4722
##  bio / rand           1.039 0.0808 23658    1   0.490  0.9965
##  bio / scramb         1.043 0.0996 23658    1   0.439  0.9979
##  bio / silhouette     0.876 0.0787 23658    1  -1.476  0.6797
##  ellipse / none       1.337 0.0739 23658    1   5.251  <.0001
##  ellipse / rand       1.230 0.0730 23658    1   3.479  0.0067
##  ellipse / scramb     1.234 0.0726 23658    1   3.580  0.0046
##  ellipse / silhouette 1.037 0.0487 23658    1   0.765  0.9732
##  none / rand          0.920 0.0497 23658    1  -1.551  0.6312
##  none / scramb        0.923 0.0491 23658    1  -1.503  0.6625
##  none / silhouette    0.775 0.0448 23658    1  -4.406  0.0002
##  rand / scramb        1.004 0.0707 23658    1   0.055  1.0000
##  rand / silhouette    0.843 0.0487 23658    1  -2.954  0.0371
##  scramb / silhouette  0.840 0.0529 23658    1  -2.771  0.0622
## 
## P value adjustment: tukey method for comparing a family of 6 estimates 
## Tests are performed on the log scale 
## [1] 2.366667
##  contrast             ratio     SE    df null t.ratio p.value
##  bio / ellipse        0.840 0.0756 24502    1  -1.938  0.3786
##  bio / none           1.113 0.0772 24502    1   1.537  0.6401
##  bio / rand           1.030 0.0789 24502    1   0.382  0.9990
##  bio / scramb         1.015 0.0924 24502    1   0.158  1.0000
##  bio / silhouette     0.872 0.0759 24502    1  -1.570  0.6184
##  ellipse / none       1.324 0.0744 24502    1   4.999  <.0001
##  ellipse / rand       1.226 0.0775 24502    1   3.222  0.0161
##  ellipse / scramb     1.208 0.0701 24502    1   3.254  0.0145
##  ellipse / silhouette 1.038 0.0632 24502    1   0.620  0.9896
##  none / rand          0.926 0.0505 24502    1  -1.417  0.7165
##  none / scramb        0.912 0.0430 24502    1  -1.955  0.3686
##  none / silhouette    0.784 0.0477 24502    1  -4.001  0.0009
##  rand / scramb        0.985 0.0626 24502    1  -0.233  0.9999
##  rand / silhouette    0.847 0.0519 24502    1  -2.705  0.0742
##  scramb / silhouette  0.860 0.0517 24502    1  -2.512  0.1206
## 
## P value adjustment: tukey method for comparing a family of 6 estimates 
## Tests are performed on the log scale 
## [1] 2.533333
##  contrast             ratio     SE    df null t.ratio p.value
##  bio / ellipse        0.831 0.0741 24525    1  -2.074  0.3011
##  bio / none           1.080 0.0749 24525    1   1.106  0.8791
##  bio / rand           0.987 0.0755 24525    1  -0.173  1.0000
##  bio / scramb         0.980 0.0834 24525    1  -0.241  0.9999
##  bio / silhouette     0.847 0.0747 24525    1  -1.880  0.4146
##  ellipse / none       1.299 0.0744 24525    1   4.571  0.0001
##  ellipse / rand       1.187 0.0760 24525    1   2.682  0.0788
##  ellipse / scramb     1.179 0.0655 24525    1   2.958  0.0366
##  ellipse / silhouette 1.019 0.0653 24525    1   0.300  0.9997
##  none / rand          0.914 0.0526 24525    1  -1.564  0.6226
##  none / scramb        0.907 0.0366 24525    1  -2.414  0.1512
##  none / silhouette    0.785 0.0503 24525    1  -3.785  0.0021
##  rand / scramb        0.993 0.0613 24525    1  -0.118  1.0000
##  rand / silhouette    0.859 0.0504 24525    1  -2.597  0.0980
##  scramb / silhouette  0.865 0.0548 24525    1  -2.291  0.1975
## 
## P value adjustment: tukey method for comparing a family of 6 estimates 
## Tests are performed on the log scale 
## [1] 2.7
##  contrast             ratio     SE    df null t.ratio p.value
##  bio / ellipse        0.865 0.0728 24563    1  -1.728  0.5134
##  bio / none           1.088 0.0731 24563    1   1.249  0.8126
##  bio / rand           1.016 0.0765 24563    1   0.208  0.9999
##  bio / scramb         0.982 0.0719 24563    1  -0.242  0.9999
##  bio / silhouette     0.864 0.0731 24563    1  -1.731  0.5112
##  ellipse / none       1.258 0.0651 24563    1   4.432  0.0001
##  ellipse / rand       1.175 0.0696 24563    1   2.720  0.0713
##  ellipse / scramb     1.136 0.0608 24563    1   2.389  0.1602
##  ellipse / silhouette 0.999 0.0631 24563    1  -0.015  1.0000
##  none / rand          0.934 0.0521 24563    1  -1.223  0.8260
##  none / scramb        0.903 0.0327 24563    1  -2.807  0.0564
##  none / silhouette    0.794 0.0526 24563    1  -3.480  0.0067
##  rand / scramb        0.967 0.0550 24563    1  -0.586  0.9920
##  rand / silhouette    0.850 0.0470 24563    1  -2.931  0.0396
##  scramb / silhouette  0.879 0.0555 24563    1  -2.041  0.3193
## 
## P value adjustment: tukey method for comparing a family of 6 estimates 
## Tests are performed on the log scale 
## [1] 2.866667
##  contrast             ratio     SE    df null t.ratio p.value
##  bio / ellipse        0.839 0.0668 24633    1  -2.202  0.2369
##  bio / none           1.061 0.0635 24633    1   0.991  0.9210
##  bio / rand           0.979 0.0640 24633    1  -0.322  0.9995
##  bio / scramb         0.958 0.0572 24633    1  -0.718  0.9799
##  bio / silhouette     0.852 0.0678 24633    1  -2.020  0.3310
##  ellipse / none       1.264 0.0649 24633    1   4.572  0.0001
##  ellipse / rand       1.167 0.0757 24633    1   2.379  0.1635
##  ellipse / scramb     1.142 0.0631 24633    1   2.399  0.1566
##  ellipse / silhouette 1.015 0.0717 24633    1   0.207  0.9999
##  none / rand          0.923 0.0551 24633    1  -1.347  0.7586
##  none / scramb        0.903 0.0317 24633    1  -2.911  0.0420
##  none / silhouette    0.802 0.0571 24633    1  -3.093  0.0243
##  rand / scramb        0.978 0.0612 24633    1  -0.348  0.9993
##  rand / silhouette    0.870 0.0526 24633    1  -2.311  0.1895
##  scramb / silhouette  0.889 0.0607 24633    1  -1.727  0.5137
## 
## P value adjustment: tukey method for comparing a family of 6 estimates 
## Tests are performed on the log scale 
## [1] 6.2
##  contrast             ratio     SE    df null t.ratio p.value
##  bio / ellipse        0.954 0.0621 23795    1  -0.717  0.9800
##  bio / none           1.042 0.0629 23795    1   0.683  0.9839
##  bio / rand           1.005 0.0843 23795    1   0.059  1.0000
##  bio / scramb         0.882 0.0682 23795    1  -1.626  0.5814
##  bio / silhouette     0.909 0.0791 23795    1  -1.093  0.8842
##  ellipse / none       1.092 0.0351 23795    1   2.734  0.0688
##  ellipse / rand       1.053 0.0760 23795    1   0.716  0.9801
##  ellipse / scramb     0.924 0.0429 23795    1  -1.701  0.5308
##  ellipse / silhouette 0.953 0.0576 23795    1  -0.801  0.9673
##  none / rand          0.964 0.0573 23795    1  -0.610  0.9903
##  none / scramb        0.846 0.0364 23795    1  -3.879  0.0015
##  none / silhouette    0.873 0.0483 23795    1  -2.461  0.1357
##  rand / scramb        0.877 0.0591 23795    1  -1.940  0.3774
##  rand / silhouette    0.905 0.0764 23795    1  -1.185  0.8440
##  scramb / silhouette  1.031 0.0764 23795    1   0.412  0.9985
## 
## P value adjustment: tukey method for comparing a family of 6 estimates 
## Tests are performed on the log scale 
## [1] 6.533333
##  contrast             ratio     SE    df null t.ratio p.value
##  bio / ellipse        0.954 0.0575 24702    1  -0.779  0.9712
##  bio / none           1.033 0.0546 24702    1   0.609  0.9904
##  bio / rand           0.971 0.0761 24702    1  -0.370  0.9991
##  bio / scramb         0.866 0.0655 24702    1  -1.905  0.3992
##  bio / silhouette     0.903 0.0797 24702    1  -1.151  0.8598
##  ellipse / none       1.082 0.0352 24702    1   2.436  0.1438
##  ellipse / rand       1.018 0.0636 24702    1   0.288  0.9997
##  ellipse / scramb     0.907 0.0439 24702    1  -2.005  0.3393
##  ellipse / silhouette 0.947 0.0582 24702    1  -0.888  0.9496
##  none / rand          0.941 0.0525 24702    1  -1.096  0.8831
##  none / scramb        0.838 0.0383 24702    1  -3.863  0.0016
##  none / silhouette    0.875 0.0519 24702    1  -2.254  0.2132
##  rand / scramb        0.891 0.0571 24702    1  -1.797  0.4679
##  rand / silhouette    0.930 0.0800 24702    1  -0.844  0.9592
##  scramb / silhouette  1.043 0.0811 24702    1   0.547  0.9942
## 
## P value adjustment: tukey method for comparing a family of 6 estimates 
## Tests are performed on the log scale
```

###### 2.2.1.1.2 Summary

```
static_results_posthoc <- static_results_posthoc[-1,]
write.csv(static_results_posthoc, paste0(tmp_path, 'displacement_staticResultsPostHoc.csv'))

kable(static_results_posthoc)
```

|  | win\_starts | win\_span | bio.ellipse | bio.none | bio.rand | bio.scramb | bio.silhouette | ellipse.none | ellipse.rand | ellipse.scramb | ellipse.silhouette | none.rand | none.scramb | none.silhouette | rand.scramb | rand.silhouette | scramb.silhouette |
| --- | --- | --- | --- | --- | --- | --- | --- | --- | --- | --- | --- | --- | --- | --- | --- | --- | --- |
| 2 | 0.0333333 | 1 | 0.9999575 | 0.0022297 | 0.9997355 | 1.0000000 | 0.8589996 | 0.0003210 | 0.9999873 | 0.9999886 | 0.8266411 | 0.0195640 | 0.0215505 | 0.0002066 | 0.9994989 | 0.9364951 | 0.8896238 |
| 3 | 0.2000000 | 1 | 0.9969836 | 0.0000822 | 0.9997157 | 0.9999991 | 0.6175199 | 0.0000457 | 0.9999997 | 0.9970441 | 0.8330855 | 0.0016615 | 0.0056983 | 0.0000014 | 0.9968903 | 0.6966777 | 0.6410534 |
| 4 | 0.3666667 | 1 | 0.9999999 | 0.0001957 | 1.0000000 | 0.9999504 | 0.9696313 | 0.0002447 | 1.0000000 | 0.9998602 | 0.9838959 | 0.0018707 | 0.0051803 | 0.0000024 | 0.9998754 | 0.9684900 | 0.9590821 |
| 5 | 0.5333333 | 1 | 1.0000000 | 0.0007788 | 0.9996014 | 0.9844658 | 0.9518432 | 0.0014519 | 0.9994057 | 0.9848201 | 0.9573530 | 0.0162269 | 0.0576496 | 0.0000032 | 0.9971991 | 0.8244931 | 0.7430547 |
| 6 | 0.7000000 | 1 | 0.9999239 | 0.0029651 | 0.9999983 | 0.9998345 | 0.9029688 | 0.0046453 | 0.9999918 | 0.9999999 | 0.8153436 | 0.0261217 | 0.0535565 | 0.0000059 | 0.9999074 | 0.8364066 | 0.8603094 |
| 7 | 0.8666667 | 1 | 0.9999987 | 0.0167525 | 0.9999727 | 0.9991520 | 0.9296339 | 0.0020400 | 0.9999989 | 0.9996185 | 0.8495370 | 0.0747851 | 0.1613866 | 0.0000707 | 0.9998388 | 0.8214696 | 0.8620013 |
| 8 | 1.0333333 | 1 | 0.9999549 | 0.0196127 | 0.9999926 | 0.9999313 | 0.9455010 | 0.0185581 | 0.9999995 | 1.0000000 | 0.8147335 | 0.0808027 | 0.0958055 | 0.0000327 | 0.9999952 | 0.9057916 | 0.9392884 |
| 9 | 1.2000000 | 1 | 0.9889902 | 0.0734804 | 0.9999914 | 0.9997860 | 0.5331372 | 0.0015805 | 0.9949250 | 0.9599096 | 0.8108915 | 0.2164305 | 0.3700428 | 0.0000016 | 0.9978816 | 0.7362780 | 0.6804912 |
| 10 | 1.3666667 | 1 | 0.5997650 | 0.2253019 | 0.9999351 | 0.9999864 | 0.2923056 | 0.0000168 | 0.6461284 | 0.6481629 | 0.9414446 | 0.3635593 | 0.5564065 | 0.0000158 | 0.9989285 | 0.3247359 | 0.4668477 |
| 11 | 1.5333333 | 1 | 0.6825453 | 0.2174533 | 0.9999989 | 0.9994646 | 0.3352689 | 0.0000038 | 0.4388280 | 0.4625648 | 0.9245463 | 0.2459957 | 0.5203714 | 0.0000031 | 0.9972714 | 0.1739313 | 0.2335414 |
| 12 | 1.7000000 | 1 | 0.6154048 | 0.2538090 | 0.9999992 | 0.9982396 | 0.3533583 | 0.0000484 | 0.1955394 | 0.2683597 | 0.9668700 | 0.2794454 | 0.5650617 | 0.0000011 | 0.9985732 | 0.0837529 | 0.0925666 |
| 13 | 1.8666667 | 1 | 0.6261407 | 0.1921711 | 0.9867205 | 0.9741076 | 0.4740692 | 0.0002709 | 0.0563420 | 0.0591674 | 0.9999987 | 0.5805800 | 0.7753131 | 0.0000035 | 0.9998505 | 0.0133581 | 0.0112466 |
| 14 | 2.0333333 | 1 | 0.6651142 | 0.3075543 | 0.9498822 | 0.9904950 | 0.6203112 | 0.0000437 | 0.0046458 | 0.0278811 | 0.9999999 | 0.7911464 | 0.5377315 | 0.0000051 | 0.9999088 | 0.0068813 | 0.0192170 |
| 15 | 2.2000000 | 1 | 0.3790526 | 0.4722305 | 0.9965410 | 0.9979452 | 0.6797013 | 0.0000019 | 0.0066951 | 0.0046356 | 0.9732500 | 0.6312221 | 0.6624763 | 0.0001534 | 0.9999999 | 0.0370744 | 0.0622426 |
| 16 | 2.3666667 | 1 | 0.3786302 | 0.6400680 | 0.9989555 | 0.9999863 | 0.6183588 | 0.0000081 | 0.0161250 | 0.0144724 | 0.9896282 | 0.7164879 | 0.3685904 | 0.0008938 | 0.9999060 | 0.0742062 | 0.1206122 |
| 17 | 2.5333333 | 1 | 0.3010874 | 0.8790746 | 0.9999787 | 0.9998903 | 0.4146234 | 0.0000711 | 0.0788397 | 0.0365669 | 0.9996760 | 0.6225948 | 0.1512049 | 0.0021312 | 0.9999969 | 0.0980135 | 0.1975488 |
| 18 | 2.7000000 | 1 | 0.5133555 | 0.8126405 | 0.9999463 | 0.9998886 | 0.5111641 | 0.0001360 | 0.0713120 | 0.1601653 | 1.0000000 | 0.8259731 | 0.0563810 | 0.0066579 | 0.9919789 | 0.0395847 | 0.3192505 |
| 19 | 2.8666667 | 1 | 0.2368869 | 0.9209524 | 0.9995442 | 0.9798533 | 0.3309871 | 0.0000706 | 0.1635144 | 0.1566497 | 0.9999482 | 0.7586495 | 0.0420055 | 0.0243120 | 0.9993276 | 0.1895107 | 0.5137318 |
| 20 | 6.2000000 | 1 | 0.9799611 | 0.9838781 | 0.9999999 | 0.5813929 | 0.8842226 | 0.0688238 | 0.9801247 | 0.5308171 | 0.9673334 | 0.9903249 | 0.0014726 | 0.1357491 | 0.3774194 | 0.8439980 | 0.9984784 |
| 21 | 6.5333333 | 1 | 0.9711829 | 0.9904338 | 0.9991012 | 0.3991659 | 0.8598175 | 0.1437736 | 0.9997365 | 0.3393267 | 0.9495668 | 0.8830938 | 0.0015706 | 0.2131548 | 0.4678757 | 0.9592314 | 0.9942057 |

###### 2.2.1.2 Moving

remaking specific dataframe

```
onlymoving <- subset(full_data, full_data$stimtype != 'none')

data <- onlymoving
```

reloading of the data and ps

```
moving <- read.csv(paste0(tmp_path, 'displacement_movingResults.csv'))

moving$adjustedp <- moving$p*sum(moving$stimtype=='bio') # create bonferroni corrected p
moving$adjustedp[static$adjustedp>1] <- 1
significant_moving <- subset(moving, moving$stimtype=='bio') #subset to have only 1 row per model

significant_moving <- subset(significant_moving, significant_moving$adjustedp < alpha)

significant_moving_wins <- significant_moving$win_starts #identify the windows that are significant
```

###### 2.2.1.2.1 Detailed

now redo the models

```
moving_results_posthoc <- data.frame(list('win_starts'=NaN,'win_span'=NaN,
                                          'bio/ellipse'=NaN, 'bio/rand'=NaN,
                                          'bio/scramb'=NaN, 'bio/silhouette'=NaN,
                                          'ellipse/rand'=NaN, 'ellipse/scramb'=NaN,
                                          'ellipse/silhouette'=NaN, 'rand/scramb'=NaN,
                                          'rand/silhouette'=NaN, 'scramb/silhouette'=NaN))

for (window_start in significant_moving_wins){
  window_end <- window_start + window_size

  window_data <- subset(data, data$time_from_start>window_start)
  window_data <- subset(window_data, window_data$time_from_start<window_end)

  full_tmb <- glmmTMB(log(midpoint_displacement) ~ stimtype + (stimtype|subj), family=gaussian, data=window_data)
  Anova(full_tmb)
  e <- emmeans(full_tmb, ~stimtype, type='response')
  prs <- as.data.frame(pairs(e))
  newrow <- c(window_start, window_size, prs$p.value[1], prs$p.value[2],
              prs$p.value[3], prs$p.value[4], prs$p.value[5], prs$p.value[6],
              prs$p.value[7], prs$p.value[8], prs$p.value[9], prs$p.value[10])
  moving_results_posthoc <- rbind(moving_results_posthoc, newrow)

  print(window_start)
  print(pairs(e))
}
```

```
## Warning in finalizeTMB(TMBStruc, obj, fit, h, data.tmb.old): Model convergence
## problem; non-positive-definite Hessian matrix. See vignette('troubleshooting')
```

```
## [1] 11.86667
##  contrast             ratio     SE    df null t.ratio p.value
##  bio / ellipse        1.378 0.1080 10454    1   4.072  0.0005
##  bio / rand           0.887 0.0472 10454    1  -2.254  0.1603
##  bio / scramb         0.984 0.0481 10454    1  -0.336  0.9972
##  bio / silhouette     0.890 0.0549 10454    1  -1.889  0.3233
##  ellipse / rand       0.644 0.0547 10454    1  -5.189  <.0001
##  ellipse / scramb     0.714 0.0616 10454    1  -3.904  0.0009
##  ellipse / silhouette 0.646 0.0512 10454    1  -5.514  <.0001
##  rand / scramb        1.109 0.0465 10454    1   2.468  0.0980
##  rand / silhouette    1.004 0.0694 10454    1   0.052  1.0000
##  scramb / silhouette  0.905 0.0673 10454    1  -1.345  0.6630
## 
## P value adjustment: tukey method for comparing a family of 5 estimates 
## Tests are performed on the log scale 
## [1] 12.03333
##  contrast             ratio     SE    df null t.ratio p.value
##  bio / ellipse        1.569 0.1370 10121    1   5.175  <.0001
##  bio / rand           0.839 0.0497 10121    1  -2.965  0.0252
##  bio / scramb         0.973 0.0501 10121    1  -0.524  0.9850
##  bio / silhouette     0.858 0.0562 10121    1  -2.343  0.1314
##  ellipse / rand       0.535 0.0494 10121    1  -6.774  <.0001
##  ellipse / scramb     0.620 0.0589 10121    1  -5.027  <.0001
##  ellipse / silhouette 0.547 0.0463 10121    1  -7.126  <.0001
##  rand / scramb        1.160 0.0554 10121    1   3.118  0.0157
##  rand / silhouette    1.023 0.0758 10121    1   0.300  0.9982
##  scramb / silhouette  0.881 0.0671 10121    1  -1.662  0.4580
## 
## P value adjustment: tukey method for comparing a family of 5 estimates 
## Tests are performed on the log scale 
## [1] 12.2
##  contrast             ratio     SE   df null t.ratio p.value
##  bio / ellipse        2.016 0.1910 9097    1   7.419  <.0001
##  bio / rand           0.783 0.0535 9097    1  -3.585  0.0031
##  bio / scramb         0.970 0.0608 9097    1  -0.479  0.9893
##  bio / silhouette     0.788 0.0563 9097    1  -3.336  0.0076
##  ellipse / rand       0.388 0.0428 9097    1  -8.584  <.0001
##  ellipse / scramb     0.481 0.0555 9097    1  -6.345  <.0001
##  ellipse / silhouette 0.391 0.0371 9097    1  -9.895  <.0001
##  rand / scramb        1.240 0.0872 9097    1   3.061  0.0188
##  rand / silhouette    1.007 0.0896 9097    1   0.076  1.0000
##  scramb / silhouette  0.812 0.0683 9097    1  -2.477  0.0960
## 
## P value adjustment: tukey method for comparing a family of 5 estimates 
## Tests are performed on the log scale 
## [1] 12.36667
##  contrast             ratio     SE   df null t.ratio p.value
##  bio / ellipse        2.560 0.2570 9009    1   9.346  <.0001
##  bio / rand           0.840 0.0673 9009    1  -2.172  0.1906
##  bio / scramb         0.997 0.0737 9009    1  -0.040  1.0000
##  bio / silhouette     0.870 0.0689 9009    1  -1.759  0.3977
##  ellipse / rand       0.328 0.0426 9009    1  -8.583  <.0001
##  ellipse / scramb     0.389 0.0516 9009    1  -7.123  <.0001
##  ellipse / silhouette 0.340 0.0342 9009    1 -10.720  <.0001
##  rand / scramb        1.186 0.0908 9009    1   2.231  0.1682
##  rand / silhouette    1.035 0.1080 9009    1   0.331  0.9974
##  scramb / silhouette  0.873 0.0813 9009    1  -1.463  0.5865
## 
## P value adjustment: tukey method for comparing a family of 5 estimates 
## Tests are performed on the log scale 
## [1] 12.53333
##  contrast             ratio     SE   df null t.ratio p.value
##  bio / ellipse        2.636 0.2630 9050    1   9.724  <.0001
##  bio / rand           0.842 0.0729 9050    1  -1.982  0.2749
##  bio / scramb         0.991 0.0774 9050    1  -0.115  1.0000
##  bio / silhouette     0.862 0.0634 9050    1  -2.017  0.2580
##  ellipse / rand       0.320 0.0431 9050    1  -8.450  <.0001
##  ellipse / scramb     0.376 0.0487 9050    1  -7.547  <.0001
##  ellipse / silhouette 0.327 0.0309 9050    1 -11.829  <.0001
##  rand / scramb        1.177 0.0991 9050    1   1.931  0.3010
##  rand / silhouette    1.023 0.1090 9050    1   0.218  0.9995
##  scramb / silhouette  0.870 0.0836 9050    1  -1.451  0.5944
## 
## P value adjustment: tukey method for comparing a family of 5 estimates 
## Tests are performed on the log scale 
## [1] 12.7
##  contrast             ratio     SE   df null t.ratio p.value
##  bio / ellipse        2.677 0.2820 8444    1   9.352  <.0001
##  bio / rand           0.851 0.0817 8444    1  -1.684  0.4441
##  bio / scramb         0.947 0.0754 8444    1  -0.688  0.9591
##  bio / silhouette     0.850 0.0688 8444    1  -2.014  0.2591
##  ellipse / rand       0.318 0.0436 8444    1  -8.350  <.0001
##  ellipse / scramb     0.354 0.0468 8444    1  -7.855  <.0001
##  ellipse / silhouette 0.317 0.0306 8444    1 -11.905  <.0001
##  rand / scramb        1.113 0.1040 8444    1   1.141  0.7848
##  rand / silhouette    0.999 0.1140 8444    1  -0.012  1.0000
##  scramb / silhouette  0.897 0.0901 8444    1  -1.078  0.8180
## 
## P value adjustment: tukey method for comparing a family of 5 estimates 
## Tests are performed on the log scale 
## [1] 12.86667
##  contrast             ratio     SE   df null t.ratio p.value
##  bio / ellipse        2.834 0.3090 8296    1   9.543  <.0001
##  bio / rand           0.864 0.0896 8296    1  -1.412  0.6198
##  bio / scramb         0.981 0.0896 8296    1  -0.212  0.9996
##  bio / silhouette     0.857 0.0806 8296    1  -1.644  0.4692
##  ellipse / rand       0.305 0.0425 8296    1  -8.521  <.0001
##  ellipse / scramb     0.346 0.0436 8296    1  -8.413  <.0001
##  ellipse / silhouette 0.302 0.0305 8296    1 -11.872  <.0001
##  rand / scramb        1.135 0.1180 8296    1   1.227  0.7356
##  rand / silhouette    0.992 0.1300 8296    1  -0.062  1.0000
##  scramb / silhouette  0.874 0.0945 8296    1  -1.250  0.7218
## 
## P value adjustment: tukey method for comparing a family of 5 estimates 
## Tests are performed on the log scale 
## [1] 13.03333
##  contrast             ratio     SE   df null t.ratio p.value
##  bio / ellipse        2.777 0.3090 7954    1   9.188  <.0001
##  bio / rand           0.841 0.0877 7954    1  -1.663  0.4573
##  bio / scramb         0.939 0.0894 7954    1  -0.656  0.9656
##  bio / silhouette     0.884 0.0833 7954    1  -1.306  0.6877
##  ellipse / rand       0.303 0.0431 7954    1  -8.389  <.0001
##  ellipse / scramb     0.338 0.0449 7954    1  -8.163  <.0001
##  ellipse / silhouette 0.318 0.0326 7954    1 -11.179  <.0001
##  rand / scramb        1.117 0.1110 7954    1   1.114  0.7992
##  rand / silhouette    1.052 0.1280 7954    1   0.415  0.9938
##  scramb / silhouette  0.941 0.0991 7954    1  -0.574  0.9788
## 
## P value adjustment: tukey method for comparing a family of 5 estimates 
## Tests are performed on the log scale 
## [1] 13.2
##  contrast             ratio     SE   df null t.ratio p.value
##  bio / ellipse        2.627 0.3190 7423    1   7.946  <.0001
##  bio / rand           0.748 0.0858 7423    1  -2.529  0.0843
##  bio / scramb         0.885 0.0923 7423    1  -1.169  0.7689
##  bio / silhouette     0.872 0.0889 7423    1  -1.341  0.6657
##  ellipse / rand       0.285 0.0423 7423    1  -8.447  <.0001
##  ellipse / scramb     0.337 0.0479 7423    1  -7.659  <.0001
##  ellipse / silhouette 0.332 0.0384 7423    1  -9.525  <.0001
##  rand / scramb        1.183 0.1250 7423    1   1.599  0.4979
##  rand / silhouette    1.166 0.1560 7423    1   1.150  0.7800
##  scramb / silhouette  0.985 0.1080 7423    1  -0.134  0.9999
## 
## P value adjustment: tukey method for comparing a family of 5 estimates 
## Tests are performed on the log scale 
## [1] 13.36667
##  contrast             ratio     SE   df null t.ratio p.value
##  bio / ellipse        2.565 0.3340 7429    1   7.240  <.0001
##  bio / rand           0.742 0.0898 7429    1  -2.467  0.0983
##  bio / scramb         0.883 0.0957 7429    1  -1.149  0.7804
##  bio / silhouette     0.862 0.0816 7429    1  -1.571  0.5163
##  ellipse / rand       0.289 0.0481 7429    1  -7.466  <.0001
##  ellipse / scramb     0.344 0.0521 7429    1  -7.051  <.0001
##  ellipse / silhouette 0.336 0.0415 7429    1  -8.821  <.0001
##  rand / scramb        1.190 0.1360 7429    1   1.518  0.5506
##  rand / silhouette    1.161 0.1590 7429    1   1.095  0.8092
##  scramb / silhouette  0.976 0.1100 7429    1  -0.215  0.9995
## 
## P value adjustment: tukey method for comparing a family of 5 estimates 
## Tests are performed on the log scale 
## [1] 13.53333
##  contrast             ratio     SE   df null t.ratio p.value
##  bio / ellipse        2.404 0.3260 7341    1   6.464  <.0001
##  bio / rand           0.703 0.0934 7341    1  -2.655  0.0610
##  bio / scramb         0.871 0.0989 7341    1  -1.218  0.7410
##  bio / silhouette     0.823 0.0811 7341    1  -1.979  0.2762
##  ellipse / rand       0.292 0.0485 7341    1  -7.408  <.0001
##  ellipse / scramb     0.362 0.0570 7341    1  -6.458  <.0001
##  ellipse / silhouette 0.342 0.0435 7341    1  -8.433  <.0001
##  rand / scramb        1.239 0.1430 7341    1   1.853  0.3431
##  rand / silhouette    1.171 0.1550 7341    1   1.189  0.7576
##  scramb / silhouette  0.945 0.1020 7341    1  -0.523  0.9851
## 
## P value adjustment: tukey method for comparing a family of 5 estimates 
## Tests are performed on the log scale 
## [1] 13.7
##  contrast             ratio     SE   df null t.ratio p.value
##  bio / ellipse        2.291 0.2950 7089    1   6.437  <.0001
##  bio / rand           0.749 0.0947 7089    1  -2.285  0.1499
##  bio / scramb         0.830 0.0918 7089    1  -1.685  0.4435
##  bio / silhouette     0.758 0.0765 7089    1  -2.747  0.0475
##  ellipse / rand       0.327 0.0549 7089    1  -6.661  <.0001
##  ellipse / scramb     0.362 0.0543 7089    1  -6.774  <.0001
##  ellipse / silhouette 0.331 0.0410 7089    1  -8.918  <.0001
##  rand / scramb        1.108 0.1160 7089    1   0.978  0.8653
##  rand / silhouette    1.012 0.1380 7089    1   0.085  1.0000
##  scramb / silhouette  0.913 0.0906 7089    1  -0.915  0.8913
## 
## P value adjustment: tukey method for comparing a family of 5 estimates 
## Tests are performed on the log scale 
## [1] 13.86667
##  contrast             ratio     SE   df null t.ratio p.value
##  bio / ellipse        2.067 0.2630 7361    1   5.714  <.0001
##  bio / rand           0.701 0.0856 7361    1  -2.909  0.0299
##  bio / scramb         0.778 0.0810 7361    1  -2.408  0.1133
##  bio / silhouette     0.691 0.0725 7361    1  -3.522  0.0039
##  ellipse / rand       0.339 0.0580 7361    1  -6.320  <.0001
##  ellipse / scramb     0.376 0.0580 7361    1  -6.342  <.0001
##  ellipse / silhouette 0.334 0.0430 7361    1  -8.518  <.0001
##  rand / scramb        1.110 0.1200 7361    1   0.963  0.8718
##  rand / silhouette    0.986 0.1230 7361    1  -0.115  1.0000
##  scramb / silhouette  0.888 0.0717 7361    1  -1.471  0.5816
## 
## P value adjustment: tukey method for comparing a family of 5 estimates 
## Tests are performed on the log scale 
## [1] 14.03333
##  contrast             ratio     SE   df null t.ratio p.value
##  bio / ellipse        1.947 0.2550 7414    1   5.085  <.0001
##  bio / rand           0.671 0.0865 7414    1  -3.095  0.0169
##  bio / scramb         0.762 0.0833 7414    1  -2.483  0.0945
##  bio / silhouette     0.629 0.0703 7414    1  -4.151  0.0003
##  ellipse / rand       0.345 0.0604 7414    1  -6.078  <.0001
##  ellipse / scramb     0.392 0.0615 7414    1  -5.969  <.0001
##  ellipse / silhouette 0.323 0.0425 7414    1  -8.595  <.0001
##  rand / scramb        1.136 0.1320 7414    1   1.103  0.8050
##  rand / silhouette    0.937 0.1210 7414    1  -0.501  0.9873
##  scramb / silhouette  0.825 0.0714 7414    1  -2.224  0.1707
## 
## P value adjustment: tukey method for comparing a family of 5 estimates 
## Tests are performed on the log scale 
## [1] 14.2
##  contrast             ratio     SE   df null t.ratio p.value
##  bio / ellipse        2.056 0.2670 7206    1   5.561  <.0001
##  bio / rand           0.708 0.0874 7206    1  -2.795  0.0415
##  bio / scramb         0.770 0.0812 7206    1  -2.482  0.0948
##  bio / silhouette     0.635 0.0653 7206    1  -4.415  0.0001
##  ellipse / rand       0.344 0.0620 7206    1  -5.919  <.0001
##  ellipse / scramb     0.374 0.0595 7206    1  -6.181  <.0001
##  ellipse / silhouette 0.309 0.0406 7206    1  -8.942  <.0001
##  rand / scramb        1.087 0.1370 7206    1   0.662  0.9644
##  rand / silhouette    0.897 0.1230 7206    1  -0.799  0.9311
##  scramb / silhouette  0.825 0.0791 7206    1  -2.008  0.2623
## 
## P value adjustment: tukey method for comparing a family of 5 estimates 
## Tests are performed on the log scale 
## [1] 14.36667
##  contrast             ratio     SE   df null t.ratio p.value
##  bio / ellipse        1.962 0.2570 7546    1   5.154  <.0001
##  bio / rand           0.635 0.0864 7546    1  -3.339  0.0075
##  bio / scramb         0.749 0.0814 7546    1  -2.659  0.0604
##  bio / silhouette     0.598 0.0669 7546    1  -4.593  <.0001
##  ellipse / rand       0.324 0.0573 7546    1  -6.377  <.0001
##  ellipse / scramb     0.382 0.0605 7546    1  -6.072  <.0001
##  ellipse / silhouette 0.305 0.0418 7546    1  -8.656  <.0001
##  rand / scramb        1.180 0.1580 7546    1   1.240  0.7277
##  rand / silhouette    0.942 0.1270 7546    1  -0.444  0.9920
##  scramb / silhouette  0.798 0.0791 7546    1  -2.275  0.1531
## 
## P value adjustment: tukey method for comparing a family of 5 estimates 
## Tests are performed on the log scale 
## [1] 14.53333
##  contrast             ratio     SE   df null t.ratio p.value
##  bio / ellipse        1.911 0.2460 7574    1   5.032  <.0001
##  bio / rand           0.592 0.0829 7574    1  -3.740  0.0017
##  bio / scramb         0.739 0.0803 7574    1  -2.783  0.0430
##  bio / silhouette     0.570 0.0653 7574    1  -4.905  <.0001
##  ellipse / rand       0.310 0.0556 7574    1  -6.525  <.0001
##  ellipse / scramb     0.387 0.0629 7574    1  -5.842  <.0001
##  ellipse / silhouette 0.298 0.0415 7574    1  -8.684  <.0001
##  rand / scramb        1.247 0.1770 7574    1   1.556  0.5261
##  rand / silhouette    0.962 0.1320 7574    1  -0.284  0.9986
##  scramb / silhouette  0.771 0.0844 7574    1  -2.377  0.1218
## 
## P value adjustment: tukey method for comparing a family of 5 estimates 
## Tests are performed on the log scale 
## [1] 14.7
##  contrast             ratio     SE   df null t.ratio p.value
##  bio / ellipse        1.777 0.2220 7330    1   4.608  <.0001
##  bio / rand           0.584 0.0812 7330    1  -3.867  0.0011
##  bio / scramb         0.749 0.0879 7330    1  -2.466  0.0987
##  bio / silhouette     0.559 0.0636 7330    1  -5.109  <.0001
##  ellipse / rand       0.329 0.0594 7330    1  -6.156  <.0001
##  ellipse / scramb     0.421 0.0705 7330    1  -5.169  <.0001
##  ellipse / silhouette 0.315 0.0445 7330    1  -8.170  <.0001
##  rand / scramb        1.281 0.1910 7330    1   1.661  0.4583
##  rand / silhouette    0.957 0.1370 7330    1  -0.309  0.9980
##  scramb / silhouette  0.747 0.0872 7330    1  -2.500  0.0906
## 
## P value adjustment: tukey method for comparing a family of 5 estimates 
## Tests are performed on the log scale 
## [1] 14.86667
##  contrast             ratio     SE   df null t.ratio p.value
##  bio / ellipse        1.725 0.2010 7572    1   4.691  <.0001
##  bio / rand           0.562 0.0848 7572    1  -3.821  0.0013
##  bio / scramb         0.773 0.0920 7572    1  -2.165  0.1932
##  bio / silhouette     0.542 0.0627 7572    1  -5.296  <.0001
##  ellipse / rand       0.326 0.0543 7572    1  -6.729  <.0001
##  ellipse / scramb     0.448 0.0648 7572    1  -5.553  <.0001
##  ellipse / silhouette 0.314 0.0401 7572    1  -9.064  <.0001
##  rand / scramb        1.375 0.2110 7572    1   2.081  0.2283
##  rand / silhouette    0.965 0.1350 7572    1  -0.255  0.9991
##  scramb / silhouette  0.702 0.0806 7572    1  -3.085  0.0174
## 
## P value adjustment: tukey method for comparing a family of 5 estimates 
## Tests are performed on the log scale 
## [1] 15.03333
##  contrast             ratio     SE   df null t.ratio p.value
##  bio / ellipse        1.724 0.1970 7572    1   4.768  <.0001
##  bio / rand           0.557 0.0908 7572    1  -3.594  0.0030
##  bio / scramb         0.792 0.0924 7572    1  -1.997  0.2676
##  bio / silhouette     0.568 0.0672 7572    1  -4.779  <.0001
##  ellipse / rand       0.323 0.0535 7572    1  -6.819  <.0001
##  ellipse / scramb     0.460 0.0667 7572    1  -5.355  <.0001
##  ellipse / silhouette 0.329 0.0418 7572    1  -8.752  <.0001
##  rand / scramb        1.424 0.2340 7572    1   2.152  0.1984
##  rand / silhouette    1.021 0.1560 7572    1   0.134  0.9999
##  scramb / silhouette  0.717 0.0866 7572    1  -2.753  0.0467
## 
## P value adjustment: tukey method for comparing a family of 5 estimates 
## Tests are performed on the log scale 
## [1] 15.2
##  contrast             ratio     SE   df null t.ratio p.value
##  bio / ellipse        1.517 0.1690 7357    1   3.738  0.0018
##  bio / rand           0.510 0.0796 7357    1  -4.315  0.0002
##  bio / scramb         0.773 0.0749 7357    1  -2.653  0.0613
##  bio / silhouette     0.555 0.0657 7357    1  -4.974  <.0001
##  ellipse / rand       0.336 0.0524 7357    1  -6.995  <.0001
##  ellipse / scramb     0.510 0.0670 7357    1  -5.125  <.0001
##  ellipse / silhouette 0.366 0.0460 7357    1  -7.989  <.0001
##  rand / scramb        1.515 0.2220 7357    1   2.837  0.0368
##  rand / silhouette    1.087 0.1540 7357    1   0.586  0.9772
##  scramb / silhouette  0.717 0.0799 7357    1  -2.986  0.0237
## 
## P value adjustment: tukey method for comparing a family of 5 estimates 
## Tests are performed on the log scale 
## [1] 15.36667
##  contrast             ratio     SE   df null t.ratio p.value
##  bio / ellipse        1.417 0.1460 7667    1   3.379  0.0066
##  bio / rand           0.498 0.0786 7667    1  -4.416  0.0001
##  bio / scramb         0.708 0.0631 7667    1  -3.881  0.0010
##  bio / silhouette     0.548 0.0615 7667    1  -5.365  <.0001
##  ellipse / rand       0.351 0.0524 7667    1  -7.009  <.0001
##  ellipse / scramb     0.500 0.0601 7667    1  -5.773  <.0001
##  ellipse / silhouette 0.387 0.0469 7667    1  -7.836  <.0001
##  rand / scramb        1.421 0.2000 7667    1   2.497  0.0913
##  rand / silhouette    1.100 0.1540 7667    1   0.678  0.9613
##  scramb / silhouette  0.774 0.0858 7667    1  -2.313  0.1408
## 
## P value adjustment: tukey method for comparing a family of 5 estimates 
## Tests are performed on the log scale 
## [1] 15.53333
##  contrast             ratio     SE   df null t.ratio p.value
##  bio / ellipse        1.424 0.1310 7704    1   3.835  0.0012
##  bio / rand           0.560 0.0777 7704    1  -4.176  0.0003
##  bio / scramb         0.734 0.0616 7704    1  -3.682  0.0022
##  bio / silhouette     0.593 0.0627 7704    1  -4.944  <.0001
##  ellipse / rand       0.394 0.0536 7704    1  -6.843  <.0001
##  ellipse / scramb     0.515 0.0561 7704    1  -6.083  <.0001
##  ellipse / silhouette 0.416 0.0497 7704    1  -7.335  <.0001
##  rand / scramb        1.310 0.1610 7704    1   2.197  0.1810
##  rand / silhouette    1.058 0.1520 7704    1   0.393  0.9950
##  scramb / silhouette  0.808 0.0896 7704    1  -1.926  0.3033
## 
## P value adjustment: tukey method for comparing a family of 5 estimates 
## Tests are performed on the log scale 
## [1] 15.7
##  contrast             ratio     SE   df null t.ratio p.value
##  bio / ellipse        1.397 0.1300 7565    1   3.595  0.0030
##  bio / rand           0.596 0.0699 7565    1  -4.413  0.0001
##  bio / scramb         0.705 0.0564 7565    1  -4.368  0.0001
##  bio / silhouette     0.635 0.0698 7565    1  -4.127  0.0004
##  ellipse / rand       0.427 0.0553 7565    1  -6.570  <.0001
##  ellipse / scramb     0.505 0.0546 7565    1  -6.319  <.0001
##  ellipse / silhouette 0.455 0.0596 7565    1  -6.010  <.0001
##  rand / scramb        1.183 0.1180 7565    1   1.688  0.4411
##  rand / silhouette    1.066 0.1660 7565    1   0.412  0.9939
##  scramb / silhouette  0.901 0.1040 7565    1  -0.904  0.8956
## 
## P value adjustment: tukey method for comparing a family of 5 estimates 
## Tests are performed on the log scale 
## [1] 15.86667
##  contrast             ratio     SE   df null t.ratio p.value
##  bio / ellipse        1.307 0.1340 7986    1   2.621  0.0666
##  bio / rand           0.596 0.0703 7986    1  -4.390  0.0001
##  bio / scramb         0.689 0.0541 7986    1  -4.743  <.0001
##  bio / silhouette     0.665 0.0732 7986    1  -3.702  0.0020
##  ellipse / rand       0.456 0.0629 7986    1  -5.696  <.0001
##  ellipse / scramb     0.527 0.0618 7986    1  -5.466  <.0001
##  ellipse / silhouette 0.509 0.0650 7986    1  -5.287  <.0001
##  rand / scramb        1.156 0.1220 7986    1   1.368  0.6481
##  rand / silhouette    1.116 0.1770 7986    1   0.691  0.9584
##  scramb / silhouette  0.966 0.1140 7986    1  -0.297  0.9983
## 
## P value adjustment: tukey method for comparing a family of 5 estimates 
## Tests are performed on the log scale 
## [1] 16.03333
##  contrast             ratio     SE   df null t.ratio p.value
##  bio / ellipse        1.324 0.1370 8049    1   2.722  0.0509
##  bio / rand           0.640 0.0712 8049    1  -4.011  0.0006
##  bio / scramb         0.703 0.0588 8049    1  -4.214  0.0002
##  bio / silhouette     0.634 0.0675 8049    1  -4.284  0.0002
##  ellipse / rand       0.483 0.0647 8049    1  -5.429  <.0001
##  ellipse / scramb     0.531 0.0602 8049    1  -5.581  <.0001
##  ellipse / silhouette 0.478 0.0605 8049    1  -5.827  <.0001
##  rand / scramb        1.098 0.1130 8049    1   0.901  0.8965
##  rand / silhouette    0.990 0.1440 8049    1  -0.072  1.0000
##  scramb / silhouette  0.902 0.0997 8049    1  -0.937  0.8824
## 
## P value adjustment: tukey method for comparing a family of 5 estimates 
## Tests are performed on the log scale 
## [1] 16.2
##  contrast             ratio     SE   df null t.ratio p.value
##  bio / ellipse        1.254 0.1310 7867    1   2.170  0.1911
##  bio / rand           0.642 0.0708 7867    1  -4.015  0.0006
##  bio / scramb         0.673 0.0607 7867    1  -4.389  0.0001
##  bio / silhouette     0.554 0.0654 7867    1  -5.005  <.0001
##  ellipse / rand       0.512 0.0666 7867    1  -5.144  <.0001
##  ellipse / scramb     0.537 0.0635 7867    1  -5.262  <.0001
##  ellipse / silhouette 0.442 0.0578 7867    1  -6.248  <.0001
##  rand / scramb        1.048 0.1040 7867    1   0.471  0.9899
##  rand / silhouette    0.863 0.1230 7867    1  -1.034  0.8399
##  scramb / silhouette  0.823 0.0940 7867    1  -1.704  0.4313
## 
## P value adjustment: tukey method for comparing a family of 5 estimates 
## Tests are performed on the log scale 
## [1] 16.36667
##  contrast             ratio     SE   df null t.ratio p.value
##  bio / ellipse        1.299 0.1460 8227    1   2.336  0.1338
##  bio / rand           0.693 0.0719 8227    1  -3.532  0.0038
##  bio / scramb         0.734 0.0665 8227    1  -3.413  0.0058
##  bio / silhouette     0.585 0.0726 8227    1  -4.317  0.0002
##  ellipse / rand       0.534 0.0702 8227    1  -4.772  <.0001
##  ellipse / scramb     0.565 0.0686 8227    1  -4.701  <.0001
##  ellipse / silhouette 0.451 0.0595 8227    1  -6.036  <.0001
##  rand / scramb        1.059 0.1060 8227    1   0.571  0.9792
##  rand / silhouette    0.844 0.1100 8227    1  -1.301  0.6907
##  scramb / silhouette  0.797 0.0936 8227    1  -1.930  0.3014
## 
## P value adjustment: tukey method for comparing a family of 5 estimates 
## Tests are performed on the log scale 
## [1] 16.53333
##  contrast             ratio     SE   df null t.ratio p.value
##  bio / ellipse        1.305 0.1520 8314    1   2.287  0.1491
##  bio / rand           0.695 0.0723 8314    1  -3.500  0.0043
##  bio / scramb         0.734 0.0672 8314    1  -3.379  0.0066
##  bio / silhouette     0.572 0.0736 8314    1  -4.339  0.0001
##  ellipse / rand       0.532 0.0726 8314    1  -4.621  <.0001
##  ellipse / scramb     0.562 0.0700 8314    1  -4.627  <.0001
##  ellipse / silhouette 0.439 0.0590 8314    1  -6.128  <.0001
##  rand / scramb        1.056 0.1120 8314    1   0.518  0.9856
##  rand / silhouette    0.824 0.1090 8314    1  -1.470  0.5819
##  scramb / silhouette  0.780 0.1020 8314    1  -1.902  0.3162
## 
## P value adjustment: tukey method for comparing a family of 5 estimates 
## Tests are performed on the log scale 
## [1] 16.7
##  contrast             ratio     SE   df null t.ratio p.value
##  bio / ellipse        1.305 0.1430 8137    1   2.424  0.1090
##  bio / rand           0.692 0.0751 8137    1  -3.388  0.0064
##  bio / scramb         0.768 0.0700 8137    1  -2.897  0.0310
##  bio / silhouette     0.564 0.0704 8137    1  -4.588  <.0001
##  ellipse / rand       0.531 0.0719 8137    1  -4.674  <.0001
##  ellipse / scramb     0.589 0.0682 8137    1  -4.572  <.0001
##  ellipse / silhouette 0.432 0.0570 8137    1  -6.361  <.0001
##  rand / scramb        1.109 0.1180 8137    1   0.978  0.8652
##  rand / silhouette    0.814 0.1020 8137    1  -1.649  0.4658
##  scramb / silhouette  0.734 0.0888 8137    1  -2.560  0.0781
## 
## P value adjustment: tukey method for comparing a family of 5 estimates 
## Tests are performed on the log scale 
## [1] 16.86667
##  contrast             ratio     SE   df null t.ratio p.value
##  bio / ellipse        1.352 0.1470 8515    1   2.771  0.0444
##  bio / rand           0.676 0.0708 8515    1  -3.734  0.0018
##  bio / scramb         0.784 0.0708 8515    1  -2.691  0.0553
##  bio / silhouette     0.577 0.0738 8515    1  -4.298  0.0002
##  ellipse / rand       0.500 0.0639 8515    1  -5.426  <.0001
##  ellipse / scramb     0.580 0.0683 8515    1  -4.624  <.0001
##  ellipse / silhouette 0.427 0.0568 8515    1  -6.395  <.0001
##  rand / scramb        1.160 0.1140 8515    1   1.513  0.5543
##  rand / silhouette    0.853 0.1050 8515    1  -1.292  0.6961
##  scramb / silhouette  0.736 0.0884 8515    1  -2.556  0.0788
## 
## P value adjustment: tukey method for comparing a family of 5 estimates 
## Tests are performed on the log scale 
## [1] 17.03333
##  contrast             ratio     SE   df null t.ratio p.value
##  bio / ellipse        1.390 0.1450 8650    1   3.162  0.0136
##  bio / rand           0.685 0.0733 8650    1  -3.535  0.0038
##  bio / scramb         0.815 0.0732 8650    1  -2.280  0.1515
##  bio / silhouette     0.608 0.0792 8650    1  -3.823  0.0013
##  ellipse / rand       0.493 0.0622 8650    1  -5.608  <.0001
##  ellipse / scramb     0.586 0.0685 8650    1  -4.569  <.0001
##  ellipse / silhouette 0.437 0.0597 8650    1  -6.058  <.0001
##  rand / scramb        1.190 0.1250 8650    1   1.659  0.4595
##  rand / silhouette    0.887 0.1100 8650    1  -0.964  0.8714
##  scramb / silhouette  0.746 0.0924 8650    1  -2.366  0.1249
## 
## P value adjustment: tukey method for comparing a family of 5 estimates 
## Tests are performed on the log scale 
## [1] 17.2
##  contrast             ratio     SE   df null t.ratio p.value
##  bio / ellipse        1.398 0.1550 8497    1   3.019  0.0214
##  bio / rand           0.716 0.0717 8497    1  -3.334  0.0077
##  bio / scramb         0.868 0.0766 8497    1  -1.599  0.4981
##  bio / silhouette     0.648 0.0861 8497    1  -3.265  0.0097
##  ellipse / rand       0.512 0.0616 8497    1  -5.565  <.0001
##  ellipse / scramb     0.621 0.0716 8497    1  -4.132  0.0004
##  ellipse / silhouette 0.463 0.0635 8497    1  -5.615  <.0001
##  rand / scramb        1.213 0.1270 8497    1   1.846  0.3469
##  rand / silhouette    0.905 0.1040 8497    1  -0.871  0.9075
##  scramb / silhouette  0.746 0.0982 8497    1  -2.226  0.1702
## 
## P value adjustment: tukey method for comparing a family of 5 estimates 
## Tests are performed on the log scale 
## [1] 17.36667
##  contrast             ratio     SE   df null t.ratio p.value
##  bio / ellipse        1.372 0.1440 8929    1   3.002  0.0225
##  bio / rand           0.723 0.0814 8929    1  -2.882  0.0324
##  bio / scramb         0.867 0.0793 8929    1  -1.563  0.5211
##  bio / silhouette     0.667 0.0797 8929    1  -3.391  0.0063
##  ellipse / rand       0.527 0.0650 8929    1  -5.194  <.0001
##  ellipse / scramb     0.632 0.0749 8929    1  -3.874  0.0010
##  ellipse / silhouette 0.486 0.0672 8929    1  -5.216  <.0001
##  rand / scramb        1.199 0.1130 8929    1   1.922  0.3056
##  rand / silhouette    0.922 0.1030 8929    1  -0.727  0.9502
##  scramb / silhouette  0.769 0.0911 8929    1  -2.216  0.1738
## 
## P value adjustment: tukey method for comparing a family of 5 estimates 
## Tests are performed on the log scale 
## [1] 17.53333
##  contrast             ratio     SE   df null t.ratio p.value
##  bio / ellipse        1.338 0.1460 9046    1   2.662  0.0599
##  bio / rand           0.723 0.0786 9046    1  -2.984  0.0239
##  bio / scramb         0.867 0.0744 9046    1  -1.659  0.4596
##  bio / silhouette     0.674 0.0784 9046    1  -3.394  0.0062
##  ellipse / rand       0.540 0.0668 9046    1  -4.976  <.0001
##  ellipse / scramb     0.648 0.0813 9046    1  -3.458  0.0050
##  ellipse / silhouette 0.504 0.0702 9046    1  -4.921  <.0001
##  rand / scramb        1.200 0.1040 9046    1   2.094  0.2225
##  rand / silhouette    0.932 0.1030 9046    1  -0.640  0.9684
##  scramb / silhouette  0.777 0.0881 9046    1  -2.227  0.1699
## 
## P value adjustment: tukey method for comparing a family of 5 estimates 
## Tests are performed on the log scale 
## [1] 17.7
##  contrast             ratio     SE   df null t.ratio p.value
##  bio / ellipse        1.330 0.1460 8840    1   2.597  0.0709
##  bio / rand           0.737 0.0849 8840    1  -2.645  0.0626
##  bio / scramb         0.887 0.0734 8840    1  -1.452  0.5939
##  bio / silhouette     0.714 0.0778 8840    1  -3.091  0.0171
##  ellipse / rand       0.555 0.0734 8840    1  -4.456  0.0001
##  ellipse / scramb     0.667 0.0852 8840    1  -3.174  0.0131
##  ellipse / silhouette 0.537 0.0733 8840    1  -4.557  0.0001
##  rand / scramb        1.202 0.1150 8840    1   1.924  0.3047
##  rand / silhouette    0.968 0.1050 8840    1  -0.297  0.9983
##  scramb / silhouette  0.805 0.0824 8840    1  -2.116  0.2132
## 
## P value adjustment: tukey method for comparing a family of 5 estimates 
## Tests are performed on the log scale 
## [1] 18.2
##  contrast             ratio     SE   df null t.ratio p.value
##  bio / ellipse        1.166 0.1270 9078    1   1.410  0.6210
##  bio / rand           0.695 0.0811 9078    1  -3.116  0.0158
##  bio / scramb         0.855 0.0716 9078    1  -1.870  0.3336
##  bio / silhouette     0.725 0.0669 9078    1  -3.484  0.0045
##  ellipse / rand       0.596 0.0908 9078    1  -3.395  0.0062
##  ellipse / scramb     0.733 0.0999 9078    1  -2.277  0.1524
##  ellipse / silhouette 0.622 0.0752 9078    1  -3.927  0.0008
##  rand / scramb        1.230 0.1170 9078    1   2.173  0.1898
##  rand / silhouette    1.043 0.1410 9078    1   0.311  0.9980
##  scramb / silhouette  0.848 0.0949 9078    1  -1.473  0.5801
## 
## P value adjustment: tukey method for comparing a family of 5 estimates 
## Tests are performed on the log scale 
## [1] 18.36667
##  contrast             ratio     SE   df null t.ratio p.value
##  bio / ellipse        1.146 0.1150 9515    1   1.359  0.6538
##  bio / rand           0.681 0.0779 9515    1  -3.359  0.0070
##  bio / scramb         0.813 0.0616 9515    1  -2.734  0.0492
##  bio / silhouette     0.750 0.0742 9515    1  -2.903  0.0304
##  ellipse / rand       0.594 0.0949 9515    1  -3.260  0.0098
##  ellipse / scramb     0.709 0.0906 9515    1  -2.687  0.0559
##  ellipse / silhouette 0.655 0.0703 9515    1  -3.944  0.0008
##  rand / scramb        1.194 0.1140 9515    1   1.852  0.3437
##  rand / silhouette    1.102 0.1700 9515    1   0.630  0.9702
##  scramb / silhouette  0.923 0.1120 9515    1  -0.661  0.9646
## 
## P value adjustment: tukey method for comparing a family of 5 estimates 
## Tests are performed on the log scale 
## [1] 18.7
##  contrast             ratio     SE   df null t.ratio p.value
##  bio / ellipse        1.167 0.1160 9273    1   1.563  0.5214
##  bio / rand           0.726 0.0767 9273    1  -3.025  0.0210
##  bio / scramb         0.794 0.0597 9273    1  -3.072  0.0181
##  bio / silhouette     0.828 0.0807 9273    1  -1.931  0.3006
##  ellipse / rand       0.622 0.0939 9273    1  -3.144  0.0144
##  ellipse / scramb     0.680 0.0888 9273    1  -2.954  0.0261
##  ellipse / silhouette 0.710 0.0730 9273    1  -3.335  0.0076
##  rand / scramb        1.093 0.0980 9273    1   0.988  0.8609
##  rand / silhouette    1.140 0.1760 9273    1   0.852  0.9141
##  scramb / silhouette  1.044 0.1390 9273    1   0.322  0.9977
## 
## P value adjustment: tukey method for comparing a family of 5 estimates 
## Tests are performed on the log scale 
## [1] 20.03333
##  contrast             ratio     SE    df null t.ratio p.value
##  bio / ellipse        1.164 0.0963 10174    1   1.840  0.3504
##  bio / rand           0.798 0.0627 10174    1  -2.870  0.0335
##  bio / scramb         0.798 0.0543 10174    1  -3.319  0.0081
##  bio / silhouette     0.853 0.0712 10174    1  -1.902  0.3164
##  ellipse / rand       0.685 0.0731 10174    1  -3.544  0.0036
##  ellipse / scramb     0.685 0.0719 10174    1  -3.604  0.0029
##  ellipse / silhouette 0.733 0.0804 10174    1  -2.832  0.0374
##  rand / scramb        1.000 0.0961 10174    1  -0.001  1.0000
##  rand / silhouette    1.069 0.1150 10174    1   0.626  0.9709
##  scramb / silhouette  1.069 0.1120 10174    1   0.641  0.9684
## 
## P value adjustment: tukey method for comparing a family of 5 estimates 
## Tests are performed on the log scale 
## [1] 20.2
##  contrast             ratio     SE   df null t.ratio p.value
##  bio / ellipse        1.105 0.0873 9955    1   1.264  0.7133
##  bio / rand           0.768 0.0543 9955    1  -3.728  0.0018
##  bio / scramb         0.777 0.0511 9955    1  -3.832  0.0012
##  bio / silhouette     0.860 0.0645 9955    1  -2.010  0.2609
##  ellipse / rand       0.695 0.0681 9955    1  -3.712  0.0019
##  ellipse / scramb     0.703 0.0657 9955    1  -3.768  0.0016
##  ellipse / silhouette 0.778 0.0747 9955    1  -2.613  0.0680
##  rand / scramb        1.011 0.0925 9955    1   0.124  0.9999
##  rand / silhouette    1.119 0.1130 9955    1   1.111  0.8008
##  scramb / silhouette  1.106 0.0930 9955    1   1.204  0.7490
## 
## P value adjustment: tukey method for comparing a family of 5 estimates 
## Tests are performed on the log scale 
## [1] 20.36667
##  contrast             ratio     SE    df null t.ratio p.value
##  bio / ellipse        1.070 0.0854 10415    1   0.848  0.9154
##  bio / rand           0.792 0.0516 10415    1  -3.573  0.0033
##  bio / scramb         0.806 0.0571 10415    1  -3.048  0.0196
##  bio / silhouette     0.844 0.0603 10415    1  -2.377  0.1215
##  ellipse / rand       0.740 0.0672 10415    1  -3.313  0.0082
##  ellipse / scramb     0.753 0.0680 10415    1  -3.139  0.0147
##  ellipse / silhouette 0.788 0.0637 10415    1  -2.945  0.0268
##  rand / scramb        1.017 0.0912 10415    1   0.190  0.9997
##  rand / silhouette    1.065 0.0955 10415    1   0.699  0.9568
##  scramb / silhouette  1.047 0.0962 10415    1   0.497  0.9876
## 
## P value adjustment: tukey method for comparing a family of 5 estimates 
## Tests are performed on the log scale 
## [1] 20.53333
##  contrast             ratio     SE    df null t.ratio p.value
##  bio / ellipse        1.095 0.0863 10472    1   1.146  0.7821
##  bio / rand           0.807 0.0587 10472    1  -2.944  0.0269
##  bio / scramb         0.863 0.0555 10472    1  -2.293  0.1471
##  bio / silhouette     0.845 0.0597 10472    1  -2.386  0.1192
##  ellipse / rand       0.738 0.0638 10472    1  -3.519  0.0040
##  ellipse / scramb     0.788 0.0731 10472    1  -2.564  0.0771
##  ellipse / silhouette 0.772 0.0585 10472    1  -3.419  0.0057
##  rand / scramb        1.069 0.0979 10472    1   0.725  0.9508
##  rand / silhouette    1.046 0.0913 10472    1   0.521  0.9853
##  scramb / silhouette  0.979 0.0914 10472    1  -0.225  0.9994
## 
## P value adjustment: tukey method for comparing a family of 5 estimates 
## Tests are performed on the log scale 
## [1] 20.7
##  contrast             ratio     SE    df null t.ratio p.value
##  bio / ellipse        1.095 0.0886 10224    1   1.119  0.7964
##  bio / rand           0.834 0.0674 10224    1  -2.242  0.1643
##  bio / scramb         0.867 0.0529 10224    1  -2.346  0.1306
##  bio / silhouette     0.810 0.0632 10224    1  -2.697  0.0544
##  ellipse / rand       0.762 0.0658 10224    1  -3.147  0.0143
##  ellipse / scramb     0.791 0.0702 10224    1  -2.637  0.0638
##  ellipse / silhouette 0.740 0.0532 10224    1  -4.184  0.0003
##  rand / scramb        1.039 0.0970 10224    1   0.406  0.9943
##  rand / silhouette    0.971 0.0923 10224    1  -0.306  0.9981
##  scramb / silhouette  0.935 0.0883 10224    1  -0.710  0.9543
## 
## P value adjustment: tukey method for comparing a family of 5 estimates 
## Tests are performed on the log scale 
## [1] 20.86667
##  contrast             ratio     SE    df null t.ratio p.value
##  bio / ellipse        1.090 0.0822 10657    1   1.140  0.7851
##  bio / rand           0.850 0.0629 10657    1  -2.201  0.1793
##  bio / scramb         0.925 0.0536 10657    1  -1.350  0.6595
##  bio / silhouette     0.778 0.0675 10657    1  -2.897  0.0309
##  ellipse / rand       0.780 0.0663 10657    1  -2.928  0.0282
##  ellipse / scramb     0.848 0.0729 10657    1  -1.912  0.3108
##  ellipse / silhouette 0.714 0.0603 10657    1  -3.996  0.0006
##  rand / scramb        1.088 0.1020 10657    1   0.900  0.8971
##  rand / silhouette    0.915 0.0985 10657    1  -0.823  0.9235
##  scramb / silhouette  0.841 0.0899 10657    1  -1.621  0.4840
## 
## P value adjustment: tukey method for comparing a family of 5 estimates 
## Tests are performed on the log scale 
## [1] 21.2
##  contrast             ratio     SE    df null t.ratio p.value
##  bio / ellipse        1.094 0.0858 10419    1   1.140  0.7852
##  bio / rand           0.855 0.0535 10419    1  -2.508  0.0890
##  bio / scramb         0.980 0.0613 10419    1  -0.317  0.9978
##  bio / silhouette     0.816 0.0606 10419    1  -2.737  0.0487
##  ellipse / rand       0.782 0.0677 10419    1  -2.845  0.0360
##  ellipse / scramb     0.896 0.0698 10419    1  -1.404  0.6251
##  ellipse / silhouette 0.746 0.0607 10419    1  -3.599  0.0030
##  rand / scramb        1.147 0.1010 10419    1   1.559  0.5237
##  rand / silhouette    0.955 0.0930 10419    1  -0.474  0.9897
##  scramb / silhouette  0.832 0.0725 10419    1  -2.105  0.2181
## 
## P value adjustment: tukey method for comparing a family of 5 estimates 
## Tests are performed on the log scale 
## [1] 21.36667
##  contrast             ratio     SE    df null t.ratio p.value
##  bio / ellipse        1.086 0.0825 10824    1   1.079  0.8174
##  bio / rand           0.860 0.0537 10824    1  -2.414  0.1117
##  bio / scramb         0.959 0.0631 10824    1  -0.642  0.9682
##  bio / silhouette     0.809 0.0531 10824    1  -3.226  0.0110
##  ellipse / rand       0.792 0.0633 10824    1  -2.911  0.0297
##  ellipse / scramb     0.883 0.0645 10824    1  -1.702  0.4330
##  ellipse / silhouette 0.745 0.0576 10824    1  -3.799  0.0014
##  rand / scramb        1.114 0.0929 10824    1   1.300  0.6913
##  rand / silhouette    0.941 0.0888 10824    1  -0.648  0.9671
##  scramb / silhouette  0.844 0.0748 10824    1  -1.912  0.3107
## 
## P value adjustment: tukey method for comparing a family of 5 estimates 
## Tests are performed on the log scale 
## [1] 21.53333
##  contrast             ratio     SE    df null t.ratio p.value
##  bio / ellipse        1.084 0.0853 10841    1   1.030  0.8418
##  bio / rand           0.853 0.0528 10841    1  -2.568  0.0765
##  bio / scramb         0.967 0.0714 10841    1  -0.461  0.9907
##  bio / silhouette     0.818 0.0540 10841    1  -3.037  0.0203
##  ellipse / rand       0.787 0.0653 10841    1  -2.890  0.0316
##  ellipse / scramb     0.891 0.0672 10841    1  -1.525  0.5460
##  ellipse / silhouette 0.755 0.0644 10841    1  -3.298  0.0087
##  rand / scramb        1.133 0.0927 10841    1   1.528  0.5439
##  rand / silhouette    0.959 0.0948 10841    1  -0.419  0.9936
##  scramb / silhouette  0.847 0.0795 10841    1  -1.772  0.3902
## 
## P value adjustment: tukey method for comparing a family of 5 estimates 
## Tests are performed on the log scale 
## [1] 21.7
##  contrast             ratio     SE    df null t.ratio p.value
##  bio / ellipse        1.064 0.0894 10492    1   0.741  0.9469
##  bio / rand           0.802 0.0532 10492    1  -3.326  0.0079
##  bio / scramb         0.957 0.0658 10492    1  -0.646  0.9674
##  bio / silhouette     0.814 0.0584 10492    1  -2.871  0.0334
##  ellipse / rand       0.754 0.0633 10492    1  -3.366  0.0069
##  ellipse / scramb     0.899 0.0697 10492    1  -1.376  0.6431
##  ellipse / silhouette 0.765 0.0766 10492    1  -2.676  0.0576
##  rand / scramb        1.193 0.0848 10492    1   2.480  0.0953
##  rand / silhouette    1.015 0.1060 10492    1   0.142  0.9999
##  scramb / silhouette  0.851 0.0805 10492    1  -1.706  0.4303
## 
## P value adjustment: tukey method for comparing a family of 5 estimates 
## Tests are performed on the log scale 
## [1] 21.86667
##  contrast             ratio     SE    df null t.ratio p.value
##  bio / ellipse        1.047 0.0834 10834    1   0.581  0.9779
##  bio / rand           0.788 0.0523 10834    1  -3.594  0.0030
##  bio / scramb         0.912 0.0562 10834    1  -1.503  0.5607
##  bio / silhouette     0.786 0.0683 10834    1  -2.776  0.0439
##  ellipse / rand       0.752 0.0597 10834    1  -3.587  0.0031
##  ellipse / scramb     0.870 0.0683 10834    1  -1.772  0.3903
##  ellipse / silhouette 0.750 0.0835 10834    1  -2.582  0.0738
##  rand / scramb        1.157 0.0767 10834    1   2.202  0.1788
##  rand / silhouette    0.997 0.1210 10834    1  -0.021  1.0000
##  scramb / silhouette  0.862 0.0902 10834    1  -1.419  0.6152
## 
## P value adjustment: tukey method for comparing a family of 5 estimates 
## Tests are performed on the log scale 
## [1] 22.03333
##  contrast             ratio     SE    df null t.ratio p.value
##  bio / ellipse        1.076 0.0865 10866    1   0.907  0.8945
##  bio / rand           0.778 0.0525 10866    1  -3.721  0.0019
##  bio / scramb         0.888 0.0586 10866    1  -1.801  0.3729
##  bio / silhouette     0.794 0.0641 10866    1  -2.862  0.0343
##  ellipse / rand       0.723 0.0583 10866    1  -4.019  0.0006
##  ellipse / scramb     0.826 0.0719 10866    1  -2.202  0.1788
##  ellipse / silhouette 0.738 0.0784 10866    1  -2.861  0.0344
##  rand / scramb        1.141 0.0775 10866    1   1.947  0.2926
##  rand / silhouette    1.020 0.1140 10866    1   0.178  0.9998
##  scramb / silhouette  0.894 0.0899 10866    1  -1.116  0.7980
## 
## P value adjustment: tukey method for comparing a family of 5 estimates 
## Tests are performed on the log scale 
## [1] 22.2
##  contrast             ratio     SE    df null t.ratio p.value
##  bio / ellipse        1.105 0.0752 10523    1   1.462  0.5874
##  bio / rand           0.787 0.0512 10523    1  -3.681  0.0022
##  bio / scramb         0.861 0.0608 10523    1  -2.122  0.2109
##  bio / silhouette     0.794 0.0787 10523    1  -2.323  0.1376
##  ellipse / rand       0.713 0.0476 10523    1  -5.070  <.0001
##  ellipse / scramb     0.779 0.0610 10523    1  -3.186  0.0126
##  ellipse / silhouette 0.719 0.0828 10523    1  -2.862  0.0342
##  rand / scramb        1.094 0.0697 10523    1   1.407  0.6229
##  rand / silhouette    1.009 0.1130 10523    1   0.083  1.0000
##  scramb / silhouette  0.923 0.0991 10523    1  -0.749  0.9448
## 
## P value adjustment: tukey method for comparing a family of 5 estimates 
## Tests are performed on the log scale 
## [1] 22.36667
##  contrast             ratio     SE    df null t.ratio p.value
##  bio / ellipse        1.184 0.0784 10928    1   2.557  0.0786
##  bio / rand           0.821 0.0505 10928    1  -3.200  0.0120
##  bio / scramb         0.911 0.0615 10928    1  -1.384  0.6380
##  bio / silhouette     0.845 0.0785 10928    1  -1.817  0.3634
##  ellipse / rand       0.693 0.0502 10928    1  -5.060  <.0001
##  ellipse / scramb     0.769 0.0602 10928    1  -3.356  0.0071
##  ellipse / silhouette 0.713 0.0769 10928    1  -3.137  0.0148
##  rand / scramb        1.109 0.0761 10928    1   1.505  0.5590
##  rand / silhouette    1.028 0.1010 10928    1   0.285  0.9986
##  scramb / silhouette  0.927 0.0971 10928    1  -0.720  0.9520
## 
## P value adjustment: tukey method for comparing a family of 5 estimates 
## Tests are performed on the log scale 
## [1] 22.7
##  contrast             ratio     SE    df null t.ratio p.value
##  bio / ellipse        1.189 0.0651 10644    1   3.169  0.0133
##  bio / rand           0.880 0.0448 10644    1  -2.512  0.0880
##  bio / scramb         0.851 0.0557 10644    1  -2.463  0.0993
##  bio / silhouette     0.867 0.0688 10644    1  -1.795  0.3764
##  ellipse / rand       0.740 0.0487 10644    1  -4.574  <.0001
##  ellipse / scramb     0.716 0.0579 10644    1  -4.139  0.0003
##  ellipse / silhouette 0.729 0.0659 10644    1  -3.498  0.0043
##  rand / scramb        0.967 0.0718 10644    1  -0.450  0.9916
##  rand / silhouette    0.985 0.0761 10644    1  -0.190  0.9997
##  scramb / silhouette  1.019 0.0948 10644    1   0.201  0.9996
## 
## P value adjustment: tukey method for comparing a family of 5 estimates 
## Tests are performed on the log scale 
## [1] 22.86667
##  contrast             ratio     SE    df null t.ratio p.value
##  bio / ellipse        1.218 0.0655 11157    1   3.670  0.0023
##  bio / rand           0.892 0.0413 11157    1  -2.466  0.0985
##  bio / scramb         0.897 0.0591 11157    1  -1.648  0.4663
##  bio / silhouette     0.893 0.0558 11157    1  -1.815  0.3645
##  ellipse / rand       0.732 0.0486 11157    1  -4.694  <.0001
##  ellipse / scramb     0.736 0.0588 11157    1  -3.834  0.0012
##  ellipse / silhouette 0.733 0.0596 11157    1  -3.824  0.0012
##  rand / scramb        1.006 0.0761 11157    1   0.075  1.0000
##  rand / silhouette    1.001 0.0660 11157    1   0.011  1.0000
##  scramb / silhouette  0.995 0.0890 11157    1  -0.055  1.0000
## 
## P value adjustment: tukey method for comparing a family of 5 estimates 
## Tests are performed on the log scale
```

```
## Warning in finalizeTMB(TMBStruc, obj, fit, h, data.tmb.old): Model convergence
## problem; non-positive-definite Hessian matrix. See vignette('troubleshooting')
```

```
## Warning in finalizeTMB(TMBStruc, obj, fit, h, data.tmb.old): Model convergence
## problem; false convergence (8). See vignette('troubleshooting'),
## help('diagnose')
```

```
## [1] 23.03333
##  contrast             ratio     SE    df null t.ratio p.value
##  bio / ellipse        1.177 0.0617 11227    1   3.106  0.0163
##  bio / rand           0.886 0.0306 11227    1  -3.498  0.0043
##  bio / scramb         0.913 0.0580 11227    1  -1.428  0.6096
##  bio / silhouette     0.867 0.0607 11227    1  -2.041  0.2465
##  ellipse / rand       0.753 0.0467 11227    1  -4.570  <.0001
##  ellipse / scramb     0.776 0.0563 11227    1  -3.498  0.0043
##  ellipse / silhouette 0.736 0.0615 11227    1  -3.665  0.0023
##  rand / scramb        1.031 0.0697 11227    1   0.445  0.9919
##  rand / silhouette    0.978 0.0702 11227    1  -0.308  0.9980
##  scramb / silhouette  0.949 0.0876 11227    1  -0.566  0.9799
## 
## P value adjustment: tukey method for comparing a family of 5 estimates 
## Tests are performed on the log scale 
## [1] 23.2
##  contrast             ratio     SE    df null t.ratio p.value
##  bio / ellipse        1.194 0.0756 10952    1   2.807  0.0401
##  bio / rand           0.867 0.0499 10952    1  -2.471  0.0973
##  bio / scramb         0.921 0.0598 10952    1  -1.273  0.7079
##  bio / silhouette     0.833 0.0626 10952    1  -2.425  0.1087
##  ellipse / rand       0.726 0.0463 10952    1  -5.017  <.0001
##  ellipse / scramb     0.771 0.0566 10952    1  -3.547  0.0036
##  ellipse / silhouette 0.698 0.0595 10952    1  -4.222  0.0002
##  rand / scramb        1.061 0.0863 10952    1   0.733  0.9488
##  rand / silhouette    0.961 0.0851 10952    1  -0.452  0.9914
##  scramb / silhouette  0.905 0.0940 10952    1  -0.959  0.8735
## 
## P value adjustment: tukey method for comparing a family of 5 estimates 
## Tests are performed on the log scale 
## [1] 23.36667
##  contrast             ratio     SE    df null t.ratio p.value
##  bio / ellipse        1.166 0.0700 11422    1   2.552  0.0795
##  bio / rand           0.851 0.0515 11422    1  -2.664  0.0595
##  bio / scramb         0.877 0.0641 11422    1  -1.792  0.3782
##  bio / silhouette     0.800 0.0686 11422    1  -2.607  0.0691
##  ellipse / rand       0.730 0.0408 11422    1  -5.635  <.0001
##  ellipse / scramb     0.753 0.0547 11422    1  -3.908  0.0009
##  ellipse / silhouette 0.686 0.0584 11422    1  -4.427  0.0001
##  rand / scramb        1.031 0.0797 11422    1   0.393  0.9950
##  rand / silhouette    0.940 0.0859 11422    1  -0.681  0.9606
##  scramb / silhouette  0.912 0.0967 11422    1  -0.873  0.9067
## 
## P value adjustment: tukey method for comparing a family of 5 estimates 
## Tests are performed on the log scale 
## [1] 23.53333
##  contrast             ratio     SE    df null t.ratio p.value
##  bio / ellipse        1.127 0.0735 11496    1   1.841  0.3502
##  bio / rand           0.822 0.0544 11496    1  -2.963  0.0254
##  bio / scramb         0.889 0.0665 11496    1  -1.575  0.5136
##  bio / silhouette     0.776 0.0709 11496    1  -2.772  0.0443
##  ellipse / rand       0.729 0.0425 11496    1  -5.421  <.0001
##  ellipse / scramb     0.788 0.0553 11496    1  -3.390  0.0063
##  ellipse / silhouette 0.689 0.0607 11496    1  -4.233  0.0002
##  rand / scramb        1.081 0.0871 11496    1   0.972  0.8679
##  rand / silhouette    0.944 0.0861 11496    1  -0.627  0.9707
##  scramb / silhouette  0.873 0.0932 11496    1  -1.269  0.7105
## 
## P value adjustment: tukey method for comparing a family of 5 estimates 
## Tests are performed on the log scale 
## [1] 23.7
##  contrast             ratio     SE    df null t.ratio p.value
##  bio / ellipse        1.115 0.0626 11207    1   1.933  0.2999
##  bio / rand           0.792 0.0535 11207    1  -3.456  0.0050
##  bio / scramb         0.901 0.0621 11207    1  -1.517  0.5512
##  bio / silhouette     0.763 0.0673 11207    1  -3.070  0.0183
##  ellipse / rand       0.710 0.0426 11207    1  -5.707  <.0001
##  ellipse / scramb     0.808 0.0481 11207    1  -3.578  0.0032
##  ellipse / silhouette 0.684 0.0598 11207    1  -4.337  0.0001
##  rand / scramb        1.137 0.0862 11207    1   1.698  0.4349
##  rand / silhouette    0.963 0.0864 11207    1  -0.417  0.9937
##  scramb / silhouette  0.847 0.0843 11207    1  -1.668  0.4537
## 
## P value adjustment: tukey method for comparing a family of 5 estimates 
## Tests are performed on the log scale 
## [1] 23.86667
##  contrast             ratio     SE    df null t.ratio p.value
##  bio / ellipse        1.098 0.0569 11648    1   1.800  0.3734
##  bio / rand           0.800 0.0512 11648    1  -3.484  0.0045
##  bio / scramb         0.874 0.0585 11648    1  -2.012  0.2600
##  bio / silhouette     0.757 0.0661 11648    1  -3.185  0.0126
##  ellipse / rand       0.729 0.0405 11648    1  -5.698  <.0001
##  ellipse / scramb     0.796 0.0417 11648    1  -4.353  0.0001
##  ellipse / silhouette 0.690 0.0585 11648    1  -4.382  0.0001
##  rand / scramb        1.093 0.0724 11648    1   1.337  0.6681
##  rand / silhouette    0.946 0.0795 11648    1  -0.657  0.9655
##  scramb / silhouette  0.866 0.0785 11648    1  -1.586  0.5066
## 
## P value adjustment: tukey method for comparing a family of 5 estimates 
## Tests are performed on the log scale 
## [1] 24.03333
##  contrast             ratio     SE    df null t.ratio p.value
##  bio / ellipse        1.121 0.0517 11669    1   2.473  0.0970
##  bio / rand           0.834 0.0503 11669    1  -3.005  0.0224
##  bio / scramb         0.891 0.0626 11669    1  -1.639  0.4724
##  bio / silhouette     0.792 0.0639 11669    1  -2.893  0.0313
##  ellipse / rand       0.745 0.0415 11669    1  -5.297  <.0001
##  ellipse / scramb     0.795 0.0450 11669    1  -4.053  0.0005
##  ellipse / silhouette 0.706 0.0557 11669    1  -4.405  0.0001
##  rand / scramb        1.068 0.0705 11669    1   0.997  0.8566
##  rand / silhouette    0.949 0.0789 11669    1  -0.632  0.9700
##  scramb / silhouette  0.888 0.0741 11669    1  -1.418  0.6159
## 
## P value adjustment: tukey method for comparing a family of 5 estimates 
## Tests are performed on the log scale
```

```
## Warning in finalizeTMB(TMBStruc, obj, fit, h, data.tmb.old): Model convergence
## problem; non-positive-definite Hessian matrix. See vignette('troubleshooting')
## Warning in finalizeTMB(TMBStruc, obj, fit, h, data.tmb.old): Model convergence
## problem; false convergence (8). See vignette('troubleshooting'),
## help('diagnose')
```

```
## [1] 24.2
##  contrast             ratio     SE    df null t.ratio p.value
##  bio / ellipse        1.101 0.0507 11297    1   2.101  0.2197
##  bio / rand           0.848 0.0549 11297    1  -2.546  0.0809
##  bio / scramb         0.861 0.0632 11297    1  -2.035  0.2492
##  bio / silhouette     0.801 0.0684 11297    1  -2.604  0.0697
##  ellipse / rand       0.770 0.0361 11297    1  -5.571  <.0001
##  ellipse / scramb     0.782 0.0451 11297    1  -4.268  0.0002
##  ellipse / silhouette 0.727 0.0553 11297    1  -4.196  0.0003
##  rand / scramb        1.016 0.0561 11297    1   0.279  0.9987
##  rand / silhouette    0.944 0.0705 11297    1  -0.772  0.9386
##  scramb / silhouette  0.929 0.0745 11297    1  -0.913  0.8921
## 
## P value adjustment: tukey method for comparing a family of 5 estimates 
## Tests are performed on the log scale 
## [1] 24.36667
##  contrast             ratio     SE    df null t.ratio p.value
##  bio / ellipse        1.085 0.0509 11742    1   1.749  0.4040
##  bio / rand           0.865 0.0478 11742    1  -2.632  0.0647
##  bio / scramb         0.871 0.0538 11742    1  -2.234  0.1672
##  bio / silhouette     0.807 0.0739 11742    1  -2.346  0.1307
##  ellipse / rand       0.797 0.0427 11742    1  -4.239  0.0002
##  ellipse / scramb     0.803 0.0432 11742    1  -4.082  0.0004
##  ellipse / silhouette 0.743 0.0627 11742    1  -3.517  0.0040
##  rand / scramb        1.008 0.0531 11742    1   0.145  0.9999
##  rand / silhouette    0.933 0.0766 11742    1  -0.846  0.9162
##  scramb / silhouette  0.926 0.0784 11742    1  -0.910  0.8931
## 
## P value adjustment: tukey method for comparing a family of 5 estimates 
## Tests are performed on the log scale 
## [1] 24.53333
##  contrast             ratio     SE    df null t.ratio p.value
##  bio / ellipse        1.088 0.0494 11755    1   1.864  0.3368
##  bio / rand           0.885 0.0452 11755    1  -2.395  0.1166
##  bio / scramb         0.848 0.0507 11755    1  -2.755  0.0465
##  bio / silhouette     0.816 0.0718 11755    1  -2.315  0.1401
##  ellipse / rand       0.813 0.0387 11755    1  -4.350  0.0001
##  ellipse / scramb     0.779 0.0439 11755    1  -4.425  0.0001
##  ellipse / silhouette 0.749 0.0617 11755    1  -3.506  0.0042
##  rand / scramb        0.959 0.0499 11755    1  -0.813  0.9267
##  rand / silhouette    0.922 0.0736 11755    1  -1.020  0.8461
##  scramb / silhouette  0.962 0.0843 11755    1  -0.447  0.9918
## 
## P value adjustment: tukey method for comparing a family of 5 estimates 
## Tests are performed on the log scale 
## [1] 24.86667
##  contrast             ratio     SE    df null t.ratio p.value
##  bio / ellipse        1.095 0.0468 11787    1   2.116  0.2131
##  bio / rand           0.972 0.0560 11787    1  -0.496  0.9878
##  bio / scramb         0.890 0.0588 11787    1  -1.770  0.3913
##  bio / silhouette     0.831 0.0582 11787    1  -2.641  0.0632
##  ellipse / rand       0.888 0.0432 11787    1  -2.449  0.1026
##  ellipse / scramb     0.813 0.0543 11787    1  -3.105  0.0164
##  ellipse / silhouette 0.759 0.0516 11787    1  -4.052  0.0005
##  rand / scramb        0.915 0.0512 11787    1  -1.582  0.5093
##  rand / silhouette    0.855 0.0604 11787    1  -2.213  0.1749
##  scramb / silhouette  0.934 0.0735 11787    1  -0.863  0.9103
## 
## P value adjustment: tukey method for comparing a family of 5 estimates 
## Tests are performed on the log scale
```

###### 2.2.1.2.2 Summary

```
moving_results_posthoc <- moving_results_posthoc[-1,]
write.csv(moving_results_posthoc, paste0(tmp_path, 'displacement_movingResultsPostHoc.csv'))
kable(moving_results_posthoc)
```

|  | win\_starts | win\_span | bio.ellipse | bio.rand | bio.scramb | bio.silhouette | ellipse.rand | ellipse.scramb | ellipse.silhouette | rand.scramb | rand.silhouette | scramb.silhouette |
| --- | --- | --- | --- | --- | --- | --- | --- | --- | --- | --- | --- | --- |
| 2 | 11.86667 | 1 | 0.0004507 | 0.1602757 | 0.9972450 | 0.3233482 | 0.0000021 | 0.0009029 | 0.0000004 | 0.0979641 | 0.9999983 | 0.6629877 |
| 3 | 12.03333 | 1 | 0.0000023 | 0.0252420 | 0.9849623 | 0.1313825 | 0.0000000 | 0.0000050 | 0.0000000 | 0.0157077 | 0.9982355 | 0.4580070 |
| 4 | 12.20000 | 1 | 0.0000000 | 0.0031235 | 0.9893137 | 0.0076164 | 0.0000000 | 0.0000000 | 0.0000000 | 0.0188247 | 0.9999924 | 0.0960024 |
| 5 | 12.36667 | 1 | 0.0000000 | 0.1906116 | 0.9999994 | 0.3977352 | 0.0000000 | 0.0000000 | 0.0000000 | 0.1681994 | 0.9974232 | 0.5865287 |
| 6 | 12.53333 | 1 | 0.0000000 | 0.2749006 | 0.9999607 | 0.2579831 | 0.0000000 | 0.0000000 | 0.0000000 | 0.3010095 | 0.9994958 | 0.5943551 |
| 7 | 12.70000 | 1 | 0.0000000 | 0.4441294 | 0.9590797 | 0.2590755 | 0.0000000 | 0.0000000 | 0.0000000 | 0.7848155 | 1.0000000 | 0.8180382 |
| 8 | 12.86667 | 1 | 0.0000000 | 0.6197789 | 0.9995539 | 0.4692411 | 0.0000000 | 0.0000000 | 0.0000000 | 0.7356299 | 0.9999966 | 0.7217863 |
| 9 | 13.03333 | 1 | 0.0000000 | 0.4572605 | 0.9655563 | 0.6876525 | 0.0000000 | 0.0000000 | 0.0000000 | 0.7991624 | 0.9937815 | 0.9788161 |
| 10 | 13.20000 | 1 | 0.0000000 | 0.0842962 | 0.7689260 | 0.6657268 | 0.0000000 | 0.0000000 | 0.0000000 | 0.4978631 | 0.7799748 | 0.9999283 |
| 11 | 13.36667 | 1 | 0.0000000 | 0.0983277 | 0.7803750 | 0.5162541 | 0.0000000 | 0.0000000 | 0.0000000 | 0.5506107 | 0.8092402 | 0.9995288 |
| 12 | 13.53333 | 1 | 0.0000000 | 0.0610377 | 0.7409972 | 0.2762295 | 0.0000000 | 0.0000000 | 0.0000000 | 0.3431043 | 0.7576111 | 0.9850904 |
| 13 | 13.70000 | 1 | 0.0000000 | 0.1499318 | 0.4434943 | 0.0474650 | 0.0000000 | 0.0000000 | 0.0000000 | 0.8653468 | 0.9999883 | 0.8913179 |
| 14 | 13.86667 | 1 | 0.0000001 | 0.0298569 | 0.1132964 | 0.0039434 | 0.0000000 | 0.0000000 | 0.0000000 | 0.8718186 | 0.9999607 | 0.5815917 |
| 15 | 14.03333 | 1 | 0.0000037 | 0.0168731 | 0.0944840 | 0.0003229 | 0.0000000 | 0.0000000 | 0.0000000 | 0.8050315 | 0.9872557 | 0.1707112 |
| 16 | 14.20000 | 1 | 0.0000003 | 0.0415115 | 0.0948406 | 0.0001001 | 0.0000000 | 0.0000000 | 0.0000000 | 0.9643603 | 0.9310900 | 0.2623115 |
| 17 | 14.36667 | 1 | 0.0000026 | 0.0075297 | 0.0603946 | 0.0000436 | 0.0000000 | 0.0000000 | 0.0000000 | 0.7276869 | 0.9919570 | 0.1531282 |
| 18 | 14.53333 | 1 | 0.0000049 | 0.0017339 | 0.0429696 | 0.0000095 | 0.0000000 | 0.0000001 | 0.0000000 | 0.5261422 | 0.9985764 | 0.1217998 |
| 19 | 14.70000 | 1 | 0.0000406 | 0.0010510 | 0.0986684 | 0.0000033 | 0.0000000 | 0.0000024 | 0.0000000 | 0.4583313 | 0.9980272 | 0.0906122 |
| 20 | 14.86667 | 1 | 0.0000272 | 0.0012649 | 0.1932051 | 0.0000012 | 0.0000000 | 0.0000003 | 0.0000000 | 0.2283213 | 0.9990748 | 0.0174160 |
| 21 | 15.03333 | 1 | 0.0000187 | 0.0030277 | 0.2676169 | 0.0000177 | 0.0000000 | 0.0000009 | 0.0000000 | 0.1983502 | 0.9999277 | 0.0466901 |
| 22 | 15.20000 | 1 | 0.0017501 | 0.0001575 | 0.0613149 | 0.0000066 | 0.0000000 | 0.0000030 | 0.0000000 | 0.0368181 | 0.9771566 | 0.0237240 |
| 23 | 15.36667 | 1 | 0.0065539 | 0.0000997 | 0.0009958 | 0.0000008 | 0.0000000 | 0.0000001 | 0.0000000 | 0.0912561 | 0.9612731 | 0.1407910 |
| 24 | 15.53333 | 1 | 0.0011951 | 0.0002899 | 0.0021718 | 0.0000078 | 0.0000000 | 0.0000000 | 0.0000000 | 0.1809603 | 0.9949765 | 0.3032848 |
| 25 | 15.70000 | 1 | 0.0030147 | 0.0001011 | 0.0001238 | 0.0003585 | 0.0000000 | 0.0000000 | 0.0000000 | 0.4411446 | 0.9939382 | 0.8956447 |
| 26 | 15.86667 | 1 | 0.0666016 | 0.0001121 | 0.0000212 | 0.0020090 | 0.0000001 | 0.0000005 | 0.0000013 | 0.6480557 | 0.9584150 | 0.9982993 |
| 27 | 16.03333 | 1 | 0.0509149 | 0.0005829 | 0.0002457 | 0.0001803 | 0.0000006 | 0.0000002 | 0.0000001 | 0.8965120 | 0.9999940 | 0.8823947 |
| 28 | 16.20000 | 1 | 0.1911211 | 0.0005754 | 0.0001125 | 0.0000056 | 0.0000027 | 0.0000015 | 0.0000000 | 0.9899491 | 0.8398557 | 0.4313038 |
| 29 | 16.36667 | 1 | 0.1337758 | 0.0038017 | 0.0058224 | 0.0001555 | 0.0000183 | 0.0000259 | 0.0000000 | 0.9792115 | 0.6907370 | 0.3014037 |
| 30 | 16.53333 | 1 | 0.1491106 | 0.0042635 | 0.0065658 | 0.0001408 | 0.0000381 | 0.0000371 | 0.0000000 | 0.9855776 | 0.5818671 | 0.3162311 |
| 31 | 16.70000 | 1 | 0.1090132 | 0.0063505 | 0.0309794 | 0.0000447 | 0.0000296 | 0.0000482 | 0.0000000 | 0.8652207 | 0.4657980 | 0.0780845 |
| 32 | 16.86667 | 1 | 0.0444440 | 0.0017725 | 0.0552986 | 0.0001695 | 0.0000006 | 0.0000376 | 0.0000000 | 0.5542846 | 0.6960931 | 0.0787555 |
| 33 | 17.03333 | 1 | 0.0136444 | 0.0037598 | 0.1515010 | 0.0012518 | 0.0000002 | 0.0000489 | 0.0000000 | 0.4594835 | 0.8713853 | 0.1248573 |
| 34 | 17.20000 | 1 | 0.0214038 | 0.0076789 | 0.4981287 | 0.0096804 | 0.0000003 | 0.0003500 | 0.0000002 | 0.3469060 | 0.9075168 | 0.1702199 |
| 35 | 17.36667 | 1 | 0.0225456 | 0.0323520 | 0.5211290 | 0.0062972 | 0.0000021 | 0.0010208 | 0.0000019 | 0.3055713 | 0.9502007 | 0.1738405 |
| 36 | 17.53333 | 1 | 0.0598854 | 0.0238839 | 0.4595868 | 0.0062173 | 0.0000066 | 0.0049647 | 0.0000087 | 0.2224818 | 0.9684404 | 0.1699069 |
| 37 | 17.70000 | 1 | 0.0708770 | 0.0626057 | 0.5938901 | 0.0171119 | 0.0000825 | 0.0131204 | 0.0000517 | 0.3046563 | 0.9983001 | 0.2131837 |
| 38 | 18.20000 | 1 | 0.6209805 | 0.0158132 | 0.3335532 | 0.0045114 | 0.0062022 | 0.1524251 | 0.0008259 | 0.1898429 | 0.9979721 | 0.5800634 |
| 39 | 18.36667 | 1 | 0.6538483 | 0.0070379 | 0.0491840 | 0.0303692 | 0.0098425 | 0.0559382 | 0.0007683 | 0.3436905 | 0.9702153 | 0.9645502 |
| 40 | 18.70000 | 1 | 0.5213668 | 0.0210344 | 0.0181377 | 0.3005800 | 0.0144203 | 0.0260925 | 0.0076366 | 0.8609449 | 0.9140639 | 0.9976875 |
| 41 | 20.03333 | 1 | 0.3504292 | 0.0334671 | 0.0080729 | 0.3163661 | 0.0036269 | 0.0029108 | 0.0374055 | 1.0000000 | 0.9708945 | 0.9683698 |
| 42 | 20.20000 | 1 | 0.7132680 | 0.0018133 | 0.0012081 | 0.2609155 | 0.0019321 | 0.0015567 | 0.0680358 | 0.9999461 | 0.8007677 | 0.7490339 |
| 43 | 20.36667 | 1 | 0.9154404 | 0.0032665 | 0.0195783 | 0.1215299 | 0.0082453 | 0.0146556 | 0.0268362 | 0.9997116 | 0.9567529 | 0.9876315 |
| 44 | 20.53333 | 1 | 0.7820823 | 0.0269350 | 0.1470996 | 0.1191542 | 0.0039842 | 0.0771296 | 0.0057002 | 0.9507585 | 0.9853088 | 0.9994310 |
| 45 | 20.70000 | 1 | 0.7964014 | 0.1642654 | 0.1306329 | 0.0544255 | 0.0143135 | 0.0638299 | 0.0002787 | 0.9942886 | 0.9980924 | 0.9542531 |
| 46 | 20.86667 | 1 | 0.7851412 | 0.1793094 | 0.6595472 | 0.0309415 | 0.0282411 | 0.3108404 | 0.0006212 | 0.8970553 | 0.9235265 | 0.4840357 |
| 47 | 21.20000 | 1 | 0.7852479 | 0.0889685 | 0.9978081 | 0.0487476 | 0.0360051 | 0.6251182 | 0.0029678 | 0.5237405 | 0.9897199 | 0.2180700 |
| 48 | 21.36667 | 1 | 0.8173869 | 0.1116832 | 0.9682051 | 0.0110267 | 0.0297218 | 0.4329687 | 0.0013765 | 0.6913136 | 0.9671361 | 0.3106534 |
| 49 | 21.53333 | 1 | 0.8417696 | 0.0764931 | 0.9907148 | 0.0202832 | 0.0315754 | 0.5459567 | 0.0086700 | 0.5438857 | 0.9935907 | 0.3901581 |
| 50 | 21.70000 | 1 | 0.9469302 | 0.0078689 | 0.9674065 | 0.0333971 | 0.0068591 | 0.6431400 | 0.0575954 | 0.0952753 | 0.9999080 | 0.4302517 |
| 51 | 21.86667 | 1 | 0.9778772 | 0.0030158 | 0.5606936 | 0.0438514 | 0.0030982 | 0.3903495 | 0.0737854 | 0.1787745 | 1.0000000 | 0.6151809 |
| 52 | 22.03333 | 1 | 0.8944801 | 0.0018671 | 0.3728968 | 0.0343179 | 0.0005630 | 0.1788494 | 0.0343994 | 0.2925549 | 0.9997762 | 0.7979979 |
| 53 | 22.20000 | 1 | 0.5874414 | 0.0021710 | 0.2108710 | 0.1376199 | 0.0000040 | 0.0125981 | 0.0342265 | 0.6229283 | 0.9999895 | 0.9448414 |
| 54 | 22.36667 | 1 | 0.0785969 | 0.0120396 | 0.6379694 | 0.3633912 | 0.0000042 | 0.0071048 | 0.0147805 | 0.5589918 | 0.9985609 | 0.9519891 |
| 55 | 22.70000 | 1 | 0.0133046 | 0.0879935 | 0.0992952 | 0.3763665 | 0.0000475 | 0.0003397 | 0.0042917 | 0.9915506 | 0.9997125 | 0.9996337 |
| 56 | 22.86667 | 1 | 0.0022637 | 0.0984931 | 0.4663333 | 0.3645335 | 0.0000268 | 0.0011960 | 0.0012453 | 0.9999929 | 1.0000000 | 0.9999979 |
| 57 | 23.03333 | 1 | 0.0162958 | 0.0042858 | 0.6096168 | 0.2465027 | 0.0000484 | 0.0043001 | 0.0023118 | 0.9918989 | 0.9980447 | 0.9799083 |
| 58 | 23.20000 | 1 | 0.0400984 | 0.0973047 | 0.7078799 | 0.1086992 | 0.0000053 | 0.0035958 | 0.0002364 | 0.9487847 | 0.9914236 | 0.8735240 |
| 59 | 23.36667 | 1 | 0.0795315 | 0.0595488 | 0.3781703 | 0.0691103 | 0.0000002 | 0.0008887 | 0.0000943 | 0.9949904 | 0.9606467 | 0.9067068 |
| 60 | 23.53333 | 1 | 0.3501934 | 0.0254199 | 0.5136403 | 0.0442764 | 0.0000006 | 0.0062976 | 0.0002252 | 0.8679077 | 0.9707236 | 0.7105274 |
| 61 | 23.70000 | 1 | 0.2998805 | 0.0049956 | 0.5511500 | 0.0182504 | 0.0000001 | 0.0031988 | 0.0001419 | 0.4349250 | 0.9936600 | 0.4537169 |
| 62 | 23.86667 | 1 | 0.3734053 | 0.0045079 | 0.2600319 | 0.0126125 | 0.0000001 | 0.0001318 | 0.0001155 | 0.6680821 | 0.9654688 | 0.5066364 |
| 63 | 24.03333 | 1 | 0.0969530 | 0.0223689 | 0.4724228 | 0.0313268 | 0.0000012 | 0.0004889 | 0.0001044 | 0.8566244 | 0.9699700 | 0.6159229 |
| 64 | 24.20000 | 1 | 0.2197149 | 0.0809122 | 0.2492108 | 0.0696742 | 0.0000003 | 0.0001930 | 0.0002647 | 0.9986766 | 0.9386123 | 0.8921173 |
| 65 | 24.36667 | 1 | 0.4039586 | 0.0646686 | 0.1672044 | 0.1306772 | 0.0002196 | 0.0004315 | 0.0039997 | 0.9999017 | 0.9162275 | 0.8930832 |
| 66 | 24.53333 | 1 | 0.3367603 | 0.1165526 | 0.0464859 | 0.1400513 | 0.0001341 | 0.0000949 | 0.0041733 | 0.9267131 | 0.8461005 | 0.9917864 |
| 67 | 24.86667 | 1 | 0.2131380 | 0.9878036 | 0.3912683 | 0.0632101 | 0.1026379 | 0.0163562 | 0.0004896 | 0.5092598 | 0.1749496 | 0.9103102 |

##### 2.2.2 Inter-eye distance

###### 2.2.2.1 Static

remaking specific dataframe

```
onlystatic <- subset(full_data, full_data$moving == 0)

data <- onlystatic
```

reloading of the data and ps

```
static <- read.csv(paste0(tmp_path, 'interEyeDist_staticResults.csv'))

static$adjustedp <- static$p*sum(static$stimtype=='bio') # create bonferroni corrected p
static$adjustedp[static$adjustedp>1] <- 1
significant_static <- subset(static, static$stimtype=='bio') #subset to have only 1 row per model

significant_static <- subset(significant_static, significant_static$adjustedp < alpha)

significant_static_wins <- significant_static$win_starts #identify the windows that are significant
```

###### 2.2.2.1.1 Detailed

now redo the models

```
static_results_posthoc <- data.frame(list('win_starts'=NaN,'win_span'=NaN,
                                          'bio/ellipse'=NaN, 'bio/none'=NaN,
                                          'bio/rand'=NaN, 'bio/scramb'=NaN,
                                          'bio/silhouette'=NaN, 'ellipse/none'=NaN,
                                          'ellipse/rand'=NaN, 'ellipse/scramb'=NaN,
                                          'ellipse/silhouette'=NaN, 'none/rand'=NaN,
                                          'none/scramb'=NaN, 'none/silhouette'=NaN,
                                          'rand/scramb'=NaN, 'rand/silhouette'=NaN,
                                          'scramb/silhouette'=NaN))

for (window_start in significant_static_wins){
  window_end <- window_start + window_size

  window_data <- subset(data, data$time_from_start>window_start)
  window_data <- subset(window_data, window_data$time_from_start<window_end)

  full_tmb <- glmmTMB(log(inter_eye_dist) ~ stimtype + (stimtype|subj), family=gaussian, data=window_data)
  Anova(full_tmb)
  e <- emmeans(full_tmb, ~stimtype, type='response')
  prs <- as.data.frame(pairs(e))
  newrow <- c(window_start, window_size, prs$p.value[1], prs$p.value[2],
              prs$p.value[3], prs$p.value[4], prs$p.value[5], prs$p.value[6],
              prs$p.value[7], prs$p.value[8], prs$p.value[9], prs$p.value[10],
              prs$p.value[11], prs$p.value[12], prs$p.value[13], prs$p.value[14],
              prs$p.value[15])
  static_results_posthoc <- rbind(static_results_posthoc, newrow)
  print(window_start)
  print(pairs(e))
}
```

###### 2.2.2.1.2 Summary

```
static_results_posthoc <- static_results_posthoc[-1,]
write.csv(static_results_posthoc, paste0(tmp_path, 'interEyeDist_staticResultsPostHoc.csv'))

kable(static_results_posthoc)
```

| win\_starts | win\_span | bio.ellipse | bio.none | bio.rand | bio.scramb | bio.silhouette | ellipse.none | ellipse.rand | ellipse.scramb | ellipse.silhouette | none.rand | none.scramb | none.silhouette | rand.scramb | rand.silhouette | scramb.silhouette |
| --- | --- | --- | --- | --- | --- | --- | --- | --- | --- | --- | --- | --- | --- | --- | --- | --- |

The data show a pattern that well fits in the wide spider literature.
Already at the first window (frame 0 to 30), and consequently upon
stimuli appearace, we see that the displacement value is higher for all
stimuli compared with no stimulus. This is probably due to the spider
shifting their gaze towards the stimuli as soon as they appear, while
instead when there is no stimulus the speed is maintained at a constant
exploration rate. Notably, we observe no difference between stimuli
here. For the scrambled, random and biological motion stimuli, the
displacement value drops back to the same level of no stimulus after
windows 20-50, 30-60 and 35-65 respectively. For the ellipse and
silhouette stimuli instead the effect is maintaned up to window 85-105.
There is no difference between the two, but they start resulting
different from the other three stmuli at around window 55-95. We do see
other effect in windows 190-230, but these are generally small and
isolated, I would not discuss them.

This data seems to suggest that spiders shift their gaze upon stimuli
appearance, but qickly lose interest in the point light displays. This
makes sense! no structure in the images mean nothing to scan. The
ellipse and silhouette are instead shapes, and as such maintain the
spider attention for the duration of a scanning procedure.

###### 2.2.2.2 Moving

remaking specific dataframe

```
onlymoving <- subset(full_data, full_data$stimtype != 'none')

data <- onlymoving
```

reloading of the data and ps

```
moving <- read.csv(paste0(tmp_path, 'interEyeDist_movingResults.csv'))

moving$adjustedp <- moving$p*sum(moving$stimtype=='bio') # create bonferroni corrected p
moving$adjustedp[moving$adjustedp>1] <- 1
significant_moving <- subset(moving, moving$stimtype=='bio') #subset to have only 1 row per model

significant_moving <- subset(significant_moving, significant_moving$adjustedp < alpha)

significant_moving_wins <- significant_moving$win_starts #identify the windows that are significant
```

###### 2.2.2.3 Detailed

now redo the models

```
moving_results_posthoc <- data.frame(list('win_starts'=NaN,'win_span'=NaN,
                                          'bio/ellipse'=NaN, 'bio/rand'=NaN,
                                          'bio/scramb'=NaN, 'bio/silhouette'=NaN,
                                          'ellipse/rand'=NaN, 'ellipse/scramb'=NaN,
                                          'ellipse/silhouette'=NaN, 'rand/scramb'=NaN,
                                          'rand/silhouette'=NaN, 'scramb/silhouette'=NaN))

for (window_start in significant_moving_wins){
  window_end <- window_start + window_size

  window_data <- subset(data, data$time_from_start>window_start)
  window_data <- subset(window_data, window_data$time_from_start<window_end)

  full_tmb <- glmmTMB(log(inter_eye_dist) ~ stimtype + (stimtype|subj), family=gaussian, data=window_data)
  Anova(full_tmb)
  e <- emmeans(full_tmb, ~stimtype, type='response')
  prs <- as.data.frame(pairs(e))
  newrow <- c(window_start, window_size, prs$p.value[1], prs$p.value[2],
              prs$p.value[3], prs$p.value[4], prs$p.value[5], prs$p.value[6],
              prs$p.value[7], prs$p.value[8], prs$p.value[9], prs$p.value[10])
  moving_results_posthoc <- rbind(moving_results_posthoc, newrow)

  print(window_start)
  print(pairs(e))
}
```

```
## [1] 11.86667
##  contrast             ratio     SE    df null t.ratio p.value
##  bio / ellipse        0.851 0.0406 11239    1  -3.374  0.0067
##  bio / rand           1.132 0.0324 11239    1   4.330  0.0001
##  bio / scramb         1.021 0.0282 11239    1   0.752  0.9441
##  bio / silhouette     1.020 0.0450 11239    1   0.451  0.9915
##  ellipse / rand       1.330 0.0702 11239    1   5.402  <.0001
##  ellipse / scramb     1.199 0.0656 11239    1   3.322  0.0080
##  ellipse / silhouette 1.198 0.0513 11239    1   4.222  0.0002
##  rand / scramb        0.902 0.0265 11239    1  -3.518  0.0040
##  rand / silhouette    0.901 0.0334 11239    1  -2.809  0.0399
##  scramb / silhouette  0.999 0.0428 11239    1  -0.021  1.0000
## 
## P value adjustment: tukey method for comparing a family of 5 estimates 
## Tests are performed on the log scale 
## [1] 12.03333
##  contrast             ratio     SE    df null t.ratio p.value
##  bio / ellipse        0.853 0.0415 10996    1  -3.274  0.0094
##  bio / rand           1.191 0.0422 10996    1   4.927  <.0001
##  bio / scramb         1.033 0.0308 10996    1   1.083  0.8153
##  bio / silhouette     1.016 0.0496 10996    1   0.323  0.9977
##  ellipse / rand       1.396 0.0764 10996    1   6.095  <.0001
##  ellipse / scramb     1.211 0.0653 10996    1   3.550  0.0035
##  ellipse / silhouette 1.191 0.0534 10996    1   3.898  0.0009
##  rand / scramb        0.867 0.0304 10996    1  -4.059  0.0005
##  rand / silhouette    0.853 0.0372 10996    1  -3.640  0.0025
##  scramb / silhouette  0.984 0.0443 10996    1  -0.367  0.9961
## 
## P value adjustment: tukey method for comparing a family of 5 estimates 
## Tests are performed on the log scale 
## [1] 12.2
##  contrast             ratio     SE    df null t.ratio p.value
##  bio / ellipse        0.849 0.0438 10058    1  -3.178  0.0129
##  bio / rand           1.275 0.0625 10058    1   4.943  <.0001
##  bio / scramb         1.047 0.0354 10058    1   1.361  0.6527
##  bio / silhouette     1.015 0.0569 10058    1   0.258  0.9990
##  ellipse / rand       1.502 0.0931 10058    1   6.556  <.0001
##  ellipse / scramb     1.234 0.0719 10058    1   3.604  0.0029
##  ellipse / silhouette 1.195 0.0664 10058    1   3.213  0.0115
##  rand / scramb        0.822 0.0386 10058    1  -4.186  0.0003
##  rand / silhouette    0.796 0.0436 10058    1  -4.166  0.0003
##  scramb / silhouette  0.969 0.0491 10058    1  -0.622  0.9716
## 
## P value adjustment: tukey method for comparing a family of 5 estimates 
## Tests are performed on the log scale 
## [1] 12.36667
##  contrast             ratio     SE    df null t.ratio p.value
##  bio / ellipse        0.836 0.0530 10095    1  -2.833  0.0373
##  bio / rand           1.354 0.0799 10095    1   5.135  <.0001
##  bio / scramb         1.048 0.0381 10095    1   1.298  0.6925
##  bio / silhouette     1.010 0.0573 10095    1   0.173  0.9998
##  ellipse / rand       1.620 0.1210 10095    1   6.467  <.0001
##  ellipse / scramb     1.255 0.0852 10095    1   3.341  0.0075
##  ellipse / silhouette 1.209 0.0792 10095    1   2.889  0.0316
##  rand / scramb        0.774 0.0387 10095    1  -5.122  <.0001
##  rand / silhouette    0.746 0.0423 10095    1  -5.167  <.0001
##  scramb / silhouette  0.963 0.0498 10095    1  -0.723  0.9514
## 
## P value adjustment: tukey method for comparing a family of 5 estimates 
## Tests are performed on the log scale 
## [1] 12.53333
##  contrast             ratio     SE    df null t.ratio p.value
##  bio / ellipse        0.851 0.0573 10113    1  -2.390  0.1181
##  bio / rand           1.356 0.0824 10113    1   5.008  <.0001
##  bio / scramb         1.044 0.0369 10113    1   1.231  0.7330
##  bio / silhouette     1.009 0.0576 10113    1   0.159  0.9999
##  ellipse / rand       1.592 0.1190 10113    1   6.223  <.0001
##  ellipse / scramb     1.227 0.0857 10113    1   2.923  0.0287
##  ellipse / silhouette 1.185 0.0774 10113    1   2.600  0.0704
##  rand / scramb        0.770 0.0390 10113    1  -5.150  <.0001
##  rand / silhouette    0.744 0.0409 10113    1  -5.372  <.0001
##  scramb / silhouette  0.966 0.0496 10113    1  -0.671  0.9627
## 
## P value adjustment: tukey method for comparing a family of 5 estimates 
## Tests are performed on the log scale 
## [1] 12.7
##  contrast             ratio     SE   df null t.ratio p.value
##  bio / ellipse        0.903 0.0638 9411    1  -1.444  0.5991
##  bio / rand           1.378 0.0963 9411    1   4.594  <.0001
##  bio / scramb         1.028 0.0394 9411    1   0.716  0.9529
##  bio / silhouette     0.991 0.0480 9411    1  -0.196  0.9997
##  ellipse / rand       1.526 0.1170 9411    1   5.534  <.0001
##  ellipse / scramb     1.138 0.0750 9411    1   1.965  0.2832
##  ellipse / silhouette 1.097 0.0663 9411    1   1.531  0.5423
##  rand / scramb        0.746 0.0374 9411    1  -5.855  <.0001
##  rand / silhouette    0.719 0.0422 9411    1  -5.624  <.0001
##  scramb / silhouette  0.964 0.0451 9411    1  -0.790  0.9337
## 
## P value adjustment: tukey method for comparing a family of 5 estimates 
## Tests are performed on the log scale 
## [1] 12.86667
##  contrast             ratio     SE   df null t.ratio p.value
##  bio / ellipse        0.974 0.0689 9263    1  -0.367  0.9961
##  bio / rand           1.426 0.1070 9263    1   4.716  <.0001
##  bio / scramb         1.056 0.0455 9263    1   1.268  0.7110
##  bio / silhouette     1.057 0.0557 9263    1   1.049  0.8324
##  ellipse / rand       1.464 0.1180 9263    1   4.740  <.0001
##  ellipse / scramb     1.084 0.0727 9263    1   1.201  0.7507
##  ellipse / silhouette 1.085 0.0675 9263    1   1.306  0.6874
##  rand / scramb        0.740 0.0388 9263    1  -5.734  <.0001
##  rand / silhouette    0.741 0.0486 9263    1  -4.568  <.0001
##  scramb / silhouette  1.001 0.0520 9263    1   0.013  1.0000
## 
## P value adjustment: tukey method for comparing a family of 5 estimates 
## Tests are performed on the log scale 
## [1] 13.03333
##  contrast             ratio     SE   df null t.ratio p.value
##  bio / ellipse        1.012 0.0750 8900    1   0.167  0.9998
##  bio / rand           1.420 0.1080 8900    1   4.631  <.0001
##  bio / scramb         1.070 0.0560 8900    1   1.286  0.7002
##  bio / silhouette     1.103 0.0774 8900    1   1.391  0.6335
##  ellipse / rand       1.403 0.1100 8900    1   4.299  0.0002
##  ellipse / scramb     1.057 0.0717 8900    1   0.810  0.9275
##  ellipse / silhouette 1.089 0.0721 8900    1   1.288  0.6986
##  rand / scramb        0.753 0.0364 8900    1  -5.870  <.0001
##  rand / silhouette    0.776 0.0538 8900    1  -3.649  0.0025
##  scramb / silhouette  1.031 0.0643 8900    1   0.485  0.9887
## 
## P value adjustment: tukey method for comparing a family of 5 estimates 
## Tests are performed on the log scale 
## [1] 13.2
##  contrast             ratio     SE   df null t.ratio p.value
##  bio / ellipse        1.005 0.0794 8306    1   0.065  1.0000
##  bio / rand           1.378 0.1020 8306    1   4.349  0.0001
##  bio / scramb         1.062 0.0574 8306    1   1.107  0.8031
##  bio / silhouette     1.126 0.0822 8306    1   1.624  0.4817
##  ellipse / rand       1.371 0.0988 8306    1   4.384  0.0001
##  ellipse / scramb     1.056 0.0771 8306    1   0.750  0.9447
##  ellipse / silhouette 1.120 0.0764 8306    1   1.664  0.4562
##  rand / scramb        0.770 0.0390 8306    1  -5.159  <.0001
##  rand / silhouette    0.817 0.0588 8306    1  -2.812  0.0395
##  scramb / silhouette  1.061 0.0686 8306    1   0.909  0.8936
## 
## P value adjustment: tukey method for comparing a family of 5 estimates 
## Tests are performed on the log scale 
## [1] 13.36667
##  contrast             ratio     SE   df null t.ratio p.value
##  bio / ellipse        0.992 0.0763 8314    1  -0.105  1.0000
##  bio / rand           1.344 0.0941 8314    1   4.216  0.0002
##  bio / scramb         1.076 0.0594 8314    1   1.319  0.6796
##  bio / silhouette     1.194 0.0949 8314    1   2.231  0.1682
##  ellipse / rand       1.354 0.1000 8314    1   4.107  0.0004
##  ellipse / scramb     1.084 0.0839 8314    1   1.045  0.8343
##  ellipse / silhouette 1.204 0.0862 8314    1   2.591  0.0722
##  rand / scramb        0.801 0.0413 8314    1  -4.315  0.0002
##  rand / silhouette    0.889 0.0575 8314    1  -1.822  0.3606
##  scramb / silhouette  1.110 0.0691 8314    1   1.680  0.4465
## 
## P value adjustment: tukey method for comparing a family of 5 estimates 
## Tests are performed on the log scale 
## [1] 13.53333
##  contrast             ratio     SE   df null t.ratio p.value
##  bio / ellipse        0.988 0.0726 8226    1  -0.168  0.9998
##  bio / rand           1.388 0.0971 8226    1   4.685  <.0001
##  bio / scramb         1.100 0.0608 8226    1   1.721  0.4209
##  bio / silhouette     1.264 0.1000 8226    1   2.958  0.0258
##  ellipse / rand       1.405 0.1070 8226    1   4.459  0.0001
##  ellipse / scramb     1.113 0.0900 8226    1   1.329  0.6733
##  ellipse / silhouette 1.280 0.0982 8226    1   3.213  0.0115
##  rand / scramb        0.792 0.0436 8226    1  -4.227  0.0002
##  rand / silhouette    0.911 0.0646 8226    1  -1.319  0.6794
##  scramb / silhouette  1.149 0.0723 8226    1   2.210  0.1759
## 
## P value adjustment: tukey method for comparing a family of 5 estimates 
## Tests are performed on the log scale 
## [1] 13.7
##  contrast             ratio     SE   df null t.ratio p.value
##  bio / ellipse        0.963 0.0713 7929    1  -0.513  0.9861
##  bio / rand           1.397 0.1020 7929    1   4.601  <.0001
##  bio / scramb         1.120 0.0762 7929    1   1.670  0.4528
##  bio / silhouette     1.342 0.1020 7929    1   3.874  0.0010
##  ellipse / rand       1.451 0.1050 7929    1   5.173  <.0001
##  ellipse / scramb     1.164 0.1030 7929    1   1.718  0.4228
##  ellipse / silhouette 1.394 0.1140 7929    1   4.069  0.0005
##  rand / scramb        0.802 0.0504 7929    1  -3.512  0.0041
##  rand / silhouette    0.961 0.0703 7929    1  -0.549  0.9821
##  scramb / silhouette  1.198 0.0816 7929    1   2.654  0.0612
## 
## P value adjustment: tukey method for comparing a family of 5 estimates 
## Tests are performed on the log scale 
## [1] 13.86667
##  contrast             ratio     SE   df null t.ratio p.value
##  bio / ellipse        0.901 0.0605 8218    1  -1.554  0.5274
##  bio / rand           1.303 0.0927 8218    1   3.717  0.0019
##  bio / scramb         1.090 0.0727 8218    1   1.290  0.6977
##  bio / silhouette     1.254 0.0856 8218    1   3.316  0.0081
##  ellipse / rand       1.446 0.0933 8218    1   5.716  <.0001
##  ellipse / scramb     1.210 0.0991 8218    1   2.323  0.1377
##  ellipse / silhouette 1.392 0.1150 8218    1   4.000  0.0006
##  rand / scramb        0.837 0.0473 8218    1  -3.153  0.0140
##  rand / silhouette    0.963 0.0635 8218    1  -0.578  0.9783
##  scramb / silhouette  1.151 0.0693 8218    1   2.328  0.1360
## 
## P value adjustment: tukey method for comparing a family of 5 estimates 
## Tests are performed on the log scale 
## [1] 14.03333
##  contrast             ratio     SE   df null t.ratio p.value
##  bio / ellipse        0.886 0.0602 8253    1  -1.779  0.3861
##  bio / rand           1.280 0.0964 8253    1   3.278  0.0093
##  bio / scramb         1.106 0.0706 8253    1   1.579  0.5109
##  bio / silhouette     1.200 0.0818 8253    1   2.679  0.0571
##  ellipse / rand       1.444 0.0919 8253    1   5.782  <.0001
##  ellipse / scramb     1.248 0.0938 8253    1   2.948  0.0266
##  ellipse / silhouette 1.354 0.1160 8253    1   3.536  0.0037
##  rand / scramb        0.864 0.0466 8253    1  -2.709  0.0527
##  rand / silhouette    0.938 0.0691 8253    1  -0.873  0.9069
##  scramb / silhouette  1.085 0.0595 8253    1   1.491  0.5683
## 
## P value adjustment: tukey method for comparing a family of 5 estimates 
## Tests are performed on the log scale 
## [1] 14.2
##  contrast             ratio     SE   df null t.ratio p.value
##  bio / ellipse        0.853 0.0591 8012    1  -2.291  0.1480
##  bio / rand           1.267 0.0984 8012    1   3.048  0.0196
##  bio / scramb         1.087 0.0649 8012    1   1.392  0.6327
##  bio / silhouette     1.173 0.0860 8012    1   2.171  0.1907
##  ellipse / rand       1.485 0.0849 8012    1   6.922  <.0001
##  ellipse / scramb     1.274 0.0875 8012    1   3.520  0.0040
##  ellipse / silhouette 1.374 0.1250 8012    1   3.504  0.0042
##  rand / scramb        0.858 0.0470 8012    1  -2.800  0.0409
##  rand / silhouette    0.925 0.0809 8012    1  -0.887  0.9020
##  scramb / silhouette  1.079 0.0611 8012    1   1.345  0.6628
## 
## P value adjustment: tukey method for comparing a family of 5 estimates 
## Tests are performed on the log scale 
## [1] 14.36667
##  contrast             ratio     SE   df null t.ratio p.value
##  bio / ellipse        0.848 0.0570 8361    1  -2.454  0.1013
##  bio / rand           1.257 0.1020 8361    1   2.809  0.0399
##  bio / scramb         1.080 0.0578 8361    1   1.429  0.6085
##  bio / silhouette     1.112 0.0770 8361    1   1.532  0.5418
##  ellipse / rand       1.483 0.0866 8361    1   6.738  <.0001
##  ellipse / scramb     1.273 0.0794 8361    1   3.877  0.0010
##  ellipse / silhouette 1.312 0.1160 8361    1   3.073  0.0181
##  rand / scramb        0.859 0.0556 8361    1  -2.349  0.1296
##  rand / silhouette    0.885 0.0857 8361    1  -1.265  0.7126
##  scramb / silhouette  1.030 0.0576 8361    1   0.528  0.9845
## 
## P value adjustment: tukey method for comparing a family of 5 estimates 
## Tests are performed on the log scale 
## [1] 14.53333
##  contrast             ratio     SE   df null t.ratio p.value
##  bio / ellipse        0.826 0.0538 8365    1  -2.931  0.0280
##  bio / rand           1.201 0.0939 8365    1   2.337  0.1333
##  bio / scramb         1.042 0.0502 8365    1   0.864  0.9101
##  bio / silhouette     1.065 0.0746 8365    1   0.906  0.8947
##  ellipse / rand       1.453 0.0887 8365    1   6.127  <.0001
##  ellipse / scramb     1.262 0.0796 8365    1   3.684  0.0022
##  ellipse / silhouette 1.290 0.1150 8365    1   2.842  0.0363
##  rand / scramb        0.868 0.0584 8365    1  -2.100  0.2200
##  rand / silhouette    0.887 0.0899 8365    1  -1.178  0.7640
##  scramb / silhouette  1.022 0.0657 8365    1   0.340  0.9971
## 
## P value adjustment: tukey method for comparing a family of 5 estimates 
## Tests are performed on the log scale 
## [1] 15.7
##  contrast             ratio     SE   df null t.ratio p.value
##  bio / ellipse        0.801 0.0501 8158    1  -3.556  0.0035
##  bio / rand           1.240 0.0829 8158    1   3.218  0.0113
##  bio / scramb         0.924 0.0583 8158    1  -1.246  0.7243
##  bio / silhouette     0.978 0.0682 8158    1  -0.316  0.9978
##  ellipse / rand       1.549 0.1330 8158    1   5.098  <.0001
##  ellipse / scramb     1.155 0.0875 8158    1   1.898  0.3186
##  ellipse / silhouette 1.222 0.0962 8158    1   2.545  0.0811
##  rand / scramb        0.745 0.0558 8158    1  -3.927  0.0008
##  rand / silhouette    0.789 0.0785 8158    1  -2.383  0.1200
##  scramb / silhouette  1.058 0.0983 8158    1   0.609  0.9737
## 
## P value adjustment: tukey method for comparing a family of 5 estimates 
## Tests are performed on the log scale 
## [1] 15.86667
##  contrast             ratio     SE   df null t.ratio p.value
##  bio / ellipse        0.816 0.0484 8578    1  -3.418  0.0057
##  bio / rand           1.224 0.0788 8578    1   3.140  0.0146
##  bio / scramb         0.933 0.0528 8578    1  -1.224  0.7372
##  bio / silhouette     1.006 0.0697 8578    1   0.092  1.0000
##  ellipse / rand       1.499 0.1220 8578    1   4.970  <.0001
##  ellipse / scramb     1.143 0.0794 8578    1   1.923  0.3051
##  ellipse / silhouette 1.233 0.0937 8578    1   2.752  0.0469
##  rand / scramb        0.762 0.0549 8578    1  -3.766  0.0016
##  rand / silhouette    0.822 0.0735 8578    1  -2.190  0.1835
##  scramb / silhouette  1.079 0.0978 8578    1   0.834  0.9201
## 
## P value adjustment: tukey method for comparing a family of 5 estimates 
## Tests are performed on the log scale 
## [1] 16.03333
##  contrast             ratio     SE   df null t.ratio p.value
##  bio / ellipse        0.813 0.0475 8651    1  -3.547  0.0036
##  bio / rand           1.206 0.0785 8651    1   2.871  0.0334
##  bio / scramb         0.935 0.0489 8651    1  -1.279  0.7044
##  bio / silhouette     1.003 0.0712 8651    1   0.047  1.0000
##  ellipse / rand       1.484 0.1190 8651    1   4.926  <.0001
##  ellipse / scramb     1.151 0.0737 8651    1   2.196  0.1810
##  ellipse / silhouette 1.235 0.0929 8651    1   2.804  0.0405
##  rand / scramb        0.776 0.0536 8651    1  -3.672  0.0023
##  rand / silhouette    0.832 0.0736 8651    1  -2.075  0.2309
##  scramb / silhouette  1.073 0.0954 8651    1   0.790  0.9335
## 
## P value adjustment: tukey method for comparing a family of 5 estimates 
## Tests are performed on the log scale 
## [1] 16.2
##  contrast             ratio     SE   df null t.ratio p.value
##  bio / ellipse        0.790 0.0494 8477    1  -3.762  0.0016
##  bio / rand           1.193 0.0797 8477    1   2.638  0.0637
##  bio / scramb         0.920 0.0505 8477    1  -1.517  0.5513
##  bio / silhouette     1.000 0.0703 8477    1   0.003  1.0000
##  ellipse / rand       1.509 0.1180 8477    1   5.242  <.0001
##  ellipse / scramb     1.164 0.0753 8477    1   2.347  0.1304
##  ellipse / silhouette 1.265 0.0896 8477    1   3.325  0.0079
##  rand / scramb        0.771 0.0518 8477    1  -3.867  0.0011
##  rand / silhouette    0.839 0.0621 8477    1  -2.379  0.1212
##  scramb / silhouette  1.087 0.0889 8477    1   1.021  0.8456
## 
## P value adjustment: tukey method for comparing a family of 5 estimates 
## Tests are performed on the log scale 
## [1] 16.36667
##  contrast             ratio     SE   df null t.ratio p.value
##  bio / ellipse        0.812 0.0491 8870    1  -3.447  0.0052
##  bio / rand           1.188 0.0875 8870    1   2.335  0.1340
##  bio / scramb         0.940 0.0606 8870    1  -0.959  0.8734
##  bio / silhouette     1.027 0.0782 8870    1   0.346  0.9969
##  ellipse / rand       1.463 0.1140 8870    1   4.885  <.0001
##  ellipse / scramb     1.158 0.0712 8870    1   2.388  0.1187
##  ellipse / silhouette 1.265 0.0794 8870    1   3.744  0.0017
##  rand / scramb        0.791 0.0530 8870    1  -3.492  0.0044
##  rand / silhouette    0.864 0.0594 8870    1  -2.122  0.2109
##  scramb / silhouette  1.092 0.0798 8870    1   1.207  0.7475
## 
## P value adjustment: tukey method for comparing a family of 5 estimates 
## Tests are performed on the log scale 
## [1] 17.53333
##  contrast             ratio     SE   df null t.ratio p.value
##  bio / ellipse        0.905 0.0527 9637    1  -1.720  0.4218
##  bio / rand           1.242 0.0720 9637    1   3.733  0.0018
##  bio / scramb         1.138 0.0632 9637    1   2.319  0.1388
##  bio / silhouette     1.190 0.0976 9637    1   2.122  0.2108
##  ellipse / rand       1.373 0.1030 9637    1   4.221  0.0002
##  ellipse / scramb     1.257 0.0980 9637    1   2.940  0.0273
##  ellipse / silhouette 1.316 0.0998 9637    1   3.616  0.0028
##  rand / scramb        0.916 0.0553 9637    1  -1.450  0.5948
##  rand / silhouette    0.958 0.0700 9637    1  -0.581  0.9779
##  scramb / silhouette  1.046 0.0899 9637    1   0.526  0.9847
## 
## P value adjustment: tukey method for comparing a family of 5 estimates 
## Tests are performed on the log scale 
## [1] 17.7
##  contrast             ratio     SE   df null t.ratio p.value
##  bio / ellipse        0.909 0.0469 9420    1  -1.854  0.3426
##  bio / rand           1.245 0.0691 9420    1   3.943  0.0008
##  bio / scramb         1.113 0.0602 9420    1   1.985  0.2735
##  bio / silhouette     1.193 0.0897 9420    1   2.346  0.1305
##  ellipse / rand       1.370 0.0938 9420    1   4.594  <.0001
##  ellipse / scramb     1.225 0.0825 9420    1   3.017  0.0216
##  ellipse / silhouette 1.313 0.0923 9420    1   3.872  0.0010
##  rand / scramb        0.895 0.0507 9420    1  -1.965  0.2832
##  rand / silhouette    0.959 0.0670 9420    1  -0.605  0.9743
##  scramb / silhouette  1.072 0.0816 9420    1   0.907  0.8942
## 
## P value adjustment: tukey method for comparing a family of 5 estimates 
## Tests are performed on the log scale 
## [1] 19.03333
##  contrast             ratio     SE    df null t.ratio p.value
##  bio / ellipse        0.885 0.0313 10272    1  -3.444  0.0052
##  bio / rand           1.177 0.0664 10272    1   2.889  0.0317
##  bio / scramb         1.102 0.0568 10272    1   1.882  0.3268
##  bio / silhouette     1.168 0.1120 10272    1   1.612  0.4899
##  ellipse / rand       1.330 0.0918 10272    1   4.124  0.0004
##  ellipse / scramb     1.245 0.0810 10272    1   3.364  0.0069
##  ellipse / silhouette 1.319 0.1170 10272    1   3.122  0.0155
##  rand / scramb        0.936 0.0550 10272    1  -1.123  0.7946
##  rand / silhouette    0.992 0.0955 10272    1  -0.081  1.0000
##  scramb / silhouette  1.060 0.1080 10272    1   0.568  0.9796
## 
## P value adjustment: tukey method for comparing a family of 5 estimates 
## Tests are performed on the log scale 
## [1] 19.2
##  contrast             ratio     SE   df null t.ratio p.value
##  bio / ellipse        0.889 0.0294 9993    1  -3.567  0.0033
##  bio / rand           1.213 0.0663 9993    1   3.533  0.0038
##  bio / scramb         1.097 0.0560 9993    1   1.811  0.3670
##  bio / silhouette     1.139 0.1120 9993    1   1.326  0.6747
##  ellipse / rand       1.365 0.0879 9993    1   4.836  <.0001
##  ellipse / scramb     1.234 0.0774 9993    1   3.360  0.0070
##  ellipse / silhouette 1.282 0.1160 9993    1   2.759  0.0459
##  rand / scramb        0.904 0.0542 9993    1  -1.680  0.4466
##  rand / silhouette    0.939 0.1000 9993    1  -0.585  0.9773
##  scramb / silhouette  1.039 0.1120 9993    1   0.355  0.9966
## 
## P value adjustment: tukey method for comparing a family of 5 estimates 
## Tests are performed on the log scale 
## [1] 19.36667
##  contrast             ratio     SE    df null t.ratio p.value
##  bio / ellipse        0.921 0.0286 10400    1  -2.645  0.0626
##  bio / rand           1.245 0.0711 10400    1   3.839  0.0012
##  bio / scramb         1.061 0.0514 10400    1   1.227  0.7358
##  bio / silhouette     1.176 0.1070 10400    1   1.781  0.3845
##  ellipse / rand       1.351 0.0796 10400    1   5.111  <.0001
##  ellipse / scramb     1.152 0.0670 10400    1   2.432  0.1070
##  ellipse / silhouette 1.276 0.1100 10400    1   2.833  0.0372
##  rand / scramb        0.852 0.0604 10400    1  -2.253  0.1605
##  rand / silhouette    0.944 0.0971 10400    1  -0.557  0.9811
##  scramb / silhouette  1.108 0.0994 10400    1   1.142  0.7842
## 
## P value adjustment: tukey method for comparing a family of 5 estimates 
## Tests are performed on the log scale 
## [1] 19.53333
##  contrast             ratio     SE    df null t.ratio p.value
##  bio / ellipse        0.930 0.0326 10493    1  -2.076  0.2304
##  bio / rand           1.255 0.0751 10493    1   3.802  0.0014
##  bio / scramb         1.071 0.0483 10493    1   1.525  0.5464
##  bio / silhouette     1.159 0.0999 10493    1   1.708  0.4292
##  ellipse / rand       1.350 0.0829 10493    1   4.894  <.0001
##  ellipse / scramb     1.152 0.0637 10493    1   2.559  0.0781
##  ellipse / silhouette 1.246 0.1010 10493    1   2.715  0.0519
##  rand / scramb        0.853 0.0596 10493    1  -2.275  0.1530
##  rand / silhouette    0.923 0.0939 10493    1  -0.789  0.9338
##  scramb / silhouette  1.082 0.0933 10493    1   0.910  0.8931
## 
## P value adjustment: tukey method for comparing a family of 5 estimates 
## Tests are performed on the log scale 
## [1] 19.7
##  contrast             ratio     SE    df null t.ratio p.value
##  bio / ellipse        0.938 0.0371 10237    1  -1.622  0.4830
##  bio / rand           1.277 0.0814 10237    1   3.839  0.0012
##  bio / scramb         1.065 0.0471 10237    1   1.434  0.6054
##  bio / silhouette     1.114 0.0885 10237    1   1.363  0.6514
##  ellipse / rand       1.362 0.0903 10237    1   4.656  <.0001
##  ellipse / scramb     1.136 0.0650 10237    1   2.231  0.1684
##  ellipse / silhouette 1.188 0.0928 10237    1   2.208  0.1768
##  rand / scramb        0.834 0.0570 10237    1  -2.653  0.0612
##  rand / silhouette    0.873 0.0873 10237    1  -1.362  0.6521
##  scramb / silhouette  1.046 0.0877 10237    1   0.534  0.9839
## 
## P value adjustment: tukey method for comparing a family of 5 estimates 
## Tests are performed on the log scale
```

###### 2.2.2.3.1 Summary

```
moving_results_posthoc <- moving_results_posthoc[-1,]
write.csv(moving_results_posthoc, paste0(tmp_path, 'interEyeDist_movingResultsPostHoc.csv'))

kable(moving_results_posthoc)
```

|  | win\_starts | win\_span | bio.ellipse | bio.rand | bio.scramb | bio.silhouette | ellipse.rand | ellipse.scramb | ellipse.silhouette | rand.scramb | rand.silhouette | scramb.silhouette |
| --- | --- | --- | --- | --- | --- | --- | --- | --- | --- | --- | --- | --- |
| 2 | 11.86667 | 1 | 0.0066766 | 0.0001464 | 0.9440746 | 0.9914775 | 0.0000007 | 0.0079919 | 0.0002360 | 0.0039854 | 0.0399043 | 1.0000000 |
| 3 | 12.03333 | 1 | 0.0094111 | 0.0000084 | 0.8152700 | 0.9976550 | 0.0000000 | 0.0035495 | 0.0009254 | 0.0004772 | 0.0025426 | 0.9961483 |
| 4 | 12.20000 | 1 | 0.0129101 | 0.0000077 | 0.6526758 | 0.9990263 | 0.0000000 | 0.0029032 | 0.0115254 | 0.0002765 | 0.0003020 | 0.9715562 |
| 5 | 12.36667 | 1 | 0.0372580 | 0.0000029 | 0.6924963 | 0.9998017 | 0.0000000 | 0.0074717 | 0.0316380 | 0.0000031 | 0.0000024 | 0.9513558 |
| 6 | 12.53333 | 1 | 0.1180768 | 0.0000056 | 0.7330217 | 0.9998565 | 0.0000000 | 0.0286636 | 0.0704264 | 0.0000026 | 0.0000008 | 0.9627145 |
| 7 | 12.70000 | 1 | 0.5991149 | 0.0000432 | 0.9528889 | 0.9996693 | 0.0000003 | 0.2831517 | 0.5422903 | 0.0000000 | 0.0000002 | 0.9337147 |
| 8 | 12.86667 | 1 | 0.9961184 | 0.0000241 | 0.7109888 | 0.8324132 | 0.0000214 | 0.7507158 | 0.6873709 | 0.0000001 | 0.0000489 | 1.0000000 |
| 9 | 13.03333 | 1 | 0.9998279 | 0.0000364 | 0.7001518 | 0.6335263 | 0.0001685 | 0.9275481 | 0.6986363 | 0.0000000 | 0.0024584 | 0.9887387 |
| 10 | 13.20000 | 1 | 0.9999960 | 0.0001349 | 0.8030782 | 0.4816777 | 0.0001152 | 0.9446786 | 0.4562127 | 0.0000025 | 0.0395405 | 0.8936325 |
| 11 | 13.36667 | 1 | 0.9999731 | 0.0002437 | 0.6795879 | 0.1681982 | 0.0003902 | 0.8343282 | 0.0721666 | 0.0001568 | 0.3605728 | 0.4464728 |
| 12 | 13.53333 | 1 | 0.9998221 | 0.0000281 | 0.4208529 | 0.0257715 | 0.0000815 | 0.6732510 | 0.0115456 | 0.0002317 | 0.6794236 | 0.1758702 |
| 13 | 13.70000 | 1 | 0.9860739 | 0.0000420 | 0.4528327 | 0.0010229 | 0.0000023 | 0.4227516 | 0.0004577 | 0.0040814 | 0.9821345 | 0.0611552 |
| 14 | 13.86667 | 1 | 0.5273674 | 0.0018965 | 0.6976550 | 0.0081491 | 0.0000001 | 0.1376746 | 0.0006103 | 0.0140167 | 0.9782934 | 0.1359656 |
| 15 | 14.03333 | 1 | 0.3861060 | 0.0092712 | 0.5109211 | 0.0571457 | 0.0000001 | 0.0266087 | 0.0037404 | 0.0526782 | 0.9068687 | 0.5683300 |
| 16 | 14.20000 | 1 | 0.1479572 | 0.0195565 | 0.6326799 | 0.1907417 | 0.0000000 | 0.0039736 | 0.0042022 | 0.0409260 | 0.9019515 | 0.6627846 |
| 17 | 14.36667 | 1 | 0.1013326 | 0.0399263 | 0.6085479 | 0.5418043 | 0.0000000 | 0.0010107 | 0.0181290 | 0.1296316 | 0.7125594 | 0.9844907 |
| 18 | 14.53333 | 1 | 0.0280045 | 0.1332706 | 0.9100866 | 0.8947377 | 0.0000000 | 0.0021514 | 0.0362906 | 0.2199815 | 0.7639670 | 0.9971420 |
| 19 | 15.70000 | 1 | 0.0034736 | 0.0113239 | 0.7243147 | 0.9978441 | 0.0000035 | 0.3186385 | 0.0811288 | 0.0008238 | 0.1199775 | 0.9737447 |
| 20 | 15.86667 | 1 | 0.0057117 | 0.0146120 | 0.7371964 | 0.9999836 | 0.0000068 | 0.3051330 | 0.0468672 | 0.0015676 | 0.1835203 | 0.9200631 |
| 21 | 16.03333 | 1 | 0.0035937 | 0.0334370 | 0.7044436 | 0.9999989 | 0.0000085 | 0.1810486 | 0.0405224 | 0.0022531 | 0.2309467 | 0.9335138 |
| 22 | 16.20000 | 1 | 0.0015905 | 0.0637284 | 0.5512818 | 1.0000000 | 0.0000016 | 0.1304259 | 0.0079149 | 0.0010521 | 0.1212448 | 0.8456003 |
| 23 | 16.36667 | 1 | 0.0051545 | 0.1339972 | 0.8733638 | 0.9969251 | 0.0000104 | 0.1187163 | 0.0017107 | 0.0043974 | 0.2108726 | 0.7475186 |
| 24 | 17.53333 | 1 | 0.4217856 | 0.0017794 | 0.1387802 | 0.2107820 | 0.0002381 | 0.0272603 | 0.0027794 | 0.5948453 | 0.9779302 | 0.9847272 |
| 25 | 17.70000 | 1 | 0.3426318 | 0.0007722 | 0.2734826 | 0.1305281 | 0.0000434 | 0.0215686 | 0.0010282 | 0.2831510 | 0.9743310 | 0.8941990 |
| 26 | 19.03333 | 1 | 0.0052079 | 0.0316792 | 0.3268395 | 0.4899125 | 0.0003611 | 0.0069157 | 0.0154869 | 0.7946490 | 0.9999902 | 0.9796243 |
| 27 | 19.20000 | 1 | 0.0033400 | 0.0037809 | 0.3669737 | 0.6746580 | 0.0000133 | 0.0070036 | 0.0459320 | 0.4465512 | 0.9773018 | 0.9966042 |
| 28 | 19.36667 | 1 | 0.0625548 | 0.0011753 | 0.7357503 | 0.3844627 | 0.0000032 | 0.1069728 | 0.0372395 | 0.1605152 | 0.9810697 | 0.7841712 |
| 29 | 19.53333 | 1 | 0.2303928 | 0.0013572 | 0.5463692 | 0.4292225 | 0.0000099 | 0.0781295 | 0.0519112 | 0.1530026 | 0.9338391 | 0.8930744 |
| 30 | 19.70000 | 1 | 0.4829994 | 0.0011718 | 0.6054172 | 0.6514178 | 0.0000322 | 0.1684496 | 0.1767818 | 0.0612207 | 0.6521044 | 0.9838515 |

##### 2.2.3 Inter-eye distance change

###### 2.2.3.1 Static

remaking specific dataframe

```
onlystatic <- subset(full_data, full_data$moving == 0)

data <- onlystatic
```

reloading of the data and ps

```
static <- read.csv(paste0(tmp_path, 'interEyeDistChange_staticResults.csv'))

static$adjustedp <- static$p*sum(static$stimtype=='bio') # create bonferroni corrected p
static$adjustedp[static$adjustedp>1] <- 1
significant_static <- subset(static, static$stimtype=='bio') #subset to have only 1 row per model

significant_static <- subset(significant_static, significant_static$adjustedp < alpha)

significant_static_wins <- significant_static$win_starts #identify the windows that are significant
```

###### 2.2.3.1.1 Detailed

now redo the models

```
static_results_posthoc <- data.frame(list('win_starts'=NaN,'win_span'=NaN,
                                          'bio/ellipse'=NaN, 'bio/none'=NaN,
                                          'bio/rand'=NaN, 'bio/scramb'=NaN,
                                          'bio/silhouette'=NaN, 'ellipse/none'=NaN,
                                          'ellipse/rand'=NaN, 'ellipse/scramb'=NaN,
                                          'ellipse/silhouette'=NaN, 'none/rand'=NaN,
                                          'none/scramb'=NaN, 'none/silhouette'=NaN,
                                          'rand/scramb'=NaN, 'rand/silhouette'=NaN,
                                          'scramb/silhouette'=NaN))

for (window_start in significant_static_wins){
  window_end <- window_start + window_size

  window_data <- subset(data, data$time_from_start>window_start)
  window_data <- subset(window_data, window_data$time_from_start<window_end)

  full_tmb <- glmmTMB(log(inter_eye_dist_change) ~ stimtype + (stimtype|subj), family=gaussian, data=window_data)
  Anova(full_tmb)
  e <- emmeans(full_tmb, ~stimtype, type='response')
  prs <- as.data.frame(pairs(e))
  newrow <- c(window_start, window_size, prs$p.value[1], prs$p.value[2],
              prs$p.value[3], prs$p.value[4], prs$p.value[5], prs$p.value[6],
              prs$p.value[7], prs$p.value[8], prs$p.value[9], prs$p.value[10],
              prs$p.value[11], prs$p.value[12], prs$p.value[13], prs$p.value[14],
              prs$p.value[15])
  static_results_posthoc <- rbind(static_results_posthoc, newrow)
  print(window_start)
  print(pairs(e))
}
```

```
## Warning in log(inter_eye_dist_change): NaNs produced
## Warning in log(inter_eye_dist_change): NaNs produced
```

```
## Warning in finalizeTMB(TMBStruc, obj, fit, h, data.tmb.old): Model convergence
## problem; non-positive-definite Hessian matrix. See vignette('troubleshooting')
```

```
## [1] 0.03333333
##  contrast             ratio     SE    df null t.ratio p.value
##  bio / ellipse        1.084 0.0856 12751    1   1.018  0.9122
##  bio / none           1.259 0.0905 12751    1   3.205  0.0171
##  bio / rand           1.035 0.0952 12751    1   0.372  0.9991
##  bio / scramb         0.983 0.0726 12751    1  -0.228  0.9999
##  bio / silhouette     0.840 0.0727 12751    1  -2.013  0.3349
##  ellipse / none       1.162 0.0670 12751    1   2.603  0.0965
##  ellipse / rand       0.955 0.0679 12751    1  -0.649  0.9872
##  ellipse / scramb     0.907 0.0759 12751    1  -1.162  0.8547
##  ellipse / silhouette 0.775 0.0637 12751    1  -3.099  0.0239
##  none / rand          0.822 0.0673 12751    1  -2.395  0.1581
##  none / scramb        0.781 0.0546 12751    1  -3.536  0.0055
##  none / silhouette    0.667 0.0475 12751    1  -5.686  <.0001
##  rand / scramb        0.950 0.0834 12751    1  -0.582  0.9923
##  rand / silhouette    0.812 0.0728 12751    1  -2.323  0.1848
##  scramb / silhouette  0.854 0.0823 12751    1  -1.633  0.5763
## 
## P value adjustment: tukey method for comparing a family of 6 estimates 
## Tests are performed on the log scale
```

```
## Warning in log(inter_eye_dist_change): NaNs produced
```

```
## Warning in log(inter_eye_dist_change): NaNs produced
```

```
## Warning in finalizeTMB(TMBStruc, obj, fit, h, data.tmb.old): Model convergence
## problem; non-positive-definite Hessian matrix. See vignette('troubleshooting')
```

```
## Warning in finalizeTMB(TMBStruc, obj, fit, h, data.tmb.old): Model convergence
## problem; singular convergence (7). See vignette('troubleshooting'),
## help('diagnose')
```

```
## [1] 0.2
##  contrast             ratio     SE    df null t.ratio p.value
##  bio / ellipse        1.090 0.0844 12215    1   1.119  0.8738
##  bio / none           1.336 0.0983 12215    1   3.934  0.0012
##  bio / rand           1.024 0.0924 12215    1   0.258  0.9998
##  bio / scramb         0.995 0.0799 12215    1  -0.057  1.0000
##  bio / silhouette     0.829 0.0739 12215    1  -2.104  0.2850
##  ellipse / none       1.225 0.0872 12215    1   2.850  0.0500
##  ellipse / rand       0.939 0.0720 12215    1  -0.825  0.9630
##  ellipse / scramb     0.913 0.0903 12215    1  -0.921  0.9411
##  ellipse / silhouette 0.760 0.0687 12215    1  -3.031  0.0294
##  none / rand          0.766 0.0685 12215    1  -2.976  0.0348
##  none / scramb        0.745 0.0596 12215    1  -3.673  0.0033
##  none / silhouette    0.621 0.0534 12215    1  -5.545  <.0001
##  rand / scramb        0.973 0.0920 12215    1  -0.294  0.9997
##  rand / silhouette    0.810 0.0750 12215    1  -2.277  0.2036
##  scramb / silhouette  0.833 0.0898 12215    1  -1.697  0.5341
## 
## P value adjustment: tukey method for comparing a family of 6 estimates 
## Tests are performed on the log scale
```

```
## Warning in log(inter_eye_dist_change): NaNs produced
```

```
## Warning in log(inter_eye_dist_change): NaNs produced
```

```
## Warning in finalizeTMB(TMBStruc, obj, fit, h, data.tmb.old): Model convergence
## problem; non-positive-definite Hessian matrix. See vignette('troubleshooting')
```

```
## Warning in finalizeTMB(TMBStruc, obj, fit, h, data.tmb.old): Model convergence
## problem; singular convergence (7). See vignette('troubleshooting'),
## help('diagnose')
```

```
## [1] 0.3666667
##  contrast             ratio     SE    df null t.ratio p.value
##  bio / ellipse        1.115 0.0958 12602    1   1.268  0.8023
##  bio / none           1.401 0.1120 12602    1   4.238  0.0003
##  bio / rand           1.074 0.0963 12602    1   0.797  0.9681
##  bio / scramb         1.049 0.0858 12602    1   0.583  0.9922
##  bio / silhouette     0.904 0.0754 12602    1  -1.204  0.8350
##  ellipse / none       1.257 0.0860 12602    1   3.337  0.0110
##  ellipse / rand       0.963 0.0700 12602    1  -0.516  0.9956
##  ellipse / scramb     0.941 0.0878 12602    1  -0.657  0.9865
##  ellipse / silhouette 0.811 0.0709 12602    1  -2.397  0.1572
##  none / rand          0.767 0.0676 12602    1  -3.014  0.0310
##  none / scramb        0.748 0.0563 12602    1  -3.849  0.0017
##  none / silhouette    0.645 0.0503 12602    1  -5.622  <.0001
##  rand / scramb        0.976 0.0875 12602    1  -0.265  0.9998
##  rand / silhouette    0.842 0.0720 12602    1  -2.009  0.3370
##  scramb / silhouette  0.862 0.0860 12602    1  -1.486  0.6736
## 
## P value adjustment: tukey method for comparing a family of 6 estimates 
## Tests are performed on the log scale
```

```
## Warning in log(inter_eye_dist_change): NaNs produced
```

```
## Warning in log(inter_eye_dist_change): NaNs produced
```

```
## Warning in finalizeTMB(TMBStruc, obj, fit, h, data.tmb.old): Model convergence
## problem; non-positive-definite Hessian matrix. See vignette('troubleshooting')
```

```
## Warning in finalizeTMB(TMBStruc, obj, fit, h, data.tmb.old): Model convergence
## problem; singular convergence (7). See vignette('troubleshooting'),
## help('diagnose')
```

```
## [1] 0.5333333
##  contrast             ratio     SE    df null t.ratio p.value
##  bio / ellipse        1.100 0.0885 12501    1   1.185  0.8441
##  bio / none           1.428 0.1090 12501    1   4.687  <.0001
##  bio / rand           1.096 0.0919 12501    1   1.090  0.8858
##  bio / scramb         1.110 0.0946 12501    1   1.227  0.8237
##  bio / silhouette     0.915 0.0804 12501    1  -1.006  0.9163
##  ellipse / none       1.298 0.0904 12501    1   3.753  0.0024
##  ellipse / rand       0.996 0.0724 12501    1  -0.055  1.0000
##  ellipse / scramb     1.009 0.0964 12501    1   0.097  1.0000
##  ellipse / silhouette 0.832 0.0786 12501    1  -1.945  0.3747
##  none / rand          0.767 0.0626 12501    1  -3.249  0.0147
##  none / scramb        0.777 0.0553 12501    1  -3.543  0.0053
##  none / silhouette    0.641 0.0523 12501    1  -5.450  <.0001
##  rand / scramb        1.013 0.0867 12501    1   0.154  1.0000
##  rand / silhouette    0.836 0.0783 12501    1  -1.918  0.3913
##  scramb / silhouette  0.825 0.0882 12501    1  -1.803  0.4641
## 
## P value adjustment: tukey method for comparing a family of 6 estimates 
## Tests are performed on the log scale
```

```
## Warning in log(inter_eye_dist_change): NaNs produced
```

```
## Warning in log(inter_eye_dist_change): NaNs produced
```

```
## [1] 0.7
##  contrast             ratio     SE    df null t.ratio p.value
##  bio / ellipse        1.076 0.0881 11985    1   0.889  0.9492
##  bio / none           1.390 0.0975 11985    1   4.693  <.0001
##  bio / rand           1.032 0.0814 11985    1   0.398  0.9987
##  bio / scramb         1.041 0.0925 11985    1   0.450  0.9977
##  bio / silhouette     0.936 0.0737 11985    1  -0.840  0.9601
##  ellipse / none       1.292 0.0823 11985    1   4.027  0.0008
##  ellipse / rand       0.959 0.0591 11985    1  -0.673  0.9849
##  ellipse / scramb     0.968 0.0826 11985    1  -0.385  0.9989
##  ellipse / silhouette 0.870 0.0767 11985    1  -1.578  0.6134
##  none / rand          0.742 0.0487 11985    1  -4.536  0.0001
##  none / scramb        0.749 0.0486 11985    1  -4.459  0.0001
##  none / silhouette    0.673 0.0520 11985    1  -5.116  <.0001
##  rand / scramb        1.009 0.0772 11985    1   0.112  1.0000
##  rand / silhouette    0.907 0.0754 11985    1  -1.175  0.8491
##  scramb / silhouette  0.899 0.0894 11985    1  -1.068  0.8940
## 
## P value adjustment: tukey method for comparing a family of 6 estimates 
## Tests are performed on the log scale
```

```
## Warning in log(inter_eye_dist_change): NaNs produced
## Warning in log(inter_eye_dist_change): NaNs produced
```

```
## Warning in finalizeTMB(TMBStruc, obj, fit, h, data.tmb.old): Model convergence
## problem; non-positive-definite Hessian matrix. See vignette('troubleshooting')
```

```
## [1] 0.8666667
##  contrast             ratio     SE    df null t.ratio p.value
##  bio / ellipse        1.101 0.0892 12230    1   1.192  0.8408
##  bio / none           1.405 0.0935 12230    1   5.115  <.0001
##  bio / rand           1.010 0.0711 12230    1   0.146  1.0000
##  bio / scramb         1.084 0.0933 12230    1   0.935  0.9376
##  bio / silhouette     1.001 0.0621 12230    1   0.011  1.0000
##  ellipse / none       1.276 0.0760 12230    1   4.091  0.0006
##  ellipse / rand       0.917 0.0606 12230    1  -1.307  0.7815
##  ellipse / scramb     0.984 0.0742 12230    1  -0.213  0.9999
##  ellipse / silhouette 0.909 0.0786 12230    1  -1.108  0.8784
##  none / rand          0.719 0.0481 12230    1  -4.931  <.0001
##  none / scramb        0.771 0.0470 12230    1  -4.262  0.0003
##  none / silhouette    0.712 0.0493 12230    1  -4.909  <.0001
##  rand / scramb        1.073 0.0874 12230    1   0.862  0.9555
##  rand / silhouette    0.991 0.0794 12230    1  -0.119  1.0000
##  scramb / silhouette  0.923 0.0835 12230    1  -0.882  0.9509
## 
## P value adjustment: tukey method for comparing a family of 6 estimates 
## Tests are performed on the log scale
```

```
## Warning in log(inter_eye_dist_change): NaNs produced
```

```
## Warning in log(inter_eye_dist_change): NaNs produced
```

```
## Warning in finalizeTMB(TMBStruc, obj, fit, h, data.tmb.old): Model convergence
## problem; non-positive-definite Hessian matrix. See vignette('troubleshooting')
```

```
## [1] 1.033333
##  contrast             ratio     SE    df null t.ratio p.value
##  bio / ellipse        1.081 0.0836 12185    1   1.004  0.9168
##  bio / none           1.359 0.0932 12185    1   4.469  0.0001
##  bio / rand           1.042 0.0705 12185    1   0.603  0.9909
##  bio / scramb         1.063 0.0865 12185    1   0.754  0.9750
##  bio / silhouette     1.055 0.0678 12185    1   0.839  0.9602
##  ellipse / none       1.257 0.0693 12185    1   4.152  0.0005
##  ellipse / rand       0.964 0.0697 12185    1  -0.509  0.9959
##  ellipse / scramb     0.984 0.0648 12185    1  -0.248  0.9999
##  ellipse / silhouette 0.977 0.0816 12185    1  -0.284  0.9998
##  none / rand          0.767 0.0537 12185    1  -3.792  0.0021
##  none / scramb        0.782 0.0466 12185    1  -4.119  0.0005
##  none / silhouette    0.777 0.0558 12185    1  -3.520  0.0058
##  rand / scramb        1.021 0.0846 12185    1   0.247  0.9999
##  rand / silhouette    1.013 0.0839 12185    1   0.158  1.0000
##  scramb / silhouette  0.993 0.0843 12185    1  -0.087  1.0000
## 
## P value adjustment: tukey method for comparing a family of 6 estimates 
## Tests are performed on the log scale
```

```
## Warning in log(inter_eye_dist_change): NaNs produced
```

```
## Warning in log(inter_eye_dist_change): NaNs produced
```

```
## Warning in finalizeTMB(TMBStruc, obj, fit, h, data.tmb.old): Model convergence
## problem; non-positive-definite Hessian matrix. See vignette('troubleshooting')
```

```
## [1] 1.2
##  contrast             ratio     SE    df null t.ratio p.value
##  bio / ellipse        1.044 0.0911 11816    1   0.496  0.9963
##  bio / none           1.317 0.0842 11816    1   4.305  0.0002
##  bio / rand           1.022 0.0726 11816    1   0.302  0.9997
##  bio / scramb         1.103 0.0912 11816    1   1.187  0.8432
##  bio / silhouette     1.010 0.0628 11816    1   0.166  1.0000
##  ellipse / none       1.261 0.0784 11816    1   3.734  0.0026
##  ellipse / rand       0.979 0.0807 11816    1  -0.264  0.9998
##  ellipse / scramb     1.057 0.0746 11816    1   0.779  0.9712
##  ellipse / silhouette 0.968 0.0835 11816    1  -0.381  0.9990
##  none / rand          0.776 0.0516 11816    1  -3.819  0.0019
##  none / scramb        0.838 0.0479 11816    1  -3.102  0.0237
##  none / silhouette    0.767 0.0508 11816    1  -4.004  0.0009
##  rand / scramb        1.080 0.0959 11816    1   0.864  0.9550
##  rand / silhouette    0.989 0.0760 11816    1  -0.145  1.0000
##  scramb / silhouette  0.916 0.0780 11816    1  -1.031  0.9076
## 
## P value adjustment: tukey method for comparing a family of 6 estimates 
## Tests are performed on the log scale
```

```
## Warning in log(inter_eye_dist_change): NaNs produced
```

```
## Warning in log(inter_eye_dist_change): NaNs produced
```

```
## Warning in finalizeTMB(TMBStruc, obj, fit, h, data.tmb.old): Model convergence
## problem; non-positive-definite Hessian matrix. See vignette('troubleshooting')
```

```
## Warning in finalizeTMB(TMBStruc, obj, fit, h, data.tmb.old): Model convergence
## problem; singular convergence (7). See vignette('troubleshooting'),
## help('diagnose')
```

```
## [1] 1.366667
##  contrast             ratio     SE    df null t.ratio p.value
##  bio / ellipse        0.966 0.0826 12301    1  -0.402  0.9987
##  bio / none           1.247 0.0716 12301    1   3.839  0.0017
##  bio / rand           0.972 0.0711 12301    1  -0.394  0.9988
##  bio / scramb         1.046 0.0898 12301    1   0.522  0.9953
##  bio / silhouette     0.942 0.0781 12301    1  -0.715  0.9802
##  ellipse / none       1.290 0.0801 12301    1   4.109  0.0006
##  ellipse / rand       1.006 0.0791 12301    1   0.070  1.0000
##  ellipse / scramb     1.082 0.0940 12301    1   0.912  0.9437
##  ellipse / silhouette 0.975 0.0857 12301    1  -0.284  0.9998
##  none / rand          0.779 0.0546 12301    1  -3.561  0.0050
##  none / scramb        0.839 0.0562 12301    1  -2.625  0.0913
##  none / silhouette    0.756 0.0627 12301    1  -3.376  0.0096
##  rand / scramb        1.076 0.1060 12301    1   0.745  0.9763
##  rand / silhouette    0.970 0.0866 12301    1  -0.341  0.9994
##  scramb / silhouette  0.901 0.0954 12301    1  -0.984  0.9231
## 
## P value adjustment: tukey method for comparing a family of 6 estimates 
## Tests are performed on the log scale
```

```
## Warning in log(inter_eye_dist_change): NaNs produced
```

```
## Warning in log(inter_eye_dist_change): NaNs produced
```

```
## Warning in finalizeTMB(TMBStruc, obj, fit, h, data.tmb.old): Model convergence
## problem; singular convergence (7). See vignette('troubleshooting'),
## help('diagnose')
```

```
## [1] 1.533333
##  contrast             ratio     SE    df null t.ratio p.value
##  bio / ellipse        0.956 0.0843 12302    1  -0.515  0.9956
##  bio / none           1.219 0.0714 12302    1   3.388  0.0092
##  bio / rand           0.954 0.0670 12302    1  -0.671  0.9851
##  bio / scramb         1.050 0.0869 12302    1   0.590  0.9917
##  bio / silhouette     0.941 0.0800 12302    1  -0.710  0.9808
##  ellipse / none       1.276 0.0790 12302    1   3.938  0.0012
##  ellipse / rand       0.998 0.0814 12302    1  -0.020  1.0000
##  ellipse / scramb     1.099 0.0912 12302    1   1.136  0.8664
##  ellipse / silhouette 0.985 0.0841 12302    1  -0.175  1.0000
##  none / rand          0.782 0.0525 12302    1  -3.657  0.0035
##  none / scramb        0.861 0.0561 12302    1  -2.294  0.1964
##  none / silhouette    0.772 0.0644 12302    1  -3.100  0.0238
##  rand / scramb        1.101 0.0980 12302    1   1.078  0.8905
##  rand / silhouette    0.987 0.0855 12302    1  -0.154  1.0000
##  scramb / silhouette  0.897 0.0914 12302    1  -1.071  0.8929
## 
## P value adjustment: tukey method for comparing a family of 6 estimates 
## Tests are performed on the log scale
```

```
## Warning in log(inter_eye_dist_change): NaNs produced
```

```
## Warning in log(inter_eye_dist_change): NaNs produced
```

```
## Warning in finalizeTMB(TMBStruc, obj, fit, h, data.tmb.old): Model convergence
## problem; non-positive-definite Hessian matrix. See vignette('troubleshooting')
```

```
## Warning in finalizeTMB(TMBStruc, obj, fit, h, data.tmb.old): Model convergence
## problem; singular convergence (7). See vignette('troubleshooting'),
## help('diagnose')
```

```
## [1] 1.7
##  contrast             ratio     SE    df null t.ratio p.value
##  bio / ellipse        0.873 0.0745 11934    1  -1.595  0.6019
##  bio / none           1.145 0.0690 11934    1   2.247  0.2166
##  bio / rand           0.955 0.0665 11934    1  -0.656  0.9865
##  bio / scramb         1.024 0.0861 11934    1   0.287  0.9997
##  bio / silhouette     0.884 0.0811 11934    1  -1.339  0.7631
##  ellipse / none       1.312 0.0810 11934    1   4.400  0.0002
##  ellipse / rand       1.095 0.0909 11934    1   1.090  0.8856
##  ellipse / scramb     1.174 0.0957 11934    1   1.967  0.3615
##  ellipse / silhouette 1.014 0.0849 11934    1   0.161  1.0000
##  none / rand          0.834 0.0607 11934    1  -2.488  0.1276
##  none / scramb        0.895 0.0579 11934    1  -1.719  0.5191
##  none / silhouette    0.772 0.0678 11934    1  -2.940  0.0386
##  rand / scramb        1.072 0.0968 11934    1   0.774  0.9720
##  rand / silhouette    0.926 0.0858 11934    1  -0.832  0.9616
##  scramb / silhouette  0.863 0.0869 11934    1  -1.459  0.6903
## 
## P value adjustment: tukey method for comparing a family of 6 estimates 
## Tests are performed on the log scale
```

```
## Warning in log(inter_eye_dist_change): NaNs produced
```

```
## Warning in log(inter_eye_dist_change): NaNs produced
```

```
## [1] 2.366667
##  contrast             ratio     SE    df null t.ratio p.value
##  bio / ellipse        0.900 0.0835 12344    1  -1.138  0.8655
##  bio / none           1.179 0.0802 12344    1   2.424  0.1481
##  bio / rand           0.988 0.0768 12344    1  -0.158  1.0000
##  bio / scramb         1.038 0.0898 12344    1   0.430  0.9982
##  bio / silhouette     0.944 0.0888 12344    1  -0.607  0.9906
##  ellipse / none       1.311 0.0905 12344    1   3.919  0.0013
##  ellipse / rand       1.098 0.0867 12344    1   1.182  0.8456
##  ellipse / scramb     1.154 0.0975 12344    1   1.690  0.5384
##  ellipse / silhouette 1.050 0.0878 12344    1   0.581  0.9923
##  none / rand          0.838 0.0539 12344    1  -2.754  0.0652
##  none / scramb        0.880 0.0480 12344    1  -2.342  0.1775
##  none / silhouette    0.801 0.0613 12344    1  -2.902  0.0431
##  rand / scramb        1.051 0.0871 12344    1   0.597  0.9913
##  rand / silhouette    0.956 0.0771 12344    1  -0.556  0.9937
##  scramb / silhouette  0.910 0.0810 12344    1  -1.060  0.8971
## 
## P value adjustment: tukey method for comparing a family of 6 estimates 
## Tests are performed on the log scale
```

```
## Warning in log(inter_eye_dist_change): NaNs produced
## Warning in log(inter_eye_dist_change): NaNs produced
```

```
## Warning in finalizeTMB(TMBStruc, obj, fit, h, data.tmb.old): Model convergence
## problem; non-positive-definite Hessian matrix. See vignette('troubleshooting')
```

```
## [1] 2.533333
##  contrast             ratio     SE    df null t.ratio p.value
##  bio / ellipse        0.905 0.0829 12387    1  -1.087  0.8866
##  bio / none           1.131 0.0793 12387    1   1.758  0.4933
##  bio / rand           0.926 0.0791 12387    1  -0.896  0.9476
##  bio / scramb         0.981 0.0863 12387    1  -0.219  0.9999
##  bio / silhouette     0.897 0.0851 12387    1  -1.148  0.8610
##  ellipse / none       1.250 0.0851 12387    1   3.271  0.0137
##  ellipse / rand       1.023 0.0774 12387    1   0.305  0.9996
##  ellipse / scramb     1.084 0.0836 12387    1   1.041  0.9042
##  ellipse / silhouette 0.991 0.0764 12387    1  -0.121  1.0000
##  none / rand          0.819 0.0548 12387    1  -2.988  0.0335
##  none / scramb        0.867 0.0440 12387    1  -2.810  0.0560
##  none / silhouette    0.793 0.0588 12387    1  -3.133  0.0215
##  rand / scramb        1.059 0.0940 12387    1   0.645  0.9876
##  rand / silhouette    0.968 0.0762 12387    1  -0.411  0.9985
##  scramb / silhouette  0.914 0.0764 12387    1  -1.072  0.8926
## 
## P value adjustment: tukey method for comparing a family of 6 estimates 
## Tests are performed on the log scale
```

```
## Warning in log(inter_eye_dist_change): NaNs produced
```

```
## Warning in log(inter_eye_dist_change): NaNs produced
```

```
## [1] 5.366667
##  contrast             ratio     SE    df null t.ratio p.value
##  bio / ellipse        0.927 0.0709 12457    1  -0.990  0.9214
##  bio / none           1.073 0.0614 12457    1   1.227  0.8240
##  bio / rand           1.060 0.0974 12457    1   0.638  0.9882
##  bio / scramb         1.034 0.0764 12457    1   0.458  0.9975
##  bio / silhouette     0.808 0.0797 12457    1  -2.164  0.2548
##  ellipse / none       1.157 0.0588 12457    1   2.870  0.0472
##  ellipse / rand       1.144 0.0836 12457    1   1.837  0.4419
##  ellipse / scramb     1.116 0.0746 12457    1   1.637  0.5735
##  ellipse / silhouette 0.871 0.0703 12457    1  -1.708  0.5266
##  none / rand          0.988 0.0712 12457    1  -0.161  1.0000
##  none / scramb        0.964 0.0503 12457    1  -0.696  0.9824
##  none / silhouette    0.753 0.0557 12457    1  -3.836  0.0018
##  rand / scramb        0.976 0.0730 12457    1  -0.330  0.9995
##  rand / silhouette    0.762 0.0697 12457    1  -2.972  0.0351
##  scramb / silhouette  0.781 0.0542 12457    1  -3.566  0.0049
## 
## P value adjustment: tukey method for comparing a family of 6 estimates 
## Tests are performed on the log scale
```

```
## Warning in log(inter_eye_dist_change): NaNs produced
## Warning in log(inter_eye_dist_change): NaNs produced
```

```
## [1] 5.866667
##  contrast             ratio     SE    df null t.ratio p.value
##  bio / ellipse        0.978 0.0605 12442    1  -0.356  0.9992
##  bio / none           1.099 0.0602 12442    1   1.731  0.5113
##  bio / rand           1.130 0.1010 12442    1   1.364  0.7485
##  bio / scramb         0.941 0.0752 12442    1  -0.764  0.9735
##  bio / silhouette     0.926 0.0748 12442    1  -0.950  0.9333
##  ellipse / none       1.124 0.0625 12442    1   2.102  0.2862
##  ellipse / rand       1.155 0.0868 12442    1   1.917  0.3913
##  ellipse / scramb     0.962 0.0781 12442    1  -0.481  0.9968
##  ellipse / silhouette 0.947 0.0790 12442    1  -0.656  0.9865
##  none / rand          1.028 0.0791 12442    1   0.355  0.9993
##  none / scramb        0.856 0.0473 12442    1  -2.818  0.0547
##  none / silhouette    0.842 0.0528 12442    1  -2.738  0.0680
##  rand / scramb        0.833 0.0788 12442    1  -1.936  0.3802
##  rand / silhouette    0.820 0.0851 12442    1  -1.915  0.3929
##  scramb / silhouette  0.984 0.0877 12442    1  -0.176  1.0000
## 
## P value adjustment: tukey method for comparing a family of 6 estimates 
## Tests are performed on the log scale
```

###### 2.2.3.1.2 Summary

```
static_results_posthoc <- static_results_posthoc[-1,]
write.csv(static_results_posthoc, paste0(tmp_path, 'interEyeDistChange_staticResultsPostHoc.csv'))

kable(static_results_posthoc)
```

|  | win\_starts | win\_span | bio.ellipse | bio.none | bio.rand | bio.scramb | bio.silhouette | ellipse.none | ellipse.rand | ellipse.scramb | ellipse.silhouette | none.rand | none.scramb | none.silhouette | rand.scramb | rand.silhouette | scramb.silhouette |
| --- | --- | --- | --- | --- | --- | --- | --- | --- | --- | --- | --- | --- | --- | --- | --- | --- | --- |
| 2 | 0.0333333 | 1 | 0.9122164 | 0.0170610 | 0.9990770 | 0.9999165 | 0.3349192 | 0.0965018 | 0.9871709 | 0.8546675 | 0.0238949 | 0.1580968 | 0.0054672 | 0.0000002 | 0.9922625 | 0.1848154 | 0.5762529 |
| 3 | 0.2000000 | 1 | 0.8738337 | 0.0011822 | 0.9998461 | 0.9999999 | 0.2850007 | 0.0500096 | 0.9630008 | 0.9411479 | 0.0294484 | 0.0347717 | 0.0032825 | 0.0000004 | 0.9997046 | 0.2036228 | 0.5340564 |
| 4 | 0.3666667 | 1 | 0.8023202 | 0.0003276 | 0.9680779 | 0.9921709 | 0.8350414 | 0.0109941 | 0.9955587 | 0.9864645 | 0.1571514 | 0.0310022 | 0.0016615 | 0.0000003 | 0.9998224 | 0.3369682 | 0.6736372 |
| 5 | 0.5333333 | 1 | 0.8440956 | 0.0000413 | 0.8857701 | 0.8236577 | 0.9162872 | 0.0024208 | 0.9999999 | 0.9999988 | 0.3747208 | 0.0147423 | 0.0053212 | 0.0000008 | 0.9999879 | 0.3912533 | 0.4640776 |
| 6 | 0.7000000 | 1 | 0.9492462 | 0.0000400 | 0.9987144 | 0.9976829 | 0.9601055 | 0.0008062 | 0.9849273 | 0.9989141 | 0.6134098 | 0.0000848 | 0.0001210 | 0.0000047 | 0.9999975 | 0.8490798 | 0.8939887 |
| 7 | 0.8666667 | 1 | 0.8407831 | 0.0000047 | 0.9999909 | 0.9375768 | 1.0000000 | 0.0006167 | 0.7814592 | 0.9999401 | 0.8783745 | 0.0000123 | 0.0002945 | 0.0000137 | 0.9555101 | 0.9999967 | 0.9509343 |
| 8 | 1.0333333 | 1 | 0.9167525 | 0.0001155 | 0.9908588 | 0.9750291 | 0.9601867 | 0.0004759 | 0.9958509 | 0.9998731 | 0.9997529 | 0.0020842 | 0.0005468 | 0.0057897 | 0.9998745 | 0.9999863 | 0.9999993 |
| 9 | 1.2000000 | 1 | 0.9963457 | 0.0002430 | 0.9996643 | 0.8432237 | 0.9999825 | 0.0026052 | 0.9998287 | 0.9711678 | 0.9989615 | 0.0018706 | 0.0236585 | 0.0008865 | 0.9550028 | 0.9999911 | 0.9075847 |
| 10 | 1.3666667 | 1 | 0.9986563 | 0.0017297 | 0.9987815 | 0.9953116 | 0.9801656 | 0.0005713 | 0.9999998 | 0.9437063 | 0.9997533 | 0.0049910 | 0.0912967 | 0.0096053 | 0.9762579 | 0.9993924 | 0.9231249 |
| 11 | 1.5333333 | 1 | 0.9956122 | 0.0092343 | 0.9851408 | 0.9917270 | 0.9807764 | 0.0011634 | 1.0000000 | 0.8663978 | 0.9999774 | 0.0034950 | 0.1964498 | 0.0237878 | 0.8904576 | 0.9999881 | 0.8929138 |
| 12 | 1.7000000 | 1 | 0.6019331 | 0.2166073 | 0.9865361 | 0.9997384 | 0.7631291 | 0.0001588 | 0.8856027 | 0.3614550 | 0.9999852 | 0.1276112 | 0.5191265 | 0.0386378 | 0.9719907 | 0.9615888 | 0.6903228 |
| 13 | 2.3666667 | 1 | 0.8654639 | 0.1481064 | 0.9999863 | 0.9981510 | 0.9905559 | 0.0012545 | 0.8455677 | 0.5383951 | 0.9922932 | 0.0652017 | 0.1774589 | 0.0431370 | 0.9912761 | 0.9937342 | 0.8971094 |
| 14 | 2.5333333 | 1 | 0.8865813 | 0.4932729 | 0.9475705 | 0.9999309 | 0.8610047 | 0.0137140 | 0.9996476 | 0.9041503 | 0.9999964 | 0.0335263 | 0.0559815 | 0.0214552 | 0.9875614 | 0.9984953 | 0.8925502 |
| 15 | 5.3666667 | 1 | 0.9213890 | 0.8239571 | 0.9881895 | 0.9974733 | 0.2547672 | 0.0472087 | 0.4419286 | 0.5735354 | 0.5265903 | 0.9999850 | 0.9824353 | 0.0017506 | 0.9994830 | 0.0351317 | 0.0048990 |
| 16 | 5.8666667 | 1 | 0.9992493 | 0.5112691 | 0.7485058 | 0.9734862 | 0.9332582 | 0.2861759 | 0.3913485 | 0.9968210 | 0.9865116 | 0.9992645 | 0.0547179 | 0.0680406 | 0.3801887 | 0.3928765 | 0.9999767 |

###### 2.2.3.2 Moving

remaking specific dataframe

```
onlymoving <- subset(full_data, full_data$stimtype != 'none')

data <- onlymoving
```

reloading of the data and ps

```
moving <- read.csv(paste0(tmp_path, 'interEyeDistChange_movingResults.csv'))

moving$adjustedp <- moving$p*sum(moving$stimtype=='bio') # create bonferroni corrected p
moving$adjustedp[moving$adjustedp>1] <- 1
significant_moving <- subset(moving, moving$stimtype=='bio') #subset to have only 1 row per model

significant_moving <- subset(significant_moving, significant_moving$adjustedp < alpha)

significant_moving_wins <- significant_moving$win_starts #identify the windows that are significant
```

###### 2.2.3.2.1 Detailed

now redo the models

```
moving_results_posthoc <- data.frame(list('win_starts'=NaN,'win_span'=NaN,
                                          'bio/ellipse'=NaN, 'bio/rand'=NaN,
                                          'bio/scramb'=NaN, 'bio/silhouette'=NaN,
                                          'ellipse/rand'=NaN, 'ellipse/scramb'=NaN,
                                          'ellipse/silhouette'=NaN, 'rand/scramb'=NaN,
                                          'rand/silhouette'=NaN, 'scramb/silhouette'=NaN))

for (window_start in significant_moving_wins){
  window_end <- window_start + window_size

  window_data <- subset(data, data$time_from_start>window_start)
  window_data <- subset(window_data, window_data$time_from_start<window_end)

  full_tmb <- glmmTMB(log(inter_eye_dist_change) ~ stimtype + (stimtype|subj), family=gaussian, data=window_data)
  Anova(full_tmb)
  e <- emmeans(full_tmb, ~stimtype, type='response')
  prs <- as.data.frame(pairs(e))
  newrow <- c(window_start, window_size, prs$p.value[1], prs$p.value[2],
              prs$p.value[3], prs$p.value[4], prs$p.value[5], prs$p.value[6],
              prs$p.value[7], prs$p.value[8], prs$p.value[9], prs$p.value[10])
  moving_results_posthoc <- rbind(moving_results_posthoc, newrow)

  print(window_start)
  print(pairs(e))
}
```

```
## Warning in log(inter_eye_dist_change): NaNs produced
## Warning in log(inter_eye_dist_change): NaNs produced
```

```
## [1] 11.86667
##  contrast             ratio     SE   df null t.ratio p.value
##  bio / ellipse        1.488 0.1310 5506    1   4.518  0.0001
##  bio / rand           0.836 0.0659 5506    1  -2.274  0.1534
##  bio / scramb         0.914 0.0620 5506    1  -1.331  0.6720
##  bio / silhouette     0.890 0.0787 5506    1  -1.312  0.6835
##  ellipse / rand       0.562 0.0488 5506    1  -6.639  <.0001
##  ellipse / scramb     0.614 0.0491 5506    1  -6.099  <.0001
##  ellipse / silhouette 0.599 0.0533 5506    1  -5.761  <.0001
##  rand / scramb        1.093 0.0767 5506    1   1.267  0.7113
##  rand / silhouette    1.065 0.1080 5506    1   0.625  0.9711
##  scramb / silhouette  0.975 0.0829 5506    1  -0.301  0.9982
## 
## P value adjustment: tukey method for comparing a family of 5 estimates 
## Tests are performed on the log scale
```

```
## Warning in log(inter_eye_dist_change): NaNs produced
## Warning in log(inter_eye_dist_change): NaNs produced
```

```
## Warning in finalizeTMB(TMBStruc, obj, fit, h, data.tmb.old): Model convergence
## problem; non-positive-definite Hessian matrix. See vignette('troubleshooting')
```

```
## Warning in finalizeTMB(TMBStruc, obj, fit, h, data.tmb.old): Model convergence
## problem; singular convergence (7). See vignette('troubleshooting'),
## help('diagnose')
```

```
## [1] 12.03333
##  contrast             ratio     SE   df null t.ratio p.value
##  bio / ellipse        1.768 0.1710 5425    1   5.909  <.0001
##  bio / rand           0.840 0.0614 5425    1  -2.380  0.1209
##  bio / scramb         0.906 0.0619 5425    1  -1.449  0.5961
##  bio / silhouette     0.872 0.0711 5425    1  -1.684  0.4439
##  ellipse / rand       0.475 0.0471 5425    1  -7.498  <.0001
##  ellipse / scramb     0.512 0.0472 5425    1  -7.259  <.0001
##  ellipse / silhouette 0.493 0.0504 5425    1  -6.916  <.0001
##  rand / scramb        1.078 0.0721 5425    1   1.120  0.7959
##  rand / silhouette    1.037 0.1020 5425    1   0.368  0.9961
##  scramb / silhouette  0.962 0.0854 5425    1  -0.434  0.9926
## 
## P value adjustment: tukey method for comparing a family of 5 estimates 
## Tests are performed on the log scale
```

```
## Warning in log(inter_eye_dist_change): NaNs produced
```

```
## Warning in log(inter_eye_dist_change): NaNs produced
```

```
## Warning in finalizeTMB(TMBStruc, obj, fit, h, data.tmb.old): Model convergence
## problem; non-positive-definite Hessian matrix. See vignette('troubleshooting')
```

```
## Warning in finalizeTMB(TMBStruc, obj, fit, h, data.tmb.old): Model convergence
## problem; singular convergence (7). See vignette('troubleshooting'),
## help('diagnose')
```

```
## [1] 12.2
##  contrast             ratio     SE   df null t.ratio p.value
##  bio / ellipse        2.491 0.2720 4980    1   8.348  <.0001
##  bio / rand           0.830 0.0679 4980    1  -2.285  0.1499
##  bio / scramb         0.937 0.0642 4980    1  -0.958  0.8740
##  bio / silhouette     0.796 0.0724 4980    1  -2.514  0.0876
##  ellipse / rand       0.333 0.0419 4980    1  -8.747  <.0001
##  ellipse / scramb     0.376 0.0455 4980    1  -8.092  <.0001
##  ellipse / silhouette 0.319 0.0350 4980    1 -10.402  <.0001
##  rand / scramb        1.129 0.0918 4980    1   1.491  0.5685
##  rand / silhouette    0.959 0.1110 4980    1  -0.362  0.9963
##  scramb / silhouette  0.849 0.0896 4980    1  -1.546  0.5322
## 
## P value adjustment: tukey method for comparing a family of 5 estimates 
## Tests are performed on the log scale
```

```
## Warning in log(inter_eye_dist_change): NaNs produced
```

```
## Warning in log(inter_eye_dist_change): NaNs produced
```

```
## Warning in finalizeTMB(TMBStruc, obj, fit, h, data.tmb.old): Model convergence
## problem; non-positive-definite Hessian matrix. See vignette('troubleshooting')
```

```
## Warning in finalizeTMB(TMBStruc, obj, fit, h, data.tmb.old): Model convergence
## problem; singular convergence (7). See vignette('troubleshooting'),
## help('diagnose')
```

```
## [1] 12.36667
##  contrast             ratio     SE   df null t.ratio p.value
##  bio / ellipse        3.049 0.3450 5039    1   9.849  <.0001
##  bio / rand           0.902 0.0875 5039    1  -1.061  0.8266
##  bio / scramb         0.937 0.0735 5039    1  -0.831  0.9210
##  bio / silhouette     0.859 0.0798 5039    1  -1.631  0.4775
##  ellipse / rand       0.296 0.0403 5039    1  -8.935  <.0001
##  ellipse / scramb     0.307 0.0432 5039    1  -8.387  <.0001
##  ellipse / silhouette 0.282 0.0317 5039    1 -11.262  <.0001
##  rand / scramb        1.038 0.0882 5039    1   0.444  0.9920
##  rand / silhouette    0.953 0.1190 5039    1  -0.388  0.9952
##  scramb / silhouette  0.917 0.1140 5039    1  -0.693  0.9580
## 
## P value adjustment: tukey method for comparing a family of 5 estimates 
## Tests are performed on the log scale
```

```
## Warning in log(inter_eye_dist_change): NaNs produced
```

```
## Warning in log(inter_eye_dist_change): NaNs produced
```

```
## Warning in finalizeTMB(TMBStruc, obj, fit, h, data.tmb.old): Model convergence
## problem; non-positive-definite Hessian matrix. See vignette('troubleshooting')
```

```
## Warning in finalizeTMB(TMBStruc, obj, fit, h, data.tmb.old): Model convergence
## problem; singular convergence (7). See vignette('troubleshooting'),
## help('diagnose')
```

```
## [1] 12.53333
##  contrast             ratio     SE   df null t.ratio p.value
##  bio / ellipse        3.157 0.3420 5042    1  10.601  <.0001
##  bio / rand           0.936 0.0931 5042    1  -0.669  0.9631
##  bio / scramb         0.949 0.0784 5042    1  -0.637  0.9690
##  bio / silhouette     0.873 0.0774 5042    1  -1.534  0.5405
##  ellipse / rand       0.296 0.0432 5042    1  -8.346  <.0001
##  ellipse / scramb     0.300 0.0426 5042    1  -8.471  <.0001
##  ellipse / silhouette 0.276 0.0305 5042    1 -11.661  <.0001
##  rand / scramb        1.014 0.0914 5042    1   0.154  0.9999
##  rand / silhouette    0.933 0.1240 5042    1  -0.525  0.9849
##  scramb / silhouette  0.920 0.1180 5042    1  -0.652  0.9663
## 
## P value adjustment: tukey method for comparing a family of 5 estimates 
## Tests are performed on the log scale
```

```
## Warning in log(inter_eye_dist_change): NaNs produced
```

```
## Warning in log(inter_eye_dist_change): NaNs produced
```

```
## [1] 12.7
##  contrast             ratio     SE   df null t.ratio p.value
##  bio / ellipse        3.104 0.3520 4690    1   9.997  <.0001
##  bio / rand           1.006 0.0983 4690    1   0.064  1.0000
##  bio / scramb         0.971 0.0844 4690    1  -0.334  0.9973
##  bio / silhouette     0.903 0.0849 4690    1  -1.081  0.8163
##  ellipse / rand       0.324 0.0477 4690    1  -7.649  <.0001
##  ellipse / scramb     0.313 0.0459 4690    1  -7.917  <.0001
##  ellipse / silhouette 0.291 0.0335 4690    1 -10.731  <.0001
##  rand / scramb        0.965 0.0783 4690    1  -0.436  0.9925
##  rand / silhouette    0.898 0.1260 4690    1  -0.767  0.9401
##  scramb / silhouette  0.930 0.1230 4690    1  -0.547  0.9824
## 
## P value adjustment: tukey method for comparing a family of 5 estimates 
## Tests are performed on the log scale
```

```
## Warning in log(inter_eye_dist_change): NaNs produced
## Warning in log(inter_eye_dist_change): NaNs produced
```

```
## [1] 12.86667
##  contrast             ratio     SE   df null t.ratio p.value
##  bio / ellipse        3.206 0.4080 4493    1   9.165  <.0001
##  bio / rand           1.119 0.1270 4493    1   0.991  0.8594
##  bio / scramb         1.066 0.1130 4493    1   0.602  0.9748
##  bio / silhouette     0.978 0.1010 4493    1  -0.217  0.9995
##  ellipse / rand       0.349 0.0544 4493    1  -6.754  <.0001
##  ellipse / scramb     0.332 0.0492 4493    1  -7.436  <.0001
##  ellipse / silhouette 0.305 0.0371 4493    1  -9.775  <.0001
##  rand / scramb        0.952 0.0921 4493    1  -0.504  0.9870
##  rand / silhouette    0.874 0.1310 4493    1  -0.899  0.8972
##  scramb / silhouette  0.918 0.1280 4493    1  -0.617  0.9725
## 
## P value adjustment: tukey method for comparing a family of 5 estimates 
## Tests are performed on the log scale
```

```
## Warning in log(inter_eye_dist_change): NaNs produced
## Warning in log(inter_eye_dist_change): NaNs produced
```

```
## [1] 13.03333
##  contrast             ratio     SE   df null t.ratio p.value
##  bio / ellipse        3.005 0.3760 4240    1   8.797  <.0001
##  bio / rand           1.044 0.1210 4240    1   0.372  0.9959
##  bio / scramb         1.006 0.1020 4240    1   0.057  1.0000
##  bio / silhouette     1.013 0.1000 4240    1   0.134  0.9999
##  ellipse / rand       0.347 0.0528 4240    1  -6.958  <.0001
##  ellipse / scramb     0.335 0.0496 4240    1  -7.381  <.0001
##  ellipse / silhouette 0.337 0.0386 4240    1  -9.500  <.0001
##  rand / scramb        0.963 0.0866 4240    1  -0.416  0.9937
##  rand / silhouette    0.971 0.1300 4240    1  -0.223  0.9995
##  scramb / silhouette  1.008 0.1290 4240    1   0.058  1.0000
## 
## P value adjustment: tukey method for comparing a family of 5 estimates 
## Tests are performed on the log scale
```

```
## Warning in log(inter_eye_dist_change): NaNs produced
## Warning in log(inter_eye_dist_change): NaNs produced
```

```
## [1] 13.2
##  contrast             ratio     SE   df null t.ratio p.value
##  bio / ellipse        2.691 0.3760 3899    1   7.093  <.0001
##  bio / rand           0.923 0.1330 3899    1  -0.555  0.9814
##  bio / scramb         0.912 0.1100 3899    1  -0.757  0.9427
##  bio / silhouette     0.978 0.1270 3899    1  -0.175  0.9998
##  ellipse / rand       0.343 0.0538 3899    1  -6.816  <.0001
##  ellipse / scramb     0.339 0.0558 3899    1  -6.576  <.0001
##  ellipse / silhouette 0.363 0.0469 3899    1  -7.847  <.0001
##  rand / scramb        0.989 0.1080 3899    1  -0.104  1.0000
##  rand / silhouette    1.059 0.1560 3899    1   0.392  0.9950
##  scramb / silhouette  1.071 0.1370 3899    1   0.540  0.9832
## 
## P value adjustment: tukey method for comparing a family of 5 estimates 
## Tests are performed on the log scale
```

```
## Warning in log(inter_eye_dist_change): NaNs produced
## Warning in log(inter_eye_dist_change): NaNs produced
```

```
## Warning in finalizeTMB(TMBStruc, obj, fit, h, data.tmb.old): Model convergence
## problem; non-positive-definite Hessian matrix. See vignette('troubleshooting')
```

```
## Warning in finalizeTMB(TMBStruc, obj, fit, h, data.tmb.old): Model convergence
## problem; singular convergence (7). See vignette('troubleshooting'),
## help('diagnose')
```

```
## [1] 13.36667
##  contrast             ratio     SE   df null t.ratio p.value
##  bio / ellipse        2.528 0.3610 3757    1   6.491  <.0001
##  bio / rand           0.870 0.1160 3757    1  -1.041  0.8361
##  bio / scramb         0.908 0.1110 3757    1  -0.792  0.9330
##  bio / silhouette     0.930 0.1300 3757    1  -0.518  0.9856
##  ellipse / rand       0.344 0.0552 3757    1  -6.653  <.0001
##  ellipse / scramb     0.359 0.0566 3757    1  -6.493  <.0001
##  ellipse / silhouette 0.368 0.0487 3757    1  -7.547  <.0001
##  rand / scramb        1.043 0.1090 3757    1   0.405  0.9944
##  rand / silhouette    1.069 0.1600 3757    1   0.446  0.9919
##  scramb / silhouette  1.025 0.1210 3757    1   0.207  0.9996
## 
## P value adjustment: tukey method for comparing a family of 5 estimates 
## Tests are performed on the log scale
```

```
## Warning in log(inter_eye_dist_change): NaNs produced
```

```
## Warning in log(inter_eye_dist_change): NaNs produced
```

```
## [1] 13.53333
##  contrast             ratio     SE   df null t.ratio p.value
##  bio / ellipse        2.393 0.3440 3717    1   6.072  <.0001
##  bio / rand           0.870 0.1200 3717    1  -1.011  0.8505
##  bio / scramb         0.907 0.1140 3717    1  -0.775  0.9378
##  bio / silhouette     0.864 0.1330 3717    1  -0.949  0.8776
##  ellipse / rand       0.364 0.0592 3717    1  -6.211  <.0001
##  ellipse / scramb     0.379 0.0591 3717    1  -6.221  <.0001
##  ellipse / silhouette 0.361 0.0512 3717    1  -7.188  <.0001
##  rand / scramb        1.042 0.1240 3717    1   0.346  0.9969
##  rand / silhouette    0.993 0.1520 3717    1  -0.045  1.0000
##  scramb / silhouette  0.953 0.1160 3717    1  -0.394  0.9949
## 
## P value adjustment: tukey method for comparing a family of 5 estimates 
## Tests are performed on the log scale
```

```
## Warning in log(inter_eye_dist_change): NaNs produced
## Warning in log(inter_eye_dist_change): NaNs produced
```

```
## Warning in finalizeTMB(TMBStruc, obj, fit, h, data.tmb.old): Model convergence
## problem; non-positive-definite Hessian matrix. See vignette('troubleshooting')
```

```
## Warning in finalizeTMB(TMBStruc, obj, fit, h, data.tmb.old): Model convergence
## problem; singular convergence (7). See vignette('troubleshooting'),
## help('diagnose')
```

```
## [1] 13.7
##  contrast             ratio     SE   df null t.ratio p.value
##  bio / ellipse        2.271 0.2840 3594    1   6.554  <.0001
##  bio / rand           0.868 0.1250 3594    1  -0.978  0.8652
##  bio / scramb         0.870 0.1100 3594    1  -1.099  0.8071
##  bio / silhouette     0.813 0.1240 3594    1  -1.354  0.6570
##  ellipse / rand       0.382 0.0656 3594    1  -5.600  <.0001
##  ellipse / scramb     0.383 0.0565 3594    1  -6.507  <.0001
##  ellipse / silhouette 0.358 0.0484 3594    1  -7.591  <.0001
##  rand / scramb        1.002 0.1260 3594    1   0.014  1.0000
##  rand / silhouette    0.936 0.1580 3594    1  -0.391  0.9951
##  scramb / silhouette  0.935 0.1090 3594    1  -0.579  0.9782
## 
## P value adjustment: tukey method for comparing a family of 5 estimates 
## Tests are performed on the log scale
```

```
## Warning in log(inter_eye_dist_change): NaNs produced
```

```
## Warning in log(inter_eye_dist_change): NaNs produced
```

```
## Warning in finalizeTMB(TMBStruc, obj, fit, h, data.tmb.old): Model convergence
## problem; non-positive-definite Hessian matrix. See vignette('troubleshooting')
```

```
## Warning in finalizeTMB(TMBStruc, obj, fit, h, data.tmb.old): Model convergence
## problem; singular convergence (7). See vignette('troubleshooting'),
## help('diagnose')
```

```
## [1] 13.86667
##  contrast             ratio     SE   df null t.ratio p.value
##  bio / ellipse        2.194 0.2840 3745    1   6.071  <.0001
##  bio / rand           0.840 0.1100 3745    1  -1.333  0.6703
##  bio / scramb         0.939 0.1230 3745    1  -0.477  0.9894
##  bio / silhouette     0.813 0.1170 3745    1  -1.446  0.5977
##  ellipse / rand       0.383 0.0682 3745    1  -5.388  <.0001
##  ellipse / scramb     0.428 0.0696 3745    1  -5.218  <.0001
##  ellipse / silhouette 0.370 0.0500 3745    1  -7.360  <.0001
##  rand / scramb        1.119 0.1420 3745    1   0.882  0.9038
##  rand / silhouette    0.968 0.1680 3745    1  -0.189  0.9997
##  scramb / silhouette  0.865 0.1040 3745    1  -1.207  0.7474
## 
## P value adjustment: tukey method for comparing a family of 5 estimates 
## Tests are performed on the log scale
```

```
## Warning in log(inter_eye_dist_change): NaNs produced
```

```
## Warning in log(inter_eye_dist_change): NaNs produced
```

```
## [1] 14.03333
##  contrast             ratio     SE   df null t.ratio p.value
##  bio / ellipse        2.000 0.2860 3765    1   4.840  <.0001
##  bio / rand           0.803 0.1130 3765    1  -1.559  0.5241
##  bio / scramb         0.959 0.1280 3765    1  -0.315  0.9979
##  bio / silhouette     0.784 0.1070 3765    1  -1.789  0.3801
##  ellipse / rand       0.401 0.0740 3765    1  -4.951  <.0001
##  ellipse / scramb     0.479 0.0809 3765    1  -4.356  0.0001
##  ellipse / silhouette 0.392 0.0577 3765    1  -6.365  <.0001
##  rand / scramb        1.194 0.1730 3765    1   1.225  0.7367
##  rand / silhouette    0.976 0.1710 3765    1  -0.137  0.9999
##  scramb / silhouette  0.817 0.0975 3765    1  -1.689  0.4406
## 
## P value adjustment: tukey method for comparing a family of 5 estimates 
## Tests are performed on the log scale
```

```
## Warning in log(inter_eye_dist_change): NaNs produced
## Warning in log(inter_eye_dist_change): NaNs produced
```

```
## [1] 14.2
##  contrast             ratio     SE   df null t.ratio p.value
##  bio / ellipse        2.077 0.2800 3675    1   5.427  <.0001
##  bio / rand           0.789 0.1190 3675    1  -1.575  0.5138
##  bio / scramb         0.948 0.1210 3675    1  -0.416  0.9937
##  bio / silhouette     0.805 0.0965 3675    1  -1.807  0.3693
##  ellipse / rand       0.380 0.0728 3675    1  -5.053  <.0001
##  ellipse / scramb     0.457 0.0780 3675    1  -4.588  <.0001
##  ellipse / silhouette 0.388 0.0628 3675    1  -5.850  <.0001
##  rand / scramb        1.203 0.1800 3675    1   1.233  0.7318
##  rand / silhouette    1.021 0.1880 3675    1   0.113  1.0000
##  scramb / silhouette  0.849 0.1100 3675    1  -1.258  0.7173
## 
## P value adjustment: tukey method for comparing a family of 5 estimates 
## Tests are performed on the log scale
```

```
## Warning in log(inter_eye_dist_change): NaNs produced
## Warning in log(inter_eye_dist_change): NaNs produced
```

```
## [1] 14.36667
##  contrast             ratio     SE   df null t.ratio p.value
##  bio / ellipse        2.053 0.2810 3881    1   5.261  <.0001
##  bio / rand           0.773 0.1090 3881    1  -1.821  0.3616
##  bio / scramb         0.931 0.1110 3881    1  -0.600  0.9752
##  bio / silhouette     0.779 0.0970 3881    1  -2.003  0.2647
##  ellipse / rand       0.377 0.0640 3881    1  -5.745  <.0001
##  ellipse / scramb     0.454 0.0751 3881    1  -4.778  <.0001
##  ellipse / silhouette 0.380 0.0639 3881    1  -5.754  <.0001
##  rand / scramb        1.204 0.1810 3881    1   1.232  0.7328
##  rand / silhouette    1.008 0.1560 3881    1   0.051  1.0000
##  scramb / silhouette  0.837 0.1040 3881    1  -1.426  0.6107
## 
## P value adjustment: tukey method for comparing a family of 5 estimates 
## Tests are performed on the log scale
```

```
## Warning in log(inter_eye_dist_change): NaNs produced
## Warning in log(inter_eye_dist_change): NaNs produced
```

```
## [1] 14.53333
##  contrast             ratio     SE   df null t.ratio p.value
##  bio / ellipse        1.962 0.2540 3885    1   5.194  <.0001
##  bio / rand           0.697 0.0934 3885    1  -2.694  0.0550
##  bio / scramb         0.889 0.1080 3885    1  -0.970  0.8686
##  bio / silhouette     0.723 0.0789 3885    1  -2.972  0.0248
##  ellipse / rand       0.355 0.0602 3885    1  -6.109  <.0001
##  ellipse / scramb     0.453 0.0749 3885    1  -4.792  <.0001
##  ellipse / silhouette 0.369 0.0580 3885    1  -6.348  <.0001
##  rand / scramb        1.275 0.1940 3885    1   1.595  0.5005
##  rand / silhouette    1.038 0.1390 3885    1   0.276  0.9987
##  scramb / silhouette  0.814 0.1040 3885    1  -1.612  0.4896
## 
## P value adjustment: tukey method for comparing a family of 5 estimates 
## Tests are performed on the log scale
```

```
## Warning in log(inter_eye_dist_change): NaNs produced
## Warning in log(inter_eye_dist_change): NaNs produced
```

```
## Warning in finalizeTMB(TMBStruc, obj, fit, h, data.tmb.old): Model convergence
## problem; non-positive-definite Hessian matrix. See vignette('troubleshooting')
```

```
## Warning in finalizeTMB(TMBStruc, obj, fit, h, data.tmb.old): Model convergence
## problem; singular convergence (7). See vignette('troubleshooting'),
## help('diagnose')
```

```
## [1] 14.7
##  contrast             ratio     SE   df null t.ratio p.value
##  bio / ellipse        1.918 0.2340 3717    1   5.342  <.0001
##  bio / rand           0.650 0.0728 3717    1  -3.848  0.0011
##  bio / scramb         0.889 0.1190 3717    1  -0.882  0.9037
##  bio / silhouette     0.707 0.0718 3717    1  -3.414  0.0058
##  ellipse / rand       0.339 0.0513 3717    1  -7.148  <.0001
##  ellipse / scramb     0.463 0.0724 3717    1  -4.924  <.0001
##  ellipse / silhouette 0.369 0.0565 3717    1  -6.511  <.0001
##  rand / scramb        1.367 0.2060 3717    1   2.077  0.2303
##  rand / silhouette    1.088 0.1100 3717    1   0.834  0.9200
##  scramb / silhouette  0.795 0.1070 3717    1  -1.707  0.4297
## 
## P value adjustment: tukey method for comparing a family of 5 estimates 
## Tests are performed on the log scale
```

```
## Warning in log(inter_eye_dist_change): NaNs produced
```

```
## Warning in log(inter_eye_dist_change): NaNs produced
```

```
## Warning in finalizeTMB(TMBStruc, obj, fit, h, data.tmb.old): Model convergence
## problem; non-positive-definite Hessian matrix. See vignette('troubleshooting')
```

```
## Warning in finalizeTMB(TMBStruc, obj, fit, h, data.tmb.old): Model convergence
## problem; singular convergence (7). See vignette('troubleshooting'),
## help('diagnose')
```

```
## [1] 14.86667
##  contrast             ratio     SE   df null t.ratio p.value
##  bio / ellipse        1.781 0.2230 3851    1   4.605  <.0001
##  bio / rand           0.619 0.0811 3851    1  -3.659  0.0024
##  bio / scramb         0.811 0.1140 3851    1  -1.490  0.5692
##  bio / silhouette     0.674 0.0725 3851    1  -3.666  0.0023
##  ellipse / rand       0.348 0.0509 3851    1  -7.211  <.0001
##  ellipse / scramb     0.455 0.0704 3851    1  -5.090  <.0001
##  ellipse / silhouette 0.378 0.0560 3851    1  -6.563  <.0001
##  rand / scramb        1.310 0.2070 3851    1   1.707  0.4298
##  rand / silhouette    1.089 0.1100 3851    1   0.842  0.9175
##  scramb / silhouette  0.831 0.1220 3851    1  -1.265  0.7127
## 
## P value adjustment: tukey method for comparing a family of 5 estimates 
## Tests are performed on the log scale
```

```
## Warning in log(inter_eye_dist_change): NaNs produced
```

```
## Warning in log(inter_eye_dist_change): NaNs produced
```

```
## Warning in finalizeTMB(TMBStruc, obj, fit, h, data.tmb.old): Model convergence
## problem; non-positive-definite Hessian matrix. See vignette('troubleshooting')
```

```
## Warning in finalizeTMB(TMBStruc, obj, fit, h, data.tmb.old): Model convergence
## problem; singular convergence (7). See vignette('troubleshooting'),
## help('diagnose')
```

```
## [1] 15.03333
##  contrast             ratio     SE   df null t.ratio p.value
##  bio / ellipse        1.703 0.1930 3880    1   4.712  <.0001
##  bio / rand           0.598 0.0702 3880    1  -4.384  0.0001
##  bio / scramb         0.763 0.1060 3880    1  -1.955  0.2887
##  bio / silhouette     0.612 0.0681 3880    1  -4.411  0.0001
##  ellipse / rand       0.351 0.0439 3880    1  -8.368  <.0001
##  ellipse / scramb     0.448 0.0665 3880    1  -5.408  <.0001
##  ellipse / silhouette 0.359 0.0480 3880    1  -7.665  <.0001
##  rand / scramb        1.277 0.1930 3880    1   1.616  0.4873
##  rand / silhouette    1.024 0.0958 3880    1   0.253  0.9991
##  scramb / silhouette  0.802 0.1140 3880    1  -1.547  0.5319
## 
## P value adjustment: tukey method for comparing a family of 5 estimates 
## Tests are performed on the log scale
```

```
## Warning in log(inter_eye_dist_change): NaNs produced
```

```
## Warning in log(inter_eye_dist_change): NaNs produced
```

```
## Warning in finalizeTMB(TMBStruc, obj, fit, h, data.tmb.old): Model convergence
## problem; non-positive-definite Hessian matrix. See vignette('troubleshooting')
```

```
## Warning in finalizeTMB(TMBStruc, obj, fit, h, data.tmb.old): Model convergence
## problem; singular convergence (7). See vignette('troubleshooting'),
## help('diagnose')
```

```
## [1] 15.2
##  contrast             ratio     SE   df null t.ratio p.value
##  bio / ellipse        1.553 0.2000 3779    1   3.416  0.0058
##  bio / rand           0.528 0.0849 3779    1  -3.970  0.0007
##  bio / scramb         0.765 0.0958 3779    1  -2.136  0.2050
##  bio / silhouette     0.583 0.0706 3779    1  -4.459  0.0001
##  ellipse / rand       0.340 0.0459 3779    1  -7.993  <.0001
##  ellipse / scramb     0.493 0.0678 3779    1  -5.146  <.0001
##  ellipse / silhouette 0.375 0.0474 3779    1  -7.768  <.0001
##  rand / scramb        1.449 0.2220 3779    1   2.424  0.1092
##  rand / silhouette    1.103 0.1290 3779    1   0.843  0.9172
##  scramb / silhouette  0.761 0.1020 3779    1  -2.039  0.2473
## 
## P value adjustment: tukey method for comparing a family of 5 estimates 
## Tests are performed on the log scale
```

```
## Warning in log(inter_eye_dist_change): NaNs produced
```

```
## Warning in log(inter_eye_dist_change): NaNs produced
```

```
## Warning in finalizeTMB(TMBStruc, obj, fit, h, data.tmb.old): Model convergence
## problem; non-positive-definite Hessian matrix. See vignette('troubleshooting')
```

```
## Warning in finalizeTMB(TMBStruc, obj, fit, h, data.tmb.old): Model convergence
## problem; singular convergence (7). See vignette('troubleshooting'),
## help('diagnose')
```

```
## [1] 15.36667
##  contrast             ratio     SE   df null t.ratio p.value
##  bio / ellipse        1.433 0.1720 3971    1   2.998  0.0230
##  bio / rand           0.510 0.0778 3971    1  -4.413  0.0001
##  bio / scramb         0.738 0.0737 3971    1  -3.046  0.0198
##  bio / silhouette     0.596 0.0594 3971    1  -5.190  <.0001
##  ellipse / rand       0.356 0.0471 3971    1  -7.805  <.0001
##  ellipse / scramb     0.515 0.0618 3971    1  -5.531  <.0001
##  ellipse / silhouette 0.416 0.0484 3971    1  -7.535  <.0001
##  rand / scramb        1.446 0.2060 3971    1   2.593  0.0718
##  rand / silhouette    1.168 0.1440 3971    1   1.258  0.7168
##  scramb / silhouette  0.808 0.0934 3971    1  -1.843  0.3487
## 
## P value adjustment: tukey method for comparing a family of 5 estimates 
## Tests are performed on the log scale
```

```
## Warning in log(inter_eye_dist_change): NaNs produced
```

```
## Warning in log(inter_eye_dist_change): NaNs produced
```

```
## Warning in finalizeTMB(TMBStruc, obj, fit, h, data.tmb.old): Model convergence
## problem; non-positive-definite Hessian matrix. See vignette('troubleshooting')
```

```
## Warning in finalizeTMB(TMBStruc, obj, fit, h, data.tmb.old): Model convergence
## problem; singular convergence (7). See vignette('troubleshooting'),
## help('diagnose')
```

```
## [1] 15.53333
##  contrast             ratio     SE   df null t.ratio p.value
##  bio / ellipse        1.469 0.1570 3980    1   3.599  0.0030
##  bio / rand           0.552 0.0699 3980    1  -4.692  <.0001
##  bio / scramb         0.743 0.0645 3980    1  -3.419  0.0057
##  bio / silhouette     0.667 0.0620 3980    1  -4.360  0.0001
##  ellipse / rand       0.376 0.0463 3980    1  -7.943  <.0001
##  ellipse / scramb     0.506 0.0541 3980    1  -6.377  <.0001
##  ellipse / silhouette 0.454 0.0539 3980    1  -6.650  <.0001
##  rand / scramb        1.346 0.1720 3980    1   2.320  0.1388
##  rand / silhouette    1.207 0.1420 3980    1   1.592  0.5029
##  scramb / silhouette  0.897 0.0972 3980    1  -1.006  0.8529
## 
## P value adjustment: tukey method for comparing a family of 5 estimates 
## Tests are performed on the log scale
```

```
## Warning in log(inter_eye_dist_change): NaNs produced
```

```
## Warning in log(inter_eye_dist_change): NaNs produced
```

```
## [1] 15.7
##  contrast             ratio     SE   df null t.ratio p.value
##  bio / ellipse        1.481 0.1490 3929    1   3.917  0.0009
##  bio / rand           0.647 0.0570 3929    1  -4.939  <.0001
##  bio / scramb         0.738 0.0673 3929    1  -3.330  0.0078
##  bio / silhouette     0.722 0.0772 3929    1  -3.052  0.0194
##  ellipse / rand       0.437 0.0506 3929    1  -7.151  <.0001
##  ellipse / scramb     0.498 0.0519 3929    1  -6.687  <.0001
##  ellipse / silhouette 0.487 0.0648 3929    1  -5.410  <.0001
##  rand / scramb        1.141 0.1250 3929    1   1.199  0.7520
##  rand / silhouette    1.115 0.1250 3929    1   0.974  0.8671
##  scramb / silhouette  0.978 0.1140 3929    1  -0.193  0.9997
## 
## P value adjustment: tukey method for comparing a family of 5 estimates 
## Tests are performed on the log scale
```

```
## Warning in log(inter_eye_dist_change): NaNs produced
## Warning in log(inter_eye_dist_change): NaNs produced
```

```
## [1] 15.86667
##  contrast             ratio     SE   df null t.ratio p.value
##  bio / ellipse        1.386 0.1270 4126    1   3.557  0.0035
##  bio / rand           0.651 0.0682 4126    1  -4.095  0.0004
##  bio / scramb         0.735 0.0584 4126    1  -3.882  0.0010
##  bio / silhouette     0.757 0.0806 4126    1  -2.613  0.0681
##  ellipse / rand       0.470 0.0586 4126    1  -6.060  <.0001
##  ellipse / scramb     0.530 0.0507 4126    1  -6.641  <.0001
##  ellipse / silhouette 0.546 0.0668 4126    1  -4.944  <.0001
##  rand / scramb        1.128 0.1230 4126    1   1.104  0.8047
##  rand / silhouette    1.162 0.1730 4126    1   1.007  0.8521
##  scramb / silhouette  1.031 0.1180 4126    1   0.263  0.9989
## 
## P value adjustment: tukey method for comparing a family of 5 estimates 
## Tests are performed on the log scale
```

```
## Warning in log(inter_eye_dist_change): NaNs produced
## Warning in log(inter_eye_dist_change): NaNs produced
```

```
## [1] 16.03333
##  contrast             ratio     SE   df null t.ratio p.value
##  bio / ellipse        1.290 0.1210 4146    1   2.722  0.0509
##  bio / rand           0.625 0.0705 4146    1  -4.166  0.0003
##  bio / scramb         0.727 0.0631 4146    1  -3.668  0.0023
##  bio / silhouette     0.666 0.0822 4146    1  -3.292  0.0089
##  ellipse / rand       0.485 0.0641 4146    1  -5.480  <.0001
##  ellipse / scramb     0.564 0.0580 4146    1  -5.571  <.0001
##  ellipse / silhouette 0.516 0.0651 4146    1  -5.242  <.0001
##  rand / scramb        1.163 0.1390 4146    1   1.264  0.7133
##  rand / silhouette    1.066 0.1580 4146    1   0.430  0.9929
##  scramb / silhouette  0.916 0.1070 4146    1  -0.751  0.9442
## 
## P value adjustment: tukey method for comparing a family of 5 estimates 
## Tests are performed on the log scale
```

```
## Warning in log(inter_eye_dist_change): NaNs produced
## Warning in log(inter_eye_dist_change): NaNs produced
```

```
## [1] 16.2
##  contrast             ratio     SE   df null t.ratio p.value
##  bio / ellipse        1.217 0.1310 4086    1   1.825  0.3589
##  bio / rand           0.637 0.0694 4086    1  -4.138  0.0003
##  bio / scramb         0.714 0.0692 4086    1  -3.479  0.0046
##  bio / silhouette     0.628 0.0834 4086    1  -3.504  0.0042
##  ellipse / rand       0.523 0.0605 4086    1  -5.599  <.0001
##  ellipse / scramb     0.586 0.0629 4086    1  -4.977  <.0001
##  ellipse / silhouette 0.516 0.0655 4086    1  -5.210  <.0001
##  rand / scramb        1.121 0.1260 4086    1   1.011  0.8505
##  rand / silhouette    0.986 0.1130 4086    1  -0.122  0.9999
##  scramb / silhouette  0.880 0.1110 4086    1  -1.011  0.8505
## 
## P value adjustment: tukey method for comparing a family of 5 estimates 
## Tests are performed on the log scale
```

```
## Warning in log(inter_eye_dist_change): NaNs produced
## Warning in log(inter_eye_dist_change): NaNs produced
```

```
## [1] 16.36667
##  contrast             ratio     SE   df null t.ratio p.value
##  bio / ellipse        1.260 0.1450 4277    1   2.015  0.2588
##  bio / rand           0.663 0.0787 4277    1  -3.460  0.0049
##  bio / scramb         0.766 0.0825 4277    1  -2.476  0.0962
##  bio / silhouette     0.668 0.0855 4277    1  -3.151  0.0141
##  ellipse / rand       0.526 0.0637 4277    1  -5.301  <.0001
##  ellipse / scramb     0.608 0.0655 4277    1  -4.620  <.0001
##  ellipse / silhouette 0.530 0.0575 4277    1  -5.848  <.0001
##  rand / scramb        1.155 0.1330 4277    1   1.247  0.7238
##  rand / silhouette    1.007 0.1180 4277    1   0.061  1.0000
##  scramb / silhouette  0.872 0.1010 4277    1  -1.179  0.7633
## 
## P value adjustment: tukey method for comparing a family of 5 estimates 
## Tests are performed on the log scale
```

```
## Warning in log(inter_eye_dist_change): NaNs produced
## Warning in log(inter_eye_dist_change): NaNs produced
```

```
## [1] 16.53333
##  contrast             ratio     SE   df null t.ratio p.value
##  bio / ellipse        1.233 0.1410 4344    1   1.832  0.3549
##  bio / rand           0.691 0.0795 4344    1  -3.210  0.0117
##  bio / scramb         0.805 0.0855 4344    1  -2.045  0.2447
##  bio / silhouette     0.660 0.0731 4344    1  -3.753  0.0017
##  ellipse / rand       0.561 0.0683 4344    1  -4.750  <.0001
##  ellipse / scramb     0.653 0.0673 4344    1  -4.142  0.0003
##  ellipse / silhouette 0.535 0.0538 4344    1  -6.217  <.0001
##  rand / scramb        1.164 0.1330 4344    1   1.328  0.6737
##  rand / silhouette    0.954 0.1020 4344    1  -0.438  0.9924
##  scramb / silhouette  0.820 0.0886 4344    1  -1.840  0.3505
## 
## P value adjustment: tukey method for comparing a family of 5 estimates 
## Tests are performed on the log scale
```

```
## Warning in log(inter_eye_dist_change): NaNs produced
## Warning in log(inter_eye_dist_change): NaNs produced
```

```
## [1] 16.7
##  contrast             ratio     SE   df null t.ratio p.value
##  bio / ellipse        1.290 0.1360 4262    1   2.427  0.1083
##  bio / rand           0.747 0.0930 4262    1  -2.340  0.1325
##  bio / scramb         0.867 0.0937 4262    1  -1.321  0.6783
##  bio / silhouette     0.649 0.0766 4262    1  -3.666  0.0023
##  ellipse / rand       0.579 0.0623 4262    1  -5.081  <.0001
##  ellipse / scramb     0.672 0.0728 4262    1  -3.671  0.0023
##  ellipse / silhouette 0.503 0.0487 4262    1  -7.105  <.0001
##  rand / scramb        1.160 0.1290 4262    1   1.337  0.6680
##  rand / silhouette    0.868 0.1010 4262    1  -1.215  0.7424
##  scramb / silhouette  0.748 0.0849 4262    1  -2.556  0.0788
## 
## P value adjustment: tukey method for comparing a family of 5 estimates 
## Tests are performed on the log scale
```

```
## Warning in log(inter_eye_dist_change): NaNs produced
## Warning in log(inter_eye_dist_change): NaNs produced
```

```
## [1] 16.86667
##  contrast             ratio     SE   df null t.ratio p.value
##  bio / ellipse        1.297 0.1370 4464    1   2.461  0.0999
##  bio / rand           0.758 0.0956 4464    1  -2.193  0.1823
##  bio / scramb         0.848 0.0973 4464    1  -1.433  0.6064
##  bio / silhouette     0.641 0.0831 4464    1  -3.433  0.0054
##  ellipse / rand       0.585 0.0600 4464    1  -5.233  <.0001
##  ellipse / scramb     0.654 0.0734 4464    1  -3.782  0.0015
##  ellipse / silhouette 0.494 0.0555 4464    1  -6.273  <.0001
##  rand / scramb        1.119 0.1300 4464    1   0.964  0.8713
##  rand / silhouette    0.845 0.1010 4464    1  -1.413  0.6196
##  scramb / silhouette  0.755 0.0868 4464    1  -2.444  0.1040
## 
## P value adjustment: tukey method for comparing a family of 5 estimates 
## Tests are performed on the log scale
```

```
## Warning in log(inter_eye_dist_change): NaNs produced
## Warning in log(inter_eye_dist_change): NaNs produced
```

```
## [1] 17.03333
##  contrast             ratio     SE   df null t.ratio p.value
##  bio / ellipse        1.306 0.1370 4526    1   2.546  0.0809
##  bio / rand           0.773 0.0939 4526    1  -2.117  0.2131
##  bio / scramb         0.860 0.0941 4526    1  -1.378  0.6418
##  bio / silhouette     0.644 0.0851 4526    1  -3.327  0.0079
##  ellipse / rand       0.592 0.0651 4526    1  -4.766  <.0001
##  ellipse / scramb     0.658 0.0697 4526    1  -3.950  0.0008
##  ellipse / silhouette 0.493 0.0561 4526    1  -6.218  <.0001
##  rand / scramb        1.112 0.1200 4526    1   0.980  0.8642
##  rand / silhouette    0.833 0.0913 4526    1  -1.666  0.4555
##  scramb / silhouette  0.749 0.0895 4526    1  -2.417  0.1110
## 
## P value adjustment: tukey method for comparing a family of 5 estimates 
## Tests are performed on the log scale
```

```
## Warning in log(inter_eye_dist_change): NaNs produced
## Warning in log(inter_eye_dist_change): NaNs produced
```

```
## [1] 17.2
##  contrast             ratio     SE   df null t.ratio p.value
##  bio / ellipse        1.290 0.1320 4390    1   2.497  0.0915
##  bio / rand           0.746 0.0970 4390    1  -2.255  0.1598
##  bio / scramb         0.861 0.0874 4390    1  -1.473  0.5800
##  bio / silhouette     0.671 0.0876 4390    1  -3.053  0.0193
##  ellipse / rand       0.578 0.0687 4390    1  -4.615  <.0001
##  ellipse / scramb     0.668 0.0698 4390    1  -3.864  0.0011
##  ellipse / silhouette 0.520 0.0658 4390    1  -5.167  <.0001
##  rand / scramb        1.155 0.1350 4390    1   1.230  0.7339
##  rand / silhouette    0.900 0.1050 4390    1  -0.905  0.8950
##  scramb / silhouette  0.780 0.0927 4390    1  -2.096  0.2221
## 
## P value adjustment: tukey method for comparing a family of 5 estimates 
## Tests are performed on the log scale
```

```
## Warning in log(inter_eye_dist_change): NaNs produced
## Warning in log(inter_eye_dist_change): NaNs produced
```

```
## Warning in finalizeTMB(TMBStruc, obj, fit, h, data.tmb.old): Model convergence
## problem; non-positive-definite Hessian matrix. See vignette('troubleshooting')
```

```
## Warning in finalizeTMB(TMBStruc, obj, fit, h, data.tmb.old): Model convergence
## problem; false convergence (8). See vignette('troubleshooting'),
## help('diagnose')
```

```
## [1] 22.7
##  contrast             ratio     SE   df null t.ratio p.value
##  bio / ellipse        1.192 0.0684 5472    1   3.065  0.0186
##  bio / rand           0.863 0.0661 5472    1  -1.926  0.3036
##  bio / scramb         0.797 0.0639 5472    1  -2.826  0.0380
##  bio / silhouette     0.964 0.0708 5472    1  -0.500  0.9873
##  ellipse / rand       0.724 0.0616 5472    1  -3.797  0.0014
##  ellipse / scramb     0.669 0.0543 5472    1  -4.958  <.0001
##  ellipse / silhouette 0.809 0.0657 5472    1  -2.614  0.0679
##  rand / scramb        0.924 0.0830 5472    1  -0.879  0.9047
##  rand / silhouette    1.117 0.0735 5472    1   1.686  0.4426
##  scramb / silhouette  1.209 0.0985 5472    1   2.329  0.1360
## 
## P value adjustment: tukey method for comparing a family of 5 estimates 
## Tests are performed on the log scale
```

```
## Warning in log(inter_eye_dist_change): NaNs produced
```

```
## Warning in log(inter_eye_dist_change): NaNs produced
```

```
## Warning in finalizeTMB(TMBStruc, obj, fit, h, data.tmb.old): Model convergence
## problem; non-positive-definite Hessian matrix. See vignette('troubleshooting')
```

```
## Warning in finalizeTMB(TMBStruc, obj, fit, h, data.tmb.old): Model convergence
## problem; singular convergence (7). See vignette('troubleshooting'),
## help('diagnose')
```

```
## [1] 22.86667
##  contrast             ratio     SE   df null t.ratio p.value
##  bio / ellipse        1.186 0.0651 5721    1   3.103  0.0165
##  bio / rand           0.897 0.0631 5721    1  -1.550  0.5301
##  bio / scramb         0.825 0.0571 5721    1  -2.779  0.0434
##  bio / silhouette     0.947 0.0640 5721    1  -0.803  0.9298
##  ellipse / rand       0.756 0.0592 5721    1  -3.570  0.0033
##  ellipse / scramb     0.696 0.0481 5721    1  -5.247  <.0001
##  ellipse / silhouette 0.799 0.0568 5721    1  -3.157  0.0139
##  rand / scramb        0.920 0.0773 5721    1  -0.991  0.8597
##  rand / silhouette    1.056 0.0754 5721    1   0.769  0.9396
##  scramb / silhouette  1.148 0.0850 5721    1   1.864  0.3369
## 
## P value adjustment: tukey method for comparing a family of 5 estimates 
## Tests are performed on the log scale
```

```
## Warning in log(inter_eye_dist_change): NaNs produced
```

```
## Warning in log(inter_eye_dist_change): NaNs produced
```

```
## [1] 23.03333
##  contrast             ratio     SE   df null t.ratio p.value
##  bio / ellipse        1.122 0.0679 5704    1   1.904  0.3153
##  bio / rand           0.886 0.0610 5704    1  -1.766  0.3940
##  bio / scramb         0.814 0.0565 5704    1  -2.962  0.0255
##  bio / silhouette     0.904 0.0718 5704    1  -1.272  0.7087
##  ellipse / rand       0.789 0.0614 5704    1  -3.042  0.0200
##  ellipse / scramb     0.726 0.0512 5704    1  -4.549  0.0001
##  ellipse / silhouette 0.806 0.0633 5704    1  -2.752  0.0469
##  rand / scramb        0.920 0.0748 5704    1  -1.032  0.8406
##  rand / silhouette    1.021 0.0835 5704    1   0.251  0.9991
##  scramb / silhouette  1.110 0.0939 5704    1   1.235  0.7306
## 
## P value adjustment: tukey method for comparing a family of 5 estimates 
## Tests are performed on the log scale
```

```
## Warning in log(inter_eye_dist_change): NaNs produced
## Warning in log(inter_eye_dist_change): NaNs produced
```

```
## [1] 23.2
##  contrast             ratio     SE   df null t.ratio p.value
##  bio / ellipse        1.019 0.0701 5558    1   0.269  0.9989
##  bio / rand           0.842 0.0570 5558    1  -2.541  0.0819
##  bio / scramb         0.763 0.0623 5558    1  -3.309  0.0084
##  bio / silhouette     0.795 0.0624 5558    1  -2.929  0.0282
##  ellipse / rand       0.826 0.0684 5558    1  -2.305  0.1435
##  ellipse / scramb     0.749 0.0625 5558    1  -3.463  0.0049
##  ellipse / silhouette 0.780 0.0610 5558    1  -3.175  0.0131
##  rand / scramb        0.906 0.0715 5558    1  -1.245  0.7247
##  rand / silhouette    0.944 0.0932 5558    1  -0.585  0.9773
##  scramb / silhouette  1.041 0.1050 5558    1   0.402  0.9945
## 
## P value adjustment: tukey method for comparing a family of 5 estimates 
## Tests are performed on the log scale
```

```
## Warning in log(inter_eye_dist_change): NaNs produced
## Warning in log(inter_eye_dist_change): NaNs produced
```

```
## Warning in finalizeTMB(TMBStruc, obj, fit, h, data.tmb.old): Model convergence
## problem; non-positive-definite Hessian matrix. See vignette('troubleshooting')
```

```
## Warning in finalizeTMB(TMBStruc, obj, fit, h, data.tmb.old): Model convergence
## problem; singular convergence (7). See vignette('troubleshooting'),
## help('diagnose')
```

```
## [1] 23.7
##  contrast             ratio     SE   df null t.ratio p.value
##  bio / ellipse        0.961 0.0590 5737    1  -0.648  0.9670
##  bio / rand           0.776 0.0532 5737    1  -3.703  0.0020
##  bio / scramb         0.794 0.0565 5737    1  -3.240  0.0105
##  bio / silhouette     0.721 0.0568 5737    1  -4.151  0.0003
##  ellipse / rand       0.807 0.0634 5737    1  -2.726  0.0503
##  ellipse / scramb     0.826 0.0545 5737    1  -2.896  0.0310
##  ellipse / silhouette 0.750 0.0644 5737    1  -3.345  0.0074
##  rand / scramb        1.023 0.0830 5737    1   0.283  0.9986
##  rand / silhouette    0.929 0.0797 5737    1  -0.854  0.9135
##  scramb / silhouette  0.908 0.0849 5737    1  -1.030  0.8417
## 
## P value adjustment: tukey method for comparing a family of 5 estimates 
## Tests are performed on the log scale
```

###### 2.2.3.2.2 Summary

```
moving_results_posthoc <- moving_results_posthoc[-1,]
write.csv(moving_results_posthoc, paste0(tmp_path, 'interEyeDistChange_movingResultsPostHoc.csv'))
kable(moving_results_posthoc)
```

|  | win\_starts | win\_span | bio.ellipse | bio.rand | bio.scramb | bio.silhouette | ellipse.rand | ellipse.scramb | ellipse.silhouette | rand.scramb | rand.silhouette | scramb.silhouette |
| --- | --- | --- | --- | --- | --- | --- | --- | --- | --- | --- | --- | --- |
| 2 | 11.86667 | 1 | 0.0000625 | 0.1534138 | 0.6719668 | 0.6834571 | 0.0000000 | 0.0000000 | 0.0000001 | 0.7113383 | 0.9710770 | 0.9982067 |
| 3 | 12.03333 | 1 | 0.0000000 | 0.1209482 | 0.5961292 | 0.4438747 | 0.0000000 | 0.0000000 | 0.0000000 | 0.7959257 | 0.9960933 | 0.9926391 |
| 4 | 12.20000 | 1 | 0.0000000 | 0.1499212 | 0.8740168 | 0.0875872 | 0.0000000 | 0.0000000 | 0.0000000 | 0.5685163 | 0.9963386 | 0.5322285 |
| 5 | 12.36667 | 1 | 0.0000000 | 0.8265526 | 0.9210221 | 0.4775375 | 0.0000000 | 0.0000000 | 0.0000000 | 0.9919608 | 0.9951907 | 0.9579731 |
| 6 | 12.53333 | 1 | 0.0000000 | 0.9631019 | 0.9689530 | 0.5405221 | 0.0000000 | 0.0000000 | 0.0000000 | 0.9998751 | 0.9848708 | 0.9663156 |
| 7 | 12.70000 | 1 | 0.0000000 | 0.9999961 | 0.9973188 | 0.8162556 | 0.0000000 | 0.0000000 | 0.0000000 | 0.9925266 | 0.9400816 | 0.9823843 |
| 8 | 12.86667 | 1 | 0.0000000 | 0.8593978 | 0.9748175 | 0.9995108 | 0.0000000 | 0.0000000 | 0.0000000 | 0.9869750 | 0.8972136 | 0.9724883 |
| 9 | 13.03333 | 1 | 0.0000000 | 0.9959206 | 0.9999976 | 0.9999276 | 0.0000000 | 0.0000000 | 0.0000000 | 0.9937244 | 0.9994548 | 0.9999974 |
| 10 | 13.20000 | 1 | 0.0000000 | 0.9813893 | 0.9426973 | 0.9997918 | 0.0000000 | 0.0000000 | 0.0000000 | 0.9999736 | 0.9950376 | 0.9831656 |
| 11 | 13.36667 | 1 | 0.0000000 | 0.8361167 | 0.9330236 | 0.9856064 | 0.0000000 | 0.0000000 | 0.0000000 | 0.9943753 | 0.9918529 | 0.9995893 |
| 12 | 13.53333 | 1 | 0.0000000 | 0.8504838 | 0.9377945 | 0.8775515 | 0.0000000 | 0.0000000 | 0.0000000 | 0.9969380 | 0.9999990 | 0.9949025 |
| 13 | 13.70000 | 1 | 0.0000000 | 0.8651732 | 0.8070965 | 0.6569802 | 0.0000002 | 0.0000000 | 0.0000000 | 1.0000000 | 0.9950817 | 0.9781998 |
| 14 | 13.86667 | 1 | 0.0000000 | 0.6702740 | 0.9894221 | 0.5977177 | 0.0000008 | 0.0000019 | 0.0000000 | 0.9037807 | 0.9997134 | 0.7474114 |
| 15 | 14.03333 | 1 | 0.0000134 | 0.5241035 | 0.9978577 | 0.3801339 | 0.0000077 | 0.0001328 | 0.0000000 | 0.7366660 | 0.9999214 | 0.4405565 |
| 16 | 14.20000 | 1 | 0.0000006 | 0.5137584 | 0.9937282 | 0.3693170 | 0.0000045 | 0.0000455 | 0.0000001 | 0.7318152 | 0.9999627 | 0.7173067 |
| 17 | 14.36667 | 1 | 0.0000015 | 0.3615763 | 0.9751538 | 0.2646673 | 0.0000001 | 0.0000181 | 0.0000001 | 0.7327903 | 0.9999984 | 0.6106966 |
| 18 | 14.53333 | 1 | 0.0000022 | 0.0549673 | 0.8685723 | 0.0248440 | 0.0000000 | 0.0000170 | 0.0000000 | 0.5005280 | 0.9987349 | 0.4896312 |
| 19 | 14.70000 | 1 | 0.0000010 | 0.0011420 | 0.9036932 | 0.0058312 | 0.0000000 | 0.0000088 | 0.0000000 | 0.2303493 | 0.9200172 | 0.4297304 |
| 20 | 14.86667 | 1 | 0.0000419 | 0.0023792 | 0.5691519 | 0.0023196 | 0.0000000 | 0.0000037 | 0.0000000 | 0.4297843 | 0.9174694 | 0.7126943 |
| 21 | 15.03333 | 1 | 0.0000250 | 0.0001165 | 0.2886747 | 0.0001032 | 0.0000000 | 0.0000007 | 0.0000000 | 0.4873025 | 0.9990995 | 0.5318826 |
| 22 | 15.20000 | 1 | 0.0057992 | 0.0006994 | 0.2050323 | 0.0000828 | 0.0000000 | 0.0000028 | 0.0000000 | 0.1092339 | 0.9171514 | 0.2473205 |
| 23 | 15.36667 | 1 | 0.0229502 | 0.0001024 | 0.0197737 | 0.0000022 | 0.0000000 | 0.0000004 | 0.0000000 | 0.0718352 | 0.7167874 | 0.3487291 |
| 24 | 15.53333 | 1 | 0.0029877 | 0.0000276 | 0.0057279 | 0.0001302 | 0.0000000 | 0.0000000 | 0.0000000 | 0.1387842 | 0.5028595 | 0.8528950 |
| 25 | 15.70000 | 1 | 0.0008675 | 0.0000081 | 0.0078057 | 0.0194118 | 0.0000000 | 0.0000000 | 0.0000007 | 0.7519569 | 0.8670511 | 0.9996878 |
| 26 | 15.86667 | 1 | 0.0034852 | 0.0004145 | 0.0009953 | 0.0681158 | 0.0000000 | 0.0000000 | 0.0000079 | 0.8047497 | 0.8521174 | 0.9989461 |
| 27 | 16.03333 | 1 | 0.0509469 | 0.0003057 | 0.0022982 | 0.0088702 | 0.0000005 | 0.0000003 | 0.0000017 | 0.7133200 | 0.9929038 | 0.9442235 |
| 28 | 16.20000 | 1 | 0.3589016 | 0.0003443 | 0.0046306 | 0.0042225 | 0.0000003 | 0.0000067 | 0.0000020 | 0.8504900 | 0.9999496 | 0.8505011 |
| 29 | 16.36667 | 1 | 0.2587517 | 0.0049448 | 0.0961591 | 0.0141480 | 0.0000012 | 0.0000389 | 0.0000001 | 0.7238101 | 0.9999969 | 0.7633184 |
| 30 | 16.53333 | 1 | 0.3548807 | 0.0116868 | 0.2447460 | 0.0016592 | 0.0000207 | 0.0003388 | 0.0000000 | 0.6736767 | 0.9923610 | 0.3505010 |
| 31 | 16.70000 | 1 | 0.1083447 | 0.1324839 | 0.6783166 | 0.0023216 | 0.0000039 | 0.0022762 | 0.0000000 | 0.6680310 | 0.7424034 | 0.0788315 |
| 32 | 16.86667 | 1 | 0.0999484 | 0.1823212 | 0.6064125 | 0.0054487 | 0.0000018 | 0.0014786 | 0.0000000 | 0.8713117 | 0.6195538 | 0.1039514 |
| 33 | 17.03333 | 1 | 0.0808568 | 0.2130640 | 0.6417722 | 0.0078742 | 0.0000192 | 0.0007565 | 0.0000000 | 0.8642118 | 0.4555103 | 0.1110082 |
| 34 | 17.20000 | 1 | 0.0914954 | 0.1598057 | 0.5799611 | 0.0193220 | 0.0000398 | 0.0010720 | 0.0000025 | 0.7339044 | 0.8949565 | 0.2220553 |
| 35 | 22.70000 | 1 | 0.0186050 | 0.3035765 | 0.0380242 | 0.9873474 | 0.0013926 | 0.0000073 | 0.0678795 | 0.9047094 | 0.4425872 | 0.1359505 |
| 36 | 22.86667 | 1 | 0.0164831 | 0.5300683 | 0.0434241 | 0.9297754 | 0.0033094 | 0.0000016 | 0.0138665 | 0.8596569 | 0.9396118 | 0.3369498 |
| 37 | 23.03333 | 1 | 0.3152863 | 0.3939890 | 0.0255029 | 0.7087083 | 0.0199553 | 0.0000541 | 0.0469429 | 0.8405979 | 0.9991243 | 0.7305938 |
| 38 | 23.20000 | 1 | 0.9988534 | 0.0818987 | 0.0083554 | 0.0281777 | 0.1434877 | 0.0048838 | 0.0130622 | 0.7246882 | 0.9772743 | 0.9945221 |
| 39 | 23.70000 | 1 | 0.9669686 | 0.0020059 | 0.0105417 | 0.0003243 | 0.0502997 | 0.0310478 | 0.0074042 | 0.9985915 | 0.9134843 | 0.8416945 |

##### 2.2.4 Dist from Center

###### 2.2.4.1 Static

remaking specific dataframe

```
onlystatic <- subset(full_data, full_data$moving == 0)

data <- onlystatic
```

reloading of the data and ps

```
static <- read.csv(paste0(tmp_path, 'distFromCenter_staticResults.csv'))

static$adjustedp <- static$p*sum(static$stimtype=='bio') # create bonferroni corrected p
static$adjustedp[static$adjustedp>1] <- 1
significant_static <- subset(static, static$stimtype=='bio') #subset to have only 1 row per model

significant_static <- subset(significant_static, significant_static$adjustedp < alpha)

significant_static_wins <- significant_static$win_starts #identify the windows that are significant
```

###### 2.2.4.1.1 Detailed

now redo the models

```
static_results_posthoc <- data.frame(list('win_starts'=NaN,'win_span'=NaN,
                                          'bio/ellipse'=NaN, 'bio/none'=NaN,
                                          'bio/rand'=NaN, 'bio/scramb'=NaN,
                                          'bio/silhouette'=NaN, 'ellipse/none'=NaN,
                                          'ellipse/rand'=NaN, 'ellipse/scramb'=NaN,
                                          'ellipse/silhouette'=NaN, 'none/rand'=NaN,
                                          'none/scramb'=NaN, 'none/silhouette'=NaN,
                                          'rand/scramb'=NaN, 'rand/silhouette'=NaN,
                                          'scramb/silhouette'=NaN))

for (window_start in significant_static_wins){
  window_end <- window_start + window_size

  window_data <- subset(data, data$time_from_start>window_start)
  window_data <- subset(window_data, window_data$time_from_start<window_end)

  full_tmb <- glmmTMB(log(dist_from_center) ~ stimtype + (stimtype|subj), family=gaussian, data=window_data)
  Anova(full_tmb)
  e <- emmeans(full_tmb, ~stimtype, type='response')
  prs <- as.data.frame(pairs(e))
  newrow <- c(window_start, window_size, prs$p.value[1], prs$p.value[2],
              prs$p.value[3], prs$p.value[4], prs$p.value[5], prs$p.value[6],
              prs$p.value[7], prs$p.value[8], prs$p.value[9], prs$p.value[10],
              prs$p.value[11], prs$p.value[12], prs$p.value[13], prs$p.value[14],
              prs$p.value[15])
  static_results_posthoc <- rbind(static_results_posthoc, newrow)
  print(window_start)
  print(pairs(e))
}
```

###### 2.2.4.1.2 Summary

```
static_results_posthoc <- static_results_posthoc[-1,]
write.csv(static_results_posthoc, paste0(tmp_path, 'distFromCenter_staticResultsPostHoc.csv'))

kable(static_results_posthoc)
```

| win\_starts | win\_span | bio.ellipse | bio.none | bio.rand | bio.scramb | bio.silhouette | ellipse.none | ellipse.rand | ellipse.scramb | ellipse.silhouette | none.rand | none.scramb | none.silhouette | rand.scramb | rand.silhouette | scramb.silhouette |
| --- | --- | --- | --- | --- | --- | --- | --- | --- | --- | --- | --- | --- | --- | --- | --- | --- |

###### 2.2.4.2 Moving

remaking specific dataframe

```
onlymoving <- subset(full_data, full_data$stimtype != 'none')

data <- onlymoving
```

reloading of the data and ps

```
moving <- read.csv(paste0(tmp_path, 'distFromCenter_movingResults.csv'))

moving$adjustedp <- moving$p*sum(moving$stimtype=='bio') # create bonferroni corrected p
moving$adjustedp[moving$adjustedp>1] <- 1
significant_moving <- subset(moving, moving$stimtype=='bio') #subset to have only 1 row per model

significant_moving <- subset(significant_moving, significant_moving$adjustedp < alpha)

significant_moving_wins <- significant_moving$win_starts #identify the windows that are significant
```

###### 2.2.4.2.1 Detailed

now redo the models

```
moving_results_posthoc <- data.frame(list('win_starts'=NaN,'win_span'=NaN,
                                          'bio/ellipse'=NaN, 'bio/rand'=NaN,
                                          'bio/scramb'=NaN, 'bio/silhouette'=NaN,
                                          'ellipse/rand'=NaN, 'ellipse/scramb'=NaN,
                                          'ellipse/silhouette'=NaN, 'rand/scramb'=NaN,
                                          'rand/silhouette'=NaN, 'scramb/silhouette'=NaN))

for (window_start in significant_moving_wins){
  window_end <- window_start + window_size

  window_data <- subset(data, data$time_from_start>window_start)
  window_data <- subset(window_data, window_data$time_from_start<window_end)

  full_tmb <- glmmTMB(log(dist_from_center) ~ stimtype + (stimtype|subj), family=gaussian, data=window_data)
  Anova(full_tmb)
  e <- emmeans(full_tmb, ~stimtype, type='response')
  prs <- as.data.frame(pairs(e))
  newrow <- c(window_start, window_size, prs$p.value[1], prs$p.value[2],
              prs$p.value[3], prs$p.value[4], prs$p.value[5], prs$p.value[6],
              prs$p.value[7], prs$p.value[8], prs$p.value[9], prs$p.value[10])
  moving_results_posthoc <- rbind(moving_results_posthoc, newrow)

  print(window_start)
  print(pairs(e))
}
```

```
## [1] 11.86667
##  contrast             ratio     SE    df null t.ratio p.value
##  bio / ellipse        0.882 0.0559 11239    1  -1.976  0.2780
##  bio / rand           1.176 0.0531 11239    1   3.587  0.0031
##  bio / scramb         1.082 0.0326 11239    1   2.612  0.0683
##  bio / silhouette     1.191 0.0686 11239    1   3.037  0.0202
##  ellipse / rand       1.333 0.0834 11239    1   4.588  <.0001
##  ellipse / scramb     1.226 0.0730 11239    1   3.425  0.0056
##  ellipse / silhouette 1.350 0.0748 11239    1   5.415  <.0001
##  rand / scramb        0.920 0.0276 11239    1  -2.775  0.0439
##  rand / silhouette    1.013 0.0572 11239    1   0.228  0.9994
##  scramb / silhouette  1.101 0.0532 11239    1   1.989  0.2714
## 
## P value adjustment: tukey method for comparing a family of 5 estimates 
## Tests are performed on the log scale 
## [1] 12.03333
##  contrast             ratio     SE    df null t.ratio p.value
##  bio / ellipse        0.795 0.0592 10996    1  -3.084  0.0175
##  bio / rand           1.219 0.0629 10996    1   3.836  0.0012
##  bio / scramb         1.084 0.0394 10996    1   2.220  0.1722
##  bio / silhouette     1.250 0.0773 10996    1   3.617  0.0028
##  ellipse / rand       1.534 0.1130 10996    1   5.789  <.0001
##  ellipse / scramb     1.364 0.0933 10996    1   4.542  0.0001
##  ellipse / silhouette 1.574 0.1160 10996    1   6.151  <.0001
##  rand / scramb        0.889 0.0300 10996    1  -3.483  0.0045
##  rand / silhouette    1.026 0.0537 10996    1   0.487  0.9885
##  scramb / silhouette  1.154 0.0510 10996    1   3.231  0.0108
## 
## P value adjustment: tukey method for comparing a family of 5 estimates 
## Tests are performed on the log scale 
## [1] 12.2
##  contrast             ratio     SE    df null t.ratio p.value
##  bio / ellipse        0.709 0.0638 10058    1  -3.820  0.0013
##  bio / rand           1.306 0.0760 10058    1   4.588  <.0001
##  bio / scramb         1.089 0.0430 10058    1   2.164  0.1938
##  bio / silhouette     1.309 0.0876 10058    1   4.016  0.0006
##  ellipse / rand       1.842 0.1760 10058    1   6.389  <.0001
##  ellipse / scramb     1.536 0.1250 10058    1   5.294  <.0001
##  ellipse / silhouette 1.845 0.1620 10058    1   6.958  <.0001
##  rand / scramb        0.834 0.0386 10058    1  -3.925  0.0008
##  rand / silhouette    1.002 0.0596 10058    1   0.031  1.0000
##  scramb / silhouette  1.201 0.0616 10058    1   3.578  0.0032
## 
## P value adjustment: tukey method for comparing a family of 5 estimates 
## Tests are performed on the log scale 
## [1] 12.36667
##  contrast             ratio     SE    df null t.ratio p.value
##  bio / ellipse        0.650 0.0604 10095    1  -4.635  <.0001
##  bio / rand           1.339 0.0880 10095    1   4.445  0.0001
##  bio / scramb         1.096 0.0512 10095    1   1.955  0.2885
##  bio / silhouette     1.360 0.1080 10095    1   3.888  0.0010
##  ellipse / rand       2.059 0.2180 10095    1   6.823  <.0001
##  ellipse / scramb     1.684 0.1510 10095    1   5.836  <.0001
##  ellipse / silhouette 2.092 0.2010 10095    1   7.678  <.0001
##  rand / scramb        0.818 0.0465 10095    1  -3.528  0.0038
##  rand / silhouette    1.016 0.0744 10095    1   0.215  0.9995
##  scramb / silhouette  1.242 0.0924 10095    1   2.910  0.0298
## 
## P value adjustment: tukey method for comparing a family of 5 estimates 
## Tests are performed on the log scale 
## [1] 12.53333
##  contrast             ratio     SE    df null t.ratio p.value
##  bio / ellipse        0.650 0.0575 10113    1  -4.867  <.0001
##  bio / rand           1.356 0.1030 10113    1   3.999  0.0006
##  bio / scramb         1.107 0.0559 10113    1   2.002  0.2651
##  bio / silhouette     1.376 0.1110 10113    1   3.954  0.0007
##  ellipse / rand       2.086 0.2260 10113    1   6.790  <.0001
##  ellipse / scramb     1.702 0.1560 10113    1   5.818  <.0001
##  ellipse / silhouette 2.117 0.2070 10113    1   7.667  <.0001
##  rand / scramb        0.816 0.0547 10113    1  -3.032  0.0206
##  rand / silhouette    1.015 0.0729 10113    1   0.203  0.9996
##  scramb / silhouette  1.244 0.0996 10113    1   2.720  0.0511
## 
## P value adjustment: tukey method for comparing a family of 5 estimates 
## Tests are performed on the log scale 
## [1] 12.7
##  contrast             ratio     SE   df null t.ratio p.value
##  bio / ellipse        0.661 0.0572 9411    1  -4.778  <.0001
##  bio / rand           1.370 0.1310 9411    1   3.283  0.0091
##  bio / scramb         1.066 0.0664 9411    1   1.027  0.8429
##  bio / silhouette     1.341 0.1090 9411    1   3.607  0.0029
##  ellipse / rand       2.071 0.2460 9411    1   6.134  <.0001
##  ellipse / scramb     1.612 0.1450 9411    1   5.300  <.0001
##  ellipse / silhouette 2.028 0.2050 9411    1   6.991  <.0001
##  rand / scramb        0.778 0.0656 9411    1  -2.974  0.0246
##  rand / silhouette    0.979 0.0775 9411    1  -0.265  0.9989
##  scramb / silhouette  1.258 0.0981 9411    1   2.943  0.0270
## 
## P value adjustment: tukey method for comparing a family of 5 estimates 
## Tests are performed on the log scale 
## [1] 12.86667
##  contrast             ratio     SE   df null t.ratio p.value
##  bio / ellipse        0.738 0.0560 9263    1  -4.004  0.0006
##  bio / rand           1.446 0.1390 9263    1   3.844  0.0012
##  bio / scramb         1.099 0.0821 9263    1   1.267  0.7116
##  bio / silhouette     1.395 0.1090 9263    1   4.263  0.0002
##  ellipse / rand       1.960 0.2400 9263    1   5.489  <.0001
##  ellipse / scramb     1.490 0.1390 9263    1   4.275  0.0002
##  ellipse / silhouette 1.890 0.1830 9263    1   6.564  <.0001
##  rand / scramb        0.760 0.0697 9263    1  -2.992  0.0232
##  rand / silhouette    0.965 0.0765 9263    1  -0.454  0.9912
##  scramb / silhouette  1.269 0.1050 9263    1   2.880  0.0325
## 
## P value adjustment: tukey method for comparing a family of 5 estimates 
## Tests are performed on the log scale 
## [1] 13.03333
##  contrast             ratio     SE   df null t.ratio p.value
##  bio / ellipse        0.794 0.0641 8900    1  -2.857  0.0347
##  bio / rand           1.416 0.1300 8900    1   3.774  0.0015
##  bio / scramb         1.128 0.1010 8900    1   1.349  0.6604
##  bio / silhouette     1.384 0.1110 8900    1   4.032  0.0005
##  ellipse / rand       1.783 0.2170 8900    1   4.751  <.0001
##  ellipse / scramb     1.421 0.1490 8900    1   3.357  0.0071
##  ellipse / silhouette 1.743 0.1630 8900    1   5.930  <.0001
##  rand / scramb        0.797 0.0743 8900    1  -2.435  0.1063
##  rand / silhouette    0.977 0.0783 8900    1  -0.286  0.9985
##  scramb / silhouette  1.226 0.1110 8900    1   2.264  0.1568
## 
## P value adjustment: tukey method for comparing a family of 5 estimates 
## Tests are performed on the log scale 
## [1] 13.2
##  contrast             ratio     SE   df null t.ratio p.value
##  bio / ellipse        0.839 0.0728 8306    1  -2.026  0.2535
##  bio / rand           1.388 0.1440 8306    1   3.151  0.0141
##  bio / scramb         1.064 0.0893 8306    1   0.737  0.9478
##  bio / silhouette     1.394 0.1210 8306    1   3.834  0.0012
##  ellipse / rand       1.655 0.2240 8306    1   3.716  0.0019
##  ellipse / scramb     1.268 0.1320 8306    1   2.278  0.1520
##  ellipse / silhouette 1.661 0.1600 8306    1   5.287  <.0001
##  rand / scramb        0.766 0.0687 8306    1  -2.969  0.0250
##  rand / silhouette    1.004 0.0897 8306    1   0.046  1.0000
##  scramb / silhouette  1.310 0.1180 8306    1   2.998  0.0228
## 
## P value adjustment: tukey method for comparing a family of 5 estimates 
## Tests are performed on the log scale 
## [1] 13.36667
##  contrast             ratio     SE   df null t.ratio p.value
##  bio / ellipse        0.895 0.0817 8314    1  -1.215  0.7428
##  bio / rand           1.347 0.1560 8314    1   2.581  0.0740
##  bio / scramb         1.035 0.0887 8314    1   0.404  0.9944
##  bio / silhouette     1.440 0.1370 8314    1   3.825  0.0012
##  ellipse / rand       1.505 0.2140 8314    1   2.874  0.0331
##  ellipse / scramb     1.157 0.1180 8314    1   1.422  0.6136
##  ellipse / silhouette 1.608 0.1520 8314    1   5.021  <.0001
##  rand / scramb        0.768 0.0860 8314    1  -2.352  0.1288
##  rand / silhouette    1.069 0.1060 8314    1   0.672  0.9625
##  scramb / silhouette  1.391 0.1290 8314    1   3.550  0.0036
## 
## P value adjustment: tukey method for comparing a family of 5 estimates 
## Tests are performed on the log scale 
## [1] 13.53333
##  contrast             ratio     SE   df null t.ratio p.value
##  bio / ellipse        0.911 0.0857 8226    1  -0.992  0.8590
##  bio / rand           1.318 0.1660 8226    1   2.183  0.1860
##  bio / scramb         1.010 0.0860 8226    1   0.114  1.0000
##  bio / silhouette     1.488 0.1450 8226    1   4.079  0.0004
##  ellipse / rand       1.447 0.2130 8226    1   2.512  0.0880
##  ellipse / scramb     1.109 0.1080 8226    1   1.059  0.8273
##  ellipse / silhouette 1.633 0.1510 8226    1   5.321  <.0001
##  rand / scramb        0.766 0.0969 8226    1  -2.106  0.2177
##  rand / silhouette    1.129 0.1260 8226    1   1.093  0.8104
##  scramb / silhouette  1.473 0.1350 8226    1   4.221  0.0002
## 
## P value adjustment: tukey method for comparing a family of 5 estimates 
## Tests are performed on the log scale 
## [1] 13.7
##  contrast             ratio     SE   df null t.ratio p.value
##  bio / ellipse        0.948 0.0883 7929    1  -0.576  0.9786
##  bio / rand           1.338 0.1630 7929    1   2.385  0.1195
##  bio / scramb         1.003 0.0910 7929    1   0.033  1.0000
##  bio / silhouette     1.570 0.1600 7929    1   4.434  0.0001
##  ellipse / rand       1.412 0.2100 7929    1   2.313  0.1406
##  ellipse / scramb     1.058 0.1020 7929    1   0.591  0.9765
##  ellipse / silhouette 1.656 0.1500 7929    1   5.569  <.0001
##  rand / scramb        0.750 0.1070 7929    1  -2.021  0.2561
##  rand / silhouette    1.173 0.1350 7929    1   1.385  0.6375
##  scramb / silhouette  1.565 0.1560 7929    1   4.483  0.0001
## 
## P value adjustment: tukey method for comparing a family of 5 estimates 
## Tests are performed on the log scale 
## [1] 13.86667
##  contrast             ratio     SE   df null t.ratio p.value
##  bio / ellipse        0.958 0.0899 8218    1  -0.461  0.9907
##  bio / rand           1.259 0.1430 8218    1   2.030  0.2517
##  bio / scramb         0.964 0.0852 8218    1  -0.419  0.9936
##  bio / silhouette     1.614 0.1650 8218    1   4.685  <.0001
##  ellipse / rand       1.315 0.2020 8218    1   1.779  0.3861
##  ellipse / scramb     1.006 0.0878 8218    1   0.071  1.0000
##  ellipse / silhouette 1.686 0.1580 8218    1   5.572  <.0001
##  rand / scramb        0.765 0.1150 8218    1  -1.773  0.3898
##  rand / silhouette    1.282 0.1410 8218    1   2.255  0.1597
##  scramb / silhouette  1.675 0.1740 8218    1   4.963  <.0001
## 
## P value adjustment: tukey method for comparing a family of 5 estimates 
## Tests are performed on the log scale 
## [1] 14.03333
##  contrast             ratio     SE   df null t.ratio p.value
##  bio / ellipse        0.959 0.0896 8253    1  -0.450  0.9915
##  bio / rand           1.248 0.1250 8253    1   2.201  0.1792
##  bio / scramb         0.903 0.0766 8253    1  -1.199  0.7522
##  bio / silhouette     1.581 0.1570 8253    1   4.620  <.0001
##  ellipse / rand       1.301 0.1930 8253    1   1.773  0.3893
##  ellipse / scramb     0.942 0.0775 8253    1  -0.724  0.9509
##  ellipse / silhouette 1.649 0.1560 8253    1   5.288  <.0001
##  rand / scramb        0.724 0.0990 8253    1  -2.362  0.1260
##  rand / silhouette    1.268 0.1410 8253    1   2.130  0.2073
##  scramb / silhouette  1.751 0.1810 8253    1   5.413  <.0001
## 
## P value adjustment: tukey method for comparing a family of 5 estimates 
## Tests are performed on the log scale 
## [1] 14.2
##  contrast             ratio     SE   df null t.ratio p.value
##  bio / ellipse        0.992 0.0861 8012    1  -0.093  1.0000
##  bio / rand           1.210 0.0880 8012    1   2.620  0.0668
##  bio / scramb         0.937 0.0694 8012    1  -0.877  0.9053
##  bio / silhouette     1.599 0.1480 8012    1   5.070  <.0001
##  ellipse / rand       1.220 0.1450 8012    1   1.669  0.4534
##  ellipse / scramb     0.945 0.0777 8012    1  -0.692  0.9583
##  ellipse / silhouette 1.612 0.1520 8012    1   5.074  <.0001
##  rand / scramb        0.774 0.0873 8012    1  -2.266  0.1560
##  rand / silhouette    1.322 0.1370 8012    1   2.686  0.0562
##  scramb / silhouette  1.706 0.1780 8012    1   5.131  <.0001
## 
## P value adjustment: tukey method for comparing a family of 5 estimates 
## Tests are performed on the log scale 
## [1] 14.36667
##  contrast             ratio     SE   df null t.ratio p.value
##  bio / ellipse        0.965 0.0862 8361    1  -0.402  0.9945
##  bio / rand           1.198 0.0831 8361    1   2.606  0.0693
##  bio / scramb         0.911 0.0658 8361    1  -1.294  0.6949
##  bio / silhouette     1.515 0.1440 8361    1   4.375  0.0001
##  ellipse / rand       1.242 0.1440 8361    1   1.870  0.3337
##  ellipse / scramb     0.944 0.0751 8361    1  -0.723  0.9512
##  ellipse / silhouette 1.571 0.1570 8361    1   4.525  0.0001
##  rand / scramb        0.760 0.0742 8361    1  -2.808  0.0401
##  rand / silhouette    1.265 0.1300 8361    1   2.290  0.1480
##  scramb / silhouette  1.664 0.1790 8361    1   4.738  <.0001
## 
## P value adjustment: tukey method for comparing a family of 5 estimates 
## Tests are performed on the log scale
```

###### 2.2.4.2.2 Summary

```
moving_results_posthoc <- moving_results_posthoc[-1,]
write.csv(moving_results_posthoc, paste0(tmp_path, 'distFromCenter_movingResultsPostHoc.csv'))
kable(moving_results_posthoc)
```

|  | win\_starts | win\_span | bio.ellipse | bio.rand | bio.scramb | bio.silhouette | ellipse.rand | ellipse.scramb | ellipse.silhouette | rand.scramb | rand.silhouette | scramb.silhouette |
| --- | --- | --- | --- | --- | --- | --- | --- | --- | --- | --- | --- | --- |
| 2 | 11.86667 | 1 | 6.25e-05 | 0.1534138 | 0.6719668 | 0.6834571 | 0.0e+00 | 0.0000000 | 1e-07 | 0.7113383 | 0.9710770 | 0.9982067 |
| 3 | 12.03333 | 1 | 0.00e+00 | 0.1209482 | 0.5961292 | 0.4438747 | 0.0e+00 | 0.0000000 | 0e+00 | 0.7959257 | 0.9960933 | 0.9926391 |
| 4 | 12.20000 | 1 | 0.00e+00 | 0.1499212 | 0.8740168 | 0.0875872 | 0.0e+00 | 0.0000000 | 0e+00 | 0.5685163 | 0.9963386 | 0.5322285 |
| 5 | 12.36667 | 1 | 0.00e+00 | 0.8265526 | 0.9210221 | 0.4775375 | 0.0e+00 | 0.0000000 | 0e+00 | 0.9919608 | 0.9951907 | 0.9579731 |
| 6 | 12.53333 | 1 | 0.00e+00 | 0.9631019 | 0.9689530 | 0.5405221 | 0.0e+00 | 0.0000000 | 0e+00 | 0.9998751 | 0.9848708 | 0.9663156 |
| 7 | 12.70000 | 1 | 0.00e+00 | 0.9999961 | 0.9973188 | 0.8162556 | 0.0e+00 | 0.0000000 | 0e+00 | 0.9925266 | 0.9400816 | 0.9823843 |
| 8 | 12.86667 | 1 | 0.00e+00 | 0.8593978 | 0.9748175 | 0.9995108 | 0.0e+00 | 0.0000000 | 0e+00 | 0.9869750 | 0.8972136 | 0.9724883 |
| 9 | 13.03333 | 1 | 0.00e+00 | 0.9959206 | 0.9999976 | 0.9999276 | 0.0e+00 | 0.0000000 | 0e+00 | 0.9937244 | 0.9994548 | 0.9999974 |
| 10 | 13.20000 | 1 | 0.00e+00 | 0.9813893 | 0.9426973 | 0.9997918 | 0.0e+00 | 0.0000000 | 0e+00 | 0.9999736 | 0.9950376 | 0.9831656 |
| 11 | 13.36667 | 1 | 0.00e+00 | 0.8361167 | 0.9330236 | 0.9856064 | 0.0e+00 | 0.0000000 | 0e+00 | 0.9943753 | 0.9918529 | 0.9995893 |
| 12 | 13.53333 | 1 | 0.00e+00 | 0.8504838 | 0.9377945 | 0.8775515 | 0.0e+00 | 0.0000000 | 0e+00 | 0.9969380 | 0.9999990 | 0.9949025 |
| 13 | 13.70000 | 1 | 0.00e+00 | 0.8651732 | 0.8070965 | 0.6569802 | 2.0e-07 | 0.0000000 | 0e+00 | 1.0000000 | 0.9950817 | 0.9781998 |
| 14 | 13.86667 | 1 | 0.00e+00 | 0.6702740 | 0.9894221 | 0.5977177 | 8.0e-07 | 0.0000019 | 0e+00 | 0.9037807 | 0.9997134 | 0.7474114 |
| 15 | 14.03333 | 1 | 1.34e-05 | 0.5241035 | 0.9978577 | 0.3801339 | 7.7e-06 | 0.0001328 | 0e+00 | 0.7366660 | 0.9999214 | 0.4405565 |
| 16 | 14.20000 | 1 | 6.00e-07 | 0.5137584 | 0.9937282 | 0.3693170 | 4.5e-06 | 0.0000455 | 1e-07 | 0.7318152 | 0.9999627 | 0.7173067 |
| 17 | 14.36667 | 1 | 1.50e-06 | 0.3615763 | 0.9751538 | 0.2646673 | 1.0e-07 | 0.0000181 | 1e-07 | 0.7327903 | 0.9999984 | 0.6106966 |

#### 2.3 Plots

##### 2.3.1 Eye displacement

```
static <- read.csv(paste0(tmp_path, 'displacement_staticResults.csv'))


ggplot(static, aes(x=win_starts, y=emmean, color=stimtype))+
  geom_ribbon(aes(x=win_starts, ymin=emmean-SE, ymax = emmean+SE, group=stimtype, fill = stimtype), alpha=0.1, colour=NA)+
    geom_line()
```

```
moving <- read.csv(paste0(tmp_path, 'displacement_movingResults.csv'))


ggplot(moving, aes(x=win_starts, y=emmean, color=stimtype))+
  geom_ribbon(aes(x=win_starts, ymin=emmean-SE, ymax = emmean+SE, group=stimtype, fill = stimtype), alpha=0.1, colour=NA)+
    geom_line()
```

##### 2.3.2 Inter-eye distance

```
static <- read.csv(paste0(tmp_path, 'interEyeDist_staticResults.csv'))


ggplot(static, aes(x=win_starts, y=emmean, color=stimtype))+
  geom_ribbon(aes(x=win_starts, ymin=emmean-SE, ymax = emmean+SE, group=stimtype, fill = stimtype), alpha=0.1, colour=NA)+
    geom_line()
```

```
moving <- read.csv(paste0(tmp_path, 'interEyeDist_movingResults.csv'))


ggplot(moving, aes(x=win_starts, y=emmean, color=stimtype))+
  geom_ribbon(aes(x=win_starts, ymin=emmean-SE, ymax = emmean+SE, group=stimtype, fill = stimtype), alpha=0.1, colour=NA)+
    geom_line()
```

##### 2.3.3 Inter-eye distance change

```
static <- read.csv(paste0(tmp_path, 'interEyeDistChange_staticResults.csv'))


ggplot(static, aes(x=win_starts, y=emmean, color=stimtype))+
  geom_ribbon(aes(x=win_starts, ymin=emmean-SE, ymax = emmean+SE, group=stimtype, fill = stimtype), alpha=0.1, colour=NA)+
    geom_line()
```

```
moving <- read.csv(paste0(tmp_path, 'interEyeDistChange_movingResults.csv'))


ggplot(moving, aes(x=win_starts, y=emmean, color=stimtype))+
  geom_ribbon(aes(x=win_starts, ymin=emmean-SE, ymax = emmean+SE, group=stimtype, fill = stimtype), alpha=0.1, colour=NA)+
    geom_line()
```

##### 2.3.4 Dist from Center

```
static <- read.csv(paste0(tmp_path, 'distFromCenter_staticResults.csv'))


ggplot(static, aes(x=win_starts, y=emmean, color=stimtype))+
  geom_ribbon(aes(x=win_starts, ymin=emmean-SE, ymax = emmean+SE, group=stimtype, fill = stimtype), alpha=0.1, colour=NA)+
    geom_line()
```

```
moving <- read.csv(paste0(tmp_path, 'distFromCenter_movingResults.csv'))


ggplot(moving, aes(x=win_starts, y=emmean, color=stimtype))+
  geom_ribbon(aes(x=win_starts, ymin=emmean-SE, ymax = emmean+SE, group=stimtype, fill = stimtype), alpha=0.1, colour=NA)+
    geom_line()
```
