## Supplement for "Biological point-light displays scanning by the principal eyes of a jumping spider": fullPvaluesTable.pdf

### Retinal speed - Sliding window GLMM for Static Section

| win_starts | win_span | p |
|---|---|---|
| 0.03333 | 1 | 0.00041 |
| 0.2 | 1 | <0.0001 |
| 0.36667 | 1 | <0.0001 |
| 0.53333 | 1 | <0.0001 |
| 0.7 | 1 | <0.0001 |
| 0.86667 | 1 | 0.00018 |
| 1.03333 | 1 | 0.00042 |
| 1.2 | 1 | <0.0001 |
| 1.36667 | 1 | <0.0001 |
| 1.53333 | 1 | <0.0001 |
| 1.7 | 1 | <0.0001 |
| 1.86667 | 1 | <0.0001 |
| 2.03333 | 1 | <0.0001 |
| 2.2 | 1 | <0.0001 |
| 2.36667 | 1 | 0.00023 |
| 2.53333 | 1 | 0.00342 |
| 2.7 | 1 | 0.00324 |
| 2.86667 | 1 | 0.00346 |
| 3.03333 | 1 | 0.06761 |
| 3.2 | 1 | 0.01472 |

| win_starts | win_span | p |
|---|---|---|
| 3.36667 | 1 | 0.03341 |
| 3.53333 | 1 | 0.20273 |
| 3.7 | 1 | 0.12835 |
| 3.86667 | 1 | 0.13698 |
| 4.03333 | 1 | 0.07253 |
| 4.2 | 1 | 0.04762 |
| 4.36667 | 1 | 0.01164 |
| 4.53333 | 1 | 0.01308 |
| 4.7 | 1 | 0.03387 |
| 4.86667 | 1 | 0.06129 |
| 5.03333 | 1 | 0.02313 |
| 5.2 | 1 | 0.01902 |
| 5.36667 | 1 | 0.09753 |
| 5.53333 | 1 | 0.12673 |
| 5.7 | 1 | 0.49437 |
| 5.86667 | 1 | 0.12778 |
| 6.03333 | 1 | 0.06102 |
| 6.2 | 1 | 0.00585 |
| 6.36667 | 1 | 0.01288 |
| 6.53333 | 1 | 0.00789 |
| 6.7 | 1 | 0.07627 |
| 6.86667 | 1 | 1 |

| win_starts | win_span | p |
|---|---|---|
| 7.03333 | 1 | 1 |
| 7.2 | 1 | 1 |
| 7.36667 | 1 | 1 |
| 7.53333 | 1 | 1 |
| 7.7 | 1 | 1 |
| 7.86667 | 1 | 0.7453 |
| 8.03333 | 1 | 0.08733 |
| 8.2 | 1 | 0.08715 |
| 8.36667 | 1 | 0.03422 |
| 8.53333 | 1 | 0.29549 |
| 8.7 | 1 | 1 |
| 8.86667 | 1 | 1 |
| 9.03333 | 1 | 0.55287 |
| 9.2 | 1 | 1 |
| 9.36667 | 1 | 1 |
| 9.53333 | 1 | 1 |
| 9.7 | 1 | 1 |
| 9.86667 | 1 | 1 |
| 10.03333 | 1 | 1 |
| 10.2 | 1 | 1 |
| 10.36667 | 1 | 1 |
| 10.53333 | 1 | 1 |

| win_starts | win_span | p |
|---|---|---|
| 10.7 | 1 | 1 |
| 10.86667 | 1 | 1 |

### Retinal speed - Post-Hoc Comparisons for Static Section

| win_starts | win_span | bio.ellipse | bio.none | bio.rand | bio.scramb | bio.silhouette | ellipse.none | none.rand | none.scramb | none.silhouette |
|---|---|---|---|---|---|---|---|---|---|---|
| 0.0333 | 1 | 1 | 0.0022 | 0.9997 | 1 | 0.859 | 0.0003 | 0.0196 | 0.0216 | 0.0002 |
| 0.2 | 1 | 0.997 | <0.0001 | 0.9997 | 1 | 0.6175 | <0.0001 | 0.0017 | 0.0057 | <0.0001 |
| 0.3667 | 1 | 1 | 0.0002 | 1 | 1 | 0.9696 | 0.0002 | 0.0019 | 0.0052 | <0.0001 |
| 0.5333 | 1 | 1 | 0.0008 | 0.9996 | 0.9845 | 0.9518 | 0.0015 | 0.0162 | 0.0576 | <0.0001 |
| 0.7 | 1 | 0.9999 | 0.003 | 1 | 0.9998 | 0.903 | 0.0046 | 0.0261 | 0.0536 | <0.0001 |
| 0.8667 | 1 | 1 | 0.0168 | 1 | 0.9992 | 0.9296 | 0.002 | 0.0748 | 0.1614 | <0.0001 |
| 1.0333 | 1 | 1 | 0.0196 | 1 | 0.9999 | 0.9455 | 0.0186 | 0.0808 | 0.0958 | <0.0001 |
| 1.2 | 1 | 0.989 | 0.0735 | 1 | 0.9998 | 0.5331 | 0.0016 | 0.2164 | 0.37 | <0.0001 |
| 1.3667 | 1 | 0.5998 | 0.2253 | 0.9999 | 1 | 0.2923 | <0.0001 | 0.3636 | 0.5564 | <0.0001 |
| 1.5333 | 1 | 0.6825 | 0.2175 | 1 | 0.9995 | 0.3353 | <0.0001 | 0.246 | 0.5204 | <0.0001 |
| 1.7 | 1 | 0.6154 | 0.2538 | 1 | 0.9982 | 0.3534 | <0.0001 | 0.2794 | 0.5651 | <0.0001 |
| 1.8667 | 1 | 0.6261 | 0.1922 | 0.9867 | 0.9741 | 0.4741 | 0.0003 | 0.5806 | 0.7753 | <0.0001 |
| 2.0333 | 1 | 0.6651 | 0.3076 | 0.9499 | 0.9905 | 0.6203 | <0.0001 | 0.7911 | 0.5377 | <0.0001 |
| 2.2 | 1 | 0.3791 | 0.4722 | 0.9965 | 0.9979 | 0.6797 | <0.0001 | 0.6312 | 0.6625 | 0.0002 |
| 2.3667 | 1 | 0.3786 | 0.6401 | 0.999 | 1 | 0.6184 | <0.0001 | 0.7165 | 0.3686 | 0.0009 |
| 2.5333 | 1 | 0.3011 | 0.8791 | 1 | 0.9999 | 0.4146 | <0.0001 | 0.6226 | 0.1512 | 0.0021 |
| 2.7 | 1 | 0.5134 | 0.8126 | 0.9999 | 0.9999 | 0.5112 | 0.0001 | 0.826 | 0.0564 | 0.0067 |
| 2.8667 | 1 | 0.2369 | 0.921 | 0.9995 | 0.9799 | 0.331 | <0.0001 | 0.7586 | 0.042 | 0.0243 |
| 6.2 | 1 | 0.98 | 0.9839 | 1 | 0.5814 | 0.8842 | 0.0688 | 0.9903 | 0.0015 | 0.1357 |
| 6.5333 | 1 | 0.9712 | 0.9904 | 0.9991 | 0.3992 | 0.8598 | 0.1438 | 0.8831 | 0.0016 | 0.2132 |

### Retinal speed - Sliding window GLMM for all stimulus duration

| win_starts | win_span | p |
|---|---|---|
| 0.03333 | 1 | 1 |
| 0.2 | 1 | 1 |
| 0.36667 | 1 | 1 |
| 0.53333 | 1 | 1 |
| 0.7 | 1 | 1 |
| 0.86667 | 1 | 1 |
| 1.03333 | 1 | 1 |
| 1.2 | 1 | 1 |
| 1.36667 | 1 | 1 |
| 1.53333 | 1 | 1 |
| 1.7 | 1 | 1 |
| 1.86667 | 1 | 0.58732 |
| 2.03333 | 1 | 0.04694 |
| 2.2 | 1 | 0.05709 |
| 2.36667 | 1 | 0.12855 |
| 2.53333 | 1 | 0.88945 |
| 2.7 | 1 | 1 |
| 2.86667 | 1 | 1 |
| 3.03333 | 1 | 1 |
| 3.2 | 1 | 1 |

| win_starts | win_span | p |
|---|---|---|
| 3.36667 | 1 | 1 |
| 3.53333 | 1 | 1 |
| 3.7 | 1 | 1 |
| 3.86667 | 1 | 1 |
| 4.03333 | 1 | 1 |
| 4.2 | 1 | 1 |
| 4.36667 | 1 | 0.50451 |
| 4.53333 | 1 | 0.21275 |
| 4.7 | 1 | 0.30076 |
| 4.86667 | 1 | 0.96721 |
| 5.03333 | 1 | 0.27651 |
| 5.2 | 1 | 0.23869 |
| 5.36667 | 1 | 0.71166 |
| 5.53333 | 1 | 0.65015 |
| 5.7 | 1 | 1 |
| 5.86667 | 1 | 1 |
| 6.03333 | 1 | 1 |
| 6.2 | 1 | 1 |
| 6.36667 | 1 | 1 |
| 6.53333 | 1 | 1 |
| 6.7 | 1 | 1 |
| 6.86667 | 1 | 1 |

| win_starts | win_span | p |
|---|---|---|
| 7.03333 | 1 | 1 |
| 7.2 | 1 | 1 |
| 7.36667 | 1 | 1 |
| 7.53333 | 1 | 1 |
| 7.7 | 1 | 1 |
| 7.86667 | 1 | 1 |
| 8.03333 | 1 | 1 |
| 8.2 | 1 | 1 |
| 8.36667 | 1 | 1 |
| 8.53333 | 1 | 1 |
| 8.7 | 1 | 1 |
| 8.86667 | 1 | 1 |
| 9.03333 | 1 | 1 |
| 9.2 | 1 | 1 |
| 9.36667 | 1 | 1 |
| 9.53333 | 1 | 1 |
| 9.7 | 1 | 1 |
| 9.86667 | 1 | 1 |
| 10.03333 | 1 | 1 |
| 10.2 | 1 | 1 |
| 10.36667 | 1 | 1 |
| 10.53333 | 1 | 1 |

| win_starts | win_span | p |
|---|---|---|
| 10.7 | 1 | 1 |
| 10.86667 | 1 | 1 |
| 11.03333 | 1 | 1 |
| 11.2 | 1 | 1 |
| 11.36667 | 1 | 1 |
| 11.53333 | 1 | 1 |
| 11.7 | 1 | 0.58048 |
| 11.86667 | 1 | 0.00017 |
| 12.03333 | 1 | <0.0001 |
| 12.2 | 1 | <0.0001 |
| 12.36667 | 1 | <0.0001 |
| 12.53333 | 1 | <0.0001 |
| 12.7 | 1 | <0.0001 |
| 12.86667 | 1 | <0.0001 |
| 13.03333 | 1 | <0.0001 |
| 13.2 | 1 | <0.0001 |
| 13.36667 | 1 | <0.0001 |
| 13.53333 | 1 | <0.0001 |
| 13.7 | 1 | <0.0001 |
| 13.86667 | 1 | <0.0001 |
| 14.03333 | 1 | <0.0001 |
| 14.2 | 1 | <0.0001 |

| win_starts | win_span | p |
|---|---|---|
| 14.36667 | 1 | <0.0001 |
| 14.53333 | 1 | <0.0001 |
| 14.7 | 1 | <0.0001 |
| 14.86667 | 1 | <0.0001 |
| 15.03333 | 1 | <0.0001 |
| 15.2 | 1 | <0.0001 |
| 15.36667 | 1 | <0.0001 |
| 15.53333 | 1 | <0.0001 |
| 15.7 | 1 | <0.0001 |
| 15.86667 | 1 | <0.0001 |
| 16.03333 | 1 | <0.0001 |
| 16.2 | 1 | <0.0001 |
| 16.36667 | 1 | <0.0001 |
| 16.53333 | 1 | <0.0001 |
| 16.7 | 1 | <0.0001 |
| 16.86667 | 1 | <0.0001 |
| 17.03333 | 1 | <0.0001 |
| 17.2 | 1 | <0.0001 |
| 17.36667 | 1 | 0.00014 |
| 17.53333 | 1 | 0.00162 |
| 17.7 | 1 | 0.00849 |
| 17.86667 | 1 | 0.02862 |

| win_starts | win_span | p |
|---|---|---|
| 18.03333 | 1 | 0.04433 |
| 18.2 | 1 | 0.00756 |
| 18.36667 | 1 | 0.00212 |
| 18.53333 | 1 | 0.01148 |
| 18.7 | 1 | 0.00808 |
| 18.86667 | 1 | 0.01546 |
| 19.03333 | 1 | 0.99083 |
| 19.2 | 1 | 1 |
| 19.36667 | 1 | 1 |
| 19.53333 | 1 | 0.69353 |
| 19.7 | 1 | 0.30804 |
| 19.86667 | 1 | 0.31786 |
| 20.03333 | 1 | 0.00955 |
| 20.2 | 1 | 0.00017 |
| 20.36667 | 1 | <0.0001 |
| 20.53333 | 1 | 0.0007 |
| 20.7 | 1 | 0.00012 |
| 20.86667 | 1 | <0.0001 |
| 21.03333 | 1 | 0.07725 |
| 21.2 | 1 | 0.00367 |
| 21.36667 | 1 | <0.0001 |
| 21.53333 | 1 | 0.00039 |

| win_starts | win_span | p |
|---|---|---|
| 21.7 | 1 | 0.00021 |
| 21.86667 | 1 | <0.0001 |
| 22.03333 | 1 | 0.00011 |
| 22.2 | 1 | 0.00039 |
| 22.36667 | 1 | 0.00097 |
| 22.53333 | 1 | 0.01889 |
| 22.7 | 1 | 0.00477 |
| 22.86667 | 1 | 0.00921 |
| 23.03333 | 1 | 0.00498 |
| 23.2 | 1 | <0.0001 |
| 23.36667 | 1 | <0.0001 |
| 23.53333 | 1 | <0.0001 |
| 23.7 | 1 | <0.0001 |
| 23.86667 | 1 | <0.0001 |
| 24.03333 | 1 | <0.0001 |
| 24.2 | 1 | <0.0001 |
| 24.36667 | 1 | 0.00652 |
| 24.53333 | 1 | 0.00333 |
| 24.7 | 1 | 0.15164 |
| 24.86667 | 1 | 0.00582 |
| 25.03333 | 1 | 0.07234 |
| 25.2 | 1 | 0.36363 |

| win_starts | win_span | p |
|---|---|---|
| 25.36667 | 1 | 1 |
| 25.53333 | 1 | 0.33202 |
| 25.7 | 1 | 0.0739 |
| 25.86667 | 1 | 0.05766 |
| 26.03333 | 1 | 0.91448 |
| 26.2 | 1 | 1 |
| 26.36667 | 1 | 1 |
| 26.53333 | 1 | 1 |
| 26.7 | 1 | 1 |
| 26.86667 | 1 | 0.06927 |
| 27.03333 | 1 | 1 |
| 27.2 | 1 | 1 |
| 27.36667 | 1 | 1 |
| 27.53333 | 1 | 1 |
| 27.7 | 1 | 1 |
| 27.86667 | 1 | 1 |
| 28.03333 | 1 | 1 |
| 28.2 | 1 | 1 |
| 28.36667 | 1 | 1 |
| 28.53333 | 1 | 1 |
| 28.7 | 1 | 1 |
| 28.86667 | 1 | 1 |

| win_starts | win_span | p |
|---|---|---|
| 29.03333 | 1 | 1 |
| 29.2 | 1 | 1 |
| 29.36667 | 1 | 1 |
| 29.53333 | 1 | 1 |
| 29.7 | 1 | 1 |
| 29.86667 | 1 | 1 |
| 30.03333 | 1 | 1 |
| 30.2 | 1 | 1 |
| 30.36667 | 1 | 1 |
| 30.53333 | 1 | 1 |
| 30.7 | 1 | 1 |
| 30.86667 | 1 | 1 |
| 31.03333 | 1 | 1 |
| 31.2 | 1 | 0.16674 |
| 31.36667 | 1 | 0.48019 |
| 31.53333 | 1 | 0.23049 |
| 31.7 | 1 | 1 |
| 31.86667 | 1 | 1 |
| 32.03333 | 1 | 1 |
| 32.2 | 1 | 1 |
| 32.36667 | 1 | 1 |
| 32.53333 | 1 | 1 |

| win_starts | win_span | p |
|---|---|---|
| 32.7 | 1 | 1 |
| 32.86667 | 1 | 1 |
| 33.03333 | 1 | 1 |
| 33.2 | 1 | 1 |
| 33.36667 | 1 | 0.09775 |
| 33.53333 | 1 | 0.38094 |
| 33.7 | 1 | 0.67654 |
| 33.86667 | 1 | 1 |
| 34.03333 | 1 | 0.86719 |
| 34.2 | 1 | 1 |
| 34.36667 | 1 | 1 |
| 34.53333 | 1 | 1 |
| 34.7 | 1 | 0.51388 |
| 34.86667 | 1 | 1 |
| 35.03333 | 1 | 1 |
| 35.2 | 1 | 1 |

### Retinal speed - Post-Hoc Comparisons for all stimulus duration

| win_starts | win_span | bio.ellipse | bio.rand | bio.scramb | bio.silhouette | ellipse.rand | ellipse.scramb | ellipse.silhouette |
|---|---|---|---|---|---|---|---|---|
| 11.8667 | 1 | 0.0005 | 0.1603 | 0.9972 | 0.3233 | <0.0001 | 0.0009 | <0.0001 |
| 12.0333 | 1 | <0.0001 | 0.0252 | 0.985 | 0.1314 | <0.0001 | <0.0001 | <0.0001 |
| 12.2 | 1 | <0.0001 | 0.0031 | 0.9893 | 0.0076 | <0.0001 | <0.0001 | <0.0001 |
| 12.3667 | 1 | <0.0001 | 0.1906 | 1 | 0.3977 | <0.0001 | <0.0001 | <0.0001 |
| 12.5333 | 1 | <0.0001 | 0.2749 | 1 | 0.258 | <0.0001 | <0.0001 | <0.0001 |
| 12.7 | 1 | <0.0001 | 0.4441 | 0.9591 | 0.2591 | <0.0001 | <0.0001 | <0.0001 |
| 12.8667 | 1 | <0.0001 | 0.6198 | 0.9996 | 0.4692 | <0.0001 | <0.0001 | <0.0001 |
| 13.0333 | 1 | <0.0001 | 0.4573 | 0.9656 | 0.6877 | <0.0001 | <0.0001 | <0.0001 |
| 13.2 | 1 | <0.0001 | 0.0843 | 0.7689 | 0.6657 | <0.0001 | <0.0001 | <0.0001 |
| 13.3667 | 1 | <0.0001 | 0.0983 | 0.7804 | 0.5163 | <0.0001 | <0.0001 | <0.0001 |
| 13.5333 | 1 | <0.0001 | 0.061 | 0.741 | 0.2762 | <0.0001 | <0.0001 | <0.0001 |
| 13.7 | 1 | <0.0001 | 0.1499 | 0.4435 | 0.0475 | <0.0001 | <0.0001 | <0.0001 |
| 13.8667 | 1 | <0.0001 | 0.0299 | 0.1133 | 0.0039 | <0.0001 | <0.0001 | <0.0001 |
| 14.0333 | 1 | <0.0001 | 0.0169 | 0.0945 | 0.0003 | <0.0001 | <0.0001 | <0.0001 |
| 14.2 | 1 | <0.0001 | 0.0415 | 0.0948 | 0.0001 | <0.0001 | <0.0001 | <0.0001 |
| 14.3667 | 1 | <0.0001 | 0.0075 | 0.0604 | <0.0001 | <0.0001 | <0.0001 | <0.0001 |
| 14.5333 | 1 | <0.0001 | 0.0017 | 0.043 | <0.0001 | <0.0001 | <0.0001 | <0.0001 |
| 14.7 | 1 | <0.0001 | 0.0011 | 0.0987 | <0.0001 | <0.0001 | <0.0001 | <0.0001 |
| 14.8667 | 1 | <0.0001 | 0.0013 | 0.1932 | <0.0001 | <0.0001 | <0.0001 | <0.0001 |
| 15.0333 | 1 | <0.0001 | 0.003 | 0.2676 | <0.0001 | <0.0001 | <0.0001 | <0.0001 |

| win_starts | win_span | bio.ellipse | bio.rand | bio.scramb | bio.silhouette | ellipse.rand | ellipse.scramb | ellipse.silhouette |
|---|---|---|---|---|---|---|---|---|
| 15.2 | 1 | 0.0018 | 0.0002 | 0.0613 | <0.0001 | <0.0001 | <0.0001 | <0.0001 |
| 15.3667 | 1 | 0.0066 | <0.0001 | 0.001 | <0.0001 | <0.0001 | <0.0001 | <0.0001 |
| 15.5333 | 1 | 0.0012 | 0.0003 | 0.0022 | <0.0001 | <0.0001 | <0.0001 | <0.0001 |
| 15.7 | 1 | 0.003 | 0.0001 | 0.0001 | 0.0004 | <0.0001 | <0.0001 | <0.0001 |
| 15.8667 | 1 | 0.0666 | 0.0001 | <0.0001 | 0.002 | <0.0001 | <0.0001 | <0.0001 |
| 16.0333 | 1 | 0.0509 | 0.0006 | 0.0002 | 0.0002 | <0.0001 | <0.0001 | <0.0001 |
| 16.2 | 1 | 0.1911 | 0.0006 | 0.0001 | <0.0001 | <0.0001 | <0.0001 | <0.0001 |
| 16.3667 | 1 | 0.1338 | 0.0038 | 0.0058 | 0.0002 | <0.0001 | <0.0001 | <0.0001 |
| 16.5333 | 1 | 0.1491 | 0.0043 | 0.0066 | 0.0001 | <0.0001 | <0.0001 | <0.0001 |
| 16.7 | 1 | 0.109 | 0.0064 | 0.031 | <0.0001 | <0.0001 | <0.0001 | <0.0001 |
| 16.8667 | 1 | 0.0444 | 0.0018 | 0.0553 | 0.0002 | <0.0001 | <0.0001 | <0.0001 |
| 17.0333 | 1 | 0.0136 | 0.0038 | 0.1515 | 0.0013 | <0.0001 | <0.0001 | <0.0001 |
| 17.2 | 1 | 0.0214 | 0.0077 | 0.4981 | 0.0097 | <0.0001 | 0.0003 | <0.0001 |
| 17.3667 | 1 | 0.0225 | 0.0324 | 0.5211 | 0.0063 | <0.0001 | 0.001 | <0.0001 |
| 17.5333 | 1 | 0.0599 | 0.0239 | 0.4596 | 0.0062 | <0.0001 | 0.005 | <0.0001 |
| 17.7 | 1 | 0.0709 | 0.0626 | 0.5939 | 0.0171 | <0.0001 | 0.0131 | <0.0001 |
| 18.2 | 1 | 0.621 | 0.0158 | 0.3336 | 0.0045 | 0.0062 | 0.1524 | 0.0008 |
| 18.3667 | 1 | 0.6538 | 0.007 | 0.0492 | 0.0304 | 0.0098 | 0.0559 | 0.0008 |
| 18.7 | 1 | 0.5214 | 0.021 | 0.0181 | 0.3006 | 0.0144 | 0.0261 | 0.0076 |
| 20.0333 | 1 | 0.3504 | 0.0335 | 0.0081 | 0.3164 | 0.0036 | 0.0029 | 0.0374 |
| 20.2 | 1 | 0.7133 | 0.0018 | 0.0012 | 0.2609 | 0.0019 | 0.0016 | 0.068 |
| 20.3667 | 1 | 0.9154 | 0.0033 | 0.0196 | 0.1215 | 0.0082 | 0.0147 | 0.0268 |

| win_starts | win_span | bio.ellipse | bio.rand | bio.scramb | bio.silhouette | ellipse.rand | ellipse.scramb | ellipse.silhouette |
|---|---|---|---|---|---|---|---|---|
| 20.5333 | 1 | 0.7821 | 0.0269 | 0.1471 | 0.1192 | 0.004 | 0.0771 | 0.0057 |
| 20.7 | 1 | 0.7964 | 0.1643 | 0.1306 | 0.0544 | 0.0143 | 0.0638 | 0.0003 |
| 20.8667 | 1 | 0.7851 | 0.1793 | 0.6595 | 0.0309 | 0.0282 | 0.3108 | 0.0006 |
| 21.2 | 1 | 0.7852 | 0.089 | 0.9978 | 0.0487 | 0.036 | 0.6251 | 0.003 |
| 21.3667 | 1 | 0.8174 | 0.1117 | 0.9682 | 0.011 | 0.0297 | 0.433 | 0.0014 |
| 21.5333 | 1 | 0.8418 | 0.0765 | 0.9907 | 0.0203 | 0.0316 | 0.546 | 0.0087 |
| 21.7 | 1 | 0.9469 | 0.0079 | 0.9674 | 0.0334 | 0.0069 | 0.6431 | 0.0576 |
| 21.8667 | 1 | 0.9779 | 0.003 | 0.5607 | 0.0439 | 0.0031 | 0.3903 | 0.0738 |
| 22.0333 | 1 | 0.8945 | 0.0019 | 0.3729 | 0.0343 | 0.0006 | 0.1788 | 0.0344 |
| 22.2 | 1 | 0.5874 | 0.0022 | 0.2109 | 0.1376 | <0.0001 | 0.0126 | 0.0342 |
| 22.3667 | 1 | 0.0786 | 0.012 | 0.638 | 0.3634 | <0.0001 | 0.0071 | 0.0148 |
| 22.7 | 1 | 0.0133 | 0.088 | 0.0993 | 0.3764 | <0.0001 | 0.0003 | 0.0043 |
| 22.8667 | 1 | 0.0023 | 0.0985 | 0.4663 | 0.3645 | <0.0001 | 0.0012 | 0.0012 |
| 23.0333 | 1 | 0.0163 | 0.0043 | 0.6096 | 0.2465 | <0.0001 | 0.0043 | 0.0023 |
| 23.2 | 1 | 0.0401 | 0.0973 | 0.7079 | 0.1087 | <0.0001 | 0.0036 | 0.0002 |
| 23.3667 | 1 | 0.0795 | 0.0595 | 0.3782 | 0.0691 | <0.0001 | 0.0009 | <0.0001 |
| 23.5333 | 1 | 0.3502 | 0.0254 | 0.5136 | 0.0443 | <0.0001 | 0.0063 | 0.0002 |
| 23.7 | 1 | 0.2999 | 0.005 | 0.5512 | 0.0183 | <0.0001 | 0.0032 | 0.0001 |
| 23.8667 | 1 | 0.3734 | 0.0045 | 0.26 | 0.0126 | <0.0001 | 0.0001 | 0.0001 |
| 24.0333 | 1 | 0.097 | 0.0224 | 0.4724 | 0.0313 | <0.0001 | 0.0005 | 0.0001 |
| 24.2 | 1 | 0.2197 | 0.0809 | 0.2492 | 0.0697 | <0.0001 | 0.0002 | 0.0003 |
| 24.3667 | 1 | 0.404 | 0.0647 | 0.1672 | 0.1307 | 0.0002 | 0.0004 | 0.004 |

| win_starts | win_span | bio.ellipse | bio.rand | bio.scramb | bio.silhouette | ellipse.rand | ellipse.scramb | ellipse.silhouette |
|---|---|---|---|---|---|---|---|---|
| 24.5333 | 1 | 0.3368 | 0.1166 | 0.0465 | 0.1401 | 0.0001 | <0.0001 | 0.0042 |
| 24.8667 | 1 | 0.2131 | 0.9878 | 0.3913 | 0.0632 | 0.1026 | 0.0164 | 0.0005 |

### Distance of retinal midpoint from stimulus center - Sliding window GLMM for Static Section

| win_starts | win_span | p |
|---|---|---|
| 0.03333 | 1 | 1 |
| 0.2 | 1 | 1 |
| 0.36667 | 1 | 1 |
| 0.53333 | 1 | 1 |
| 0.7 | 1 | 1 |
| 0.86667 | 1 | 1 |
| 1.03333 | 1 | 1 |
| 1.2 | 1 | 1 |
| 1.36667 | 1 | 1 |
| 1.53333 | 1 | 1 |
| 1.7 | 1 | 1 |
| 1.86667 | 1 | 1 |
| 2.03333 | 1 | 1 |
| 2.2 | 1 | 1 |
| 2.36667 | 1 | 1 |
| 2.53333 | 1 | 1 |
| 2.7 | 1 | 1 |
| 2.86667 | 1 | 1 |
| 3.03333 | 1 | 1 |
| 3.2 | 1 | 1 |

| win_starts | win_span | p |
|---|---|---|
| 3.36667 | 1 | 1 |
| 3.53333 | 1 | 1 |
| 3.7 | 1 | 1 |
| 3.86667 | 1 | 0.4339 |
| 4.03333 | 1 | 0.25167 |
| 4.2 | 1 | 1 |
| 4.36667 | 1 | 1 |
| 4.53333 | 1 | 1 |
| 4.7 | 1 | 1 |
| 4.86667 | 1 | 1 |
| 5.03333 | 1 | 0.59184 |
| 5.2 | 1 | 1 |
| 5.36667 | 1 | 1 |
| 5.53333 | 1 | 1 |
| 5.7 | 1 | 1 |
| 5.86667 | 1 | 1 |
| 6.03333 | 1 | 1 |
| 6.2 | 1 | 1 |
| 6.36667 | 1 | 1 |
| 6.53333 | 1 | 1 |
| 6.7 | 1 | 1 |
| 6.86667 | 1 | 1 |

| win_starts | win_span | p |
|---|---|---|
| 7.03333 | 1 | 1 |
| 7.2 | 1 | 1 |
| 7.36667 | 1 | 1 |
| 7.53333 | 1 | 1 |
| 7.7 | 1 | 1 |
| 7.86667 | 1 | 1 |
| 8.03333 | 1 | 1 |
| 8.2 | 1 | 1 |
| 8.36667 | 1 | 1 |
| 8.53333 | 1 | 1 |
| 8.7 | 1 | 1 |
| 8.86667 | 1 | 1 |
| 9.03333 | 1 | 1 |
| 9.2 | 1 | 1 |
| 9.36667 | 1 | 1 |
| 9.53333 | 1 | 1 |
| 9.7 | 1 | 1 |
| 9.86667 | 1 | 1 |
| 10.03333 | 1 | 1 |
| 10.2 | 1 | 1 |
| 10.36667 | 1 | 1 |
| 10.53333 | 1 | 1 |

| win_starts | win_span | p |
| --- | --- | --- |
| 10.7 | 1 | 1 |
| 10.86667 | 1 | 0.46112 |

### Distance of retinal midpoint from stimulus center - Post-Hoc Comparisons for Static Section

| win_starts | win_span | bio.ellipse | bio.none | bio.rand | bio.scramb | bio.silhouette | ellipse.none | none.rand | none.scramb | none.silhouette |
|---|---|---|---|---|---|---|---|---|---|---|

### Distance of retinal midpoint from stimulus center - Sliding window GLMM for all stimulus duration

| win_starts | win_span | p |
|---|---|---|
| 0.03333 | 1 | 1 |
| 0.2 | 1 | 1 |
| 0.36667 | 1 | 1 |
| 0.53333 | 1 | 1 |
| 0.7 | 1 | 1 |
| 0.86667 | 1 | 1 |
| 1.03333 | 1 | 1 |
| 1.2 | 1 | 1 |
| 1.36667 | 1 | 1 |
| 1.53333 | 1 | 1 |
| 1.7 | 1 | 1 |
| 1.86667 | 1 | 1 |
| 2.03333 | 1 | 1 |
| 2.2 | 1 | 1 |
| 2.36667 | 1 | 1 |
| 2.53333 | 1 | 1 |
| 2.7 | 1 | 1 |
| 2.86667 | 1 | 1 |
| 3.03333 | 1 | 1 |
| 3.2 | 1 | 1 |

| win_starts | win_span | p |
| --- | --- | --- |
| 3.36667 | 1 | 1 |
| 3.53333 | 1 | 1 |
| 3.7 | 1 | 1 |
| 3.86667 | 1 | 1 |
| 4.03333 | 1 | 1 |
| 4.2 | 1 | 1 |
| 4.36667 | 1 | 1 |
| 4.53333 | 1 | 1 |
| 4.7 | 1 | 1 |
| 4.86667 | 1 | 1 |
| 5.03333 | 1 | 1 |
| 5.2 | 1 | 1 |
| 5.36667 | 1 | 1 |
| 5.53333 | 1 | 1 |
| 5.7 | 1 | 1 |
| 5.86667 | 1 | 1 |
| 6.03333 | 1 | 1 |
| 6.2 | 1 | 1 |
| 6.36667 | 1 | 1 |
| 6.53333 | 1 | 1 |
| 6.7 | 1 | 1 |
| 6.86667 | 1 | 1 |

| win_starts | win_span | p |
|---|---|---|
| 7.03333 | 1 | 1 |
| 7.2 | 1 | 1 |
| 7.36667 | 1 | 1 |
| 7.53333 | 1 | 1 |
| 7.7 | 1 | 1 |
| 7.86667 | 1 | 1 |
| 8.03333 | 1 | 1 |
| 8.2 | 1 | 1 |
| 8.36667 | 1 | 1 |
| 8.53333 | 1 | 1 |
| 8.7 | 1 | 1 |
| 8.86667 | 1 | 1 |
| 9.03333 | 1 | 1 |
| 9.2 | 1 | 1 |
| 9.36667 | 1 | 1 |
| 9.53333 | 1 | 1 |
| 9.7 | 1 | 1 |
| 9.86667 | 1 | 1 |
| 10.03333 | 1 | 1 |
| 10.2 | 1 | 1 |
| 10.36667 | 1 | 1 |
| 10.53333 | 1 | 1 |

| win_starts | win_span | p |
|---|---|---|
| 10.7 | 1 | 1 |
| 10.86667 | 1 | 1 |
| 11.03333 | 1 | 1 |
| 11.2 | 1 | 1 |
| 11.36667 | 1 | 1 |
| 11.53333 | 1 | 1 |
| 11.7 | 1 | 0.03745 |
| 11.86667 | 1 | <0.0001 |
| 12.03333 | 1 | <0.0001 |
| 12.2 | 1 | <0.0001 |
| 12.36667 | 1 | <0.0001 |
| 12.53333 | 1 | <0.0001 |
| 12.7 | 1 | <0.0001 |
| 12.86667 | 1 | <0.0001 |
| 13.03333 | 1 | 0.00011 |
| 13.2 | 1 | 0.00114 |
| 13.36667 | 1 | 0.0032 |
| 13.53333 | 1 | 0.00025 |
| 13.7 | 1 | <0.0001 |
| 13.86667 | 1 | <0.0001 |
| 14.03333 | 1 | <0.0001 |
| 14.2 | 1 | 0.00013 |

| win_starts | win_span | p |
|---|---|---|
| 14.36667 | 1 | 0.00144 |
| 14.53333 | 1 | 0.0166 |
| 14.7 | 1 | 0.09145 |
| 14.86667 | 1 | 1 |
| 15.03333 | 1 | 1 |
| 15.2 | 1 | 1 |
| 15.36667 | 1 | 1 |
| 15.53333 | 1 | 1 |
| 15.7 | 1 | 1 |
| 15.86667 | 1 | 1 |
| 16.03333 | 1 | 1 |
| 16.2 | 1 | 1 |
| 16.36667 | 1 | 1 |
| 16.53333 | 1 | 1 |
| 16.7 | 1 | 1 |
| 16.86667 | 1 | 1 |
| 17.03333 | 1 | 1 |
| 17.2 | 1 | 1 |
| 17.36667 | 1 | 1 |
| 17.53333 | 1 | 1 |
| 17.7 | 1 | 1 |
| 17.86667 | 1 | 1 |

| win_starts | win_span | p |
|---|---|---|
| 18.03333 | 1 | 1 |
| 18.2 | 1 | 1 |
| 18.36667 | 1 | 1 |
| 18.53333 | 1 | 1 |
| 18.7 | 1 | 1 |
| 18.86667 | 1 | 1 |
| 19.03333 | 1 | 1 |
| 19.2 | 1 | 1 |
| 19.36667 | 1 | 1 |
| 19.53333 | 1 | 1 |
| 19.7 | 1 | 1 |
| 19.86667 | 1 | 1 |
| 20.03333 | 1 | 1 |
| 20.2 | 1 | 1 |
| 20.36667 | 1 | 1 |
| 20.53333 | 1 | 1 |
| 20.7 | 1 | 1 |
| 20.86667 | 1 | 1 |
| 21.03333 | 1 | 1 |
| 21.2 | 1 | 1 |
| 21.36667 | 1 | 1 |
| 21.53333 | 1 | 1 |

| win_starts | win_span | p |
|---|---|---|
| 21.7 | 1 | 1 |
| 21.86667 | 1 | 1 |
| 22.03333 | 1 | 1 |
| 22.2 | 1 | 1 |
| 22.36667 | 1 | 1 |
| 22.53333 | 1 | 1 |
| 22.7 | 1 | 1 |
| 22.86667 | 1 | 1 |
| 23.03333 | 1 | 1 |
| 23.2 | 1 | 1 |
| 23.36667 | 1 | 1 |
| 23.53333 | 1 | 1 |
| 23.7 | 1 | 1 |
| 23.86667 | 1 | 1 |
| 24.03333 | 1 | 1 |
| 24.2 | 1 | 1 |
| 24.36667 | 1 | 1 |
| 24.53333 | 1 | 1 |
| 24.7 | 1 | 1 |
| 24.86667 | 1 | 1 |
| 25.03333 | 1 | 1 |
| 25.2 | 1 | 1 |

| win_starts | win_span | p |
|---|---|---|
| 25.36667 | 1 | 1 |
| 25.53333 | 1 | 1 |
| 25.7 | 1 | 1 |
| 25.86667 | 1 | 1 |
| 26.03333 | 1 | 1 |
| 26.2 | 1 | 1 |
| 26.36667 | 1 | 1 |
| 26.53333 | 1 | 1 |
| 26.7 | 1 | 1 |
| 26.86667 | 1 | 1 |
| 27.03333 | 1 | 1 |
| 27.2 | 1 | 1 |
| 27.36667 | 1 | 1 |
| 27.53333 | 1 | 1 |
| 27.7 | 1 | 1 |
| 27.86667 | 1 | 1 |
| 28.03333 | 1 | 1 |
| 28.2 | 1 | 1 |
| 28.36667 | 1 | 1 |
| 28.53333 | 1 | 1 |
| 28.7 | 1 | 1 |
| 28.86667 | 1 | 1 |

| win_starts | win_span | p |
|---|---|---|
| 29.03333 | 1 | 1 |
| 29.2 | 1 | 1 |
| 29.36667 | 1 | 1 |
| 29.53333 | 1 | 1 |
| 29.7 | 1 | 1 |
| 29.86667 | 1 | 1 |
| 30.03333 | 1 | 1 |
| 30.2 | 1 | 1 |
| 30.36667 | 1 | 1 |
| 30.53333 | 1 | 1 |
| 30.7 | 1 | 1 |
| 30.86667 | 1 | 1 |
| 31.03333 | 1 | 1 |
| 31.2 | 1 | 1 |
| 31.36667 | 1 | 1 |
| 31.53333 | 1 | 1 |
| 31.7 | 1 | 1 |
| 31.86667 | 1 | 1 |
| 32.03333 | 1 | 1 |
| 32.2 | 1 | 1 |
| 32.36667 | 1 | 1 |
| 32.53333 | 1 | 0.79157 |

| win_starts | win_span | p |
|---|---|---|
| 32.7 | 1 | 0.21655 |
| 32.86667 | 1 | 0.17937 |
| 33.03333 | 1 | 0.98574 |
| 33.2 | 1 | 1 |
| 33.36667 | 1 | 1 |
| 33.53333 | 1 | 1 |
| 33.7 | 1 | 1 |
| 33.86667 | 1 | 1 |
| 34.03333 | 1 | 1 |
| 34.2 | 1 | 1 |
| 34.36667 | 1 | 1 |
| 34.53333 | 1 | 1 |
| 34.7 | 1 | 1 |
| 34.86667 | 1 | 1 |
| 35.03333 | 1 | 1 |
| 35.2 | 1 | 1 |

### Distance of retinal midpoint from stimulus center - Post-Hoc Comparisons for all stimulus duration

| win_starts | win_span | bio.ellipse | bio.rand | bio.scramb | bio.silhouette | ellipse.rand | ellipse.scramb | ellipse.silhouette |
|---|---|---|---|---|---|---|---|---|
| 11.8667 | 1 | <0.0001 | 0.1534 | 0.672 | 0.6835 | <0.0001 | <0.0001 | <0.0001 |
| 12.0333 | 1 | <0.0001 | 0.1209 | 0.5961 | 0.4439 | <0.0001 | <0.0001 | <0.0001 |
| 12.2 | 1 | <0.0001 | 0.1499 | 0.874 | 0.0876 | <0.0001 | <0.0001 | <0.0001 |
| 12.3667 | 1 | <0.0001 | 0.8266 | 0.921 | 0.4775 | <0.0001 | <0.0001 | <0.0001 |
| 12.5333 | 1 | <0.0001 | 0.9631 | 0.969 | 0.5405 | <0.0001 | <0.0001 | <0.0001 |
| 12.7 | 1 | <0.0001 | 1 | 0.9973 | 0.8163 | <0.0001 | <0.0001 | <0.0001 |
| 12.8667 | 1 | <0.0001 | 0.8594 | 0.9748 | 0.9995 | <0.0001 | <0.0001 | <0.0001 |
| 13.0333 | 1 | <0.0001 | 0.9959 | 1 | 0.9999 | <0.0001 | <0.0001 | <0.0001 |
| 13.2 | 1 | <0.0001 | 0.9814 | 0.9427 | 0.9998 | <0.0001 | <0.0001 | <0.0001 |
| 13.3667 | 1 | <0.0001 | 0.8361 | 0.933 | 0.9856 | <0.0001 | <0.0001 | <0.0001 |
| 13.5333 | 1 | <0.0001 | 0.8505 | 0.9378 | 0.8776 | <0.0001 | <0.0001 | <0.0001 |
| 13.7 | 1 | <0.0001 | 0.8652 | 0.8071 | 0.657 | <0.0001 | <0.0001 | <0.0001 |
| 13.8667 | 1 | <0.0001 | 0.6703 | 0.9894 | 0.5977 | <0.0001 | <0.0001 | <0.0001 |
| 14.0333 | 1 | <0.0001 | 0.5241 | 0.9979 | 0.3801 | <0.0001 | 0.0001 | <0.0001 |
| 14.2 | 1 | <0.0001 | 0.5138 | 0.9937 | 0.3693 | <0.0001 | <0.0001 | <0.0001 |
| 14.3667 | 1 | <0.0001 | 0.3616 | 0.9752 | 0.2647 | <0.0001 | <0.0001 | <0.0001 |

### Inter-retinal distance - Sliding window GLMM for Static Section

| win_starts | win_span | p |
|---|---|---|
| 0.03333 | 1 | 1 |
| 0.2 | 1 | 1 |
| 0.36667 | 1 | 1 |
| 0.53333 | 1 | 1 |
| 0.7 | 1 | 1 |
| 0.86667 | 1 | 1 |
| 1.03333 | 1 | 1 |
| 1.2 | 1 | 1 |
| 1.36667 | 1 | 1 |
| 1.53333 | 1 | 1 |
| 1.7 | 1 | 1 |
| 1.86667 | 1 | 1 |
| 2.03333 | 1 | 1 |
| 2.2 | 1 | 1 |
| 2.36667 | 1 | 1 |
| 2.53333 | 1 | 1 |
| 2.7 | 1 | 1 |
| 2.86667 | 1 | 1 |
| 3.03333 | 1 | 1 |
| 3.2 | 1 | 1 |

| win_starts | win_span | p |
|---|---|---|
| 3.36667 | 1 | 1 |
| 3.53333 | 1 | 1 |
| 3.7 | 1 | 1 |
| 3.86667 | 1 | 1 |
| 4.03333 | 1 | 1 |
| 4.2 | 1 | 1 |
| 4.36667 | 1 | 1 |
| 4.53333 | 1 | 1 |
| 4.7 | 1 | 1 |
| 4.86667 | 1 | 1 |
| 5.03333 | 1 | 1 |
| 5.2 | 1 | 1 |
| 5.36667 | 1 | 1 |
| 5.53333 | 1 | 1 |
| 5.7 | 1 | 1 |
| 5.86667 | 1 | 1 |
| 6.03333 | 1 | 1 |
| 6.2 | 1 | 1 |
| 6.36667 | 1 | 1 |
| 6.53333 | 1 | 1 |
| 6.7 | 1 | 1 |
| 6.86667 | 1 | 1 |

| win_starts | win_span | p |
|---|---|---|
| 7.03333 | 1 | 1 |
| 7.2 | 1 | 1 |
| 7.36667 | 1 | 1 |
| 7.53333 | 1 | 1 |
| 7.7 | 1 | 1 |
| 7.86667 | 1 | 1 |
| 8.03333 | 1 | 1 |
| 8.2 | 1 | 1 |
| 8.36667 | 1 | 1 |
| 8.53333 | 1 | 1 |
| 8.7 | 1 | 1 |
| 8.86667 | 1 | 1 |
| 9.03333 | 1 | 1 |
| 9.2 | 1 | 1 |
| 9.36667 | 1 | 1 |
| 9.53333 | 1 | 1 |
| 9.7 | 1 | 1 |
| 9.86667 | 1 | 1 |
| 10.03333 | 1 | 1 |
| 10.2 | 1 | 1 |
| 10.36667 | 1 | 1 |
| 10.53333 | 1 | 1 |

| win_starts | win_span | p |
|---|---|---|
| 10.7 | 1 | 1 |
| 10.86667 | 1 | 1 |

### Inter-retinal distance - Post-Hoc Comparisons for Static Section

| win_starts | win_span | bio.ellipse | bio.none | bio.rand | bio.scramb | bio.silhouette | ellipse.none | none.rand | none.scramb | none.silhouette |
|---|---|---|---|---|---|---|---|---|---|---|

### Inter-retinal distance - Sliding window GLMM for all stimulus duration

| win_starts | win_span | p |
|---|---|---|
| 0.03333 | 1 | 1 |
| 0.2 | 1 | 1 |
| 0.36667 | 1 | 1 |
| 0.53333 | 1 | 1 |
| 0.7 | 1 | 1 |
| 0.86667 | 1 | 1 |
| 1.03333 | 1 | 1 |
| 1.2 | 1 | 1 |
| 1.36667 | 1 | 1 |
| 1.53333 | 1 | 1 |
| 1.7 | 1 | 1 |
| 1.86667 | 1 | 1 |
| 2.03333 | 1 | 1 |
| 2.2 | 1 | 1 |
| 2.36667 | 1 | 1 |
| 2.53333 | 1 | 1 |
| 2.7 | 1 | 1 |
| 2.86667 | 1 | 1 |
| 3.03333 | 1 | 1 |
| 3.2 | 1 | 1 |

| win_starts | win_span | p |
| --- | --- | --- |
| 3.36667 | 1 | 1 |
| 3.53333 | 1 | 1 |
| 3.7 | 1 | 1 |
| 3.86667 | 1 | 1 |
| 4.03333 | 1 | 1 |
| 4.2 | 1 | 1 |
| 4.36667 | 1 | 1 |
| 4.53333 | 1 | 1 |
| 4.7 | 1 | 1 |
| 4.86667 | 1 | 1 |
| 5.03333 | 1 | 1 |
| 5.2 | 1 | 1 |
| 5.36667 | 1 | 1 |
| 5.53333 | 1 | 1 |
| 5.7 | 1 | 1 |
| 5.86667 | 1 | 1 |
| 6.03333 | 1 | 1 |
| 6.2 | 1 | 1 |
| 6.36667 | 1 | 1 |
| 6.53333 | 1 | 1 |
| 6.7 | 1 | 1 |
| 6.86667 | 1 | 1 |

| win_starts | win_span | p |
|---|---|---|
| 7.03333 | 1 | 1 |
| 7.2 | 1 | 1 |
| 7.36667 | 1 | 1 |
| 7.53333 | 1 | 1 |
| 7.7 | 1 | 1 |
| 7.86667 | 1 | 1 |
| 8.03333 | 1 | 1 |
| 8.2 | 1 | 1 |
| 8.36667 | 1 | 1 |
| 8.53333 | 1 | 1 |
| 8.7 | 1 | 1 |
| 8.86667 | 1 | 1 |
| 9.03333 | 1 | 1 |
| 9.2 | 1 | 1 |
| 9.36667 | 1 | 1 |
| 9.53333 | 1 | 1 |
| 9.7 | 1 | 1 |
| 9.86667 | 1 | 1 |
| 10.03333 | 1 | 1 |
| 10.2 | 1 | 1 |
| 10.36667 | 1 | 1 |
| 10.53333 | 1 | 1 |

| win_starts | win_span | p |
|---|---|---|
| 10.7 | 1 | 1 |
| 10.86667 | 1 | 1 |
| 11.03333 | 1 | 1 |
| 11.2 | 1 | 1 |
| 11.36667 | 1 | 1 |
| 11.53333 | 1 | 1 |
| 11.7 | 1 | 0.1842 |
| 11.86667 | 1 | 0.00037 |
| 12.03333 | 1 | <0.0001 |
| 12.2 | 1 | <0.0001 |
| 12.36667 | 1 | <0.0001 |
| 12.53333 | 1 | <0.0001 |
| 12.7 | 1 | <0.0001 |
| 12.86667 | 1 | <0.0001 |
| 13.03333 | 1 | <0.0001 |
| 13.2 | 1 | <0.0001 |
| 13.36667 | 1 | 0.00071 |
| 13.53333 | 1 | <0.0001 |
| 13.7 | 1 | <0.0001 |
| 13.86667 | 1 | <0.0001 |
| 14.03333 | 1 | <0.0001 |
| 14.2 | 1 | <0.0001 |

| win_starts | win_span | p |
|---|---|---|
| 14.36667 | 1 | <0.0001 |
| 14.53333 | 1 | <0.0001 |
| 14.7 | 1 | 0.08252 |
| 14.86667 | 1 | 0.51881 |
| 15.03333 | 1 | 0.91988 |
| 15.2 | 1 | 1 |
| 15.36667 | 1 | 0.4854 |
| 15.53333 | 1 | 0.37499 |
| 15.7 | 1 | 0.00115 |
| 15.86667 | 1 | 0.00085 |
| 16.03333 | 1 | 0.00275 |
| 16.2 | 1 | 0.00087 |
| 16.36667 | 1 | 0.00173 |
| 16.53333 | 1 | 0.01126 |
| 16.7 | 1 | 0.10069 |
| 16.86667 | 1 | 0.59841 |
| 17.03333 | 1 | 0.49827 |
| 17.2 | 1 | 0.34635 |
| 17.36667 | 1 | 0.03253 |
| 17.53333 | 1 | 0.00471 |
| 17.7 | 1 | 0.00747 |
| 17.86667 | 1 | 0.06161 |

| win_starts | win_span | p |
|---|---|---|
| 18.03333 | 1 | 1 |
| 18.2 | 1 | 1 |
| 18.36667 | 1 | 1 |
| 18.53333 | 1 | 0.78263 |
| 18.7 | 1 | 0.04574 |
| 18.86667 | 1 | 0.01871 |
| 19.03333 | 1 | 0.00708 |
| 19.2 | 1 | 0.00049 |
| 19.36667 | 1 | <0.0001 |
| 19.53333 | 1 | 0.00067 |
| 19.7 | 1 | 0.00677 |
| 19.86667 | 1 | 0.01071 |
| 20.03333 | 1 | 0.54348 |
| 20.2 | 1 | 1 |
| 20.36667 | 1 | 1 |
| 20.53333 | 1 | 0.85141 |
| 20.7 | 1 | 0.46271 |
| 20.86667 | 1 | 1 |
| 21.03333 | 1 | 0.60698 |
| 21.2 | 1 | 0.18681 |
| 21.36667 | 1 | 0.03953 |
| 21.53333 | 1 | 0.03242 |

| win_starts | win_span | p |
|---|---|---|
| 21.7 | 1 | 0.04201 |
| 21.86667 | 1 | 0.23283 |
| 22.03333 | 1 | 0.99102 |
| 22.2 | 1 | 1 |
| 22.36667 | 1 | 1 |
| 22.53333 | 1 | 1 |
| 22.7 | 1 | 1 |
| 22.86667 | 1 | 1 |
| 23.03333 | 1 | 1 |
| 23.2 | 1 | 1 |
| 23.36667 | 1 | 1 |
| 23.53333 | 1 | 1 |
| 23.7 | 1 | 1 |
| 23.86667 | 1 | 1 |
| 24.03333 | 1 | 1 |
| 24.2 | 1 | 0.572 |
| 24.36667 | 1 | 1 |
| 24.53333 | 1 | 1 |
| 24.7 | 1 | 1 |
| 24.86667 | 1 | 1 |
| 25.03333 | 1 | 1 |
| 25.2 | 1 | 1 |

| win_starts | win_span | p |
|---|---|---|
| 25.36667 | 1 | 1 |
| 25.53333 | 1 | 1 |
| 25.7 | 1 | 0.3427 |
| 25.86667 | 1 | 1 |
| 26.03333 | 1 | 0.28634 |
| 26.2 | 1 | 0.95009 |
| 26.36667 | 1 | 1 |
| 26.53333 | 1 | 0.2956 |
| 26.7 | 1 | 1 |
| 26.86667 | 1 | 1 |
| 27.03333 | 1 | 1 |
| 27.2 | 1 | 0.66428 |
| 27.36667 | 1 | 0.68881 |
| 27.53333 | 1 | 0.43173 |
| 27.7 | 1 | 1 |
| 27.86667 | 1 | 1 |
| 28.03333 | 1 | 1 |
| 28.2 | 1 | 1 |
| 28.36667 | 1 | 1 |
| 28.53333 | 1 | 1 |
| 28.7 | 1 | 1 |
| 28.86667 | 1 | 1 |

| win_starts | win_span | p |
|---|---|---|
| 29.03333 | 1 | 1 |
| 29.2 | 1 | 1 |
| 29.36667 | 1 | 1 |
| 29.53333 | 1 | 1 |
| 29.7 | 1 | 1 |
| 29.86667 | 1 | 1 |
| 30.03333 | 1 | 1 |
| 30.2 | 1 | 1 |
| 30.36667 | 1 | 1 |
| 30.53333 | 1 | 1 |
| 30.7 | 1 | 1 |
| 30.86667 | 1 | 1 |
| 31.03333 | 1 | 1 |
| 31.2 | 1 | 1 |
| 31.36667 | 1 | 1 |
| 31.53333 | 1 | 1 |
| 31.7 | 1 | 1 |
| 31.86667 | 1 | 1 |
| 32.03333 | 1 | 1 |
| 32.2 | 1 | 1 |
| 32.36667 | 1 | 1 |
| 32.53333 | 1 | 1 |

| win_starts | win_span | p |
|---|---|---|
| 32.7 | 1 | 1 |
| 32.86667 | 1 | 1 |
| 33.03333 | 1 | 1 |
| 33.2 | 1 | 1 |
| 33.36667 | 1 | 1 |
| 33.53333 | 1 | 1 |
| 33.7 | 1 | 1 |
| 33.86667 | 1 | 1 |
| 34.03333 | 1 | 1 |
| 34.2 | 1 | 1 |
| 34.36667 | 1 | 1 |
| 34.53333 | 1 | 1 |
| 34.7 | 1 | 1 |
| 34.86667 | 1 | 1 |
| 35.03333 | 1 | 1 |
| 35.2 | 1 | 1 |

### Inter-retinal distance - Post-Hoc Comparisons for all stimulus duration

| win_starts | win_span | bio.ellipse | bio.rand | bio.scramb | bio.silhouette | ellipse.rand | ellipse.scramb | ellipse.silhouette |
|---|---|---|---|---|---|---|---|---|
| 11.8667 | 1 | 0.0067 | 0.0001 | 0.9441 | 0.9915 | <0.0001 | 0.008 | 0.0002 |
| 12.0333 | 1 | 0.0094 | <0.0001 | 0.8153 | 0.9977 | <0.0001 | 0.0035 | 0.0009 |
| 12.2 | 1 | 0.0129 | <0.0001 | 0.6527 | 0.999 | <0.0001 | 0.0029 | 0.0115 |
| 12.3667 | 1 | 0.0373 | <0.0001 | 0.6925 | 0.9998 | <0.0001 | 0.0075 | 0.0316 |
| 12.5333 | 1 | 0.1181 | <0.0001 | 0.733 | 0.9999 | <0.0001 | 0.0287 | 0.0704 |
| 12.7 | 1 | 0.5991 | <0.0001 | 0.9529 | 0.9997 | <0.0001 | 0.2832 | 0.5423 |
| 12.8667 | 1 | 0.9961 | <0.0001 | 0.711 | 0.8324 | <0.0001 | 0.7507 | 0.6874 |
| 13.0333 | 1 | 0.9998 | <0.0001 | 0.7002 | 0.6335 | 0.0002 | 0.9275 | 0.6986 |
| 13.2 | 1 | 1 | 0.0001 | 0.8031 | 0.4817 | 0.0001 | 0.9447 | 0.4562 |
| 13.3667 | 1 | 1 | 0.0002 | 0.6796 | 0.1682 | 0.0004 | 0.8343 | 0.0722 |
| 13.5333 | 1 | 0.9998 | <0.0001 | 0.4209 | 0.0258 | <0.0001 | 0.6733 | 0.0115 |
| 13.7 | 1 | 0.9861 | <0.0001 | 0.4528 | 0.001 | <0.0001 | 0.4228 | 0.0005 |
| 13.8667 | 1 | 0.5274 | 0.0019 | 0.6977 | 0.0081 | <0.0001 | 0.1377 | 0.0006 |
| 14.0333 | 1 | 0.3861 | 0.0093 | 0.5109 | 0.0571 | <0.0001 | 0.0266 | 0.0037 |
| 14.2 | 1 | 0.148 | 0.0196 | 0.6327 | 0.1907 | <0.0001 | 0.004 | 0.0042 |
| 14.3667 | 1 | 0.1013 | 0.0399 | 0.6085 | 0.5418 | <0.0001 | 0.001 | 0.0181 |
| 14.5333 | 1 | 0.028 | 0.1333 | 0.9101 | 0.8947 | <0.0001 | 0.0022 | 0.0363 |
| 15.7 | 1 | 0.0035 | 0.0113 | 0.7243 | 0.9978 | <0.0001 | 0.3186 | 0.0811 |
| 15.8667 | 1 | 0.0057 | 0.0146 | 0.7372 | 1 | <0.0001 | 0.3051 | 0.0469 |
| 16.0333 | 1 | 0.0036 | 0.0334 | 0.7044 | 1 | <0.0001 | 0.181 | 0.0405 |

| win_starts | win_span | bio.ellipse | bio.rand | bio.scramb | bio.silhouette | ellipse.rand | ellipse.scramb | ellipse.silhouette |
|---|---|---|---|---|---|---|---|---|
| 16.2 | 1 | 0.0016 | 0.0637 | 0.5513 | 1 | <0.0001 | 0.1304 | 0.0079 |
| 16.3667 | 1 | 0.0052 | 0.134 | 0.8734 | 0.9969 | <0.0001 | 0.1187 | 0.0017 |
| 17.5333 | 1 | 0.4218 | 0.0018 | 0.1388 | 0.2108 | 0.0002 | 0.0273 | 0.0028 |
| 17.7 | 1 | 0.3426 | 0.0008 | 0.2735 | 0.1305 | <0.0001 | 0.0216 | 0.001 |
| 19.0333 | 1 | 0.0052 | 0.0317 | 0.3268 | 0.4899 | 0.0004 | 0.0069 | 0.0155 |
| 19.2 | 1 | 0.0033 | 0.0038 | 0.367 | 0.6747 | <0.0001 | 0.007 | 0.0459 |
| 19.3667 | 1 | 0.0626 | 0.0012 | 0.7358 | 0.3845 | <0.0001 | 0.107 | 0.0372 |
| 19.5333 | 1 | 0.2304 | 0.0014 | 0.5464 | 0.4292 | <0.0001 | 0.0781 | 0.0519 |
| 19.7 | 1 | 0.483 | 0.0012 | 0.6054 | 0.6514 | <0.0001 | 0.1684 | 0.1768 |

### Speed of change in inter-retinal distance - Sliding window GLMM for Static Section

| win_starts | win_span | p |
|---|---|---|
| 0.03333 | 1 | <0.0001 |
| 0.2 | 1 | <0.0001 |
| 0.36667 | 1 | <0.0001 |
| 0.53333 | 1 | <0.0001 |
| 0.7 | 1 | <0.0001 |
| 0.86667 | 1 | <0.0001 |
| 1.03333 | 1 | <0.0001 |
| 1.2 | 1 | <0.0001 |
| 1.36667 | 1 | <0.0001 |
| 1.53333 | 1 | 0.00047 |
| 1.7 | 1 | 0.00888 |
| 1.86667 | 1 | 0.09016 |
| 2.03333 | 1 | 0.07613 |
| 2.2 | 1 | 0.01214 |
| 2.36667 | 1 | 0.00823 |
| 2.53333 | 1 | 0.0065 |
| 2.7 | 1 | 0.27783 |
| 2.86667 | 1 | 1 |
| 3.03333 | 1 | 1 |
| 3.2 | 1 | 0.12368 |

| win_starts | win_span | p |
|---|---|---|
| 3.36667 | 1 | 1 |
| 3.53333 | 1 | 1 |
| 3.7 | 1 | 1 |
| 3.86667 | 1 | 1 |
| 4.03333 | 1 | 1 |
| 4.2 | 1 | 1 |
| 4.36667 | 1 | 0.15872 |
| 4.53333 | 1 | 0.27154 |
| 4.7 | 1 | 0.23708 |
| 4.86667 | 1 | 0.13073 |
| 5.03333 | 1 | 0.05414 |
| 5.2 | 1 | 0.15109 |
| 5.36667 | 1 | 0.00553 |
| 5.53333 | 1 | 0.04573 |
| 5.7 | 1 | 0.08291 |
| 5.86667 | 1 | 0.00951 |
| 6.03333 | 1 | 0.07284 |
| 6.2 | 1 | 0.09024 |
| 6.36667 | 1 | 1 |
| 6.53333 | 1 | 1 |
| 6.7 | 1 | 1 |
| 6.86667 | 1 | 1 |

| win_starts | win_span | p |
|---|---|---|
| 7.03333 | 1 | 1 |
| 7.2 | 1 | 1 |
| 7.36667 | 1 | 1 |
| 7.53333 | 1 | 1 |
| 7.7 | 1 | 0.8583 |
| 7.86667 | 1 | 1 |
| 8.03333 | 1 | 1 |
| 8.2 | 1 | 1 |
| 8.36667 | 1 | 0.99749 |
| 8.53333 | 1 | 0.8158 |
| 8.7 | 1 | 1 |
| 8.86667 | 1 | 1 |
| 9.03333 | 1 | 1 |
| 9.2 | 1 | 1 |
| 9.36667 | 1 | 1 |
| 9.53333 | 1 | 1 |
| 9.7 | 1 | 1 |
| 9.86667 | 1 | 1 |
| 10.03333 | 1 | 1 |
| 10.2 | 1 | 1 |
| 10.36667 | 1 | 1 |
| 10.53333 | 1 | 1 |

| win_starts | win_span | p |
|---|---|---|
| 10.7 | 1 | 1 |
| 10.86667 | 1 | 1 |

### Speed of change in inter-retinal distance - Post-Hoc Comparisons for Static Section

| win_starts | win_span | bio.ellipse | bio.none | bio.rand | bio.scramb | bio.silhouette | ellipse.none | none.rand | none.scramb | none.silhouette |
|---|---|---|---|---|---|---|---|---|---|---|
| 0.0333 | 1 | 0.9122 | 0.0171 | 0.9991 | 0.9999 | 0.3349 | 0.0965 | 0.1581 | 0.0055 | <0.0001 |
| 0.2 | 1 | 0.8738 | 0.0012 | 0.9998 | 1 | 0.285 | 0.05 | 0.0348 | 0.0033 | <0.0001 |
| 0.3667 | 1 | 0.8023 | 0.0003 | 0.9681 | 0.9922 | 0.835 | 0.011 | 0.031 | 0.0017 | <0.0001 |
| 0.5333 | 1 | 0.8441 | <0.0001 | 0.8858 | 0.8237 | 0.9163 | 0.0024 | 0.0147 | 0.0053 | <0.0001 |
| 0.7 | 1 | 0.9492 | <0.0001 | 0.9987 | 0.9977 | 0.9601 | 0.0008 | <0.0001 | 0.0001 | <0.0001 |
| 0.8667 | 1 | 0.8408 | <0.0001 | 1 | 0.9376 | 1 | 0.0006 | <0.0001 | 0.0003 | <0.0001 |
| 1.0333 | 1 | 0.9168 | 0.0001 | 0.9909 | 0.975 | 0.9602 | 0.0005 | 0.0021 | 0.0005 | 0.0058 |
| 1.2 | 1 | 0.9963 | 0.0002 | 0.9997 | 0.8432 | 1 | 0.0026 | 0.0019 | 0.0237 | 0.0009 |
| 1.3667 | 1 | 0.9987 | 0.0017 | 0.9988 | 0.9953 | 0.9802 | 0.0006 | 0.005 | 0.0913 | 0.0096 |
| 1.5333 | 1 | 0.9956 | 0.0092 | 0.9851 | 0.9917 | 0.9808 | 0.0012 | 0.0035 | 0.1964 | 0.0238 |
| 1.7 | 1 | 0.6019 | 0.2166 | 0.9865 | 0.9997 | 0.7631 | 0.0002 | 0.1276 | 0.5191 | 0.0386 |
| 2.3667 | 1 | 0.8655 | 0.1481 | 1 | 0.9982 | 0.9906 | 0.0013 | 0.0652 | 0.1775 | 0.0431 |
| 2.5333 | 1 | 0.8866 | 0.4933 | 0.9476 | 0.9999 | 0.861 | 0.0137 | 0.0335 | 0.056 | 0.0215 |
| 5.3667 | 1 | 0.9214 | 0.824 | 0.9882 | 0.9975 | 0.2548 | 0.0472 | 1 | 0.9824 | 0.0018 |
| 5.8667 | 1 | 0.9992 | 0.5113 | 0.7485 | 0.9735 | 0.9333 | 0.2862 | 0.9993 | 0.0547 | 0.068 |

### Speed of change in inter-retinal distance - Sliding window GLMM for all stimulus duration

| win_starts | win_span | p |
|---|---|---|
| 0.03333 | 1 | 1 |
| 0.2 | 1 | 1 |
| 0.36667 | 1 | 1 |
| 0.53333 | 1 | 1 |
| 0.7 | 1 | 1 |
| 0.86667 | 1 | 1 |
| 1.03333 | 1 | 1 |
| 1.2 | 1 | 1 |
| 1.36667 | 1 | 1 |
| 1.53333 | 1 | 1 |
| 1.7 | 1 | 1 |
| 1.86667 | 1 | 1 |
| 2.03333 | 1 | 1 |
| 2.2 | 1 | 1 |
| 2.36667 | 1 | 1 |
| 2.53333 | 1 | 1 |
| 2.7 | 1 | 1 |
| 2.86667 | 1 | 1 |
| 3.03333 | 1 | 1 |
| 3.2 | 1 | 1 |

| win_starts | win_span | p |
|---|---|---|
| 3.36667 | 1 | 1 |
| 3.53333 | 1 | 1 |
| 3.7 | 1 | 1 |
| 3.86667 | 1 | 1 |
| 4.03333 | 1 | 1 |
| 4.2 | 1 | 1 |
| 4.36667 | 1 | 0.41044 |
| 4.53333 | 1 | 0.44723 |
| 4.7 | 1 | 0.63788 |
| 4.86667 | 1 | 0.53885 |
| 5.03333 | 1 | 0.41197 |
| 5.2 | 1 | 1 |
| 5.36667 | 1 | 0.37867 |
| 5.53333 | 1 | 1 |
| 5.7 | 1 | 1 |
| 5.86667 | 1 | 1 |
| 6.03333 | 1 | 1 |
| 6.2 | 1 | 1 |
| 6.36667 | 1 | 1 |
| 6.53333 | 1 | 1 |
| 6.7 | 1 | 1 |
| 6.86667 | 1 | 1 |

| win_starts | win_span | p |
|---|---|---|
| 7.03333 | 1 | 1 |
| 7.2 | 1 | 1 |
| 7.36667 | 1 | 1 |
| 7.53333 | 1 | 1 |
| 7.7 | 1 | 1 |
| 7.86667 | 1 | 1 |
| 8.03333 | 1 | 1 |
| 8.2 | 1 | 1 |
| 8.36667 | 1 | 1 |
| 8.53333 | 1 | 1 |
| 8.7 | 1 | 1 |
| 8.86667 | 1 | 1 |
| 9.03333 | 1 | 1 |
| 9.2 | 1 | 1 |
| 9.36667 | 1 | 1 |
| 9.53333 | 1 | 1 |
| 9.7 | 1 | 1 |
| 9.86667 | 1 | 1 |
| 10.03333 | 1 | 1 |
| 10.2 | 1 | 1 |
| 10.36667 | 1 | 1 |
| 10.53333 | 1 | 1 |

| win_starts | win_span | p |
|---|---|---|
| 10.7 | 1 | 1 |
| 10.86667 | 1 | 1 |
| 11.03333 | 1 | 1 |
| 11.2 | 1 | 1 |
| 11.36667 | 1 | 1 |
| 11.53333 | 1 | 1 |
| 11.7 | 1 | 0.15753 |
| 11.86667 | 1 | <0.0001 |
| 12.03333 | 1 | <0.0001 |
| 12.2 | 1 | <0.0001 |
| 12.36667 | 1 | <0.0001 |
| 12.53333 | 1 | <0.0001 |
| 12.7 | 1 | <0.0001 |
| 12.86667 | 1 | <0.0001 |
| 13.03333 | 1 | <0.0001 |
| 13.2 | 1 | <0.0001 |
| 13.36667 | 1 | <0.0001 |
| 13.53333 | 1 | <0.0001 |
| 13.7 | 1 | <0.0001 |
| 13.86667 | 1 | <0.0001 |
| 14.03333 | 1 | <0.0001 |
| 14.2 | 1 | <0.0001 |

| win_starts | win_span | p |
|---|---|---|
| 14.36667 | 1 | <0.0001 |
| 14.53333 | 1 | <0.0001 |
| 14.7 | 1 | <0.0001 |
| 14.86667 | 1 | <0.0001 |
| 15.03333 | 1 | <0.0001 |
| 15.2 | 1 | <0.0001 |
| 15.36667 | 1 | <0.0001 |
| 15.53333 | 1 | <0.0001 |
| 15.7 | 1 | <0.0001 |
| 15.86667 | 1 | <0.0001 |
| 16.03333 | 1 | <0.0001 |
| 16.2 | 1 | <0.0001 |
| 16.36667 | 1 | <0.0001 |
| 16.53333 | 1 | <0.0001 |
| 16.7 | 1 | <0.0001 |
| 16.86667 | 1 | <0.0001 |
| 17.03333 | 1 | <0.0001 |
| 17.2 | 1 | 0.00036 |
| 17.36667 | 1 | 0.02096 |
| 17.53333 | 1 | 0.64478 |
| 17.7 | 1 | 0.685 |
| 17.86667 | 1 | 1 |

| win_starts | win_span | p |
|---|---|---|
| 18.03333 | 1 | 0.438 |
| 18.2 | 1 | 0.44751 |
| 18.36667 | 1 | 0.85659 |
| 18.53333 | 1 | 0.79627 |
| 18.7 | 1 | 0.67223 |
| 18.86667 | 1 | 1 |
| 19.03333 | 1 | 1 |
| 19.2 | 1 | 1 |
| 19.36667 | 1 | 1 |
| 19.53333 | 1 | 1 |
| 19.7 | 1 | 0.31885 |
| 19.86667 | 1 | 1 |
| 20.03333 | 1 | 1 |
| 20.2 | 1 | 1 |
| 20.36667 | 1 | 1 |
| 20.53333 | 1 | 0.22497 |
| 20.7 | 1 | 1 |
| 20.86667 | 1 | 1 |
| 21.03333 | 1 | 1 |
| 21.2 | 1 | 1 |
| 21.36667 | 1 | 1 |
| 21.53333 | 1 | 1 |

| win_starts | win_span | p |
|---|---|---|
| 21.7 | 1 | 1 |
| 21.86667 | 1 | 0.07944 |
| 22.03333 | 1 | 0.01989 |
| 22.2 | 1 | 0.3504 |
| 22.36667 | 1 | 0.8412 |
| 22.53333 | 1 | 0.05372 |
| 22.7 | 1 | 0.00082 |
| 22.86667 | 1 | 0.00252 |
| 23.03333 | 1 | 0.00121 |
| 23.2 | 1 | 0.00483 |
| 23.36667 | 1 | 0.01326 |
| 23.53333 | 1 | 0.01096 |
| 23.7 | 1 | 0.00081 |
| 23.86667 | 1 | 0.04318 |
| 24.03333 | 1 | 0.05121 |
| 24.2 | 1 | 1 |
| 24.36667 | 1 | 1 |
| 24.53333 | 1 | 1 |
| 24.7 | 1 | 1 |
| 24.86667 | 1 | 1 |
| 25.03333 | 1 | 1 |
| 25.2 | 1 | 1 |

| win_starts | win_span | p |
|---|---|---|
| 25.36667 | 1 | 1 |
| 25.53333 | 1 | 1 |
| 25.7 | 1 | 1 |
| 25.86667 | 1 | 1 |
| 26.03333 | 1 | 1 |
| 26.2 | 1 | 1 |
| 26.36667 | 1 | 1 |
| 26.53333 | 1 | 1 |
| 26.7 | 1 | 1 |
| 26.86667 | 1 | 1 |
| 27.03333 | 1 | 1 |
| 27.2 | 1 | 1 |
| 27.36667 | 1 | 1 |
| 27.53333 | 1 | 1 |
| 27.7 | 1 | 1 |
| 27.86667 | 1 | 1 |
| 28.03333 | 1 | 1 |
| 28.2 | 1 | 1 |
| 28.36667 | 1 | 1 |
| 28.53333 | 1 | 1 |
| 28.7 | 1 | 1 |
| 28.86667 | 1 | 1 |

| win_starts | win_span | p |
|---|---|---|
| 29.03333 | 1 | 1 |
| 29.2 | 1 | 1 |
| 29.36667 | 1 | 1 |
| 29.53333 | 1 | 1 |
| 29.7 | 1 | 1 |
| 29.86667 | 1 | 1 |
| 30.03333 | 1 | 1 |
| 30.2 | 1 | 1 |
| 30.36667 | 1 | 1 |
| 30.53333 | 1 | 1 |
| 30.7 | 1 | 1 |
| 30.86667 | 1 | 1 |
| 31.03333 | 1 | 1 |
| 31.2 | 1 | 1 |
| 31.36667 | 1 | 1 |
| 31.53333 | 1 | 1 |
| 31.7 | 1 | 1 |
| 31.86667 | 1 | 1 |
| 32.03333 | 1 | 1 |
| 32.2 | 1 | 1 |
| 32.36667 | 1 | 1 |
| 32.53333 | 1 | 1 |

| win_starts | win_span | p |
|---|---|---|
| 32.7 | 1 | 1 |
| 32.86667 | 1 | 1 |
| 33.03333 | 1 | 1 |
| 33.2 | 1 | 1 |
| 33.36667 | 1 | 1 |
| 33.53333 | 1 | 1 |
| 33.7 | 1 | 1 |
| 33.86667 | 1 | 1 |
| 34.03333 | 1 | 1 |
| 34.2 | 1 | 1 |
| 34.36667 | 1 | 1 |
| 34.53333 | 1 | 1 |
| 34.7 | 1 | 0.86922 |
| 34.86667 | 1 | 1 |
| 35.03333 | 1 | 1 |
| 35.2 | 1 | 1 |

### Speed of change in inter-retinal distance - Post-Hoc Comparisons for all stimulus duration

| win_starts | win_span | bio.ellipse | bio.rand | bio.scramb | bio.silhouette | ellipse.rand | ellipse.scramb | ellipse.silhouette |
|---|---|---|---|---|---|---|---|---|
| 11.8667 | 1 | <0.0001 | 0.1534 | 0.672 | 0.6835 | <0.0001 | <0.0001 | <0.0001 |
| 12.0333 | 1 | <0.0001 | 0.1209 | 0.5961 | 0.4439 | <0.0001 | <0.0001 | <0.0001 |
| 12.2 | 1 | <0.0001 | 0.1499 | 0.874 | 0.0876 | <0.0001 | <0.0001 | <0.0001 |
| 12.3667 | 1 | <0.0001 | 0.8266 | 0.921 | 0.4775 | <0.0001 | <0.0001 | <0.0001 |
| 12.5333 | 1 | <0.0001 | 0.9631 | 0.969 | 0.5405 | <0.0001 | <0.0001 | <0.0001 |
| 12.7 | 1 | <0.0001 | 1 | 0.9973 | 0.8163 | <0.0001 | <0.0001 | <0.0001 |
| 12.8667 | 1 | <0.0001 | 0.8594 | 0.9748 | 0.9995 | <0.0001 | <0.0001 | <0.0001 |
| 13.0333 | 1 | <0.0001 | 0.9959 | 1 | 0.9999 | <0.0001 | <0.0001 | <0.0001 |
| 13.2 | 1 | <0.0001 | 0.9814 | 0.9427 | 0.9998 | <0.0001 | <0.0001 | <0.0001 |
| 13.3667 | 1 | <0.0001 | 0.8361 | 0.933 | 0.9856 | <0.0001 | <0.0001 | <0.0001 |
| 13.5333 | 1 | <0.0001 | 0.8505 | 0.9378 | 0.8776 | <0.0001 | <0.0001 | <0.0001 |
| 13.7 | 1 | <0.0001 | 0.8652 | 0.8071 | 0.657 | <0.0001 | <0.0001 | <0.0001 |
| 13.8667 | 1 | <0.0001 | 0.6703 | 0.9894 | 0.5977 | <0.0001 | <0.0001 | <0.0001 |
| 14.0333 | 1 | <0.0001 | 0.5241 | 0.9979 | 0.3801 | <0.0001 | 0.0001 | <0.0001 |
| 14.2 | 1 | <0.0001 | 0.5138 | 0.9937 | 0.3693 | <0.0001 | <0.0001 | <0.0001 |
| 14.3667 | 1 | <0.0001 | 0.3616 | 0.9752 | 0.2647 | <0.0001 | <0.0001 | <0.0001 |
| 14.5333 | 1 | <0.0001 | 0.055 | 0.8686 | 0.0248 | <0.0001 | <0.0001 | <0.0001 |
| 14.7 | 1 | <0.0001 | 0.0011 | 0.9037 | 0.0058 | <0.0001 | <0.0001 | <0.0001 |
| 14.8667 | 1 | <0.0001 | 0.0024 | 0.5692 | 0.0023 | <0.0001 | <0.0001 | <0.0001 |
| 15.0333 | 1 | <0.0001 | 0.0001 | 0.2887 | 0.0001 | <0.0001 | <0.0001 | <0.0001 |

| win_starts | win_span | bio.ellipse | bio.rand | bio.scramb | bio.silhouette | ellipse.rand | ellipse.scramb | ellipse.silhouette |
|---|---|---|---|---|---|---|---|---|
| 15.2 | 1 | 0.0058 | 0.0007 | 0.205 | <0.0001 | <0.0001 | <0.0001 | <0.0001 |
| 15.3667 | 1 | 0.023 | 0.0001 | 0.0198 | <0.0001 | <0.0001 | <0.0001 | <0.0001 |
| 15.5333 | 1 | 0.003 | <0.0001 | 0.0057 | 0.0001 | <0.0001 | <0.0001 | <0.0001 |
| 15.7 | 1 | 0.0009 | <0.0001 | 0.0078 | 0.0194 | <0.0001 | <0.0001 | <0.0001 |
| 15.8667 | 1 | 0.0035 | 0.0004 | 0.001 | 0.0681 | <0.0001 | <0.0001 | <0.0001 |
| 16.0333 | 1 | 0.0509 | 0.0003 | 0.0023 | 0.0089 | <0.0001 | <0.0001 | <0.0001 |
| 16.2 | 1 | 0.3589 | 0.0003 | 0.0046 | 0.0042 | <0.0001 | <0.0001 | <0.0001 |
| 16.3667 | 1 | 0.2588 | 0.0049 | 0.0962 | 0.0141 | <0.0001 | <0.0001 | <0.0001 |
| 16.5333 | 1 | 0.3549 | 0.0117 | 0.2447 | 0.0017 | <0.0001 | 0.0003 | <0.0001 |
| 16.7 | 1 | 0.1083 | 0.1325 | 0.6783 | 0.0023 | <0.0001 | 0.0023 | <0.0001 |
| 16.8667 | 1 | 0.0999 | 0.1823 | 0.6064 | 0.0054 | <0.0001 | 0.0015 | <0.0001 |
| 17.0333 | 1 | 0.0809 | 0.2131 | 0.6418 | 0.0079 | <0.0001 | 0.0008 | <0.0001 |
| 17.2 | 1 | 0.0915 | 0.1598 | 0.58 | 0.0193 | <0.0001 | 0.0011 | <0.0001 |
| 22.7 | 1 | 0.0186 | 0.3036 | 0.038 | 0.9873 | 0.0014 | <0.0001 | 0.0679 |
| 22.8667 | 1 | 0.0165 | 0.5301 | 0.0434 | 0.9298 | 0.0033 | <0.0001 | 0.0139 |
| 23.0333 | 1 | 0.3153 | 0.394 | 0.0255 | 0.7087 | 0.02 | <0.0001 | 0.0469 |
| 23.2 | 1 | 0.9989 | 0.0819 | 0.0084 | 0.0282 | 0.1435 | 0.0049 | 0.0131 |
| 23.7 | 1 | 0.967 | 0.002 | 0.0105 | 0.0003 | 0.0503 | 0.031 | 0.0074 |
