## Supplementary figures and images for "Biological point-light displays scanning by the principal eyes of a jumping spider"

### unnamed-chunk-3-1.png

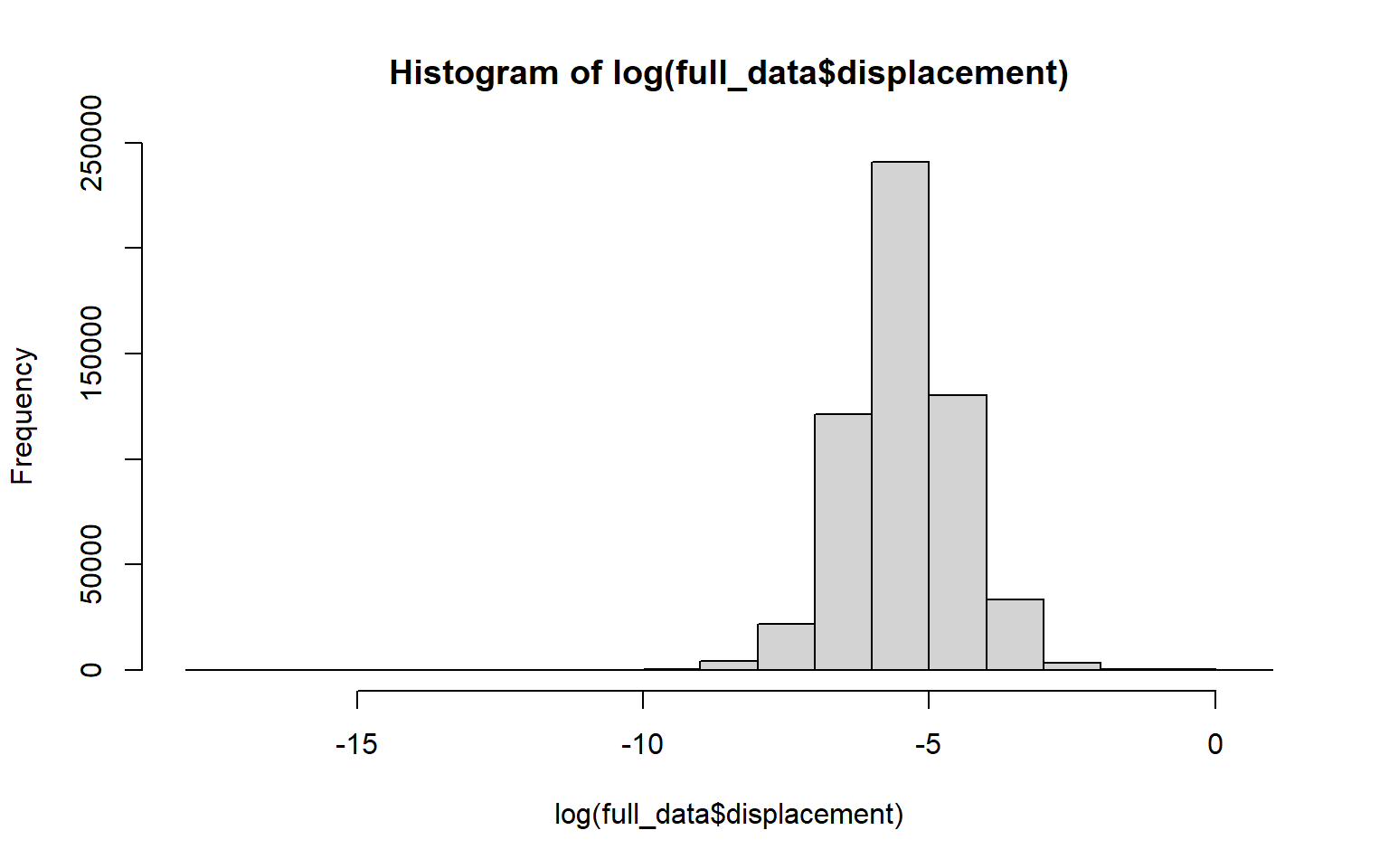

### unnamed-chunk-3-2.png

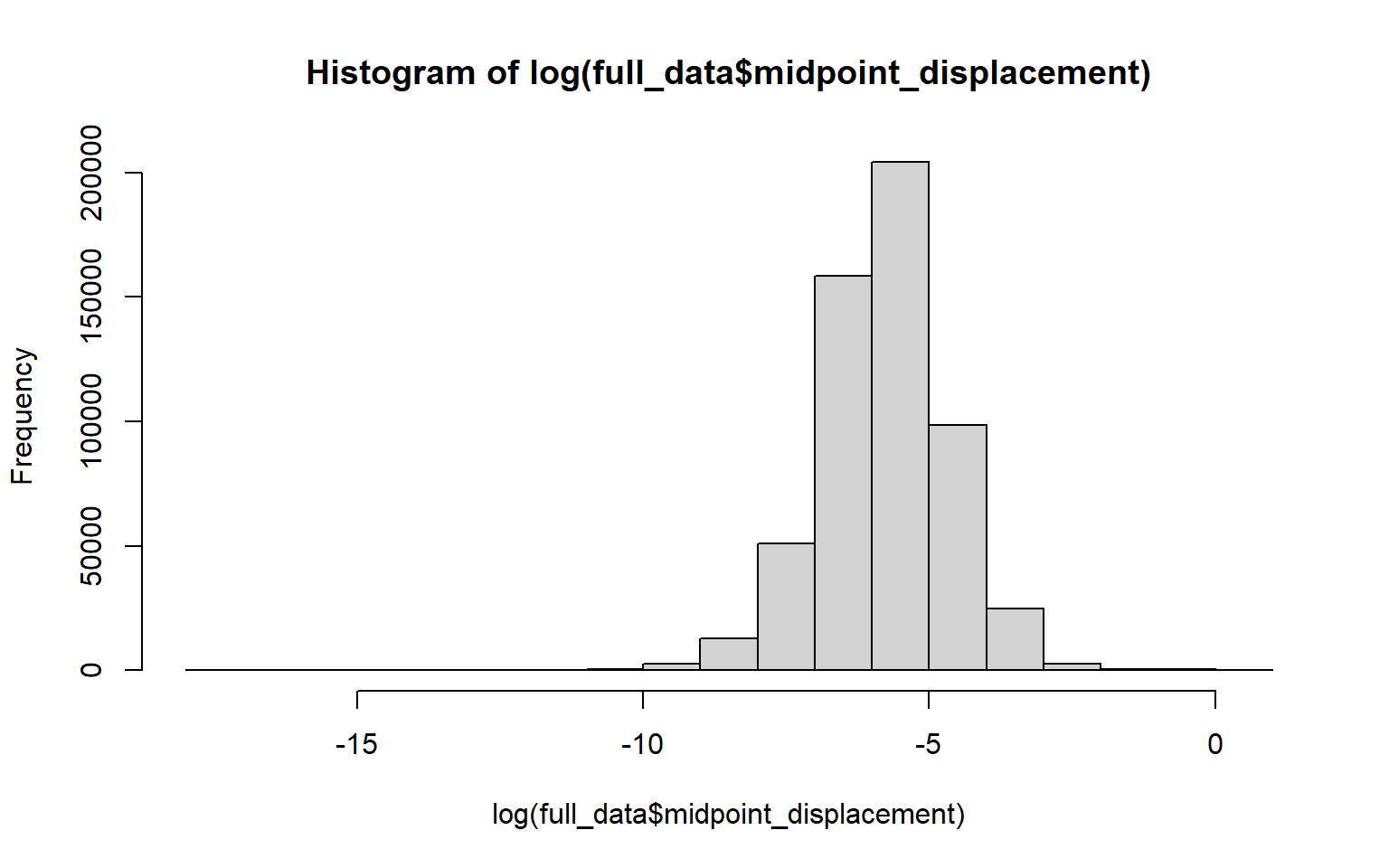

### unnamed-chunk-3-3.png

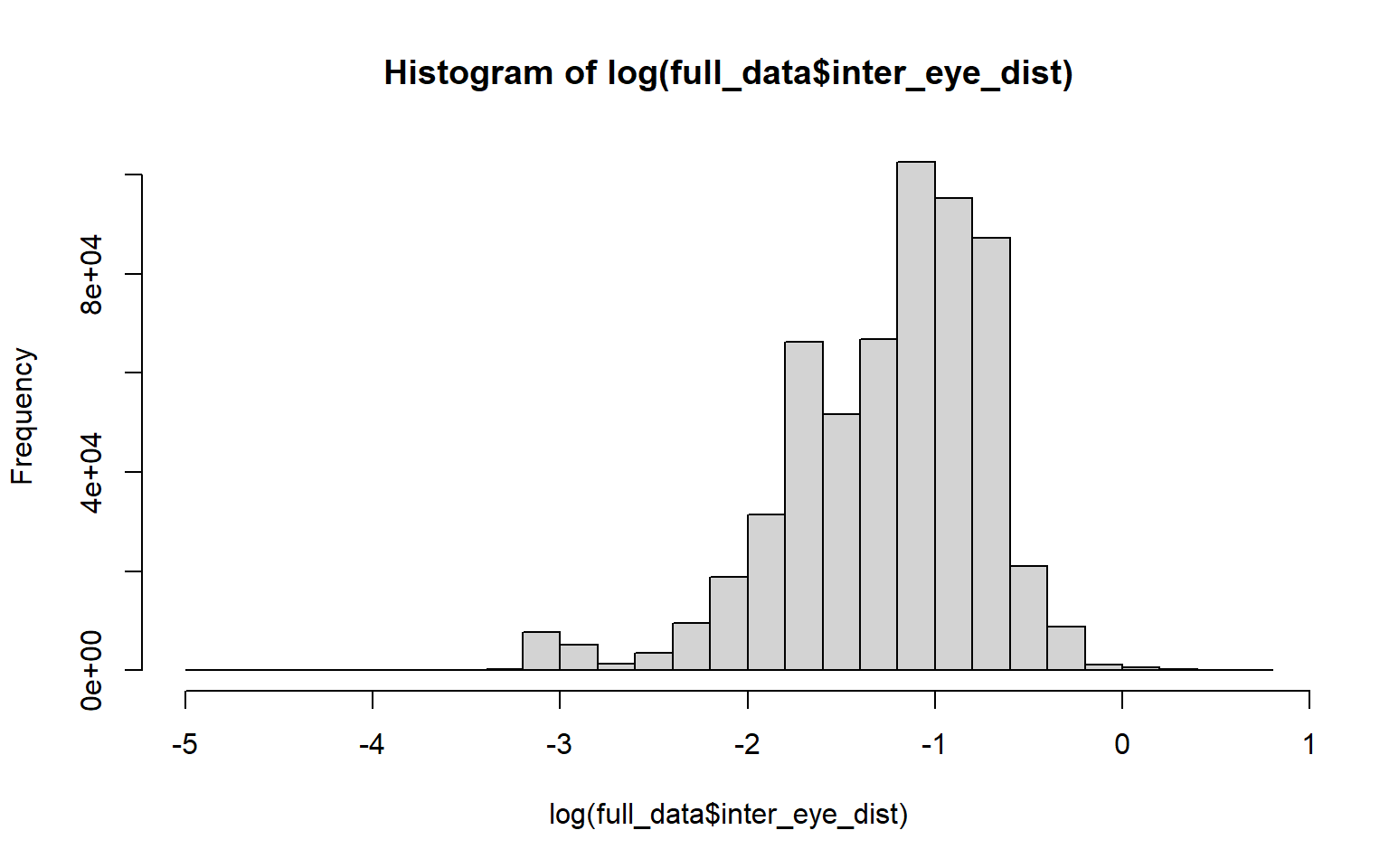

### unnamed-chunk-3-4.png

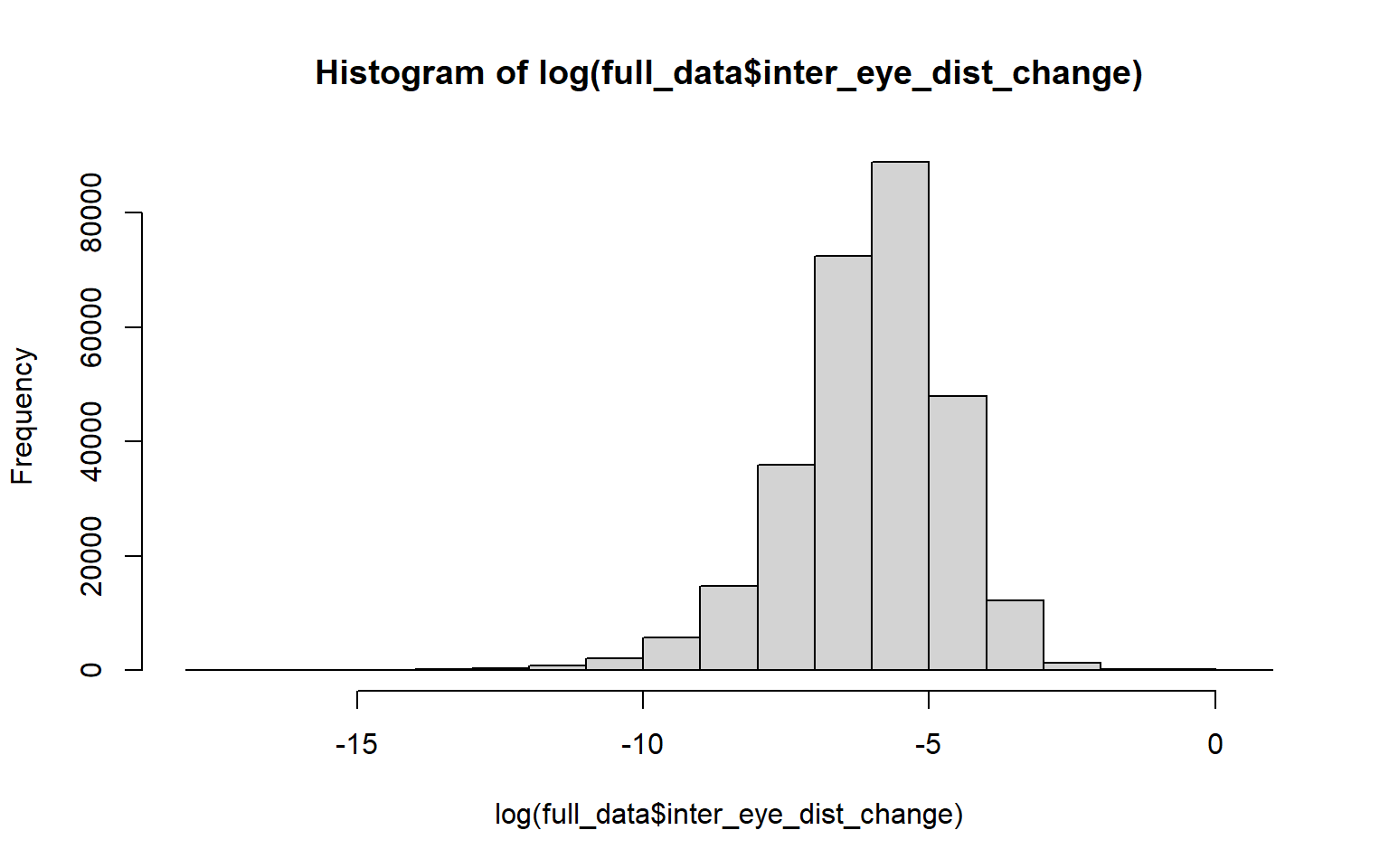

### unnamed-chunk-3-5.png

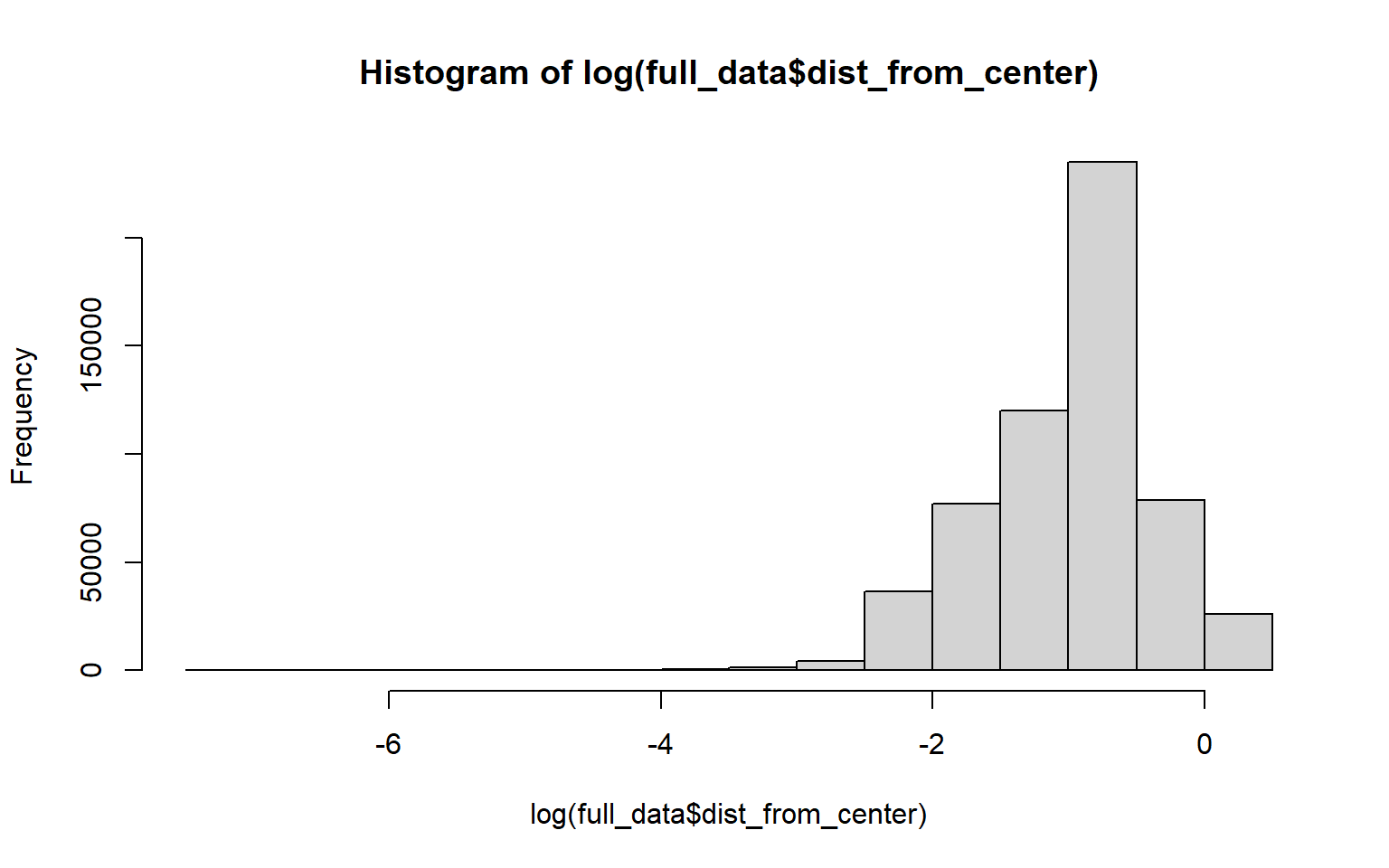

### unnamed-chunk-55-1.png

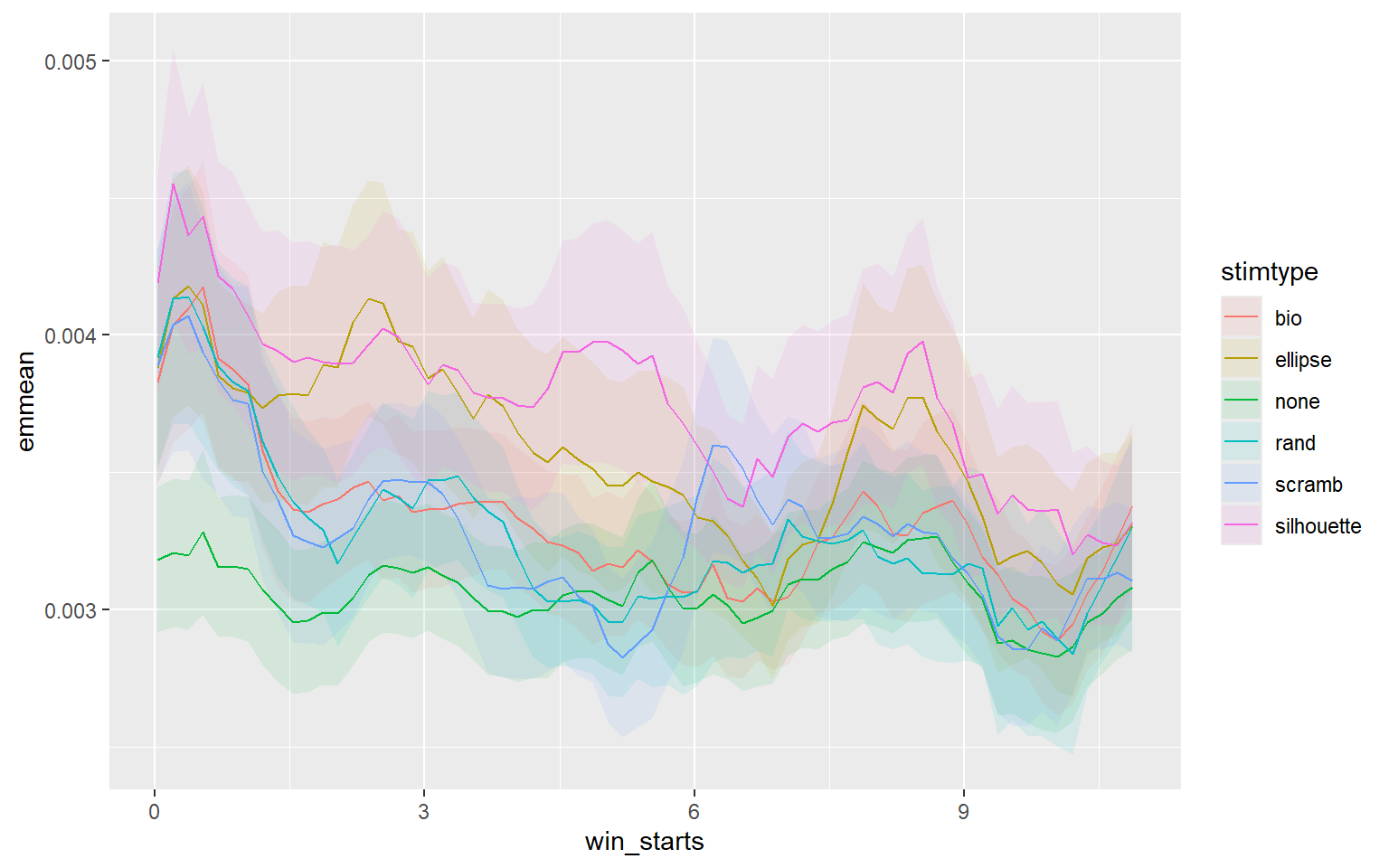

### unnamed-chunk-56-1.png

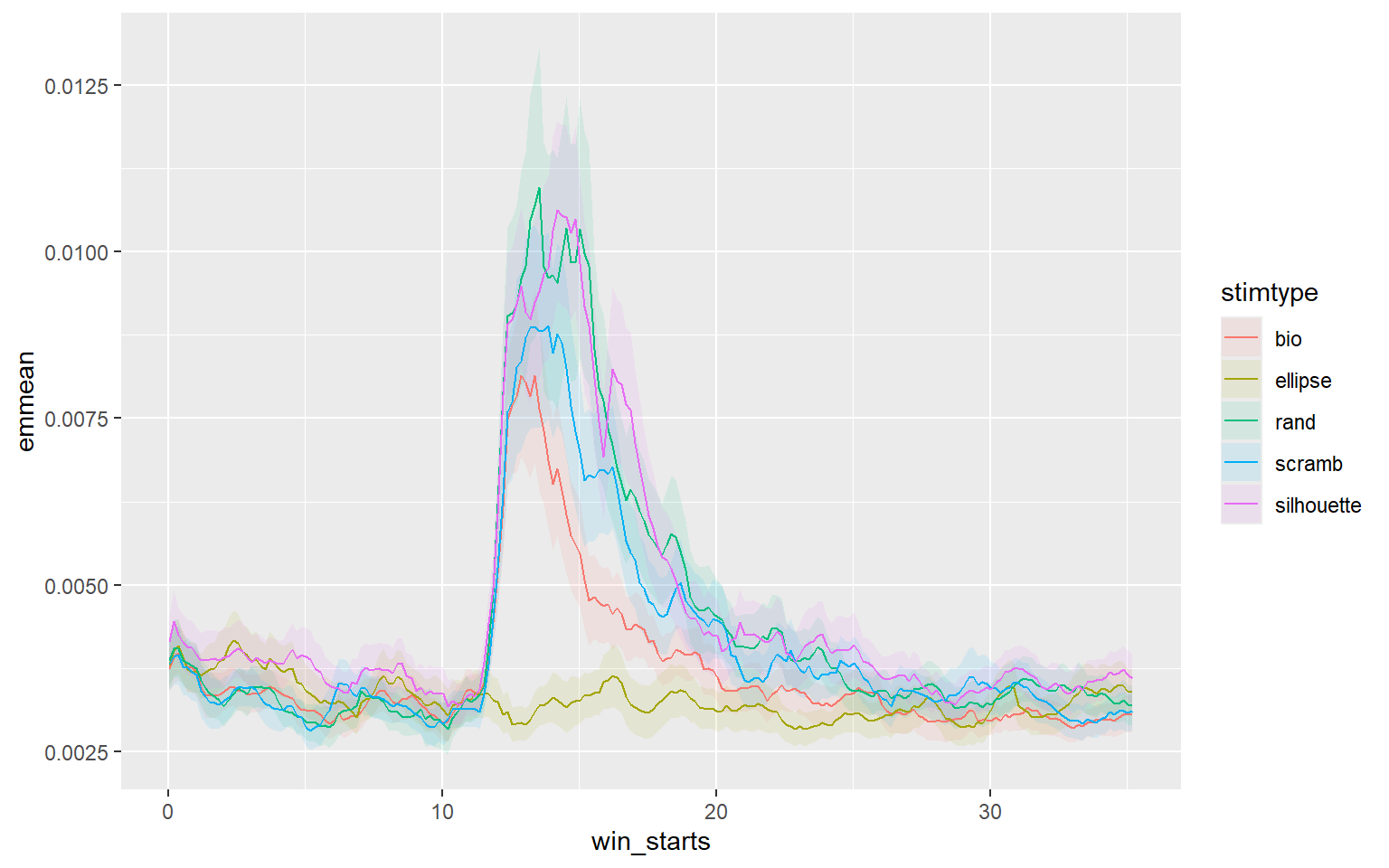

### unnamed-chunk-57-1.png

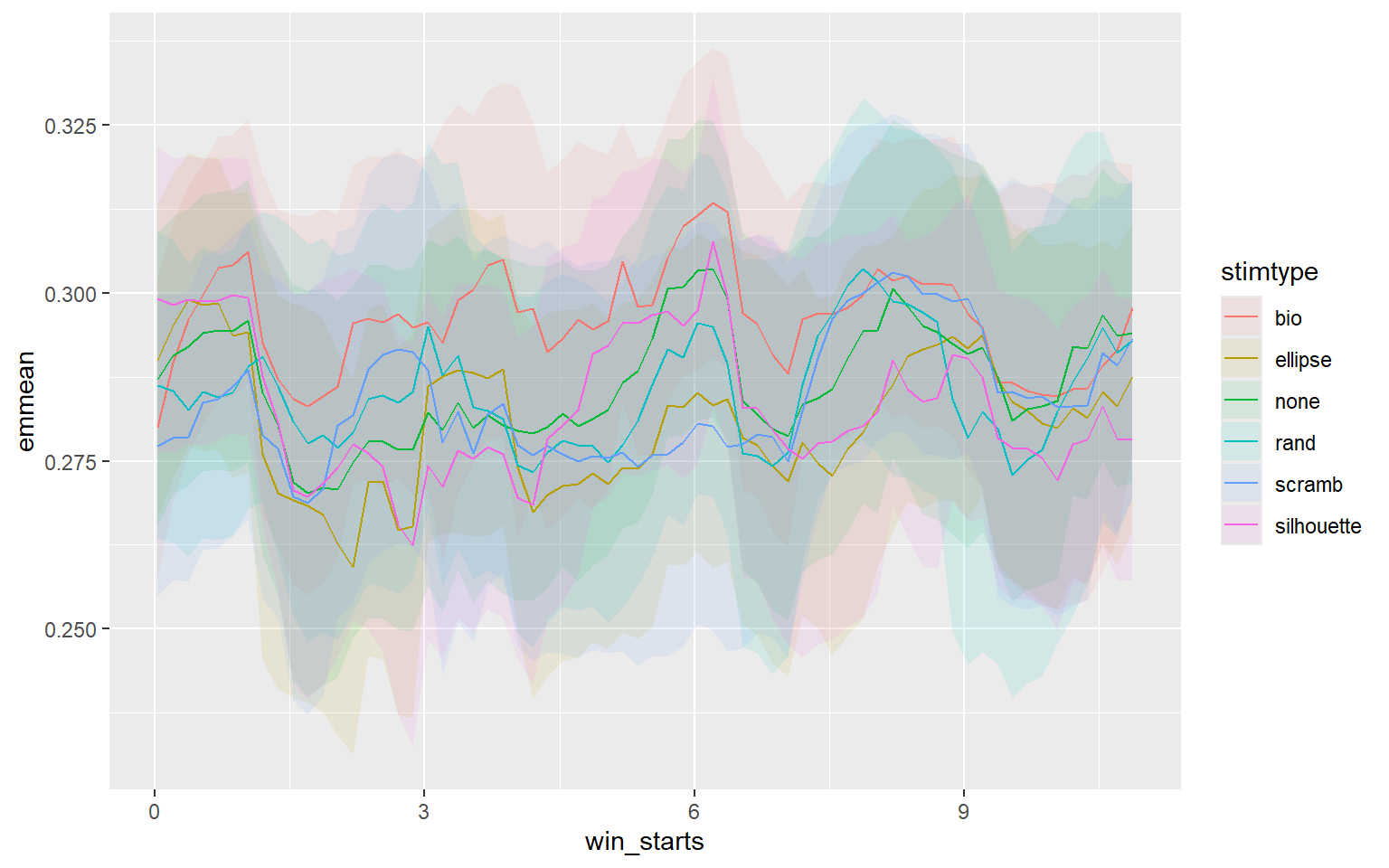

### unnamed-chunk-58-1.png

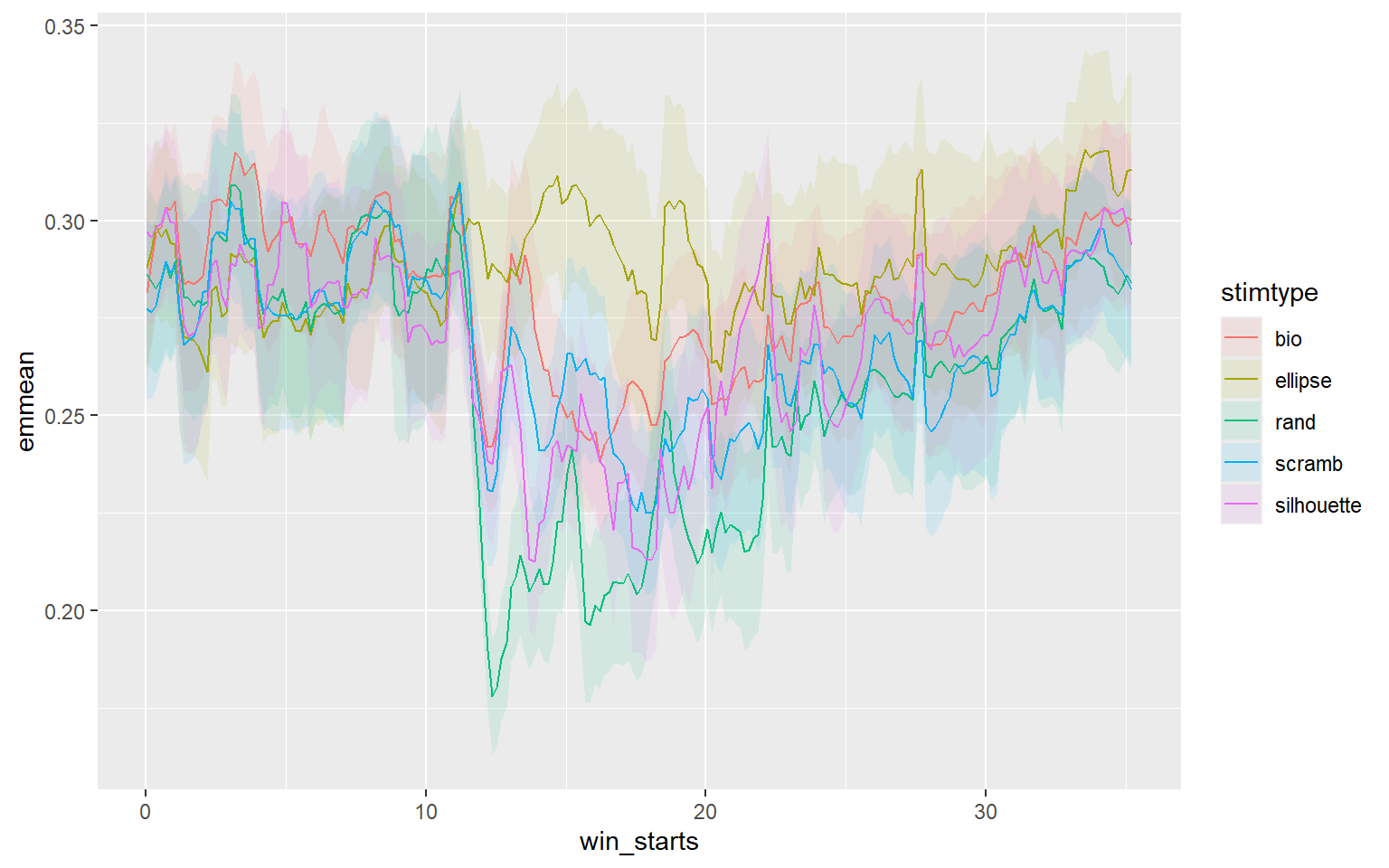

### unnamed-chunk-59-1.png

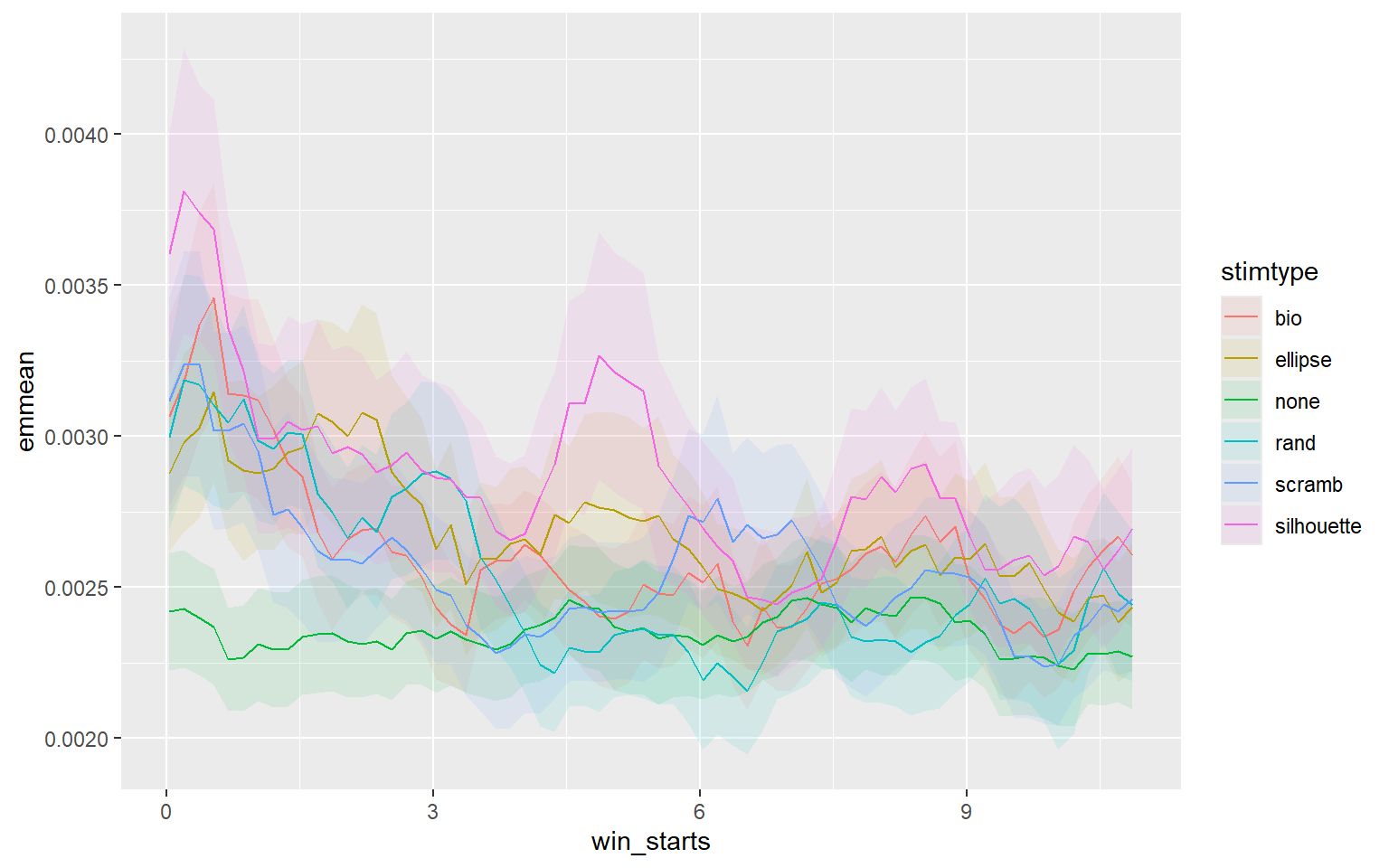

### unnamed-chunk-60-1.png

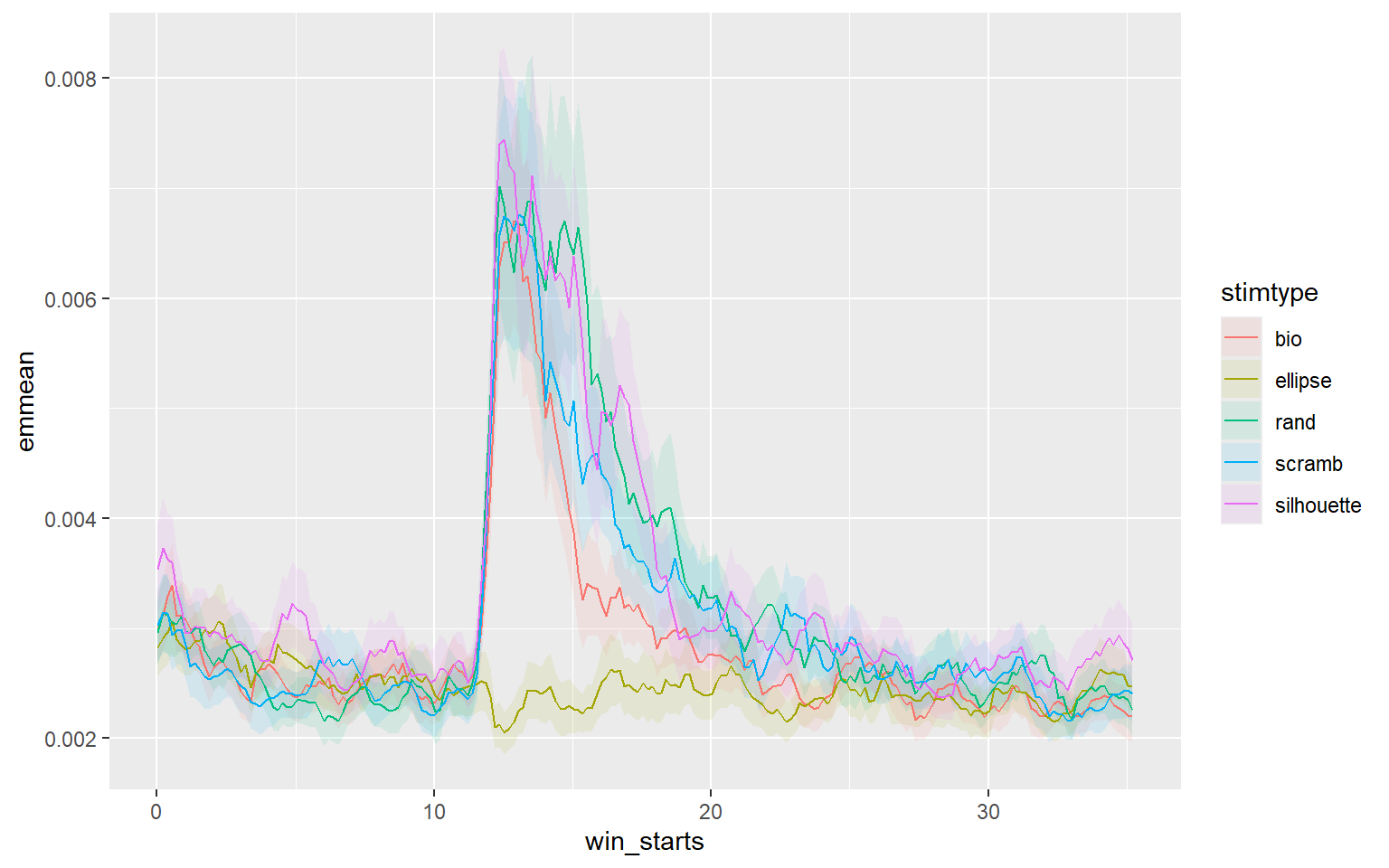

### unnamed-chunk-61-1.png

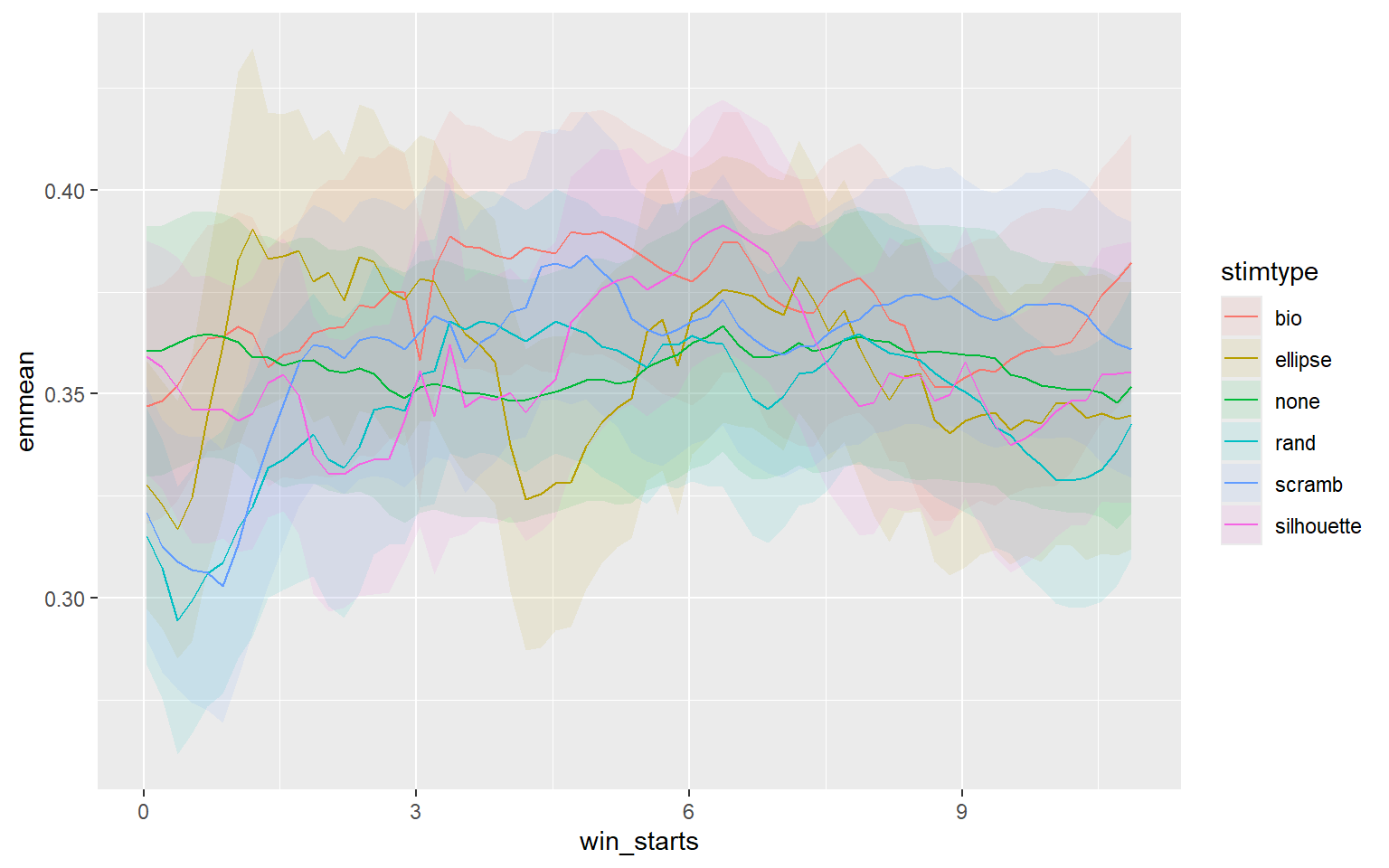

### unnamed-chunk-62-1.png

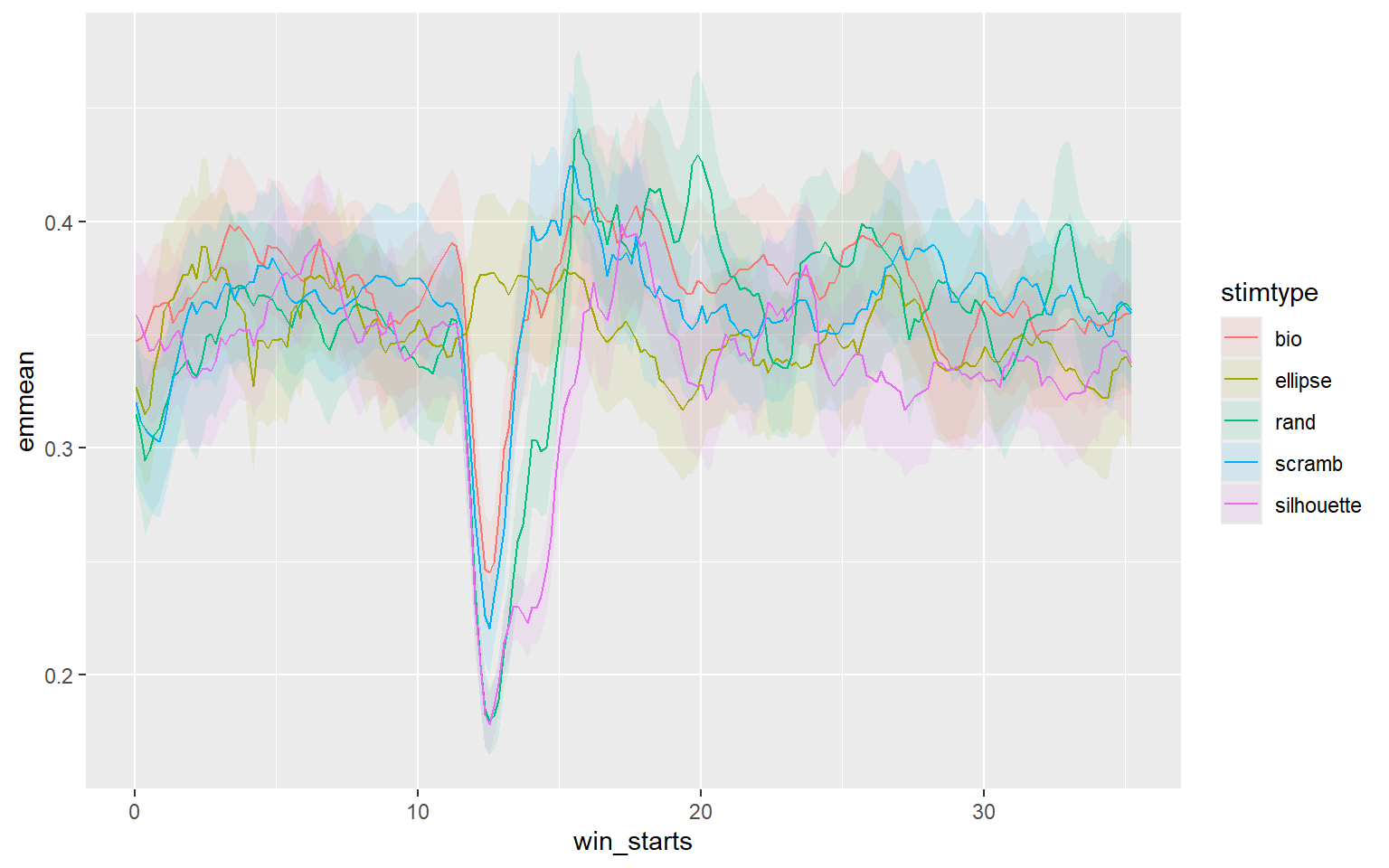
